# SNAP-tag2: faster and brighter protein labeling

**DOI:** 10.1101/2024.08.28.610127

**Authors:** Stefanie Kühn, Veselin Nasufovic, Jonas Wilhelm, Julian Kompa, Eline M.F. de Lange, Yin-Hsi Lin, Cornelia Egoldt, Jonas Fischer, Artem Lennoi, Miroslaw Tarnawski, Jochen Reinstein, Rifka Vlijm, Julien Hiblot, Kai Johnsson

## Abstract

SNAP-tag is a powerful tool for labeling proteins with synthetic fluorophores in bioimaging. However, its utility in live-cell applications can be constrained by its relatively slow labeling kinetics and the limited cell permeability of its substrates. Here we introduce new labeling substrates and an engineered SNAP-tag for faster labeling *in vitro* and in live cells. SNAP-tag2 presents a second-order rate constant with rhodamine substrates that approaches 10^7^ s^-1^ M^-1^, a 100-fold improvement over the corresponding SNAP-tag-substrate pairs. When labeled with highly fluorogenic dyes, SNAP-tag2 also shows a 5-fold increase in fluorescence brightness relative to currently used SNAP-tag. The increased labeling kinetics and brightness of SNAP-tag2 translates into a greatly improved performance in various live-cell (super-resolution) imaging applications.

## Introduction

Self-labeling protein tags (SLPs) can be specifically and covalently labeled with synthetic probes *in vitro* and in live cells^[1]^. Popular examples of such tags are SNAP-tag^[2-3]^, HaloTag7^[4]^ and, to a lesser extent, CLIP-tag^[5]^. Their main field of use has been in bioimaging, where these tags offer the opportunity to attach bright and photostable synthetic fluorophores to proteins of interest. The development of fluorogenic probes for protein labeling, i.e. probes that only become highly fluorescent upon binding to the protein of interest, has further facilitated the use of SLPs in live-cell bioimaging as it decreases background signal from unbound dye^[6-7]^.

HaloTag7 is at present the most popular SLP for live-cell imaging. HaloTag7 was engineered to react fast and efficiently with chloroalkane (CA)-rhodamine substrates^[8]^ (Supplementary Figure 1). Rhodamines are well-suited synthetic fluorophores for live-cell imaging as they cover a broad spectral range, possess excellent spectroscopic properties^[9]^, can be fluorogenic^[6, 10]^ and are live-cell permeable^[11-12]^. The fluorogenicity and permeability of rhodamines can probably be attributed to the reversible spirocyclization of the zwitterionic, fluorescent rhodamine to a nonpolar and non-fluorescent spirolactone^[7, 12-13]^. For selected CA-rhodamines, labeling kinetics of HaloTag7 are close to the diffusion limit. The high reactivity of HaloTag7 towards CA-rhodamines is at least partially due to specific interactions of the protein surface with the rhodamine dye.^[14]^ The combination of the high labeling velocity of HaloTag7 with CA-rhodamines, their relatively good permeability as well as the excellent performance of these dyes in various microscopy applications have made the approach a powerful tool in bioimaging^[15]^.

SNAP-tag undergoes a nucleophilic substitution reaction with *O*^*6*^-benzylguanine (BG) or chloropyrimidine (CP) derivatives, in which the benzyl group of the substrate is irreversibly transferred to an active site reactive cysteine residue (Fig. 1a). SNAP-tag is a highly engineered human *O*^*6*^-alkylguanine-DNA alkyltransferase (hAGT) that was optimized for reaction with BG-derivatives^[2, 16-18]^. While BG-derivatives show faster labeling kinetics with SNAP-tag than the corresponding CP-derivatives^[14]^, the less polar CP-derivatives generally perform better in live-cell imaging^[19]^. However, compared to HaloTag7, SNAP-tag labeling kinetics with rhodamine-based substrates are in general at least two orders of magnitude slower^[14]^. Additionally, HaloTag7 shows higher fluorogenicity with rhodamines than SNAP-tag^[7, 20]^. This is presumably due to specific interactions between the fluorophore and the HaloTag7 surface, which promote an equilibrium shift from the non-fluorescent spirolactone to the fluorescent zwitterionic state^[21]^. No such specific interactions between SNAP-tag and rhodamine-based fluorophores have been identified^[14]^.

**Fig. 1.**
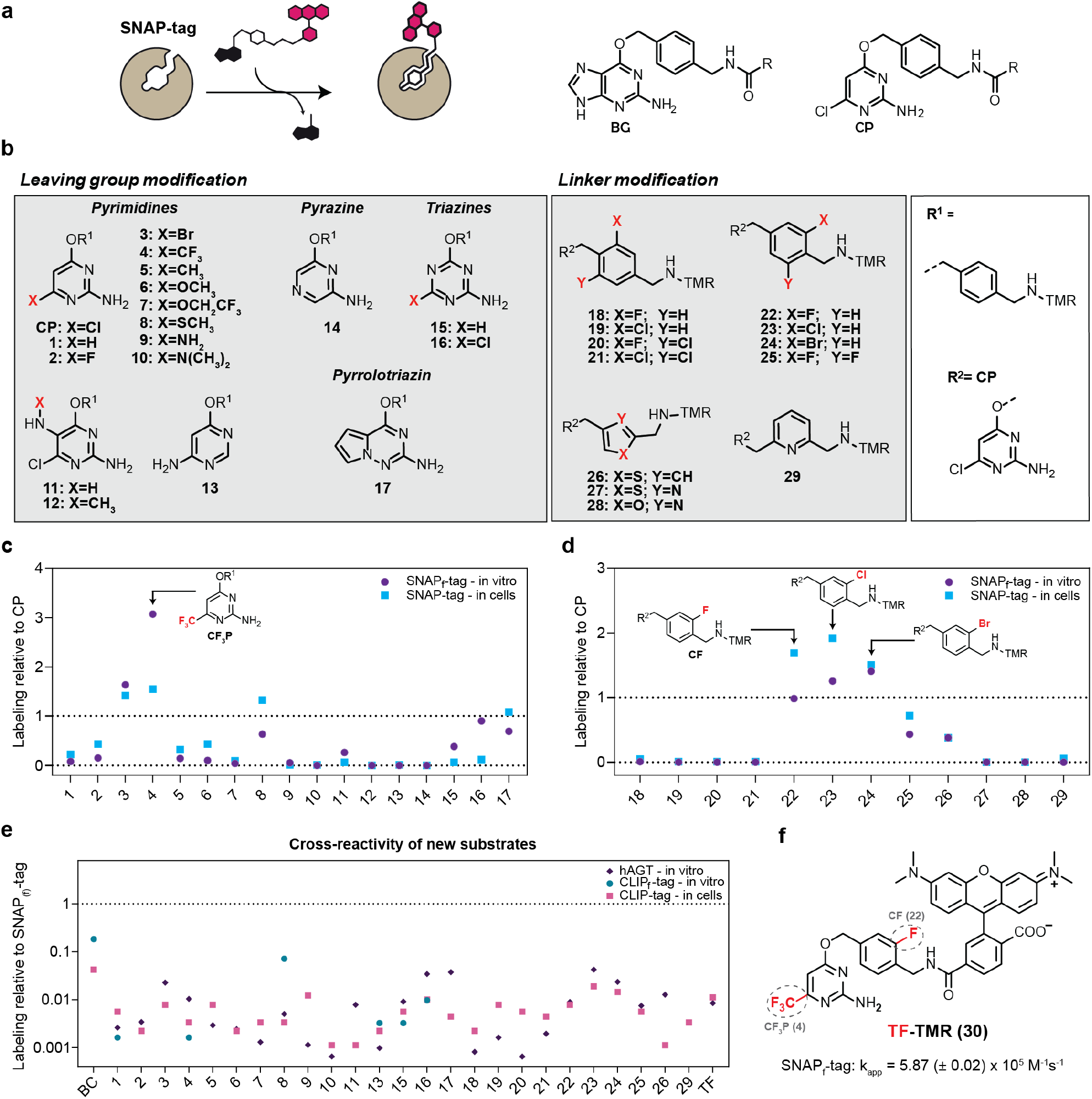
SNAP-tag substrate screening for more efficient labeling *in vitro* and in live mammalian cells. **a**, Scheme of SNAP-tag labeling reaction with fluorophore substrates. The chemical structures of SNAP-tag substrates BG and CP are shown on the right. R represents the functional moiety to be linked to SNAP-tag. **b**, Chemical structures of modified SNAP-tag substrates divided into two groups: leaving group modification and linker modification. Modified leaving groups were linked to tetramethylrhodamine (TMR) over linker R^1^. Modified linkers coupled to TMR were tested on the CP (R^2^) scaffold. **c**, Comparison of substrates **1**-**17** and **d**, substrates **18**-**29** relative to SNAP_(f)_-tag labeling with CP-TMR regarding their *in vitro* labeling kinetics and performance in live-cell labeling. *In vitro* labeling kinetics of SNAP_f_-tag with new substrates were measured recording fluorescence polarization traces over time. Apparent second-rate constants (k_app_) were calculated (Supplementary Table 1) and normalized to the k_app_ of SNAP_f_-tag with CP-TMR. Live-cell performance of new substrates was tested by labeling of U2OS cells that stably express a mEGFP-SNAP-tag fusion protein with TMR-substrates at [100 nM] for 2 h. Cells were washed and analyzed *via* flow cytometry. Fluorescence intensity ratios of TMR/mEGFP were calculated (Supplementary Table 1) and normalized to the ratio obtained for SNAP-tag with CP-TMR. Leaving group substrate **4** and linker substrates **22, 23** and **24** showed the most promising results relative to CP-TMR. **e**, Reactivity of selected SNAP-tag substrates with hAGT and CLIP_(f)_-tag compared to SNAP_(f)_-tag labeling with CP-TMR. Experiments were conducted as previously described. **f**, Chemical structure of TF-TMR (**30**) found through the combination of leaving group **4** and linker **22**. The apparent second-order rate constant (k_app_) of SNAP_f_-tag labeling with TF-TMR is depicted below.

The limited cell permeability of SNAP-tag’s substrate as well as SNAP-tag’s reduced labeling kinetics and fluorogenicity can impede its use in certain live-cell imaging applications. Here, we present SNAP-tag2, a highly engineered SNAP-tag mutant that rapidly reacts with newly developed pyrimidine-based substrates optimized for live-cell applications, and demonstrate the effectiveness of SNAP-tag2 and its substrates in a range of bioimaging applications.

## Results

To improve the performance of SNAP-tag in live-cell imaging, we decided to first develop new substrate scaffolds for higher intrinsic reactivity *in vitro* and improved live-cell compatibility, followed by engineering of SNAP-tag for faster reaction kinetics and enhanced fluorescence brightness with these new substrates. SNAP-tag substrates can be divided into two parts: the leaving group and the linker that connects the synthetic probe to the reactive cysteine (Fig. 1b). We synthesized 17 different heterocycles connected through a benzyl linker to tetramethylrhodamine (TMR) and measured their labeling kinetics *in vitro* with SNAP_f_-tag, a single point mutant of SNAPtag with increased labeling kinetics^[14, 22]^, as well as their performance of labeling SNAP-tag in a cellular assay (Fig. 1c-d, Supplementary Table 1). The selection of heterocycles was based on previous observations that pyrimidines can be good substrates for SNAP-tag^[19, 23]^, but also comprised pyrazines, triazines and a pyrrolotriazine. Of these compounds, the trifluoromethyl-substituted pyrimidine **4** (CF_3_P) showed the best results (Fig. 1c), with a ∼3-fold increase in labeling kinetics *in vitro* and a ∼1.5-fold higher fluorescent labeling in live cells. Additionally, we investigated 12 different linkers connecting CP as leaving group and TMR as fluorescent probe. The fluoro- and chloro-derivatives **22** and **23** showed an increased fluorescent labeling of ∼2-fold in the cellular assay (Fig. 1d). However, the chloro-analogue **23** showed an increased reactivity against parental hAGT (Fig. 1e). To reduce the risk of unwanted background reactions, it was therefore excluded. Combining the features of **4** (CF_3_P) and **22** (CF) yielded trifluoromethyl fluorobenzyl pyrimidine **30** (TF), which as a TMR-derivative showed an apparent second-order rate constant (k_app_) of 5.87 (± 0.02) x 10^5^ M^-1^s^-1^ for the labeling of SNAP_f_-tag (Fig. 1f), which corresponds to a 3.9-fold faster labeling kinetic than its reaction with CP-TMR (k_app_ = 1.51 (± 0.01) x 10^5^ M^-1^s^-1^)^[14]^. Moreover, TF-TMR showed a ∼1.7-fold higher fluorescent labeling than CP-TMR in live cells (Supplementary Fig. 2).

In addition to substrate optimization, we used protein engineering to increase the reactivity of SNAP-tag towards pyrimidine-based substrates and to increase its brightness when labeled with fluorogenic rhodamines such as MaP618^[7]^. Specifically, we used mutagenesis of selected residues, computational design and directed evolution to identify mutants with increased reactivity and fluorogenicity (Fig. 2 and Supplementary Fig. 3). The choice of positions for mutagenesis was guided by the available X-ray structures of SNAP-tag in its apo (PDB: 3KZY), benzylated (PDB: 3L00), BG-bound (PDB: 3KZZ) and TMR-labeled (PDB: 6Y8P) states as well as by past engineering efforts^[2, 17, 22, 24]^ (Fig. 2a). Furthermore, we used the sequence- and structure-based design method PROSS^[25]^ to identify mutations that might increase the thermal stability of SNAP-tag (Fig. 2a). For directed evolution experiments, protein libraries were screened using yeast surface display combined with fluorescence-activated cell sorting (Supplementary Fig. 3a). Libraries of two distinct regions were created by saturation mutagenesis (Fig. 2a): the β-strand proximal to the active site (residues 29-36) and the active-site loop (residues 155-161). In addition, a synthetic deep mutational scanning library (sDMSL)^[26]^ was screened, in which each residue in SNAP-tag was individually replaced by all other 19 amino acids, excluding cysteines but including deletions. Moreover, we used RosettaRemodel^[27]^ to design a short eight-residue loop to replace a long, unstructured region in SNAP-tag (residues 37-54; Fig. 2a-c). Protein variants were screened for enhanced labeling kinetics and/or increased fluorescence intensity using either TMR- or MaP618-derivatives (Supplementary Fig. 1 and 3). These combined engineering efforts led to the creation of SNAP-tag2, which carries 11 substitutions and a replacement of 18 amino acids (residues 37-54) by a peptide of eight amino acids (Fig. 2b-c), without destabilizing the protein (T_m_ ≈ 65 °C; Supplementary Fig. 4). The k_app_ of its reaction with TF-TMR is of 8.22 (± 0.79) × 10^6^ M^-1^s^-1^ (Table 1, Supplementary Table 2 and 3), which is about 100-fold faster than the reaction of SNAP-tag with CP-TMR (Fig. 2d) and close to the rate constant for the reaction of HaloTag7 with CA-TMR (k_app_ = 1.88 (± 0.01) × 10^7^ M^-1^s^-1^)^[14]^. SNAP-tag2 also shows increased reaction kinetics with CP- and CF-based substrates and reaches its fastest labeling reaction with TF-CPY *in vitro* with a k_app_ of 1.04 (± 0.12) x 10^7^ M^-1^s^-1^ (Table 1, Supplementary Table 2 and 3). Furthermore, SNAP-tag2 shows increased reactivity towards non-fluorescent probes (Fig. 2d, Supplementary Table 4). The extinction coefficients and quantum yields of SNAP-tag2 labeled with different rhodamines are similar to those of its predecessor, except for very fluorogenic MaP618-substrates (Supplementary Table 5). For these, SNAP-tag2 shows a significantly higher absorbance than SNAP_f_-tag, (Fig. 2e), indicating that SNAP-tag2 might have a higher capacity to shift the dye towards its zwitterionic, fluorescent state.

**Table 1:** Second-order rate constants (k_app_) of SNAP-tag2 labeling with different TMR and CPY substrates. Values represent the average k_app_ values calculated from experimental replicates (Supplementary Table 2).

| TMR substrate | $k_{app}$ (value ± s.d.) [$M^{-1}s^{-1}$] | CPY substrate | $k_{app}$ (value ± s.d.) [$M^{-1}s^{-1}$] |
| --- | --- | --- | --- |
| TF-TMR | $8.22 (\pm 0.79) \times 10^6$ | TF-CPY | $1.04 (\pm 0.12) \times 10^7$ |
| CF-TMR | $6.21 (\pm 3.65) \times 10^6$ | CF-CPY | $3.62 (\pm 0.73) \times 10^6$ |
| CP-TMR | $7.82 (\pm 2.78) \times 10^6$ | CP-CPY | $5.29 (\pm 1.62) \times 10^6$ |

**Fig. 2.**
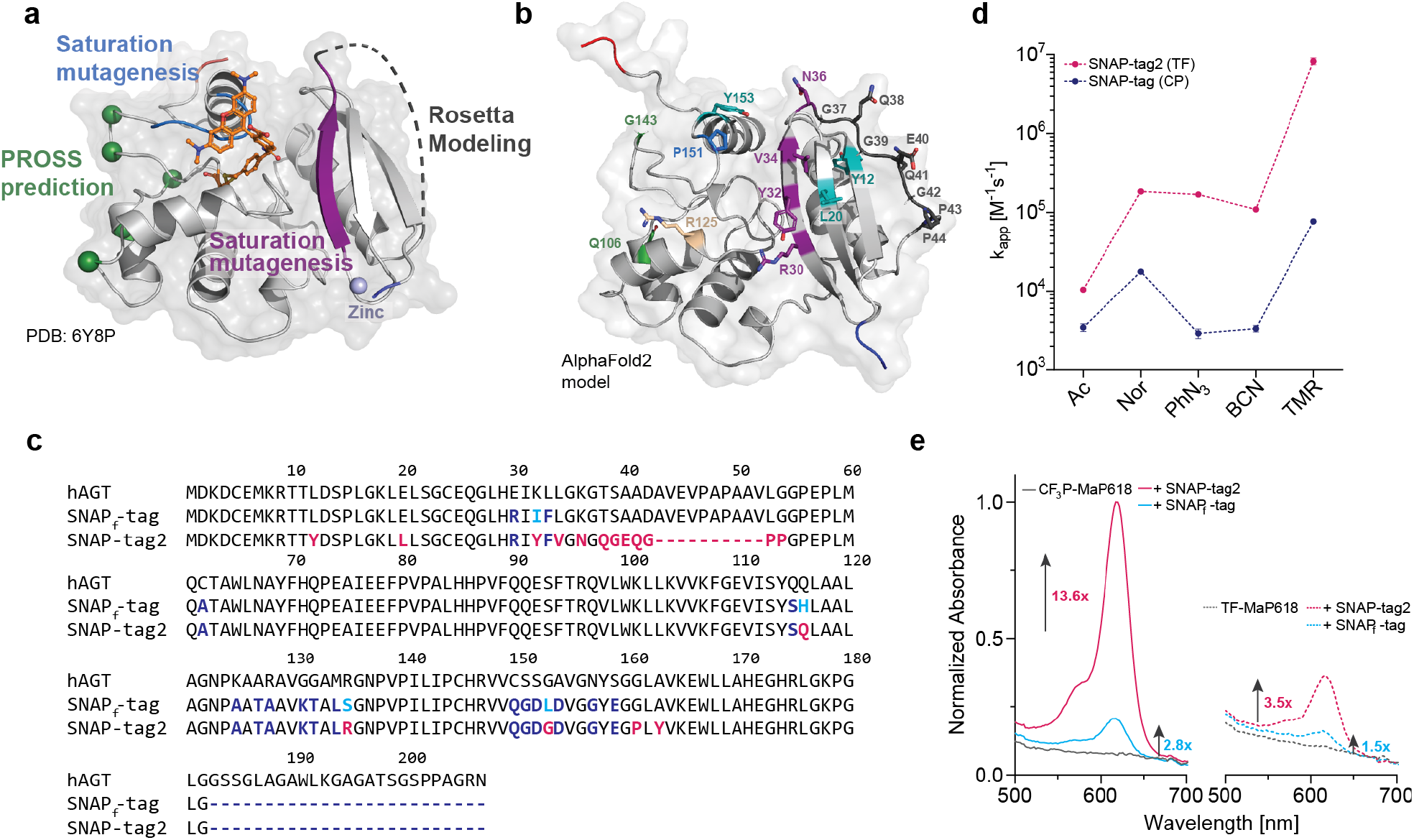
SNAP-tag engineering aiming for faster and brighter fluorescence labeling. **a**, Crystal structure of SNAP-tag labeled with TMR (PDB: 6Y8P) with engineered regions highlighted. SNAP-tag is represented as light-gray cartoon and the TMR-ligand is represented as sticks. Saturation mutagenesis libraries were constructed on the C-terminal loop region (residues 155-161) and the β-strand proximal to the active site (residues 29-36), highlighted in blue and violet, respectively. Substitutions predicted by PROSS^[25]^ to increase the protein thermal stability are highlighted as green spheres. The unstructured region in SNAP-tag (residues 37-55) highlighted as gray dotted lines was redesigned using RosettaRemodel^[27]^. Termini are highlighted in blue (N-terminus) and red (C-terminus) and the coordinated zinc ion is illustrated as light-blue sphere. **b**, AlphaFold2^[35]^ model of SNAP-tag2. Introduced substitutions are highlighted as sticks and color-coded based on the engineering rational. Green: PROSS^[25]^ prediction. Marine blue and deep-purple: saturation mutagenesis libraries. Wheat: rational design. Teal: deep mutational scanning library. Dark gray: Rosetta modeled loop. Termini are highlighted in blue (N-terminus) and red (C-terminus). **c**, Sequence alignment of hAGT, SNAP_f_-tag and SNAP-tag2. Dark blue: common differences of SNAP_f_-tag and SNAP-tag2 vs. hAGT Turquoise: unique differences SNAP_f_-tag vs. hAGT. Pink: unique differences SNAP-tag2 vs hAGT. Amino acid deletions are presented as dotted lines. **d**, Comparison of labeling kinetics between SNAP-tag2 with TF- and SNAP-tag with CP-substrates. Apparent second order rate constants (k_app_) of SNAP-tag2 with TF-substrates demonstrate one to two orders of magnitude faster labeling kinetics compared to the reaction of parental SNAP-tag with CP-substrates (CP results taken from Wilhelm et al.)^[14]^. Abbreviations: Ac, acetate; Nor, (1S,4S)-5-methylbicyclo[2.2.1]hept-2-ene (norbornene), PhN_3_, phenylazide; BCN, biscyclononyne. **e**, Normalized absorbance spectra of fluorogenic CF_3_P- and TF-MaP618 substrates (15 µM) in presence or absence of SNAP-tag2/SNAP_f_-tag protein (30 µM). Fold changes are referred to the absorbance of the dye only. SNAP-tag2 shows an improved increase in absorbance of fluorogenic substrates by around 4.9-fold for CF_3_P- and 2.3- fold for TF-MaP618 compared to SNAP_f_-tag.

We then tested the performance of SNAP-tag2 and its new substrates in live-cell imaging applications. We first compared the labeling kinetics of SNAP-tag2 and SNAP_f_-tag with different fluorescent probes (i.e., TMR, carbopyronine (CPY) and silicon rhodamine (SiR)) in live U2OS cells. The fluorophores were coupled to either CP, CF or TF (Supplementary Fig. 1) to identify the best substrate-fluorophore pair for each fluorophore individually to use in live-cell labeling. We used U2OS cells stably co-expressing nuclear-localized HaloTag7-SNAP-tag2 or HaloTag7-SNAP_f_-tag fusions together with mTurquoise2, which was used for normalization of the protein expression level. Cells were incubated with the respective fluorescent probes and the fluorescence intensity changes were recorded as a function of time. For all tested fluorescent substrates, SNAP-tag2 shows significantly faster labeling than SNAP_f_-tag with CP-substrates (Fig. 3a-d). Among the various fluorophores, the differences between SNAP-tag2 and SNAP_f_-tag are most pronounced for the labeling with SiR-derivatives (Fig. 3c-d). The half-labeling time (t_1/2_) for reaching saturation of the fluorescence signal was around 20 min for SNAP-tag2 labeled with CF-SiR, whereas SNAP_f_-tag shows only relatively weak labeling with CP-SiR under the same experimental conditions, not reaching a plateau within 90 min. (Fig. 3c-d). For all CPY substrates, SNAP-tag2 shows ∼4-fold faster labeling than SNAP_f_-tag. The t_1/2_ values for SNAP-tag2 labeling with different CPY substrates are similar (t_1/2_ = 10 - 12 min.), highlighting a small influence of the substrate core for live cell labeling. For TMR substrates, the differences between SNAP-tag2 and SNAP_f_-tag are less pronounced (1.6 - 2.2-fold). This suggests that live cell labeling with TMR-substrates is limited by the entry of the substrate into the cell.

**Fig. 3.**
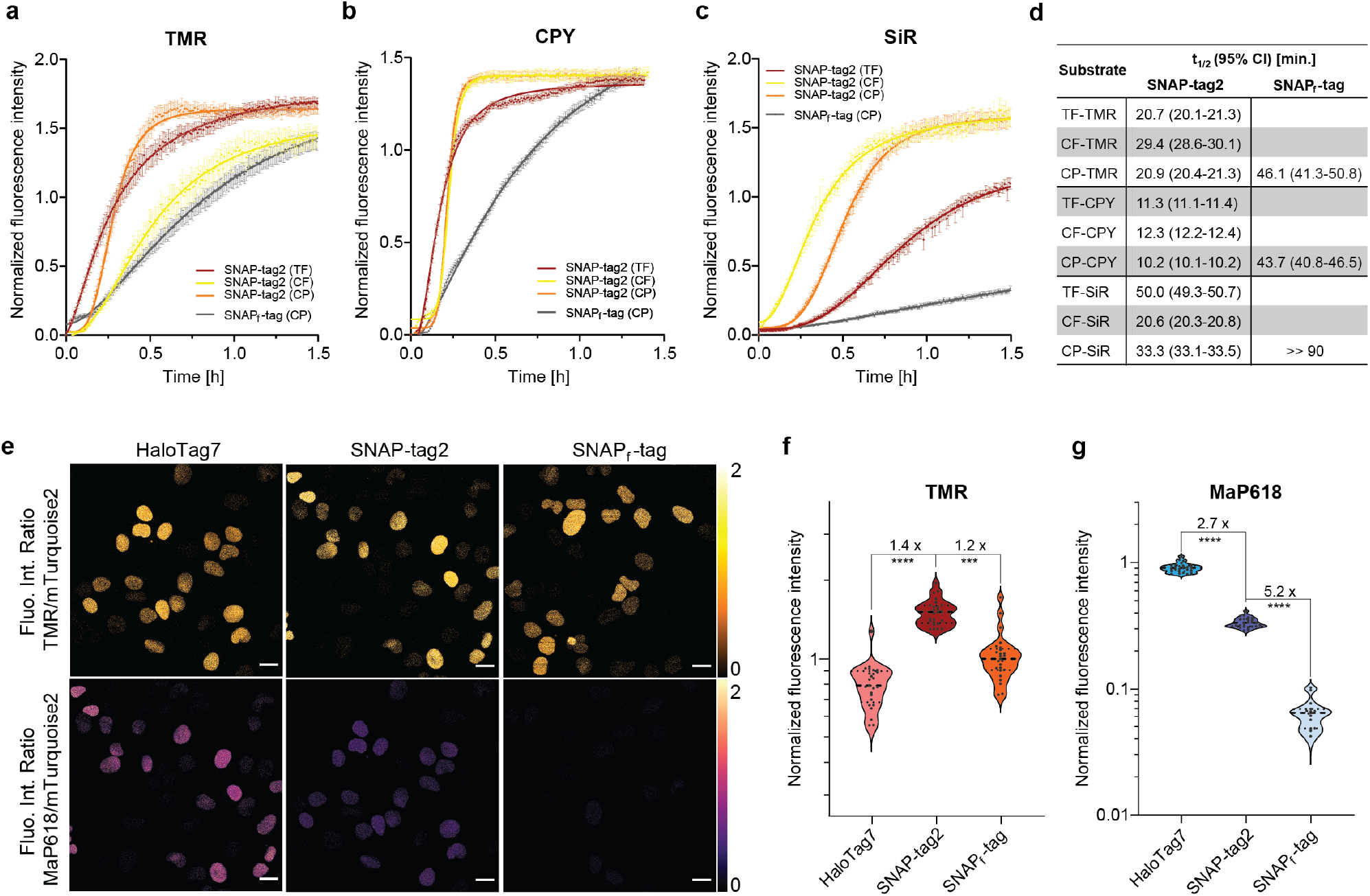
Labeling kinetics and fluorescence brightness for SNAP-tag2 in live cells. Experiments were conducted in live U2OS cells stably co-expressing HaloTag7-SNAP_f_-tag or HaloTag7-SNAP-tag2 together with mTurquoise2 (expression marker) in the nucleus. **a-c**, Kinetic traces of SNAP-tag2 and SNAP_f_-tag fluorescent labeling in live cells with TMR (a), CPY (b) and SiR (c) substrates. U2OS cells were labeled with TF-, CF- and CP-fluorophore substrates (50 nM for TMR and CPY, 100 nM for SiR) and the labeling reaction was followed *via* confocal fluorescence microscopy. Fluorescence intensity changes of the substrates were normalized to the mTurquoise2 fluorescence over time. Data was fitted to a sigmoidal curve. Representative result from a biological duplicate. **d**, Half-labeling times (t_1/2_) of SNAP-tag2 and SNAP_f_-tag calculated from in-cell kinetic measurements as described in a-c. Average from biological duplicate. **e**, Comparison of HaloTag7, SNAP-tag2 and SNAP_f_-tag fluorescence brightness in confocal fluorescence microscopy with non-fluorogenic TMR and highly fluorogenic MaP618 substrates. U2OS cells were labeled with SLP-respective CA-/TF-TMR and -MaP618 substrates (100 nM overnight). Ratiometric projections are presented corresponding to fluorescence intensities (Fluo. Int.) of label/mTurquoise2 using orange-hot (TMR) and mpl-magma (MaP618) look-up tables. Scale bar: 20 µm. **f-g**, Quantitative analysis of single cells shown in e represented as violin plots. Numbers represent fold-changes between the different SLPs (n = 15-25 cells, Welch t-test: **** ≙ p < 0.0001, *** ≙ p = 0.0003).

Similar results were obtained in a flow cytometry-based assay for the labeling of mEGFP-SNAP-tag2 and mEGFP-SNAP_f_-tag fusion proteins in U2OS cells. All tested substrates lead to a more efficient labeling of SNAP-tag2 than SNAP_f_-tag with the most pronounced differences observed for SiR- and MaP618-substrates (Supplementary Fig. 5).

Moreover, we compared the fluorescence brightness of labeled SNAP-tag2 in live cells to that of SNAP_f_-tag and HaloTag7. Previously described U2OS cells stably co-expressing nuclear-localized HaloTag7-SNAP-tag2 or HaloTag7-SNAP_f_-tag fusions together with mTurquoise2 were labeled overnight with either CA-/TF-TMR or -MaP618 to achieve full labeling (Fig. 3e). Quantification of the SLP fluorescence signal normalized to mTurquoise2 expression shows smaller differences between all three SLPs for TMR labeling (Fig. 3f, Supplementary Fig. 6c). Similar results were obtained for SiR-substrates (Supplementary Fig. 6a-b, f) and CPY-substrates (Supplementary Fig. 6d), for which differences in the fluorescence intensities between all SLPs, including Hal-oTag7, remain below 2-fold. For labeling with TF-MaP618, SNAP-tag2 shows a 5.2-fold higher fluorescence intensity than SNAP_f_-tag in cells (Fig. 3g, Supplementary Fig. 6e), which is in line with the measured absorbance difference *in vitro* (Fig. 2e). However, the brightness of MaP618-labeled SNAP-tag2 remains 2.7-fold dimmer than that of MaP618-labeled HaloTag7. Thus, SNAP-tag2 probably shifts the equilibrium of spirocyclization of very closed rhodamine dyes more towards the zwitterionic form than SNAP_f_-tag.

We further tested the performance of SNAP-tag2 in super-resolution stimulated emission-depletion (STED)^[28]^ microscopy. HeLa cells stably co-expressing HaloTag7, SNAP-tag2 or SNAP_f_-tag fused to the Cox8a pre-sequence (mitochondrial matrix localization) and mEGFP as an expression marker (no specific localization) were labeled with their respective CA- or TF-SiR substrates (Fig. 4a). SNAP-tag2 shows much stronger fluorescence labeling inside mitochondria, both in confocal laser scanning microscopy (CLSM) and STED microscopy than observed for SNAP_f_-tag under the same experimental conditions, even though a cell with stronger mEGFP fluorescence (≙higher expression level) was chosen for SNAP_f_-tag. Detection of the dim SNAP_f_-tag labeling required a significant increase in laser power (Supplementary Fig. 7). The observed fluorescence signal for SNAP-tag2 is comparable to the signal obtained for the corresponding HaloTag7 labeling at similar expression levels. Similar differences in fluorescence intensities were observed for SNAP-tag2 and SNAP_f_-tag labeling of intermediate filaments (Vimentin fusion; Fig. 4b). Because of the increased photon-count, SNAP-tag2 shows an improved STED resolution as approximated by the reduced full-width half-maximum (FWHM) of vimentin fibers (108 ± 23 nm) compared to SNAP_f_-tag (146 ± 7.0 nm). We also tested the STED performance of SNAP-tag2 with other fusion proteins such as Sec61b (outer endoplasmic reticulum membrane; Fig. 4c) and TOMM20 (outer mitochondrial membrane; Fig. 4d) and observed bright and specific labeling of SNAP-tag2 with CF-SiR. Furthermore, we performed dual-color live-cell STED imaging in live U2OS cells expressing Vim-SNAP-tag2 and LifeAct-HaloTag7 (Actin) using CF-SiR and CA-MaP618, respectively (Fig. 4e). These results demonstrate the superiority of SNAP-tag2 over previous SNAP-tag versions in live-cell imaging experiments.

**Fig. 4.**
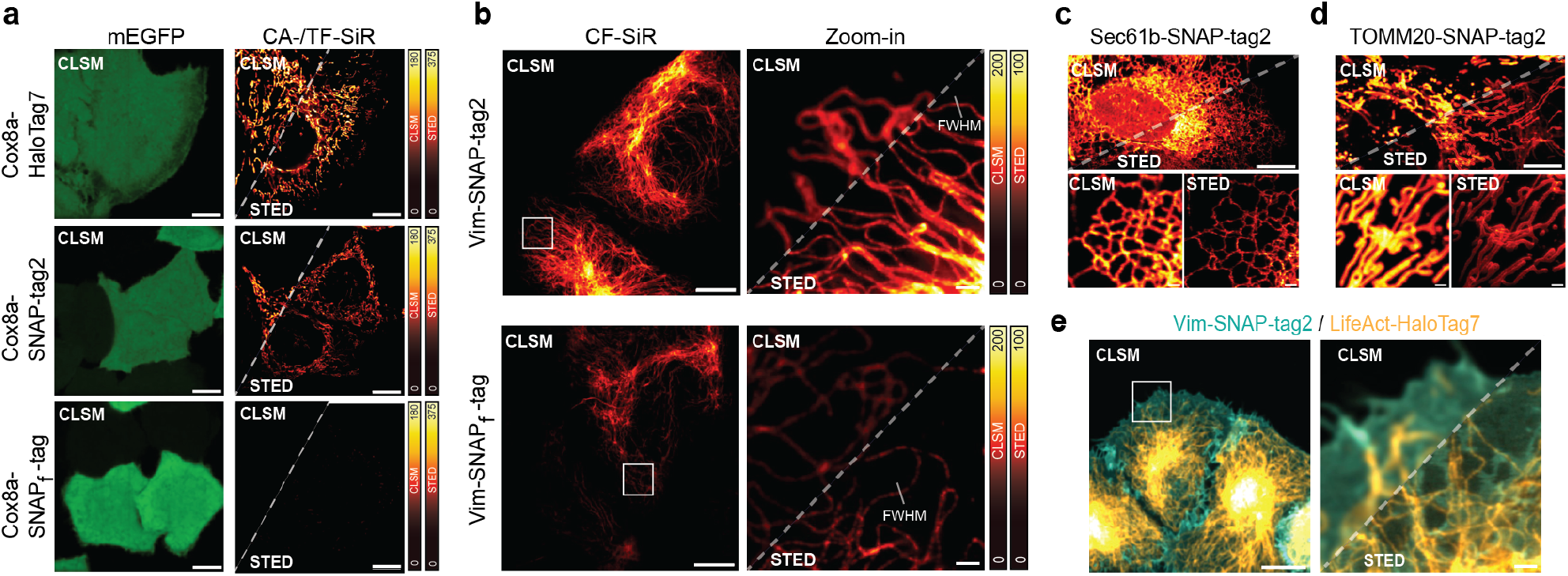
SNAP-tag2 performance in live cell super-resolution microscopy. **a**, Comparison of HaloTag7, SNAP-tag2 and SNAP_f_-tag performance in confocal laser scanning microscopy (CLSM) and stimulated emission depletion (STED) microscopy. HeLa cells stably co-expressing HaloTag7, SNAP-tag2 or SNAP_f_-tag in the mitochondria (Cox8a localization sequence) together with mEGFP (no specific localization) were labeled with CA-/TF-SiR (100 nM) for 1 h. SNAP-tag2 and HaloTag7 show comparable performance in STED imaging, while SNAP_f_-tag shows insufficient signal under the same imaging conditions. Scale bars: 10 µm. LUTs: green (mEGFP), red-hot (SiR) **b**, CLSM and STED images of U2OS cells stably expressing Vim-SNAP-tag2 (b) or Vim-SNAP_f_-tag (c) labeled with CF-SiR (100 nM) for 1 h. White squares in the overview images (left) highlight area chosen for magnification and STED imaging (right images). The Full Width at Half Maximum (FWHM) of single intermediate filament fibers highlighted in b was determined to be 108 (± 23) nm and 146 (± 7.0) nm, respectively, underlining a higher resolution achieved with SNAP-tag2 compared to SNAP_f_-tag (15 individual fibrils and 3 individual images each). Scale bars: 5 µm (overview), 1 µm (magnification). **c-d**, CLSM and STED images of U2OS cells stably expressing SNAP-tag2 (**c**) in the endoplasmic reticulum (ER) and (**d**) on the outer mitochondrial membrane (TOMM20) labeled with CF-SiR (100 nm) for 1 h, demonstrating the versatility of using SNAP-tag2 in different cellular compartments. Scale bars: 5 µm (overview), 1 µm (magnification). **e**, Dual-color CLSM and STED images of U2OS cells expressing Vim-SNAP-tag2 and LifeAct-HaloTag7. SNAP-tag2 was labeled with CF-SiR (100 nM) and HaloTag7 with CA-MaP618 (100 nM) for 1 h. Scale bars: 5 µm (overview), 1 µm (magnification). LUTs: cyan (SNAP-tag2-SiR), orange-hot (HaloTag7-MaP618).

We then investigated, if the improved SNAP-tag2 system translates to more efficient fluorescence labeling in yeast. Chemical labeling in yeast is challenging, presumably due to the presence of its cell wall, an additional permeability barrier, as well as the expression of various multidrug efflux pumps.^[29]^ The peroxisomal membrane protein Pex3 plays a key role in peroxisomal biogenesis, inheritance and degradation in yeast^[30]^. To study the precise localization, dynamics and distribution of Pex3 and its binding partners, live-cell STED nanoscopy is a suitable technique, since it allows to quantify the exact number and size of peroxisomes in live yeast.^[31]^ Thus, we expressed either Pex3-SNAP-tag2 or Pex3-SNAP_f_-tag in the methylotrophic yeast species *Hansenula poly-morpha* (*H. polymorpha*) and tested different SiR-substrates for labeling of peroxisomes in live-cells (Fig. 5a). *H. polymorpha* expressing Pex3-SNAP-tag2 showed homogenous and highly efficient labeling of peroxisomes, whereas for yeast cells expressing Pex3-SNAP_f_-tag the fluorescence signal was much weaker (Fig. 5a-b). The correct localization of Pex3-SNAP-tag2 was furthermore confirmed through STED microscopy (Fig. 5c, Supplementary Fig. 9a-b). Similar results were obtained for labeling with MaP555^[7]^-substrates, for which SNAP-tag2 works best in combination with CP-MaP555 (Supplementary Fig. 8 and 9c-d). These results demonstrate that SNAP-tag2 allows for more efficient labeling in *H. polymorpha* yeast than its predecessor.

**Fig. 5.**
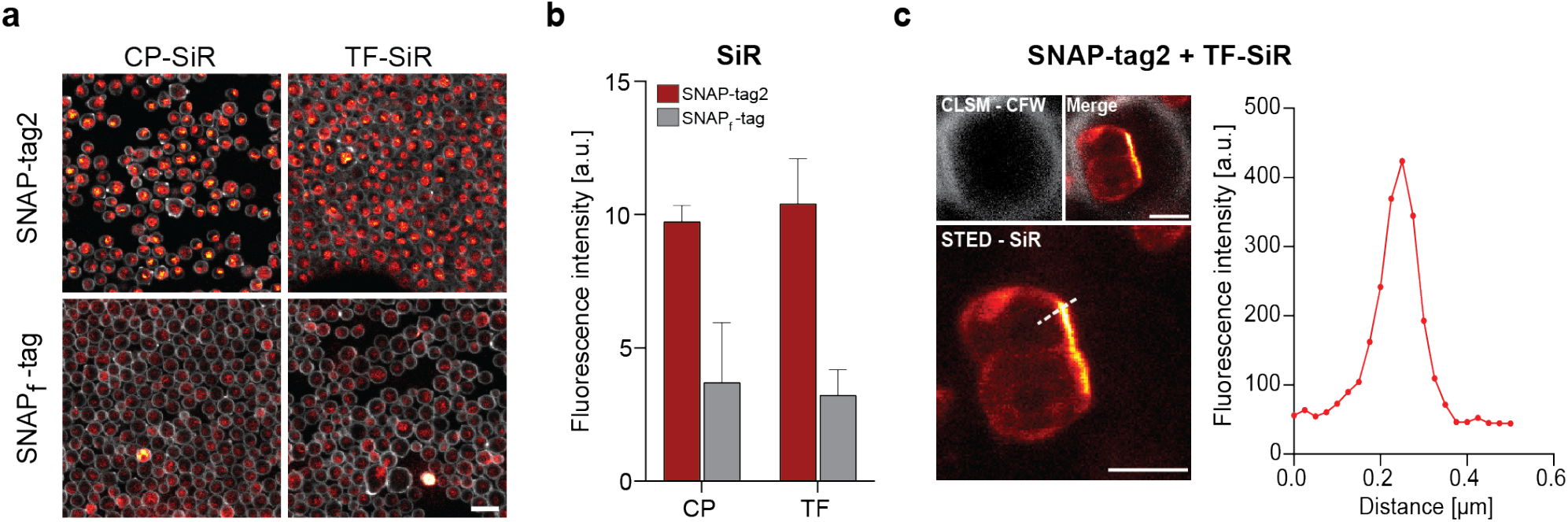
Comparison of SNAP-tag2 and SNAP_f_-tag labeling of yeast peroxisomes. **a**, CLSM images of *H. polymorpha* yeast cells expressing Pex3-SNAP-tag2 or Pex3-SNAP_f_-tag fusion proteins labeled with different SiR substrates. Yeast cells were labeled with CP- or TF-SiR (250 nM) for 18 h and the cell wall was stained with Calcofluor White (CFW; 25 µg/mL) for 15 min. Scale bar: 10 µm. **b**, Bar plot representing the quantitative analysis of SNAP-tag2 and SNAP_f_-tag labeling with SiR substrates in yeast. Experiments were conducted in biological triplicates and the mean fluorescence intensity of SiR substrates was calculated for 3 × 125 cells (Supplementary Fig. 10). SNAP-tag2 greatly outperforms SNAP_f_-tag in labeling of live yeast peroxisomes. **c**, STED image of Pex3-SNAP-tag2 labeled with TF-SiR (lower panel), CLSM image of CFW-stained cell wall and merge of both channels (upper panels). Scale bar: 1 µm. The right plot shows the line profile of labeled peroxisomes in STED imaging (highlighted as dashed line in the image). SNAP-tag2 with TF-SiR is suitable to perform live cell STED microscopy in yeast.

## Discussion

In this work we used a combination of substrate optimization and protein engineering to develop SNAP-tag2, an improved version of the SNAP-tag labeling system. SNAP-tag2 reacts faster with a new set of pyrimidine-based substrates. An increased labeling speed is observed for a large variety of probes, including rhodamines as well as different reactive biorthogonal groups. The apparent second-order rate constant for the reaction of SNAP-tag2 with the new substrate TF-TMR *in vitro* approaches 10^7^ M^-1^s^-1^, which is about a 100-fold faster than the labeling reaction of SNAP-tag with CP-TMR and is close to that of HaloTag7 with CA-TMR. SNAP-tag2 remained a very stable protein with a melting temperature of ∼65 °C, which might facilitate further engineering attempts such as circular permutations and the generation of split-versions of SNAP-tag2. This could allow for the development of novel biosensors and recorders for studying biological activities in cells and/or *in vivo*.^[32]^ Its size of 172 amino acids is slightly below that of its predecessor and significantly below that of HaloTag7 with 297 amino acids. SNAP-tag2 displaces the spirocyclization equilibrium of bound rhodamines more towards the open form compared to SNAP_f_-tag, a property for which we directly screened in the directed evolution experiments. We assume that a possible interaction of SNAP-tag2 with the zwitterionic, fluorescent configuration of rhodamines results in higher brightness of SNAP-tag2 when labeled with the very fluorogenic dye MaP618. Nevertheless, SNAP-tag2 displaces the spirocyclization equilibrium still to a lower degree than HaloTag7. Since very closed rhodamine-derivatives tend to show reduced labeling performances in cellular experiments^[13]^ and *in vivo*^[33]^, we believe that this is not a major limitation of SNAP-tag2 relative to HaloTag7.

Most importantly, the increased reactivity of SNAP-tag2 towards its new substrates translates into a more efficient fluorescence labeling in live cells. This was most pronounced for fluorescence labeling with the far-red SiR, which enabled efficient fluorescence labeling at substrate concentrations that only resulted in weak labeling of SNAP_f_-tag. The relatively weak SiR-labeling observed for SNAP_f_-tag is in line with previous reports by others^[20]^. In live-cell imaging, the different fluorophores showed some preferences among the different SNAP-tag substrates. For SiR, CF-SiR showed faster labeling in U2OS cells than TF-SiR and CP-SiR, even though all reached the same brightness after longer incubation periods. The labeling of SNAP-tag2 in live cells with TMR- and CPY-substrates showed less dependency on the nature of the substrate (TF, CF and CP). For the preparation of any new probes for live cell applications with SNAP-tag2 or for new applications of the here described probes, we thus recommend to first test the CF-based probes. The greater performance of SNAP-tag2 for fluorescent labeling in live cells also extends to super-resolution microscopy imaging techniques such as STED microscopy as well as to other cell types known to be refractory to chemical labeling such as yeast cells. As SNAP-tag2 together with its new substrates is superior to the previously used SNAP-tag versions in every application we have tested, we expect that it will also excel in *in vivo* applications. Finally, implementation of the new SNAP-tag2 system to already existing SNAP-tag-based biosensors such as Snifits^[34]^ should certainly enhance their performance.

In summary, the improvements introduced in SNAP-tag2 are expected to advance the utility of this already widely adopted tool for live-cell imaging and other applications in life sciences.

## Methods

### Synthesis and general methods

Detailed procedures for the synthesis and chemical characterization of all compounds are given in the Supplementary Methods. Buffer, media and reagent compositions can be found in Supplementary Table 12.

### Plasmids for bacterial and mammalian protein expression

A pET51b(+) vector (Novagen) was used for protein production in *Escherichia coli*. Proteins were N-terminally Strep- and C-terminally Hisx10-tagged for affinity purification. pcDNA5/FRT or pcDNA5/FRT/TO vectors (ThermoFisher Scientific) were used for protein expression in mammalian cells *via* transient transfection or for stable cell line generation. Cloning was performed with Gibson assembly or point-mutations were introduced by site-directed mutagenesis using the Q5®-site-directed mutagenesis kit (NEB) according to the manufacturer protocol. Plasmid sequences were verified by Sanger sequencing (Eurofins Genomics or Microsynth) and plasmids were stored at -20 °C.

### Recombinant protein production and purification

pET51b(+) plasmids were transformed in *E. coli* BL21(DE3)-pLysS (Novagen) cells. LB-Amp expression cultures were grown at 37 °C to an optical density at 600 nm (OD_600_) of 0.6-0.8 and protein expression was subsequently induced at 16°C by addition of isopropyl β-thiogalacopyranoside (0.5 mM). After overnight expression, cells were harvested by centrifugation (4’000 ×g, 4 °C, 15 min.), resuspended in His-tag extraction buffer (20 – 30 mL) and lysed by sonication (50 % duty, 70 % power, 7 min.) on wet ice. All proteins were purified *via* immobilized metal ion affinity chromatography (IMAC) either by gravity column or over an ÄktaPure FPLC instrument (Cytiva) equipped with a HisTrap FF crude column (Cytiva, Marlborough, MA). Buffer was exchanged to activity buffer (50 mM HEPES, 50 mM NaCl, pH = 7.3) either using Zeba™ Spin desalting columns (7 K MWCO, 0.5 mL, Thermo Scientific) or over a HiPrep 26/10 desalting column (Cytiva) on the FPLC system. Purified proteins were concentrated using Amicon Ultra centrifugal filters (10 kDa MWCO). The purity and correct size of the proteins were assessed by sodium dodecyl sulfate–polyacrylamide gel electrophoresis (SDS– PAGE) and high-resolution mass spectrometry (HRMS). Purified proteins were aliquoted, flash frozen in liquid nitrogen and stored at -80 °C.

### Plasmids for yeast surface display (YSD)

For protein expression on the yeast surface, either a pCTcon2 vector (Addgene, #73152) with SNAP-tag proteins fused in-between Aga2p-HA-tag (N-terminal) and cMyc-tag (C-terminal) or a pJYDNg vector (Addgene, #162452) with a C-terminal fusion to Aga2p-HA-tag-cMyc-tag-eUnaG2 was used. Epitope-tags HA- and cMyc-tag were used for immunostaining and the fluorescent reporter protein eUnaG2 was labeled with bilirubin for monitoring the protein expression level.

### DNA library preparation for YSD

Libraries were created either on pCTcon2-SNAP-tag1.1 (SNAP-E30R-S135R-L153G) *via* one-pot saturation mutagenesis as published by Wrenbeck et al.^[36]^ or on pJYDNg-SNAP-tag1.5 (SNAP-E30R-I32Y-L34V-K36N-^50^GHPEPQ-S135R-L153G-G161P) using site-directed saturation mutagenesis in combination with assembly PCR.^[37]^ Libraries were generated using a mixture of degenerate primers, each including one or four NNK over the target regions (amino acids H29-K36 and 156-161; Supplementary Tables 14-17). Library DNA was transformed into *E. cloni* cells and cells were recovered in 2 mL of SOC medium (New England Biolabs) for 1 h. Serial dilutions were spread on agar plates containing kanamycin (50 µg/mL) and incubated at 37 °C overnight. Remaining cells were transferred to 3 mL of LB-Kan medium and the culture was grown at 37 °C overnight. Colonies on plates were counted to determine the library size and the isolated plasmids from ten single picked colonies were sequenced to control the library diversity. The plasmid library was isolated at culture saturation using a QIAprep Spin Miniprep kit (Qiagen). Libraries on the active site loop of pJYDNg-SNAP-tag1.5 were generated using degenerated primers with three NNKs at different positions throughout the targeted region (156-GGYEGP-161). Primers were designed with central NNKs and 20 bp flanking regions for assembly *via* PCR. Forward and reverse primers were designed to be completely overlapping to each other. The protocol for assembly PCR was adjusted from Routh et al. Library inserts were purified using a QIAquick PCR Purification kit (Qiagen). A synthetic deep mutational scanning library (sDMSL) designed on the SNAP-tag1.5 scaffold was purchased from Twist Bioscience. The library consists of single point variants, having each amino acid along the SNAP-tag1.5 sequence substituted to all other 19 amino acids, excluding cysteines and including deletions. Libraries were used for homologous recombination in EBY100 yeast cells for YSD screening.

### Electrocompetent yeast cell preparation for YSD

Single colonies of *S. cerevisiae* strain EBY100 (ATCC) spread on a YPD agar plate were used for inoculation of YPD liquid cultures. YPD expression cultures were grown at 250 rpm and 30 °C until an OD_600_ of 1.3-1.5. Afterwards, Tris-DTT buffer (800 µL) and Tris-LiAc buffer (2 mL) were added and the culture was incubated at 250 rpm and 30 °C for 15 min. Cells were harvested by centrifugation (2’500×g, 4 °C, 3 min.) and washed with ice-cold electroporation buffer (25 mL), followed by cell harvesting (2’500 ×g, 4 °C, 3 min.). Cells were resuspended in electroporation buffer to a total volume of 300 µL. Cells were aliquoted (50 µL) and directly used for transformation by electroporation. Remaining cells were frozen and stored at -80 °C.

### Yeast cell transformation and homologous recombination for YSD

For circular plasmid DNA transformation, DNA (1-3 µg) was diluted in ddH2O (< 10 µL) and kept on ice. For homologous recombination (HR), amplified vector (1 µg) and insert DNA (2-3 µg) were mixed to a final volume of 100 µL. The DNA was precipitated by addition of NaOAc (10 µL, 3 M), glycogen (0.5 µg/µL final conc., ThermoFisher) and isopropanol (100 µL) and stored at -20 °C overnight. Afterwards, the mixture was pelleted by centrifugation (10’000 rpm, 20 min.) and the DNA pellet was washed with 70 % EtOH (200 µl) followed by centrifugation (10’000 rpm, 20 min.). The supernatant was removed and the pellet was air-dried. Electrocompetent EBY100 yeast cells (50 µL) were used to resuspend the DNA and transformation was done *via* electroporation. Electroporation cuvettes (Gene Pulser® Cuvette, 0.2 cm electrode gap, Bio-Rad Laboratories) were pre-chilled on ice. The cell-DNA mix was transferred to the pre-chilled electroporation cuvette and electroporation was done on a Bio-Rad GenePulser Xcell™ device (0.54 kV, 25 µF, infinite resistance with an exponential decay wave form). After electroporation, pre-warmed YPDS medium (1 mL) was immediately added to the cuvette and the cell suspension was transferred to culture tubes (15 mL). The cuvette was washed with 1 mL of YPDS medium and added to the culture. The culture was incubated at 250 rpm and 30 °C for 1 h. Yeast cells were harvested by centrifugation (2’500×g, 5 min.) and resuspended in SDCAA medium (1 mL). Serial dilutions were plated on SDCAA agar plates to determine the transformation efficiency. Plates were incubated at 30 °C for 2-3 days. The remaining cells suspension was used to inoculate SDCAA culture (100 mL) and grown at 30 °C for 2 days. The culture was directly used for yeast surface display screens or aliquots were frozen in a 1:1 mixture of culture:50% glycerol in Tris-buffer (pH = 8).

### Protein expression on the yeast surface

SDCAA medium (5 mL final volume) was inoculated either using a single colony from a SDCAA plate or by addition of fully grown yeast liquid culture (0.5 mL) after transformation. The culture was incubated at 250 rpm and 30 °C overnight. This pre-culture (0.5 mL) was used to inoculate 4.5 mL of SGCAA medium for protein expression on the yeast surface. Protein expression was conducted for at least 20 h at 250 rpm and 30 °C.

### Protein labeling on yeast surface for fluorescence-activated cell sorting (FACS)

For fluorescent labeling, 10^7^ cells were harvested by centrifugation (14’000×g, 1 min.). The concentration of the yeast culture expressing the protein(s) of interest on the surface was determined by accounting that an OD_600_ = 1 corresponds to about 10^7^ cells/mL.^[38]^ For antibody-based expression staining of yeast cells transformed with pCTcon2 encoded protein libraries, yeast cells were resuspended in 1:10-diluted primary mouse anti-cMyc antibody (#OP10, EMD Millipore, Merck) in PBS (50 μL) and incubated on a rotating wheel at 4 °C for 1 h. The cells were pelleted (14’000×g, 1 min.) and washed twice with PBS (125 μL) with centrifugation in-between (14’000×g, 1 min.). Afterwards, the cell pellet was resuspended in 1:50-diluted secondary goat anti-mouse-Alexa647 antibody (#A-21236, Invitrogen, ThermoFisher Scientific) in PBS (50 μL) and incubated on a rotating wheel at 4 °C for 1 h. The cells were again washed twice with PBS (125 μL) prior to labeling with SNAP-tag substrates. For protein libraries encoded on the pJYDNg expression vectors, expression control was monitored over labeling of eUnaG2 with bilirubin. Yeast cells were resuspended in PBS (50 μL) containing bilirubin [10 μM] and BSA [1 mg/mL], gently vortexed and incubated on ice for 10 min. The cells were pelleted and washed twice with PBS (150 μL) with centrifugation in-between (14’000×g, 1 min.) prior to SNAP-tag labeling. Cells expressing SNAP-tag variants were labeled in PBS (50 μL) using different substrates (CF_3_P-TMR/-MaP618, TF-TMR/MaP618) with varying concentrations [10-500nM] and incubation times (10-60 min) in order to tune screening stringency. Labeling was performed at r.t. on a rotating wheel. Cells were washed with PBS (125 μL), resuspended in 1 mL of PBS and filtered through 5 mL round bottom polystyrene test tubes with cell strainer snap caps (#352235, Falcon®) for FACS.

### Protein library screening using FACS

Labeled yeast libraries were analyzed and subsequently sorted on a BD FACSMelody™ Cell Sorter (BD Biosciences) using the appropriate filter settings for each fluorophore (Supplementary Table 6). For yeast cell sorting, a 100-micron sorting nozzle and a 1.5 neutral density filter was used. Photomultiplier tube (PMT) detector voltages were optimized based on single and double labeled samples and negative control samples. The range for positive signal was set to be approximately 10^3^ – 10^4^, while negative and background staining was set to ≤10^2^. Gating strategies are depicted in Supplementary Fig. 11. Cells were gated for live and single cell events based on their size in the forward scatter (FCS) and sideward scatter (SSC). The single cell population was then gated for double positive labeling signal and the top 0.5 – 1 % of double labeled cells (10^4^ cells) were sorted in bulk. After sorting, yeast cells were grown in SDCAA media (5 mL) supplement with penicillin-streptomycin (Gibco, 5’000 U/mL, 1:100 dilution) for 2 days. Plasmids were isolated from fully grown cultures (1 mL) using a Zymoprep™ Yeast Plasmid Miniprep II kit (Zymo Research) according to the manufacturer’s protocol and the remaining culture was propagated to start the next screening round.

### Computational redesign of the unstructured loop in SNAP-tag

To redesign the unstructured loop (residues 37-54), the TMR labeled SNAP-tag structure (PDB-ID: 6Y8P)^[14]^ was used as input. The covalent benzyl-TMR ligand was parameterized and a Rosetta constraint file was set up to fix the covalent bond between the ligand the protein analogous to published procedures.^[32]^ Point mutations E30R, I32Y, L34V, K36N, S135R, L153G and G161P were modeled and the structure was minimized using a RosettaScripts^[39]^ protocol. The top scoring structure was used as input for the loop design using the RosettaRemodel^[27]^ application. Blueprint files for 7, 8 and 9 amino acid loops were set up, allowing any amino acids except cysteine in the designed regions (example script can be found in the Supplementary information). The top 5 scoring sequences for each loop length were tested experimentally.

### Next-Generation Sequencing (NGS)

Randomized regions from isolated plasmids were amplified with NGS primers (Supplementary Table 18) using standard PCR conditions and purified (Qiagen PCR purification kit). NGS was performed using adaptor ligation Ilumina sequencing at Eurofins Genomics. The NGS package for each sample included 10 million total reads comprising of 2 × 150 bp paired-end reads (5 million read pairs). NGS amplicon sizes varied from 150-300 bp and were pooled in equivalent quantities, since the NGS primers’ unique adapter sequences enable to sort them at data analysis. NGS samples comprised 2 μg of DNA in 100 μL of ddH_2_O.

### Analysis of NGS data

Illumina sequencing results were first split into separate files based on their experiment- and selection-round-specific sequence barcodes using “je demultiplex”^[40]^. Forward and reverse reads were combined into a single file for each barcode. Primer and adapter sequences were trimmed using cutadapt^[41]^. Trimmed reads were then aligned to the template sequence with bowtie2^[42]^. The resulting ‘sam’ file was converted to a ‘bam’ file using samtools^[43]^. The alignments were then analyzed with a custom R script to calculate amino acid frequencies for randomized positions (for saturation mutagenesis libraries) or point mutation frequencies (for deep mutational scanning libraries). Reads were trimmed to match the translation frame of the template sequence. Reads containing deletions, insertions relative to the template sequence, or ambiguous bases were removed. The remaining reads were translated into amino acid sequences. For saturation mutagenesis libraries, amino acid frequencies at the positions of interest were calculated by dividing the number of reads featuring a specific amino acid at a particular position by the total number of reads. For deep mutational scanning libraries, mutation frequencies were calculated by dividing the number of reads featuring a specific mutation by the number of reads that cover the corresponding position. The script used for NGS data analysis can be found in the Supplementary Information.

### Protein thermal stability analysis

Thermal stabilities of final protein variants were measured on a Prometheus NT48 nanoscale differential scanning fluorimeter (NanoDSF). Protein samples were prepared in activity buffer (0.8 mg/mL) and changes in tryptophan fluorescence at 330 nm were followed over a temperature range of 20 - 95 °C with a temperature increase of 1 °C/min.

Measurements were done in technical triplicates. The inflection point of the first derivative corresponds to the proteins melting temperature.

### *In vitro* labeling kinetics measured *via* a microplate reader fluorescence polarization assay

Labeling kinetics were measured in black non-binding flat bottom 96-well plates (200 μL final reaction volume) or in black non-binding low volume 384-well plates (20 μL final reaction volume) (Corning Inc.) on a microplate reader (Spark 20M, Tecan) by recording fluorescence polarization (FP) traces at 37 °C over time. All measurements were performed in technical triplicates in FP buffer. Labeling reactions were started by either addition of substrate to the protein using a multi-channel pipette or using the injector module of the plate reader for fast reaction conditions. Filter settings were chosen according to the fluorophore substrate (Supplementary Table 7). The G-factors were calculated using a buffer only (blank) and free fluorophore substrate (reference) control. The optical gain, an amplification factor for the photomultiplier tube, was adjusted to 50 %. Baselines were determined by recording FP kinetic traces of the free fluorophore substrates.

For screening purposes, labeling kinetics were recorded using 50 nM of protein and 20 nM of fluorophore substrate. For fast labeling kinetics, FP kinetics were conducted with 10 nM protein and 2 nM substrate in 384-well plates. A one-phase association equation (2) was fitted to the data using GraphPad Prism and apparent second-order rate constants (k_app_) were calculated using equation (1).

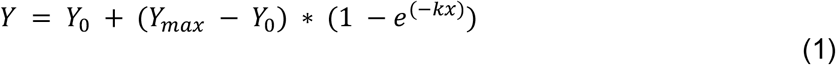

With Y = fluorescence polarization [mFP], Y_max_ = FP plateau [mFP], Y_0_ = y-intercept [mFP], k = labeling rate constant [s^-1^] and x = time [s].

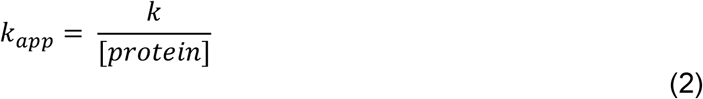

For more accurate determination of apparent second-order rate reactions (k_app_), a global fit approach to kinetic models was used as published in Wilhelm et al.^[14]^ In brief, kinetics were measured at fixed fluorophore substrate concentration [20 nM] and various protein concentrations [0 & 4.60 – 900 nM]. Fast kinetics were measured at fluorophore substrate concentrations of 2 nM and protein concentrations varying between 0 & 0.78 - 50 nM. Kinetic data were pre-processed using a custom R script ^[44-45]^and subsequently, kinetic model 1 (equation (3)) was fitted to the data using DynaFit.^[46]^ The delay time, FP baseline value and protein concentration were fixed parameters, while the fluorophore substrate concentration was adjustable to account for potential quantification errors. Including limiting protein concentration into the experimental conditions, allowed to ensure accurate concentration fitting by impacting the final FP value (decreasing plateau). Standard deviations and confidence intervals were obtained using a Monte Carlo simulation^[47]^ with standard settings (N = 1000, 5 % worst fits excluded). Representative graphics were generated using a custom-made R script.

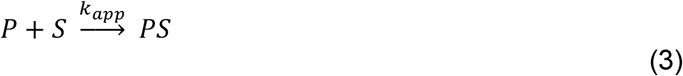

With P = SNAP-tag2 protein, S = fluorophore substrate, and PS = fluorescently labeled SNAP-tag2.

Labeling kinetics of non-fluorescent substrates were determined in a competition assay as described in Wilhelm et al.^[14]^

### *In vitro* labeling kinetics of SNAP-tag2 using a stopped-flow device

Labeling kinetics of SNAP-tag2 with TF-, CF-, and CP-TMR and -CPY substrates were measured were measured by recording fluorescence anisotropy changes over time using a BioLogic SFM-400 stopped-flow instrument (BioLogic Science Instruments, Claix, France) in a single-mixing configuration at 37 °C as described in Wilhelm et al.^[14]^ However, the final substrate concentration was fixed to 0.5 or 1 μM, and the protein concentration was varied from 0.42 to 3.13 μM in activity buffer (50 mM HEPES, 50 mM NaCl, pH = 7.3) supplemented with 1 mM DTT and 0.1 mg/mL BSA. The anisotropy of the free substrate was measured to obtain a baseline. The sampling time was set to 1 ms for a total recording time of 2 s or varied from 200 µs for 0.2 s followed by 2 ms to a total duration of 3 s. For each condition, 15 technical replicates were recorded. Data analysis was conducted as described in Wilhelm et al.^[14]^, fitting model 2 (equation (4)) to the data.

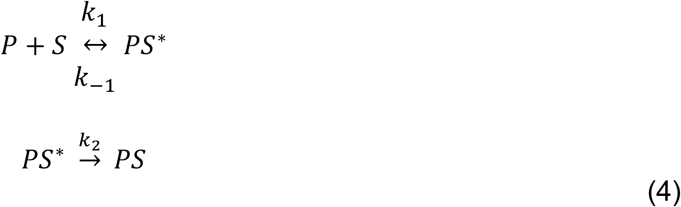

With P = SNAP-tag2 protein, S = fluorophore substrate, PS* = substrate-bound protein complex and PS = fluorescently labeled SNAP-tag2.

Multiple replicates of the experiment were conducted for TF-TMR (4 replicates), CF-TMR (8 replicates) and CP-TMR (3 replicates) as well as their corresponding CPY analogues (2 replicates each). Results of the individual kinetic measurements are depicted in Supplementary Table 2.

Replicates were averaged and standard deviations were calculated (**Error! Reference source not found**., Supplementary Table 3).

### Photophysical properties of labeled SNAP-tag2 and SNAP_f_-tag

For determination of the extinction coefficients (ε) of labeled SNAP-tag proteins, recombinant SNAP-tag2 and SNAP_f_-tag at [30 µM] were labeled with fluorophore substrates at [7.5 µM] in FP buffer overnight. For measurements, a serial dilution of the labeling reaction was performed, targeting final substrate concentrations of 7.5 – 2.22 µM. Absorbance spectra were recorded on a V-770 spectrophotometer (Jasco) in a Quartz cuvette (3 mm). All measurements were performed in technical triplicates and absorbance values were corrected for absorbance at 800 nm using SpectraGryph v1.2. For all fluorophores, the maximal absorbance value at each concentration was fitted to a linear function (equation (5)) and extinction coefficients were calculated from the slope *b*.

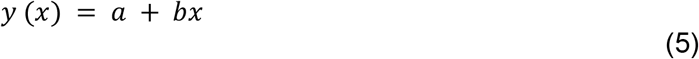

Absolute quantum yields (QYs) of fluorescently labeled SNAP-tag2 and SNAP_f_-tag were measured on a Quantaurs-QY spectrometer (model C11347, Hamamatsu). Measurements were carried out with diluted samples from the same reaction solution used for ε-measurements (A = 0.03-0.08).

### Mammalian cell culture

All media were filtered prior to use employing sterile Nalgene™ Rapid-Flow™ filter units (0.22 μm pore size, 500 mL, Thermo Fisher Scientific). All experimental steps were performed under sterile conditions. Mammalian U2OS Flp In™ T-REx™ cells^[48]^ (hereafter named U2OS cells) or Hela Kyoto Flp-In cells^[32]^ (hereafter referred to as HeLa cells) were cultured in cell growth medium (Dulbecco’s modified Eagle’s medium (DMEM) + GlutaMAX™ supplemented with phenol red, high glucose (4.5 g/L), pyruvate (Gibco Life Technologies, Ref. 31966-021) and 10 % FBS (Gibco Life Technologies)) in either T-25 or T-75 culture flasks (Sarstedt AG & Co.KG). The cultured cells were cultivated in a humidified incubator at 37 °C and 5 % CO_2_ and passaged when reaching 80-90 % confluency. Cell handling involved washing with PBS (1 ×, pH = 7.4, Gibco Life Technologies, Ref. 10010-015), detachment (trypsin 0.05 % + EDTA, TrypLE™ Express, Gibco Life Technologies, Ref. 12604-013), resuspension in DMEM + 10 % FBS and transfer into fresh flasks. For imaging experiments, cells were handled in DMEM without phenol red, referred to as imaging medium (Gibco Life Technologies, Ref. 31053-028) supplemented with GlutaMAX™ (Gibco Life Technologies, Ref. 35050-038), pyruvate (Gibco Life Technologies, Ref. 11360-070) and 10 % FBS (Gibco Life Technologies). For seeding cells at desired densities, cells were counted using a fluidlab R-300 cell counter (anvajo GmbH) with acella 100 chambers (20 μL sample volume, anvajo GmbH). Cells were seeded either in 96-well tissue culture plates (TPP, Ref. 92096) or 8- or 96-well μ-plates with a glass bottom (ibidi GmbH, 8-well Ref. 80827, 96-well Ref. 89627) in a total volume of 100-150 μL of medium.

### Generation of mammalian stable cell lines

Stable cell lines were generated using the Flp In™ T-REx™ system (Thermo Fisher Scientific) with U2OS or HeLa cells. Cells were grown to ∼80 % confluency in T-25 flasks. Lipofectamine™ 3000 (8 μL, Thermo Fisher Scientific, Ref. L3000008) was diluted in Opti-MEM™ I Reduced Serum Medium + GlutaMAX™ (200 μL, Thermo Fisher Scientific, Ref. 51985026) in a 1.5 mL reaction tube and thoroughly mixed. In a separate tube, the plasmid DNA encoding the gene of interest on a pcDNA/FRT or pcDNA/FRT/TO vector (440 ng) and pOG44 (3560 ng; encoding the Flp-In recombinase) were diluted in Opti-MEM™ I Reduced Serum Medium + GlutaMAX™ (200 μL). P3000 (8.0 μL) was added to the DNA mix After thorough mixing, the DNA-P3000 mix incubated at r.t. for 10 min. Subsequently, the DNA-P3000 mix was combined with the diluted Lipofectamine™ 3000, mixed and left to incubate at r.t. for additional 15 min. The resulting DNA-P3000-Lipofectamine™ 3000 mix was added to the T-25 flask and incubated overnight. The following day, the medium was exchanged to cell growth medium supplemented with 100 μg/mL hygromycin B (Gibco Life Technologies), selecting cells for gene of interest incorporation. Selection was performed for 48-72 h. Survived cells were recovered in cell growth medium to reach confluency. Cells were sorted *via* FACS for expression levels by positive fluorescent protein signal (mEGFP, FITC filter) or SLP labeling (SiR, APC filter). Generated stable cell lines can be found in Supplementary Table 13.

### rAAV production and transduction

For dual-color STED imaging, recombinant AAVs (rAAVs) were generated as described earlier.^[49]^ In brief, the AAV plasmid containing the CAG-Lifeact-HT7 expression cassette flanked by AAV2 packaging signals (ITRs) were co-transfected with plasmids pRV1 (AAV2 Rep and Cap sequences), pH21 (AAV1 Rep and Cap sequences) and pFD6 (Adenovirus helper plasmid) using Lipofectamine 3000 into HEK293 cells. 5 days post transfection, the cells were harvested and lysed using TNT extraction buffer (20 mM Tris pH = 7.5, 150 mM NaCl, 1% TX-100, 10 mM MgCl_2_) for 10 min. The cell debris was removed by centrifugation (3000 x g, 5 min, 4 °C), the cell supernatant was treated with Benzonase (4.5 µL, 30 min., 37 °C) and the rAAVs were purified *via* FPLC using AVB Sepharose columns. Purified rAAVs were concentrated using Amicon® Ultra 15 mL 100 kDa MWCO Centrifugal Filters (Merck KGaA, Darmstadt, Germany) and the buffer was exchanged to PBS pH 7.3.

U2OS Flp-IN T-REx™ cells stably expressing Vim-SNAP-tag2 were transduced with rAVVs 18 h prior to STED imaging by adding 0.5 μL purified rAAVs (∼ 10^9^-10^10^ rAAV particles) into 200 μL imaging medium (DMEM GlutaMAX™, 10% FCS, phenol-red free, Gibco).

### Labeling performance of SNAP-tag proteins determined with flowcytometry

Flow cytometry was conducted on a BD Fortessa™ X-20 Cell Analyzer (BD Biosciences) equipped with a highthroughput screening (HTS) module for 96-well plates. Cells were seeded on transparent 96-well cell culture plates and treated according to desired experiment in a reaction volume of 100 µL. After treatment, labeling reaction was stopped by addition of recombinant SNAP-tag2 (2 µM, 100 µL, 10 min. incubation), cells were washed twice with PBS (150 μL, 10 min. incubation), trypsinized (50 μL trypsin, 10 min. incubation) and resuspended in FACS buffer (2 % FBS in PBS) to a final volume of 200 μL. The cell suspension was transferred to non-binding u-bottom 96-well plates (Falcon) and analyzed on the flow cytometer using the HTS module. Flow rates were set to 3 μL/s with 3 mixing steps and subsequent washing volume of 400 μL. Cells were gated for live cells (SSC-A/FSC-A) and single cells (FSC-H/FSC-A) as exemplified in Supplementary Fig. 12. Excitation laser and emission filter settings were adjusted according to the fluorophore properties (Supplementary Table 8). Photomultiplier tubes (PMTs) settings were adjusted to positive and negative controls. Data was analyzed using FlowJo™ software (BD Biosciences).

### Substrate screening for SNAP-tag labeling in live mammalian cells

U2OS Flp In™ T-REx™ (Thermo Fisher) cells stably expressing mEGFP-SNAP-tag or mEGFP-CLIP-tag fusion proteins were seeded into 96-well cell culture plates (10’000 cells/well) the day prior to experiment. The cells were then incubated with substrates **1 – 30, CP or BC** [100 nM] for 2 h at 37 °C. All substrates were tested in technical triplicates. Cells were washed twice with cell growth medium for 15 min incubation at 37°°C and 1x with sterile PBS (pH = 7.4) prior to detachment with trypsin (50 µL, 10 min., 37 °C). Cells were resuspended in FACS buffer to a final volume of 200 µL. The cell suspension was transferred to non-binding u-bottom 96-well plates (Falcon) and analyzed on the flow cytometer using the HTS module, as previously described. All single cell events with a mEGFP signal in the FITC channel ≥1000 were analyzed for the labeling signal (TMR, PE channel) over expression level (mEGFP) and the medians of the ratios were derived using FlowJo™ software (BD Biosciences).

### Confocal fluorescence microscopy of SNAP-tag2 in live mammalian cells

Confocal fluorescence microscopy of U2OS cells stably expressing HaloTag7-P30-SNAP-tag2/SNAP_f_-tag-NLS-P2A-NLS-mTurquoise2 was conducted on a Stellaris 5 microscope as previously described. Live cell imaging was performed in either 8-well or 96-well glass bottom dishes (ibidi) at 37 °C and 5 % CO_2_ in a humidified chamber. Images were acquired either with a 20x (HC PL APO CS2 20x/0.75 IMM) or 40x (HC PL APO CS2 40x/1.10) water objective, a scan speed of 400 − 700 Hz, line average of 2, optical zoom of 1 − 1.3, image size 1024 × 1024 μm and either 12-bit or 16-bit pixel depth. Z-stacks were recorded with 2-5 μm step size. HaloTag7 and SNAP_f_-tag/SNAP-tag2 were labeled with their respective TF-/CA-fluorophore substrates at [100 nM] overnight. Cells were washed twice with PBS prior to imaging in Imaging medium. Laser settings are described in Supplementary Table 9. Image analysis was performed using FIJI software^[50]^. For visualization purposes, maximum intensity projections of z-stacked images were calculated. For quantification, intensities of multi-plane images were summed and region of interests (ROIs) were defined manually. Mean fluorescence intensities of ROI’s fluorescent channels were calculated (multi-ROI measurement) and ratios calculated from fluorescent labels to respective expression signals were compared. Ratiometric projections were conceived using the BRET-analyzer plug-in.^[51]^

### In cell kinetic measurements of SNAP-tag2 and SNAP_f_-tag

In cell kinetics of SNAP-tag2 and SNAP_f_-tag labeling with TF-, CF- and CP-fluorophore substrates were performed in U2OS cells stably expressing HaloTag7-P30-SNAP-tag2/SNAP_f_-tag-NLS-P2A-NLS-mTurquoise2 on a Stellaris 5 microscope as previously described. Fluorophore substrates were added at a final concentration of 50 nM (for TMR and CPY) or 100 nM (for SiR) to the cells and images with three z-stacks were acquired every 30 seconds over time. For quantification, fluorescence intensities of multiplane images were summed and the ratios of fluorophore substrate signals normalized to mTurquoise2 expression signals were analyzed using CellProfiler.^[52]^ Additional image acquisition parameters: 20x water objective (HC PL APO CS2 20x/0.75 IMM), 1.28 x optical zoom, image size 1024 × 1024 μm, scan speed: 400 Hz, line average: 2. Laser settings are described in Supplementary Table 10. Kinetic data was analyzed fitting a sigmoidal curve (equation (6)) in GraphPad Prism 10.2.3.

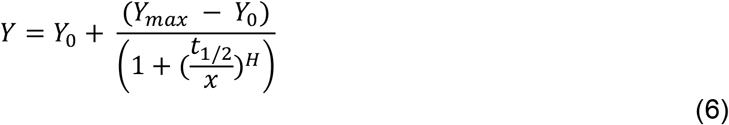

With Y = fluorescence intensity ratio (signal/mTurquoise2), Y_max_ = plateau of fluorescence intensity ratio, Y_0_ = y-intercept, t_1/2_ = half-labeling time [min.], H = Hill Slope and x = time [min.].

Experiment was conducted in biological duplicated and the average half labeling time (t_1/2_) of both replicates was calculated. Respective confidence intervals were propagated by equations (7-10) as followed:

1) Calculate mean t_1/2_ of both replicates:

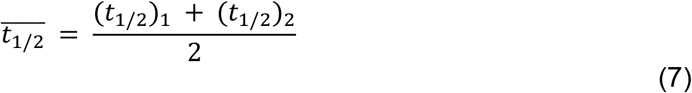

2) Computed the standard errors for each t_1/2_ of a replicate:

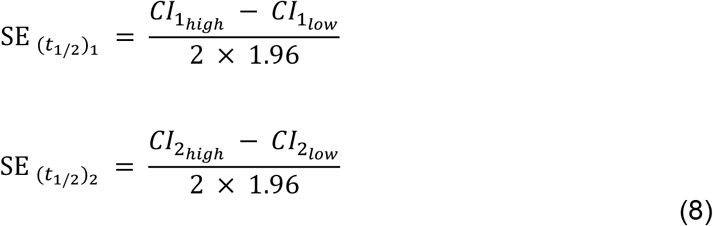

3) calculated the standard error of the mean

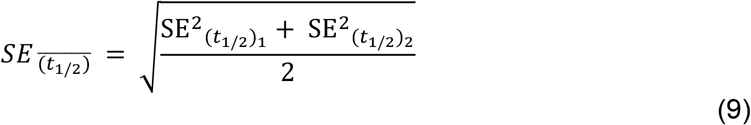

4) Compute the 95% confidence interval for the average t_1/2_

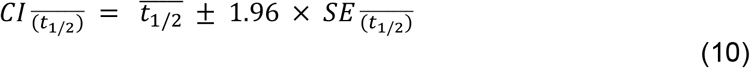

with 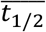= average half-labeling time of duplicate, CI = 95 % confidence interval, CI_high_ = higher 95 % confidence interval, CI_low_ = lower 95 % confidence interval, SE = standard error.

### Live-cell STED microscopy in mammalian cells

HeLa cells stably expressing Cox8a-SLP fusion proteins together with mEGFP were labeled with respective CA- or TF-SiR (100 nM) substrates for 1 h and washed twice with imaging medium prior to microscopy experiment. U2OS cells expressing Vim-SNAP_f_-tag/Vim-SNAP-tag2, TOMM20-SNAP-tag2, Sec61b-SNAP-tag2 or co-expressing Vim-SNAP-tag2 and LifeAct-HaloTag7 were labeled with CF-SiR and CA-MaP618 (100 nM), respectively, for 1 h and wished twice with imaging medium prior to microscopy experiment. Live-cell STED nanoscopy was performed using an Abberior STED Expert Line 595/775/RESOLFT QUAD scanning microscope (Abberior Instruments, Göttingen, Germany). SiR was excited using a 640 nm excitation line and depleted using a 775 nm STED line. For dual-color imaging, MaP618 was excited with a 561 nm excitation line and depleted with the same STED line. The microscope was equipped with a UPlanSApo 100x/1.4 oil immersion objective lens (Olympus, Tokyo, Japan) and the fluorescence signal was detected using avalanche photodiodes (APD). The pinhole was set to 0.9 Airy units, a gating of 0.75 – 8 ns was applied, dwell times of 7 – 10 µs and a pixel size of 25 nm were used. For STED images, each line was scanned 4 to 8 times and the signal was accumulated. Imaging settings are further detailed in Supplementary Table 11. Lookup tables used for images **a-d** in Figure 4: green (mEGFP), red-hot (SiR); for **e**: cyan (SNAP-tag2-SiR) and orange-hot (HaloTag7-MaP618).

The Full Width at Half Maximum (FWHM) of single intermediate filament fibers (Vimentin-SNAP_f_- tag/ SNAP-tag2) was determined by extracting fluorescence intensity profiles perpendicular to single filaments using the Fiji software. Mean filament diameters were calculated from 15 individual fibrils and from three individual images (n = 3) by fitting a gaussian function (equation (11)), yielding FWHM from equation (12).

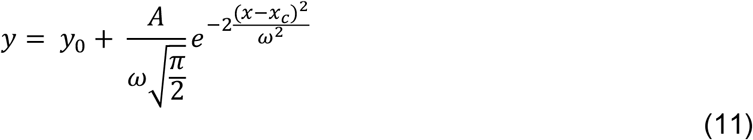

y: normalized fluorescence intensity, x: x-coordinate [μm], y_0_: offset, A: area, ω: width, x_c_: center.

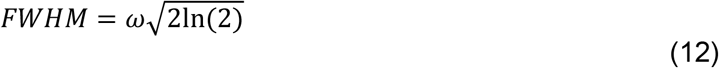

FWHM: Full Width at Half Maximum, ω: Gauss width.

### Strain and growth conditions of *Hansenula polymorpha* (*H. polymorpha*) yeast cells

*H. polymorpha* ku80 yeast cells were grown at 37 °C with shaking (200 rpm). For fluorescence microscopy studies, the cultures were grown in mineral medium (MM)^[53]^ containing 30 µg/ml leucine, 1x vitamin solution, 0.25% ammonium sulphate and 0.5% MeOH.

### Construction of *H. polymorpha* strains expressing Pex3-SNAP-tag2/SNAP_f_-tag

SNAP_f_-tag DNA sequence was synthesized by Genscript and SNAP-tag2 was PCR amplified from pET51b-SNAP-tag2 using primers Fw_BglII_SNAP-tag2 (GATGCTAGATCTGATAAA-GACTGCGAAATGAAACG) and Rv_SalI_SNAP-tag2 (CTCTAGAGTCGACCTAGCCCAGGCCTGGTTTACCC). Both SNAP-tag inserts were cloned into pUC57 plasmid vector. pHIPZ_Pex3_mGFP^[54]^ and pUC57_SNAP_f_-tag/SNAP-tag2 were restricted using SalI and BglII according to the supplier protocol (ThermoFisher Scientific) and ligated to generate pHIPZ_Pex3_SNAP_f_-tag or pHIPZ_Pex3_SNAP-tag2 plasmids for peroxisome targeting in yeast. Plasmids were transformed into *E. coli* DH5α cells selected on LB with Ampicillin. Positive colonies were checked using XhoI restriction. The plasmids were linearized with EcoRI or MluI respectively and transformed into *H. polymorpha* yku80 cells *via* electroporation as described in Faber et al.^[55]^ Positive integrations in yeast were tested with colony PCR using primers Ppex3_Fw (CCTGTT-GCGGCAAGATATAG) and Seq_SNAP_Rv (CGTAGGAAGGCTGGATGTC). Plasmids were confirmed by LightRun sequencing (Eurofins Genomics) and compared using Clone Manager 9 (Scientific and Educational Software). Correct yeast integration was checked *via* colony PCR with DreamTaq enzyme (Thermo Fisher Scientific) of zymolyase treated cells, and imaged using the Gel Doc XR+ System (Bio-Rad).

### Live-cell labeling of yeast peroxisomes

*H. polymorpha* cells were grown until stationary phase in MM supplemented with 0.5% glucose at 37 °C shaking (200 rpm). The culture was diluted to OD_660_ = 0.1 and grown until an OD_660_ >1.6. This culture was again diluted to OD_660_ = 0.1 in MM/MeOH and the cells were simultaneously stained using TF-, CP- or BG-SiR or -MaP555 at a final concentration of 250 nM for 18 h at 37 °C shaking, reaching a final OD_660_ of ∼3.0. To stain the yeast cell walls, cells were incubated with 25 µg/ml Calcofluor White (Fluorescent Brightner 28, Sigma-Aldrich Cat.No 910090) for 15 min at 37 °C shaking. Cells were washed 3x with PBS and immediately used for live-cell imaging in MM/MeOH medium.

### Live-cell STED microscopy of yeast peroxisomes

STED nanoscopy was conducted on a commercial microscope (Abberior Instruments GmbH) equipped with a STED depletion laser (775 nm), four excitation lasers (640, 561, 488 and 405 nm), a CoolLED pE-2 excitation system and a 100 × oil immersion objective (Olympus UPLSAPO/1.40). Confocal overview images (80 × 80 µm) were taken using a pixel size of 180 nm, dwell time of 20 µs, 6% laser power using the 561 nm excitation laser with 570 – 680 nm detection for MaP555 substrates, or 0.5% laser power using the 640nm excitation laser with 650 – 730 nm detection for SiR substrates. STED images were taken with a pixel size of 25nm, dwell time of 20 µs, 30% laser power using the 561 nm excitation laser with 570 – 680 nm detection for MaP555 substrates, or 2% laser power using the 640 nm excitation laser with 650 – 730 nm detection for SiR substrates. A depletion laser power of 50% or 10% at 775 nm was applied for 561 or 640 nm excitation respectively. Calcofluor White was imaged using 3% laser power with the 405 nm excitation laser and 410 – 480 nm detection. Image acquisition was carried out using Imspector software (v16.3, Abberior Instruments). Analysis was performed using Fiji software (ImageJ 2.14.014). To compare the fluorescence intensities of SNAP_f_-tag and SNAP-tag2 labeled with different fluorophore substrates, the mean and maximum fluorescent intensity of 3 × 125 cells (rectangles of 3 × 3 µm as ROI) were measured in biological triplicates.

## Acknowledgements

This work was supported by the Max Planck Society, the École Polytechnique Fédérale de Lausanne (EPFL) and the Deutsche Forschungsgemeinschaft (DFG, German Research Foundation), TRR 186. S.K., J.W. and J.K. acknowledge support from the Heidelberg Biosciences International Graduate School (HBIGS). J.W. was supported by the Max Planck School Matter to Life. Y.-H.L. was supported by EPFL Doctoral Program in Biotechnology and Bioengineering (EDBB). The authors thank A. Bergner and B. Réssy for providing reagents and materials. We thank Dr. B. Koch, J. Kress and P. Breuer for help with stable cell line generation. We acknowledge the mass spectrometry facility (S. Fabritz, T. Rudi and J. Kling) for its support. The authors thank all members of the Johnsson lab for critical discussions.

## Author Contributions

S.K., V.N., J.H., and K.J. designed and interpreted the experiments. V.N., A.L. and S.K. designed and synthesized the compounds used in this paper. S.K., J.W., Y-H.L., C.E. and J.H. performed protein engineering. V.N. and J.F. performed substrate screening and measured their photophysical properties. M.T. performed NanoDSF measurements. S.K. and J.R. designed, performed and analyzed *in vitro* kinetic studies. S.K. performed *in vitro* characterization, confocal microscopy and flowcytometry experiments of mammalian cells. J.K. performed STED imaging in mammalian cells. E.M.F.d.L and R.V. designed and performed experiments in yeast. S.K. and K.J. wrote the manuscript with input from all authors.

