## Supplementary Information for "SNAP-tag2: faster and brighter protein labeling"

for

#### For Table of Contents use only


**page**

##### Supplementary Figures

**Supplementary Fig. 1**: Chemical structures of SLP substrates used in this study. 4
**Supplementary Fig. 2:** Comparison of SNAP$_{(f)}$-tag labeling with novel TF-TMR substrate (**30**) relative to labeling with CP-TMR regarding their labeling kinetics *in vitro* and labeling performance in live cells. 5
**Supplementary Fig. 3:** Schematic overview of SNAP-tag engineering process. 5
**Supplementary Fig. 4:** Thermal stability measurement of SNAP-tag2 and SNAP$_f$-tag *via* NanoDSF. 6
**Supplementary Fig. 5:** Comparison of SNAP-tag2 und SNAP$_f$-tag for labeling with different fluorescent substrates in live cells. 6
**Supplementary Fig. 6:** Comparison of HaloTag7, SNAP-tag2 and SNAP$_f$-tag performances in confocal fluorescence microscopy. 7
**Supplementary Fig. 7:** SNAP$_f$-tag performance in CLSM and STED microscopy. 7
**Supplementary Fig. 8**: SNAP-tag2 and SNAP$_f$-tag labeling of live yeast peroxisomes. 8
**Supplementary Fig. 9:** Comparison of SNAP-tag2 and SNAP$_f$-tag performance with different substrates in STED imaging of live *H. polymorpha* yeast peroxisomes. 9
**Supplementary Fig. 10:** CLSM images of labeled yeast peroxisomes used for quantification of fluorescence intensities of SNAP-tag2 and SNAP$_f$-tag labeled with different SiR and MaP555 substrates. 10
**Supplementary Fig. 11:** Gating strategies for YSD library screening *via* FACS. 11
**Supplementary Fig. 12:** Exemplified gating strategy for determination of SNAP variant labeling performance in cells *via* flow cytometry. 11

##### Supplementary Tables

**Supplementary Table 1:** Substrate screening for reaction kinetics *in vitro* and in cell labeling of SLPs with new substrates. 12
**Supplementary Table 2:** Kinetical parameters of independent experimental replicates for SNAP-tag2 labeling with different TMR and CPY substrates analyzed by model 2. 13
**Supplementary Table 3:** Average of kinetical parameters for SNAP-tag2 labeling with different TMR and CPY substrates calculated from data in Supplementary Table 2. 13
**Supplementary Table 4:** Kinetic parameters of SNAP-tag2 labeling with non-fluorescent substrates. 14
**Supplementary Table 5:** Comparison of extinction coefficients (ε) and quantum yields (QY) for fluorescently labeled SNAP$_f$-tag and SNAP-tag2. 14

##### Supplementary Methods

**Supplementary Table 6:** Settings used for YSD library screening *via* FACS. 15
**Supplementary Table 7:** Filter settings used in FP measurements. 15
**Supplementary Table 8:** Settings used for measuring the labeling performance of SNAP-tag proteins in live mammalian cells *via* flowcytometry. 15
**Supplementary Table 9:** Settings used for confocal fluorescence imaging of SLPs in live mammalian cells 15
**Supplementary Table 10:** Microscopy settings for measuring labeling kinetics of SNAP-tag2 and SNAP$_f$-tag in live mammalian cells. 15
**Supplementary Table 11:** Settings used for CLSM and STED microscopy of SLPs in live mammalian cells. 16
**Supplementary Table 12:** Buffers, media and reagents used. 17
**Supplementary Table 13:** Most important plasmids and generated stable mammalian cell lines used. 18
**Supplementary Table 14:** Primers used for generation of library 1 on SNAP-tag1.1 *via* one-pot saturation mutagenesis. 19
**Supplementary Table 15:** Primers used for generation of library 2 on SNAP-tag1.1 *via* one-pot saturation mutagenesis. 19
**Supplementary Table 16:** Primers used for generation of library 3 *via* site-directed saturation mutagenesis. 19
**Supplementary Table 17:** Primers used for generation of site-directed saturation mutagenesis libraries 4-9 *via* assembly PCR. 20
**Supplementary Table 18:** NGS primers used for amplification of libraries after FACS screen. 20

**Protein Sequences** 21-24

**Scripts**
- Rosetta Scripts 25-27
- NGS scripts 28
- Example DynaFit Scripts 29-31

**Chemical synthesis of SNAP-tag2 substrates**
- General remarks 32
- Building blocks – nucleobases 33-36
- Building blocks – benzyl alcohol 37-39
- Compound precursors 40-67
- Fluorophore compounds – TMR substrates 68-78
- Fluorophore compounds – other fluorophores 79-81
- Non-fluorescent compounds 82-83

**NMR spectra** 84-158

**References** 159

#### Supplementary Figures

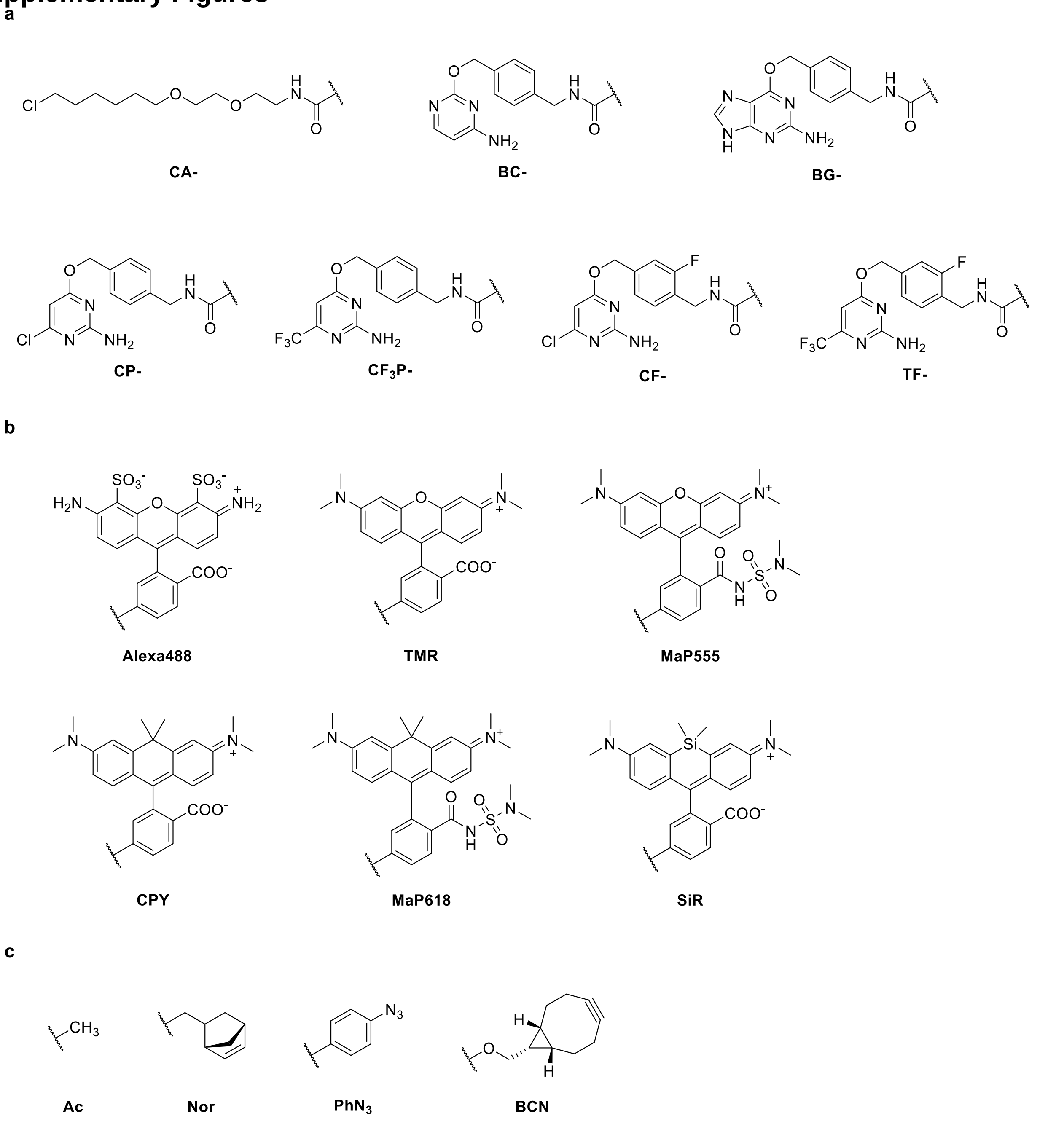


**Supplementary Fig. 1**: Chemical structures of SLP substrates used in this study. **a**, Chemical structures of HaloTag7 (CA), CLIP-tag (BC) and SNAP-tag (BG, CP, $CF_3P$, CF and TF) core substrates. **b**, Chemical structures of fluorescent substituents. **c**, Chemical structures of non-fluorescent substituents.

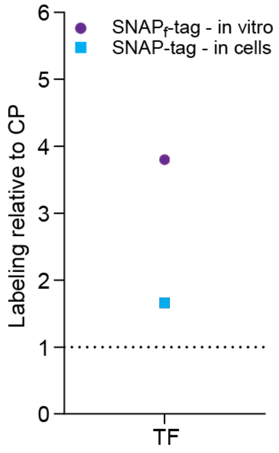


**Supplementary Fig. 2:** Comparison of SNAP$_{(f)}$-tag labeling with novel TF-TMR substrate (**30**) relative to labeling with CP-TMR regarding their labeling kinetics *in vitro* and labeling performance in live cells. *In vitro* labeling kinetics were measured recording fluorescence polarization (FP) traces over time using TF-TMR at [20 nM] and SNAP$_f$-tag protein at [50 nM]. A one-phase association was fitted to the data, apparent second-rate constants ($k_{app}$) were calculated ($k_{app}$ = k/[protein]) and normalized to $k_{app}$ of SNAP$_f$-tag labeling with CP-TMR. In-cell characterization was conducted using U2OS cells stably expressing a mEGFP-SNAP-tag fusion protein. Cells were labeled with TF-TMR at [100 nM] for 2 h, washed and analyzed *via* flow cytometry. Ratios of TMR/mEGFP were calculated and normalized to the ratio obtained for SNAP-tag with CP-TMR.

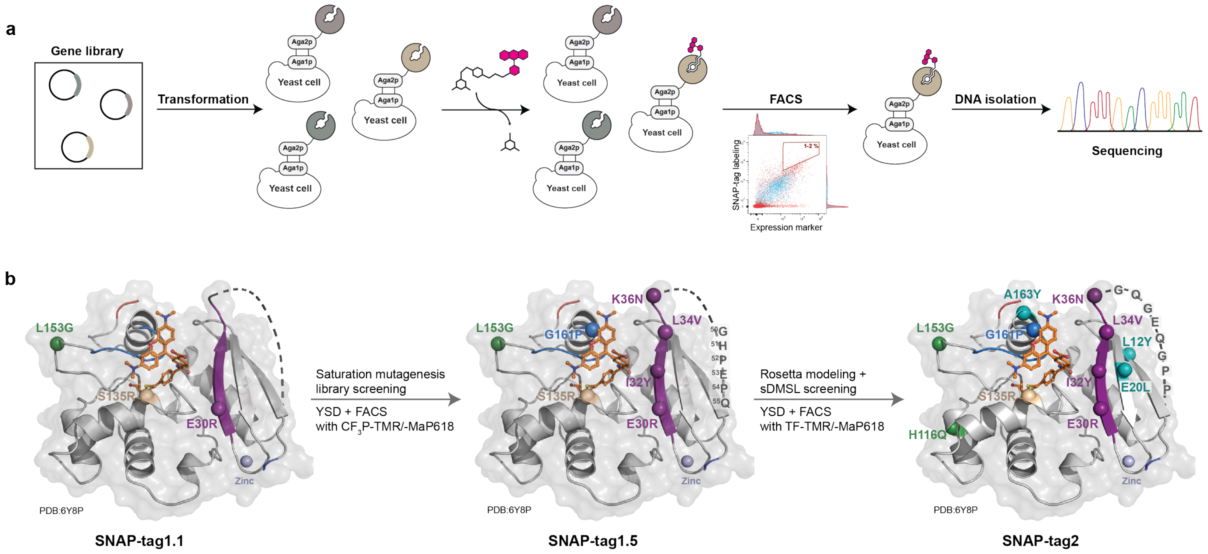


**Supplementary Fig. 3:** Schematic overview of SNAP-tag engineering process. **a**, Schematic overview of directed evolution process coupled to YSD. Gene libraries were generated and transformed into EBY100 yeast cells for surface display. Yeast cells were incubated with new SNAP-tag substrates and fluorescent antibodies or bilirubin (eUnaG2 substrate) for expression control. Yeast cells were sorted for double positive labeling signal. DNA was isolated and subsequently analyzed by Sanger Sequencing of single clones or NGS of the library pool. This process was repeated for several rounds increasing the stringency in each round. **b**, Crystal structures of TMR-labeled SNAP (PDB:6Y8P) highlighting mutations of intermediate SNAP-tag versions used for library generation. SNAP-tag1.1 was used for library generation on the active site loop (res. 155-161) and on the proximal β-strand to the active site (res. 29-36). Libraries were screened with $CF_3$P-TMR and -MaP618 and combination of hit mutations resulted in SNAP-tag1.5. SNAP-tag1.5 was used for creation of a custom-made synthetic deep mutational scanning library (sDMSL) and for Rosetta calculations on the unresolved region (res.37-54). The sDMSL library was screened with TF-TMR and -MaP618 substrates. Combination of hit mutations and implementation of the Rosetta-modeled 8 amino acid loop resulted in the final SNAP-tag2 protein. Colors represent the origin of different mutations: Forest green: PROSS-predicted; Wheat: rational design, deep-purple and marine blue: saturation mutagenesis screens; gray: mutations in the unresolved loop (for SNAP-tag1.5 serendipitously found during screening of β-strand library, for SNAP-tag2 Rosetta modeled), Teal: sDMSL screen. Termini are highlighted in blue (N-terminus) and red (C-terminus) and the coordinated zinc ion is illustrated as light-blue sphere.

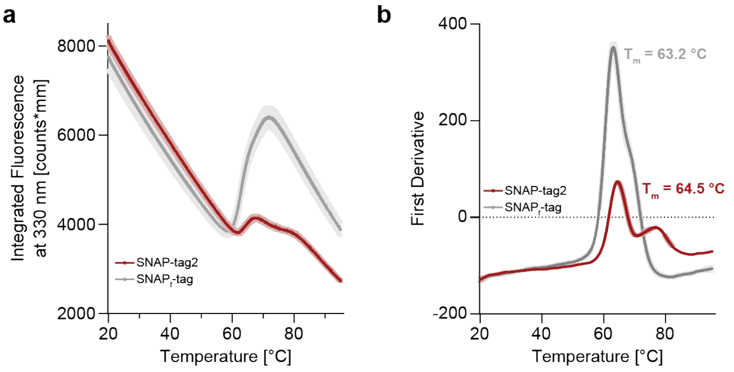


**Supplementary Fig. 4:** Thermal stability measurement of SNAP-tag2 and SNAPf-tag *via* NanoDSF. NanoDSF thermograms representing **a**, the integrated fluorescence at 330 nm with increasing temperature and **b** the corresponding first derivative of the integrated fluorescence at 330 nm for SNAP-tag2 and SNAPf-tag. SNAP-tag2 has a melting temperature ($T_m$) of ~ 65 °C, which is slightly increased compared to SNAPf-tag ($T_m$ ≈ 63 °C).

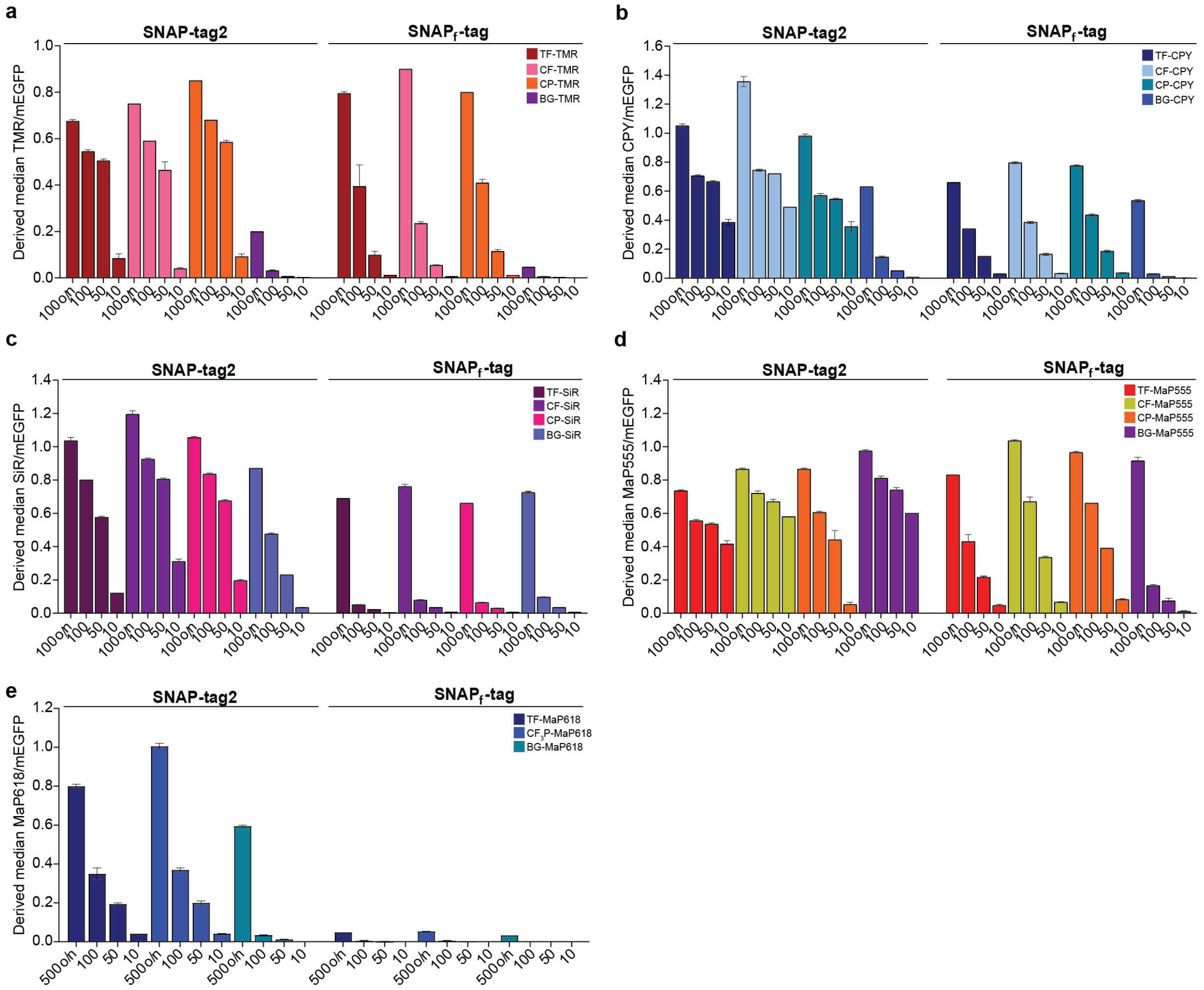


**Supplementary Fig. 5:** Comparison of SNAP-tag2 und SNAPf-tag for labeling with different fluorescent substrates in live cells. U2OS cell stably expressing mEGFP-SNAP-tag2 or mEGFP-SNAPf-tag were incubated with different **a** TMR, **b** CPY, **c** SiR, **d** MaP555 or **e** MaP618 substrates at [100 nM] or [500 nM] overnight or at [100 nM, 50 nM and 10 nM] for 1 h. Cells were analyzed by flow cytometry. Analysis was done using FlowJo by gating for single cells and double positive labeling signal (as depicted in Supplementary Fig. 12) and the median of the fluorescent label/mEGFP was derived. SNAP-tag2 showed for most of the substrates a higher labeling ratio.

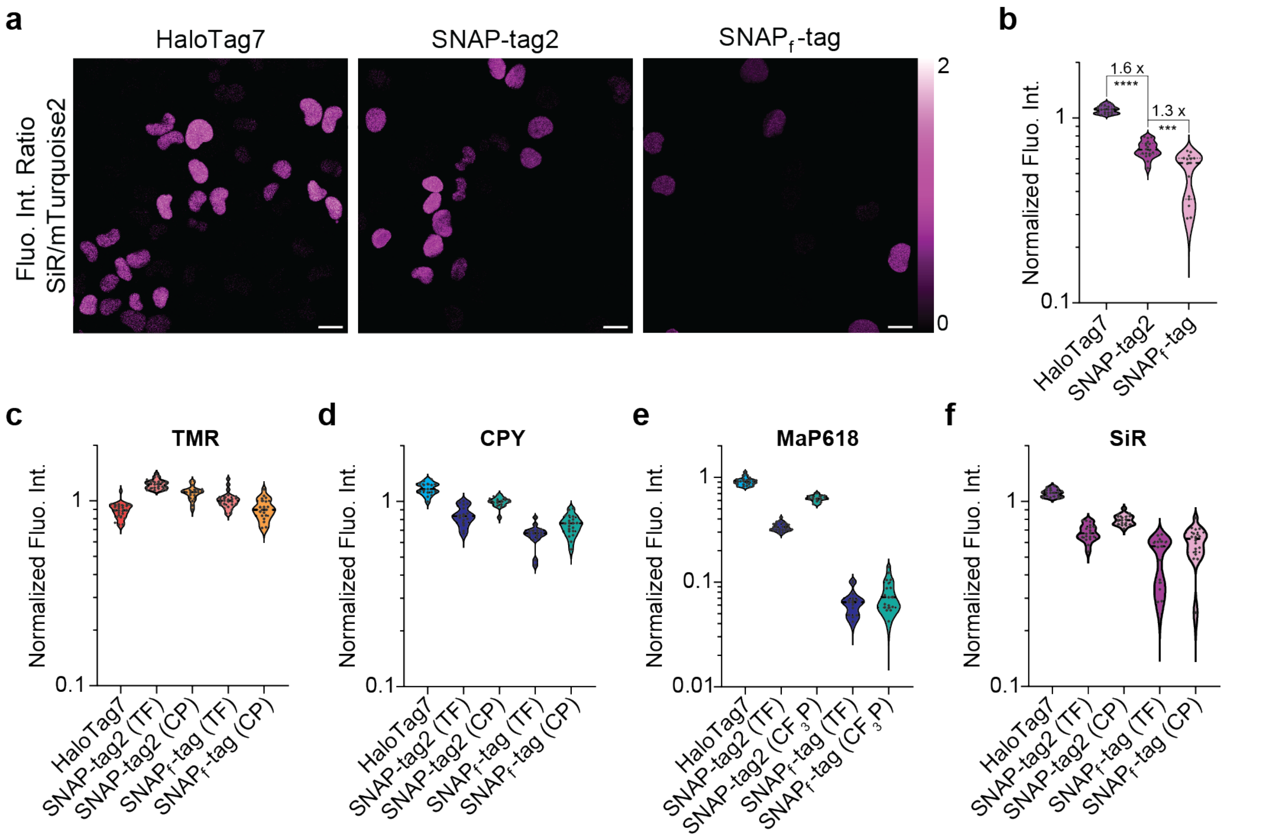


**Supplementary Fig. 6:** Comparison of HaloTag7, SNAP-tag2 and SNAP$_f$-tag performances in confocal fluorescence microscopy. Experiments were conducted in live U2OS cells stably co-expressing HaloTag7-SNAP$_f$-tag or HaloTag7-SNAP-tag2 together with mTurquoise2 (expression marker) in the nucleus. **a**, Comparison of HaloTag7, SNAP-tag2 and SNAPf-tag performances in confocal fluorescence microscopy with SiR substrates. U2OS cells were labeled with SLP-respective CA-/TF-SiR substrates at [100 nM] overnight. Ratiometric projections are presented corresponding to SiR label/mTurquoise2 on a magenta-hot look-up table. Scale bar: 20 µm. **b**, Violin plots representing the quantitative analysis of single cells shown in a. Numbers represent fold-changes between the different SLPs (n = 15 - 20 cells, Welch t-test: **** ≙ $p < 0.0001$, *** ≙ $p = 0.0003$). **c-f,** Violin plots representing the quantitative analysis of different fluorescent substrates/mTurquoise2 for **c** TMR, **d** CPY, **e**, MaP618 and **f** SiR (n = 13 - 25 cells).

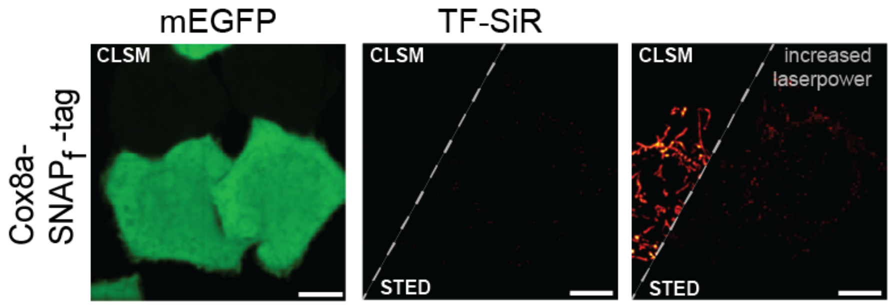


**Supplementary Fig. 7:** SNAP$_f$-tag performance in CLSM and STED microscopy. HeLa cells stably co-expressing SNAP$_f$-tag in the mitochondria (Cox8a localization sequence) together with mEGFP (no specific localization) were labeled with TF-SiR (100 nM) for 1 h. SNAP$_f$-tag showed insufficient signal under the same imaging conditions used for SNAP-tag2 and HaloTag7 and required the use of an increase laser power to see labeling signal. Scale bars: 10 µm. LUTs: green (mEGFP), red-hot (SiR).

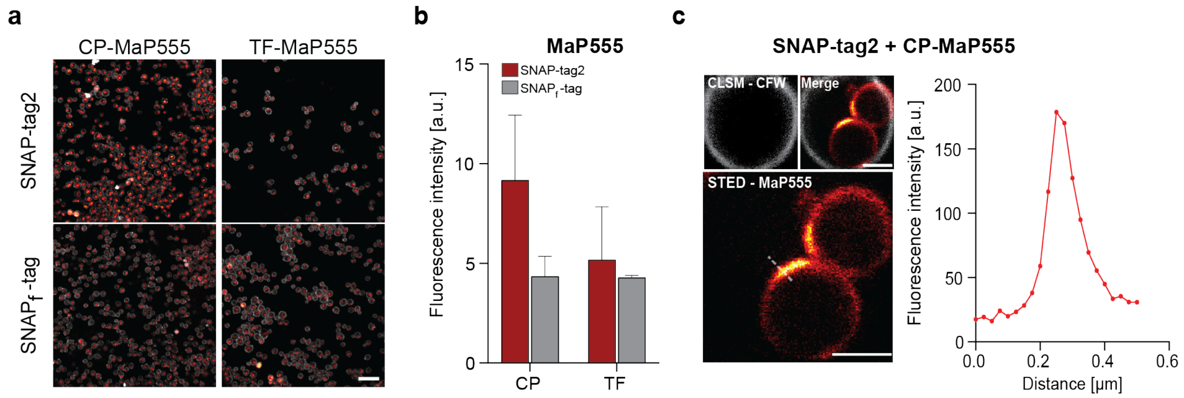


**Supplementary Fig. 8**: SNAP-tag2 and SNAP$_f$-tag labeling of live yeast peroxisomes. **a**, CLSM images of *H. polymorpha* yeast cells expressing Pex3-SNAP-tag2 or Pex3-SNAP$_f$-tag fusion proteins labeled with different MaP555 substrates. Yeast cells were labeled with CP- or TF-MaP555 (250 nM) for 18 h and the cell wall was stained with Calcofluor White (CFW; 25 µg/mL) for 15 min. Scale bar: 10 µm. **b**, Bar plot representing the quantitative analysis of SNAP-tag2 and SNAP$_f$-tag labeling with MaP555 substrates in yeast. Experiments were conducted in biological triplicates and the mean fluorescence intensity of MaP555 substrates was calculated for 3 x 125 cells (Supplementary Fig. 10). Combination of SNAP-tag2 with CP-MaP555 show the best results for labeling of yeast peroxisomes. **c,** STED image of Pex3-SNAP-tag2 labeled with CP-MaP555 (lower panel), CLSM image of CFW stained cell wall and merge of both channels (upper panels). LUTs: red-hot (MaP555) and gray (CFW). Scale bar: 1 µm. The right plot shows the line profile of labeled peroxisomes in STED (highlighted as white dashed line in the image). SNAP-tag2 with CP-MaP555 is well suited to perform live cell STED in yeast.

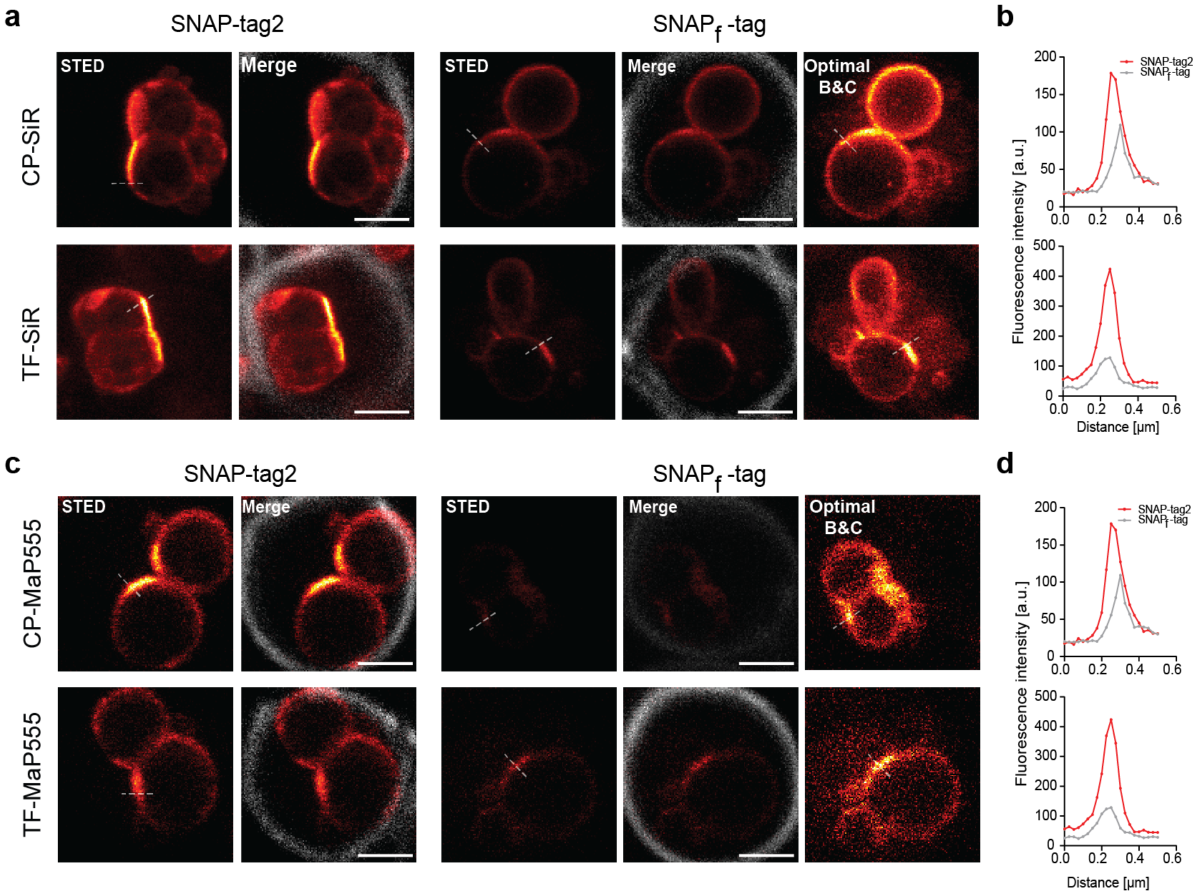


**Supplementary Fig. 9:** Comparison of SNAP-tag2 and SNAP$_f$-tag performance with different substrates in STED imaging of live *H. polymorpha* yeast peroxisomes. **a**, STED images of Pex3-SNAP-tag2 and Pex3-SNAPf-tag labeled with CP- or TF-SiR (left panel) and merged images with CLSM image of Calcofluor White stained cell wall (right panel). Due to the low fluorescence signal for SNAP$_f$-tag under the same experimental conditions as SNAP-tag2, a third panel showing the STED image adjusted to optimal brightness and contrast (B & C) settings was added to show labeling of SNAP$_f$-tag. LUTs: red-hot (SiR) and gray (Calcofluor White). Scale bar: 1 µm. **b**, Plots representing the line profiles of SiR-labeled peroxisomes in STED imaging of cells shown in a (highlighted as dashed lines in images) for SNAP-tag2 and SNAP$_f$-tag. **c,** STED images Pex3-SNAP-tag2 and Pex3-SNAPf-tag labeled with CP- or TF-MaP555 (left panel) and merged images with CLSM image of Calcofluor White stained cell wall (right panel) similar to images shown in a. **d** Plots representing the line profiles of MaP555-labeled peroxisomes in STED imaging of cells shown in c (highlighted as dashed lines in images) for SNAP-tag2 and SNAP$_f$-tag. SNAP-tag2 shows a sharper line profile with a higher fluorescence intensity compared to SNAP$_f$-tag for both SiR and MaP555 substrates.

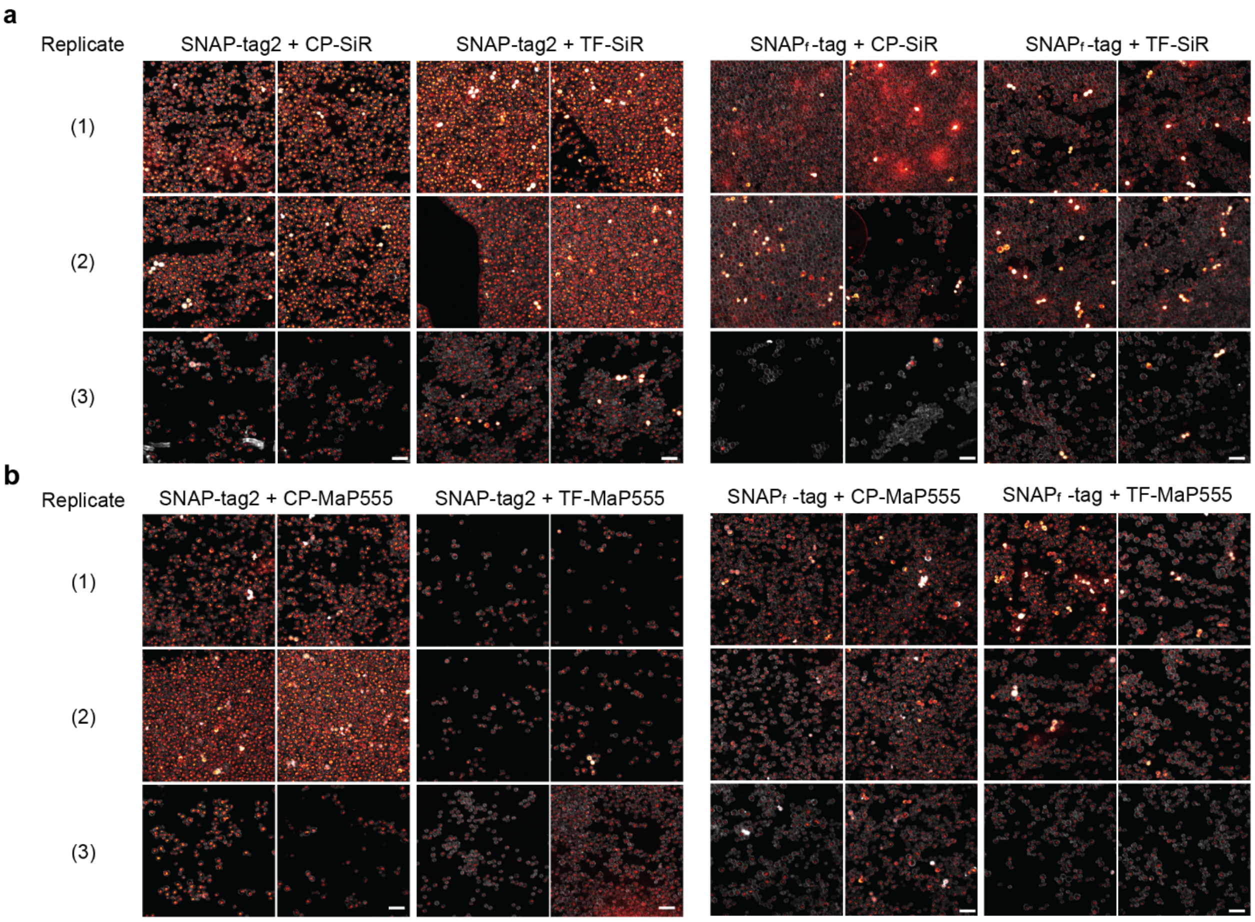


**Supplementary Fig. 10:** CLSM images of labeled yeast peroxisomes used for quantification of fluorescence intensities of SNAP-tag2 and SNAP$_f$-tag labeled with different **a** SiR and **b** MaP555 substrates (Fig. 5b and Supplementary Fig. 9b). Experiments were conducted in biological triplicates and the fluorescence intensities of labeled peroxisomes of 3 x 125 cells from two field of views were analyzed.

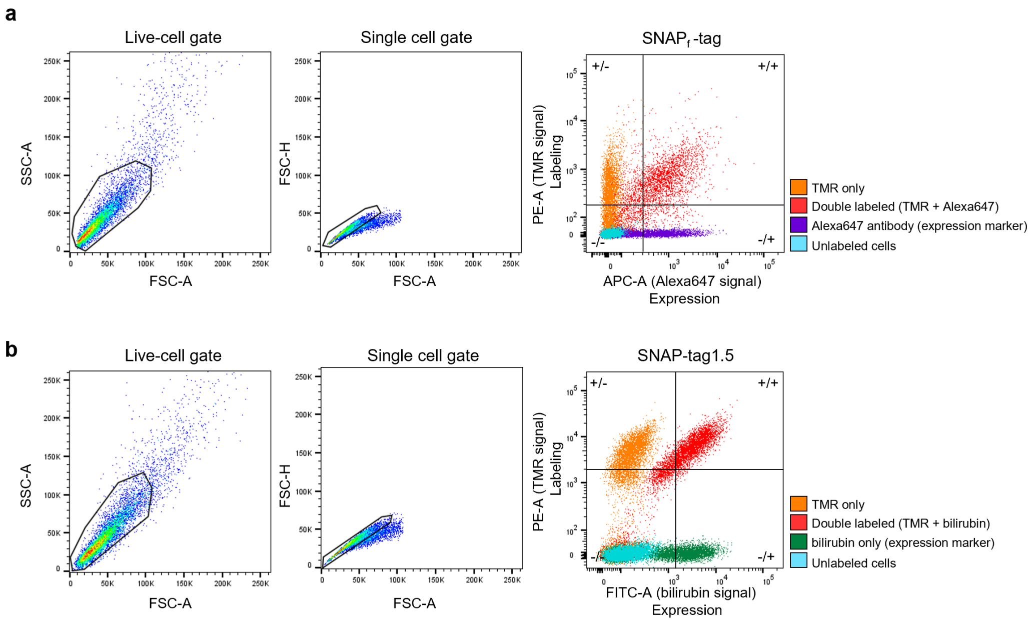


**Supplementary Fig. 11:** Gating strategies for YSD library screening *via* FACS. Hierarchical gating (left to right) of live cells (SSC-A/FSC-A), single cells (FSC-H/FSC-A) and SNAP labeling (label/expression). **a,** Gating strategy for libraries 1-3 using the pCTcon2 expression vector for YSD. Gating strategy for labeling is here exemplified for the SNAP$_f$-tag control labeled with TMR. Expression levels were marked by immunostaining of cmyc-tag with Alexa647 antibody (x-axis; APC-A) and plotted against the TMR labeling signal (y-axis; PE-A). Control samples of yeast cells displaying SNAP$_f$-tag were used for setting the gates based on labeling with TMR only (+/-; orange population), Alexa647 only (-/+; purple population), double labeled (+/+, red population) or unlabeled (-/-, blue population). The top 0.5-3 % of the double labeled population (+/+) were sorted based on the condition and selection round. **b**, Gating strategy for libraries 4-9 using the pJYDNg expression vector for YSD. Gating strategy for labeling is here exemplified for the SNAP-tag1.5 control labeled with TMR. Expression levels were reported by eUnaG2 labeling with bilirubin (x-axis; FITC-A) and plotted against the TMR signal (y-axis; PE-A). Control samples of yeast cells displaying SNAP1.5 were used for setting the gates based on labeling with TMR only (+/-; orange population), bilirubin only (-/+; green population), double labeled (+/+, red population) or unlabeled (-/-, blue population). The top 0.5 - 1 % of the double labeled population (+/+) were sorted based on the condition and selection round.

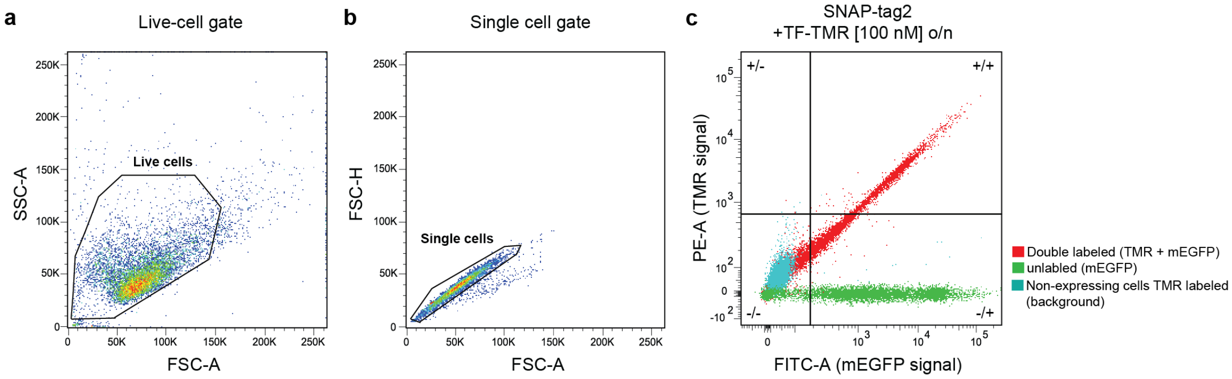


**Supplementary Fig. 12:** Exemplified gating strategy for determination of SNAP variant labeling performance in cells *via* flow cytometry. **a**, Gating for live cell events (SSC-A/FSC-A). **b**, Gating for single cell events (FSC-H/FSC-A). **c**, Gating for double positive labeled cells (+/+) exemplified for SNAP-tag2 labeling with TF-TMR [100 nM] overnight. Gates were adjusted according to control samples to eliminate background labeling.

#### Supplementary Tables

**Supplementary Table 1:** Substrate screening for reaction kinetics *in vitro* and in cell labeling of SLPs with new substrates. $k_{app}$ = apparent second-order rate constant; n.d. = not determined/not reactive; s.d. = standard deviation; Fluo. Int. = fluorescence intensity.

| Compound | | SNAP$_f$-tag | SNAP-tag | CLIP$_f$-tag | CLIP-tag | hAGT |
|---|---|---|---|---|---|---|
| | | *In vitro* | In cells | *In vitro* | In cells | *In vitro* |
| | | **$k_{app}$ (± s.d.) × $10^4$ [$M^{-1}$ $s^{-1}$]** | **Fluo. Int. Ratio TMR/GFP** | **$k_{app}$ (± s.d.) × $10^4$ [$M^{-1}$ $s^{-1}$]** | **Fluo. Int. Ratio TMR/GFP** | **$k_{app}$ (± s.d.) × $10^3$ [$M^{-1}$ $s^{-1}$]** |
| **Leaving group modifications** | **BC-TMR** | n.d. | 0.003 ± 0.001 | 1.13 ± 0.02 | 0.038 ± 0.001 | n.d. |
| | **BG-TMR** | 34.9 ± 3.43 | 0.005 ± 0.001 | 0.02 ± 0.01 | 0.001 ± 0.001 | 0.97 ± 0.02 |
| | **CP-TMR** | 6.17 ± 0.50 | 0.898 ± 0.024 | n.d. | 0.003 ± 0.001 | 0.80 ± 0.04 |
| | **1** | 0.52 ± 0.03 | 0.194 ± 0.005 | 0.01 ± 0.01 | 0.005 ± 0.001 | 0.16 ± 0.01 |
| | **2** | 0.91 ± 0.06 | 0.385 ± 0.005 | n.d. | 0.002 ± 0.001 | 0.21 ± 0.01 |
| | **3** | 10.1 ± 0.40 | 1.268 ± 0.025 | n.d. | 0.007 ± 0.001 | 1.40 ± 0.04 |
| | **4 ($CF_3P$)** | 18.9 ± 2.57 | 1.380 ± 0.019 | n.d. | 0.003 ± 0.001 | 0.64 ± 0.04 |
| | **5** | 0.88 ± 0.01 | 0.287 ± 0.007 | 0.01 ± 0.01 | 0.007 ± 0.001 | 0.18 ± 0.01 |
| | **6** | 0.62 ± 0.03 | 0.389 ± 0.009 | n.d. | 0.002 ± 0.001 | 0.15 ± 0.01 |
| | **7** | 0.27 ± 0.02 | 0.080 ± 0.007 | n.d. | 0.003 ± 0.001 | 0.08 ± 0.01 |
| | **8** | 3.89 ± 0.19 | 1.186 ± 0.031 | n.d. | 0.003 ± 0.001 | 0.31 ± 0.01 |
| | **9** | 0.30 ± 0.03 | 0.008 ± 0.001 | 0.44 ± 0.02 | 0.011 ± 0.001 | 0.07 ± 0.01 |
| | **10** | n.d. | 0.008 ± 0.001 | n.d. | 0.001 ± 0.001 | 0.04 ± 0.01 |
| | **11** | 1.61 ± 0.02 | 0.054 ± 0.002 | n.d. | 0.001 ± 0.001 | 0.48 ± 0.01 |
| | **12** | n.d. | 0.002 ± 0.001 | n.d. | n.d. | n.d. |
| | **13** | n.d. | 0.007 ± 0.001 | 0.02 ± 0.01 | 0.002 ± 0.001 | 0.06 ± 0.01 |
| | **14** | n.d. | 0.001 ± 0.001 | n.d. | n.d. | n.d. |
| | **15** | 2.33 ± 0.16 | 0.055 ± 0.001 | 0.02 ± 0.01 | 0.005 ± 0.001 | 0.56 ± 0.16 |
| | **16** | 5.55 ± 0.39 | 0.107 ± 0.007 | 0.06 ± 0.01 | 0.009 ± 0.001 | 2.11 ± 0.03 |
| | **17** | 4.28 ± 0.73 | 0.973 ± 0.140 | n.d. | 0.004 ± 0.001 | 2.30 ± 0.07 |
| **Linker modifications** | **18** | 0.06 ± 0.03 | 0.044 ± 0.001 | n.d. | 0.002 ± 0.001 | 0.05 ± 0.01 |
| | **19** | n.d. | 0.013 ± 0.001 | n.d. | 0.007 ± 0.001 | 0.10 ± 0.01 |
| | **20** | n.d. | 0.008 ± 0.001 | n.d. | 0.005 ± 0.001 | 0.04 ± 0.02 |
| | **21** | n.d. | 0.009 ± 0.001 | n.d. | 0.004 ± 0.001 | 0.12 ± 0.01 |
| | **22 (CF)** | 6.06 ± 1.14 | 1.514 ± 0.017 | n.d. | 0.007 ± 0.001 | 0.55 ± 0.05 |
| | **23** | 7.70 ± 1.49 | 1.721 ± 0.028 | n.d. | 0.017 ± 0.001 | 2.61 ± 0.06 |
| | **24** | 8.68 ± 1.02 | 1.356 ± 0.037 | n.d. | 0.013 ± 0.001 | 1.45 ± 0.11 |
| | **25** | 2.68 ± 0.19 | 0.646 ± 0.034 | n.d. | 0.005 ± 0.001 | 0.46 ± 0.03 |
| | **26** | 2.37 ± 0.46 | 0.340 ± 0.006 | n.d. | 0.001 ± 0.001 | 0.78 ± 0.04 |
| | **27** | n.d. | 0.002 ± 0.001 | n.d. | n.d. | n.d. |
| | **28** | n.d. | 0.002 ± 0.001 | n.d. | n.d. | n.d. |
| | **29** | n.d. | 0.058 ± 0.002 | n.d. | 0.003 ± 0.001 | n.d. |
| | **30 (TF)** | 23.5 ± 4.87 | 1.491 ± 0.109 | n.d. | 0.010 ± 0.001 | 0.52 ± 0.06 |

**Supplementary Table 2** Kinetical parameters **of** independent experimental replicates for SNAP-tag2 labeling with different TMR and CPY substrates analyzed by model 2.

| **Substrate** | **$k_1$ (± s.d.) [$M^{-1}$ $s^{-1}$]** | **$k_{-1}$ (± s.d.) [$s^{-1}$]** | **$k_2$ (± s.d.) [$s^{-1}$]** | **$k_{app}$ (± s.d.) [$M^{-1}$ $s^{-1}$]** | **$K_d$ (± s.d.) [nM]** |
|---|---|---|---|---|---|
| **TF-TMR** | 1.34 (± 0.01) × $10^7$ | 0.57 (± 0.03) | 0.71 (± 0.05) | 7.45 (± 0.16) × $10^6$ | 42.2 (± 0.18) |
| | 1.78 (± 0.01) × $10^7$ | 0.60 (± 0.02) | 0.43 (± 0.02) | 7.42 (± 0.17) × $10^6$ | 33.9 (± 0.01) |
| | 1.31 (± 0.01) × $10^7$ | 0.43 (± 0.05) | 0.97 (± 0.13) | 9.08 (± 0.33) × $10^6$ | 32.6 (± 0.36) |
| | 1.60 (± 0.01) × $10^7$ | 0.31 (± 0.03) | 0.39 (± 0.07) | 8.93 (± 0.54) × $10^6$ | 19.1 (± 0.20) |
| **CF-TMR** | 1.90 (± 0.03) × $10^7$ | 1.88 (± 0.07) | 0.77 (± 0.03) | 5.51 (± 0.22) × $10^6$ | 99.2 (± 0.33) |
| | 1.91 (± 0.03) × $10^7$ | 0.71 (± 0.05) | 0.96 (± 0.08) | 1.09 (± 0.03) × $10^7$ | 37.2 (± 0.24) |
| | 1.91 (± 0.03) × $10^7$ | 5.62 (± 0.18) | 0.56 (± 0.02) | 1.72 (± 0.01) × $10^6$ | 294 (± 0.10) |
| | 1.80 (± 0.02) × $10^7$ | 4.75 (± 0.14) | 0.57 (± 0.02) | 1.95 (± 0.10) × $10^6$ | 263 (± 0.08) |
| | 2.29 (± 0.03) × $10^7$ | 5.07 (± 0.16) | 0.52 (± 0.02) | 2.13 (± 0.12) × $10^6$ | 222 (± 0.08) |
| | 2.59 (± 0.03) × $10^7$ | 3.60 (± 0.14) | 1.86 (± 0.10) | 8.85 (± 0.33) × $10^6$ | 139 (± 0.04) |
| | 1.89 (± 0.03) × $10^7$ | 1.36 (± 0.11) | 1.03 (± 0.11) | 8.15 (± 0.58) × $10^6$ | 71.6 (± 0.06) |
| | 2.42 (± 0.03) × $10^7$ | 2.08 (± 0.09) | 1.57 (± 0.09) | 1.04 (± 0.04) × $10^7$ | 85.6 (± 0.03) |
| **CP-TMR** | 2.03 (± 0.03) × $10^7$ | 1.48 (± 0.08) | 1.80 (± 0.09) | 1.11 (± 0.02) × $10^7$ | 73.2 (± 0.32) |
| | 2.13 (± 0.02) × $10^7$ | 1.56 (± 0.04) | 0.94 (± 0.02) | 8.03 (± 0.13) × $10^6$ | 73.2 (± 0.15) |
| | 1.66 (± 0.03) × $10^7$ | 5.08 (± 0.23) | 1.78 (± 0.09) | 4.31 (± 0.18) × $10^6$ | 306 (± 0.11) |
| **TF-CPY** | 1.56 (± 0.03) × $10^7$ | 1.71 (± 0.23) | 4.81 (± 0.48) | 1.15 (± 0.02) × $10^7$ | 109 (± 0.13) |
| | 1.69 (± 0.04) × $10^7$ | 4.06 (± 0.44) | 4.88 (± 0.55) | 9.19 (± 0.34) × $10^6$ | 240 (± 0.21) |
| **CF-CPY** | 2.50 (± 0.05) × $10^7$ | 3.32 (± 0.12) | 0.43 (± 0.02) | 2.89 (± 0.13) × $10^6$ | 133 (± 0.04) |
| | 3.54 (± 0.06) × $10^7$ | 4.18 (± 0.20) | 0.59 (± 0.04) | 4.35 (± 0.33) × $10^6$ | 118 (± 0.06) |
| **CP-CPY** | 1.53 (± 0.03) × $10^7$ | 3.39 (± 0.12) | 1.07 (± 0.04) | 3.67 (± 0.11) × $10^6$ | 222 (± 0.05) |
| | 2.87 (± 0.05) × $10^7$ | 7.26 (± 0.26) | 2.31 (± 0.11) | 6.92 (± 0.25) × $10^6$ | 253 (± 0.06) |

**Supplementary Table 3:** Average of kinetical parameters for SNAP-tag2 labeling with different TMR and CPY substrates calculated from data in Supplementary Table 2.

| **Substrate** | **$k_1$ (± s.d.) [$M^{-1}$ $s^{-1}$]** | **$k_{-1}$ (± s.d.) [$s^{-1}$]** | **$k_2$ (± s.d.) [$s^{-1}$]** | **$k_{app}$ (± s.d.) [$M^{-1}$ $s^{-1}$]** | **$K_d$ (± s.d.) [nM]** |
|---|---|---|---|---|---|
| **TF-TMR** | 1.51 (± 0.19) × $10^7$ | 0.48 (± 0.12) | 0.62 (± 0.23) | 8.22 (± 0.79) × $10^6$ | 32.0 (± 0.83) |
| **CF-TMR** | 2.09 (± 0.20) × $10^7$ | 3.13 (± 1.75) | 0.98 (± 0.47) | 6.21 (± 3.65) × $10^6$ | 151 (± 0.90) |
| **CP-TMR** | 1.94 (± 0.20) × $10^7$ | 2.71 (± 1.68) | 1.51 (± 0.40) | 7.82 (± 2.78) × $10^6$ | 151 (± 1.10) |
| **TF-CPY** | 1.62 (± 0.06) × $10^7$ | 2.88 (± 1.18) | 4.84 (± 0.04) | 1.04 (± 0.12) × $10^7$ | 175 (± 0.66) |
| **CF-CPY** | 3.02 (± 0.52) × $10^7$ | 3.75 (± 0.43) | 0.51 (± 0.08) | 3.62 (± 0.73) × $10^6$ | 126 (± 0.07) |
| **CP-CPY** | 2.20 (± 0.67) × $10^7$ | 5.33 (± 1.93) | 1.69 (± 0.62) | 5.29 (± 1.62) × $10^6$ | 237 (± 0.16) |

**Supplementary Table 4:** Kinetic parameters of SNAP-tag2 labeling with non-fluorescent substrates.

| Substrate | $k_{app}$ (± s.d.) [$M^{-1}$ $s^{-1}$] |
|---|---|
| **TF-Ac** | 1.04 (± 0.08) × $10^4$ |
| **TF-Nor** | 1.84 (± 0.03) × $10^5$ |
| **TF-PhN$_3$** | 1.68 (± 0.03) × $10^5$ |
| **TF-BCN** | 1.09 (± 0.02) × $10^5$ |

**Supplementary Table 5:** Comparison of extinction coefficients (ε) and quantum yields (QY) for fluorescently labeled SNAP$_f$-tag and SNAP-tag2.

| Substrate | SNAP$_f$-tag | | SNAP-tag2 | |
|---|---|---|---|---|
| | ε [$M^{-1}cm^{-1}$] | QY | ε [$M^{-1}cm^{-1}$] | QY |
| **CP-TMR** | 93133 (± 642) | 0.52 (± 0.01) | 98697 (± 1412) | 0.49 (± 0.01) |
| **TF-TMR** | 62560 (± 2136) | 0.48 (± 0.02) | 62120 (± 1272) | 0.46 (± 0.02) |
| **CP-MaP555** | 94353 (± 1577) | 0.50 (± 0.01) | 97707 (± 4280) | 0.46 (± 0.01) |
| **TF-MaP555** | 79223 (± 3133) | 0.46 (± 0.01) | 77027 (± 4660) | 0.46 (± 0.01) |
| **CP-CPY** | 147933 (± 3840) | 0.64 (± 0.03) | 158157 (± 1638) | 0.60 (± 0.03) |
| **TF-CPY** | 119760 (± 3036) | 0.68 (± 0.02) | 131307 (± 2614) | 0.66 (± 0.01) |
| **CP-SiR** | 75997 (± 2106) | 0.56 (± 0.02) | 83240 (± 6997) | 0.49 (± 0.02) |
| **TF-SiR** | 123533 (± 9063) | 0.53 (± 0.01) | 115953 (± 7203) | 0.51 (± 0.01) |
| **CF$_3$P-MaP618*** | ~2452 | n.d. | ~11829 | n.d. |
| **TF-MaP618*** | ~1809 | n.d. | ~4264 | n.d. |

*The ε for MaP618-substrates is estimated based on the maximum absorbance shown in Figure 2e.

### Supplementary Methods

#### Laser, detector and filter settings used for fluorescence-based experiments.

**Supplementary Table 6:** Settings used for YSD library screening *via* FACS.

| Fluorophore substrate | Excitation Laser | Filter | Emission Filter |
|---|---|---|---|
| Bilirubin | 488 nm | FITC | 527/32 nm |
| TMR | 561 nm | PE | 582/15 nm |
| MaP618 | 561 nm | PE-CF594 | 613/18 nm |
| Alexa647 | 640 nm | APC | 660/10 nm |

**Supplementary Table 7:** Filter settings used in FP measurements.

| Fluorophore substrate | Excitation filter | Emission filter |
|---|---|---|
| Alexa488, Fluorescein | 485/20 | 527/32 |
| TMR, MaP555 | 535/25 | 595/35 |
| CPY, MaP618, SiR | 620/20 | 680/30 |

**Supplementary Table 8:** Settings used for measuring the labeling performance of SNAP-tag proteins in live mammalian cells *via* flowcytometry.

| Fluorophore substrate | Excitation Laser | Filter | Emission Filter |
|---|---|---|---|
| mTurquoise2 | 405 nm | BV-421 | 450/40 nm |
| GFP | 488 nm | FITC | 530/30 nm |
| TMR | 561 nm | PE | 575/26 nm |
| CPY, MaP618 | 561 nm | PE-TexasRed | 610/20 nm |
| SiR | 640 nm | APC | 660/10 nm |

**Supplementary Table 9:** Settings used for confocal fluorescence imaging of SLPs in live mammalian cells

| Fluorophore | Excitation | Emission | HyD detector gain | Laser power |
|---|---|---|---|---|
| mTurquoise2 | 448 nm | 458-520 nm | 100 | 0.2-1.5 % |
| TMR | 552 nm | 562-625 nm | 100 | 0.2-2.0 % |
| CPY, MaP618 | 615 nm | 625-715 nm | 85-100 | 0.2-1.8 % |
| SiR | 652 nm | 662-805 nm | 100 | 0.70 % |

**Supplementary Table 10:** Microscopy settings for measuring labeling kinetics of SNAP-tag2 and $SNAP_f$-tag in live mammalian cells.

| Fluorophore | Excitation | Emission | HyD detector gain | Laser power |
|---|---|---|---|---|
| mTurquoise2 | 448 nm | 458-520 nm | 60-100 | 0.20 % |
| TMR | 552 nm | 562-625 nm | 60 | 0.30 % |
| CPY | 615 nm | 625-715 nm | 100 | 0.20 % |
| SiR | 652 nm | 662-805 nm | 100 | 0.30 % |

.

**Supplementary Table 11:** Settings used for CLSM and STED microscopy of SLPs in live mammalian cells.

| Figure | Dye | Excitation [nm] (%) | Emission [nm] | STED [nm] (%) | Pixel dwell time [µs] | Pixel size x-y [nm] | Size [µm] | Comment |
|---|---|---|---|---|---|---|---|---|
| **4a** | SiR | 640 (0.5) | 650 - 746 | - | 8 | 80 | 60 x 60 | CLSM |
| | | 640 (7) | 650 - 746 | 775 (30) | 8 | 25 | 60 x 60 | STED, 3 Line accu. |
| **4b** | SiR | 640 (1) | 650 - 746 | - | 7 | 25 | 10 x 10 | CLSM |
| | | 640 (7) | 650 - 746 | 775 (10) | 7 | 25 | 10 x 10 | STED, 8 Line accu. |
| | SiR | 640 (1) | 650 - 746 | - | 7 | 25 | 10 x 10 | CLSM |
| | | 640 (7) | 650 - 746 | 775 (10) | 7 | 25 | 10 x 10 | STED, 8 Line accu. |
| **4c** | SiR | 640 (6) | 650 - 746 | - | 7 | 100 | 60 x 60 | CLSM, 8 Line accu. |
| | | 640 (15) | 650 - 746 | 775 (15) | 7 | 80 | 60 x 60 | STED, 8 Line accu. |
| | | 640 (3) | 650 - 746 | - | 7 | 25 | 10 x 10 | CLSM, 5 Line accu. |
| | | 640 (12) | 650 - 746 | 775 (15) | 7 | 25 | 10 x 10 | STED, 5 Line accu. |
| **4d** | SiR | 640 (1) | 650 - 746 | - | 7 | 100 | 60 x 60 | CLSM, 8 Line accu. |
| | | 640 (9) | 650 - 746 | 775 (15) | 7 | 80 | 60 x 60 | STED 8 Line accu. |
| | | 640 (3) | 650 - 746 | - | 7 | 25 | 10 x 10 | CLSM, 5 Line accu. |
| | | 640 (12) | 650 - 746 | 775 (15) | 7 | 25 | 10 x 10 | STED, 5 Line accu. |
| **4e** | MaP618 | 561 (15) | 571 - 630 | - | 7 | 20 | 10 x 10 | CLSM, 10 Line accu. |
| | | 561 (26) | 571 - 630 | 775 (20) | 7 | 20 | 10 x 10 | STED, 10 Line accu. |
| | SiR | 640 (2) | 670 - 746 | - | 7 | 20 | 10 x 10 | CLSM, 10 Line accu. |
| | | 640 (3) | 670 - 746 | 775 (10) | 7 | 20 | 10 x 10 | STED, 10 Line accu. |

#### Buffers, media and reagents

**Supplementary Table 12:** Buffers, media and reagents used.

| Buffer/Media/Reagent | Composition |
|---|---|
| Activity buffer | 50 mM HEPES, 50 mM NaCl, pH = 7.3 |
| Cell growth medium | DMEM with phenol red + GlutaMAX™ (1 x), 4.5 g/L glucose, sodium pyruvate (1 x), 10 % FBS |
| DNA lysis buffer | 0.45 % Tween 20, 0.45 % TritonX100, 2.5 mM $MgCl_2$, 50 mM KCl, 10 mM Tris-HCl, 100 µg/mL proteinase K, pH = 8.3 |
| Electroporation buffer | 10 mM Tris-base, 270 mM Sucrose, 2.1 mM $MgCl_2 \times 6\ H_2O$, pH = 7.5, autoclaved |
| FACS buffer | 2 % FBS in PBS |
| FP buffer | Activity buffer, 0.1 mg/mL BSA, 1 mM DTT, pH = 7.3 |
| Gibson Assembly master mix (60 x) | 0.64 µL T5 exonuclease (10 U/µL), 20 µL Phusion polymerase (2 U/µL), 160 µL Taq ligase (40 U/µL), 700 µL $H_2O$, 320 µL 5 × ISO buffer, aliquoted to 20 µL/reaction |
| His-tag extraction buffer | 50 mM $KH_2PO_4$, 300 mM NaCl, 5 mM imidazole, 1 mM PMSF, 0.25 mg/mL lysozyme, pH = 8.0 |
| His-tag wash buffer | 50 mM $KH_2PO_4$, 300 mM NaCl, 10 mM imidazole, pH = 7.5 |
| His-tag elution buffer | 50 mM $KH_2PO_4$, 300 mM NaCl, 500 mM imidazole, pH = 7.5 |
| ISO buffer (5 x) | 25 % PEG-8000, 500 mM Tris–HCl, 50 mM $MgCl_2$, 50 mM DTT, 1 mM each dNTP, 5 mM NAD, pH = 7.5 |
| Imaging medium | DMEM without phenol red + GlutaMAX™ (1 x), 4.5 g/L glucose, sodium pyruvate (1 x), 10 % FBS |
| LB medium | 10 g/L bacto tryptone, 24 g/L yeast extract, 10 g/L NaCl, pH = 7.5 |
| SDCAA drop out medium | 20 g/L glucose, 6.7 g/L Difco yeast nitrogen base, 5 g/L Bacto casamino acids (without tryptophan), 38 mM $Na_2HPO_4 \times 12\ H_2O$, 62 mM $NaH_2PO_4 \times H_2O$, autoclaved (prior to addition of glucose) |
| SDCAA drop out agar plates | SDCAA drop out medium, 3 g/L agar, 1 M sorbitol |
| SGCAA drop out medium | 20 g/L galactose, 6.7 g/L Difco yeast nitrogen base, 5 g/L Bacto casamino acids (without tryptophan), 38 mM $Na_2HPO_4 \times 12\ H_2O$, 62 mM $NaH_2PO_4 \times H_2O$, autoclaved (prior to addition of galactose) |
| Tris buffer | 1 M Tris-HCl, pH = 8 |
| Tris-DTT buffer | 0.39 g DTT (2.5 M) dissolved in Tris buffer (800 µL), filtered |
| Tris-LiAc buffer | 1.02 g LiAc × 2 $H_2O$ dissolved in Tris buffer (2 mL), filtered |
| YPD medium | 20 g/L glucose, 20 g/L peptone, 10 g/L yeast extract, autoclaved (prior to addition of glucose) |
| YPD agar plates | YPD medium, 15 g/L agar (1.5 % w/v) |
| YPDS | 1:1 mixture of YPD and 1 M sorbitol |

#### Used plasmids and generated stable cell lines:

**Supplementary Table 13:** Most important plasmids and generated stable mammalian cell lines used.

| Name | Plasmid | Gene | Stable cell lines |
|---|---|---|---|
| pET51b(+)_SNAP$_f$-tag | pET51b(+) | SNAP$_f$-tag | |
| pET51b(+)_SNAP-tag2 | pET51b(+) | SNAP-tag2 | |
| pET51b(+)_CLIP$_f$-tag | pET51b(+) | CLIP$_f$-tag | |
| pET51b(+)_hAGT | pET51b(+) | hAGT | |
| pcDNA5/FRT_mEGFP-SNAP-tag | pCDNA5/FRT | SNAP-tag | U2OS Flp-In TREx |
| pcDNA5/FRT_mEGFP-CLIP-tag | pCDNA5/FRT | mEGFP-CLIP-tag | U2OS Flp-In TREx |
| pcDNA5/FRT_mEGFP-SNAP$_f$-tag | pCDNA5/FRT | mEGFP-SNAP$_f$-tag | U2OS Flp-In TREx |
| pcDNA5/FRT-HaloTag7-SNAP$_f$-P2A-NLS-mTurquoise2 | pCDNA5/FRT | HaloTag7-SNAP$_f$ and NLS-mTurquoise2 | U2OS Flp-In TREx |
| pcDNA5/FRT-HaloTag7-SNAP-tag2-P2A-NLS-mTurquoise2 | pCDNA5/FRT | HaloTag7-SNAP-tag2 and NLS-mTurquoise2 | U2OS Flp-In TREx |
| pcDNA5/FRT/TO-Cox8a(4x)-HaloTag7-T2A-EGFP | pCDNA5/FRT/TO | Cox8a(4x)-HaloTag7 and EGFP | HeLa Kyoto Flp-In |
| pcDNA5/FRT/TO-Cox8a(4x)-SNAP$_f$-tag-T2A-EGFP | pCDNA5/FRT/TO | Cox8a(4x)-SNAP$_f$-tag and EGFP | HeLa Kyoto Flp-In |
| pcDNA5/FRT/TO-Cox8a(4x)-SNAP-tag2-T2A-EGFP | pCDNA5/FRT/TO | Cox8a(4x)-SNAP-tag2 and EGFP | HeLa Kyoto Flp-In |
| pcDNA5/FRT/TO_TOMM20-SNAP-tag2-T2A-EGFP | pCDNA5/FRT/TO | TOMM20-SNAP-tag2 and EGFP | U2OS Flp-In TREx |
| pcDNA5/FRT/TO_CalR-SNAP-tag2-KDEL | pCDNA5/FRT/TO | CalR-SNAP-tag2_KDEL | U2OS Flp-In TREx |
| pcDNA5/FRT/TO_Vim-SNAP-tag2 | pCDNA5/FRT/TO | Vim-SNAP-tag2 | U2OS Flp-In TREx |
| pcDNA5/FRT/TO_Vim- SNAP$_f$-tag | pCDNA5/FRT/TO | Vim-SNAP$_f$-tag | U2OS Flp-In TREx |
| pCTCON2_HA-TEV-SNAP$_f$-tag-cMyc-Nt.BbvCI | pCTCON2 | SNAP$_f$-tag | |
| pCTCON2_ HA-TEV-SNAP-tag1.1-cMyc-Nt.BbvCI | pCTCON2 | SNAP-tag1.1 | |
| pJYDNg-appS4-SNAP-tag1.5 -Linker-Aga2p-HA-cMyc-eUnaG2 | pJYDNg | SNAP-tag1.5-Linker-Aga2p-HA-Myc-eUnaG2 | |
| pJYDNg-appS4-SNAP$_f$-tag-Linker-Aga2p-HA-cMyc-eUnaG2 | pJYDNg | SNAP$_f$-tag-Linker-Aga2p-HA-Myc-eUnaG2 | |
| pHIPZ_Pex3-SNAP$_f$-tag | pHIPZ | Pex3-SNAP$_f$-tag | |
| pHIPZ_Pex3-SNAP-tag2 | pHIPZ | Pex3-SNAP-tag2 | |

#### Degenerated primers used for protein library construction.

**Supplementary Table 14:** Primers used for generation of library 1 on SNAP-tag1.1 *via* one-pot saturation mutagenesis.

| Primer sequence | Target position in SNAP-tag1.1 |
|---|---|
| GCGACGGCGACGTA**NNK**GGCTACGAAGGCGG | G156X |
| GACGGCGACGTAGGT**NNK**TACGAAGGCGGCCTG | G157X |
| GCGACGTAGGTGGC**NNK**GAAGGCGGCCTGG | Y158X |
| GCGACGTAGGTGGCTAC**NNK**GGCGGCCTGGC | E159X |
| GACGTAGGTGGCTACGAA**NNK**GGCCTGGCTGTGAAAG | G160X |
| GGTGGCTACGAAGGC**NNK**CTGGCTGTGAAAGAATGGC | G161X |

**Supplementary Table 15:** Primers used for generation of library 2 on SNAP-tag1.1 *via* one-pot saturation mutagenesis.

| Primer sequence | Target positions in SNAP-tag1.1 |
|---|---|
| GAGCAGGGCCTGCAC**NNK**ATC**NNK**TTC**NNK**GGC**NNK**GGTACGAGCGCTGCG | E30X, I32X, L34X, K36X |

**Supplementary Table 16:** Primers used for generation of library 3 *via* site-directed saturation mutagenesis.

| Forward primer sequence | Reverse primer sequence | Target position |
|---|---|---|
| CGTGGGCAATGGTACGAGCGCTGCGGATG | AAATAGATGCG**MNN**CAGGCCCTGCTCGC | H29X |
| CGTGGGCAATGGTACGAGCGCTGCGGATG | AAATA**MNN**GCGGTGCAGGCCCTGCTCGC | I31X |
| T**NNK**GTGGGCAATGGTACGAGCGC | TAGATGCGGTGCAGGCCC | F33X |
| TTTC**NNK**GGCAATGGTACGAGCGCTGC | TAGATGCGGTGCAGGCCC | V34X |
| CGTG**NNK**AATGGTACGAGCGCTGCGGATGCGG | AAATAGATGCGGTGCAGGCCCTGCTCGCA | G35X |

**Supplementary Table 17:** Primers used for generation of site-directed saturation mutagenesis libraries 4-9 *via* assembly PCR.

| Library | Forward primer sequence | Reverse primer sequence | Target mutation in SNAP-tag1.5 |
|---|---|---|---|
| **4** | GCGACGGCGACGTA**NNKNNKNNK**GAAGGCCCTCTGGCTGTGAAAGAATGGC | GCCATTCTTTCACAGCCAGAGGGCCTTC**MNNMNNMNN**TACGTCGCCGTCGCC | G156X, G157X, Y158X |
| **5** | GGCGACGGCGACGTAGGT**NNKNNKNNK**GGCCCTCTGGCTGTGAAAGAATGGC | GCCATTCTTTCACAGCCAGAGGGCC**MNNMNNMNN**ACCTACGTCGCCGTCGCC | G157X, Y158X, E159X |
| **6** | GGCGACGGCGACGTAGGTGGC**NNKNNKNNK**CCTCTGGCTGTGAAAGAATGGC | GCCATTCTTTCACAGCCAGAGG**MNNMNNMNN**GCCACCTACGTCGCCGTCGCC | Y158X, E159X, G160X |
| **7** | GGCGACGGCGACGTAGGTGGCTAC**NNKNNKNNK**CTGGCTGTGAAAGAATGGC | GCCATTCTTTCACAGCCAG**MNNMNNMNN**GTAGCCACCTACGTCGCCGTCGCC | E159X, G160X, P161X |
| **8** | GGCGACGGCGACGTAGGT**NNK**TAC**NNK**GGC**NNK**CTGGCTGTGAAAGAATGGC | GCCATTCTTTCACAGCCAG**MNN**GCC**MNN**GTA**MNN**ACCTACGTCGCCGTCGCC | G157X, E159X, P161X |
| **9** | GGCGACGGCGACGTA**NNK**GGC**NNK**GAA**NNK**CCTCTGGCTGTGAAAGAATGGC | GCCATTCTTTCACAGCCAGAGG**MNN**TTC**MNN**GCC**MNN**TACGTCGCCGTCGCC | G156X, Y158X, G160X |

| Flanking region for assembly | Forward primer sequence | Reverse primer sequence |
|---|---|---|
| N-terminus | GAGGTTCCCATCTATTTTCACCGC | TACGTCGCCGTCGCCCTG |
| C-terminus | CTGGCTGTGAAAGAATGGCTGC | CCTTCACCATGATATGAAGAGTTGACAACG |

**Supplementary Table 18:** NGS primers used for amplification of libraries after FACS screen.

| Library | Adapter sequence | Primer-binding sequence |
|---|---|---|
| **sDMSL** | CGAGATCTCT | GCTGCTTCTTCTGCTTTGG |
| | GATACTTGCA | |
| | TGCAGCTAAG | |
| | CCGAAGTGGA | |
| | CCGCCTGTTA | GCATGGCTGAACGCATATTTTC |
| | ATCAGCGAGG | |
| | CGGTCTAACG | |
| | GAGCTCGCTA | |
| | GCGTAATTAC | CCGTGCCACCGTG |
| | TCCGCGAAGT | |
| | AGACCGTTAT | |
| | AGACGGAGAG | |
| **Libraries 4-9** | GATATCAGGA | TCTGATCCCGTGCCAC |
| | ATCGAGTTCC | |
| | GTTAGCAGCC | |
| | AATTAGGCCG | |

### Protein sequences

#### Bacterial expression

General color code: Strep-tag – Enterokinase CDS - Hisx10-tag – TEV cleavage site – |Protein| sequence – **Introduced mutation** – **Catalytic residue** - linkers

>pET51b-SNAP-tag
MHHHHHHHHHHENLYFQ|GMDKDCEMKRTTLDSPLGKLELSGCEQGLHEIIFLGKGTSAADAVEVPAPAAVLGGPEP
LMQATAWLNAYFHQPEAIEEFPVPALHHPVFQQESFTRQVLWKLLKVVKFGEVISYSHLAALAGNPAATAAVKTALSG
NPVPILIP**C**HRVVQGDLDVGGYEGGLAVKEWLLAHEGHRLGKPGLG

>pET51b-SNAP$_f$-tag
MASWSHPQFEKGADDDDKVPH|MDKDCEMKRTTLDSPLGKLELSGCEQGLH**R**IIFLGKGTSAADAVEVPAPAAVLGG
PEPLMQATAWLNAYFHQPEAIEEFPVPALHHPVFQQESFTRQVLWKLLKVVKFGEVISYSHLAALAGNPAATAAVKTA
LSGNPVPILIP**C**HRVVQGDLDVGGYEGGLAVKEWLLAHEGHRLGKPGLG|APGFSSISAHHHHHHHHHH

>pET51b-SNAP-tag2

MASWSHPQFEKGADDDDKVPH|MDKDCEMKRTT**Y**DSPLGKL**L**LSGCEQGLH**R**I**Y**F**V**G**NGQGEQGPP**GPEPLMQAT
AWLNAYFHQPEAIEEFPVPALHHPVFQQESFTRQVLWKLLKVVKFGEVISYS**Q**LAALAGNPAATAAVKTAL**R**GNPVPIL
IP**C**HRVVQGD**G**DVGGYEG**P**L**Y**VKEWLLAHEGHRLGKPGLG|APGFSSISAHHHHHHHHHH

>pET51b-hAGT
MASWSHPQFEKGADDDDKVPH|MDKDCEMKRTTLDSPLGKLELSGCEQGLHEIKLLGKGTSAADAVEVPAPAAVLG
GPEPLMQCTAWLNAYFHQPEAIEEFPVPALHHPVFQQESFTRQVLWKLLKVVKFGEVISYQQLAALAGNPKAARAVG
GAMRGNPVPILIP**C**HRVVCSSGAVGNYSGGLAVKEWLLAHEGHRLGKPGLGGSSGLAGAWLKGAGATSGSPPAGR
N|APGFSSISAHHHHHHHHHH

>pET51b-CLIP-tag
MHHHHHHHHHHENLYFQG|MDKDCEMKRTTLDSPLGKLELSGCEQGLHEIIFLGKGTSAADAVEVPAPAAVLGGPEP
LIQATAWLNAYFHQPEAIEEFPVPALHHPVFQQESFTRQVLWKLLKVVKFGEVISESHLAALVGNPAATAAVNTALDG
NPVPILIP**C**HRVVQGDSDVGPYLGGLAVKEWLLAHEGHRLGKPGLGG

>pET51b-CLIP$_f$-tag
MHHHHHHHHHHENLYFQG|MDKDCEMKRTTLDSPLGKLELSGCEQGLH**R**IIFLGKGTSAADAVEVPAPAAVLGGPEP
LIQATAWLNAYFHQPEAIEEFPVPALHHPVFQQESFTRQVLWKLLKVVKFGEVISESHLAALVGNPAATAAVNTALDG
NPVPILIP**C**HRVVQGDSDVGPYLGGLAVKEWLLAHEGHRLGKPGLGG

#### Expression in mammalian cells

General color code: Kozak – Localization/fusion protein sequence –Cleavage sequence– |SLP sequence| –
**Introduced mutation – Catalytic residue -** linkers

>mEGFP (N-terminally fused)

MVSKGEELFTGVVPILVELDGDVNGHKFSVSGEGEGDATYGKLTLKFICTTGKLPVPWPTLVTTLTYGVQCFSRYPDH
MKQHDFFKSAMPEGYVQERTIFFKDDGNYKTRAEVKFEGDTLVNRIELKGIDFKEDGNILGHKLEYNYNSHNVYIMAD
KQKNGIKVNFKIRHNIEDGSVQLADHYQQNTPIGDGPVLLPDNHYLSTQSKLSKDPNEKRDHMVLLEFVTAAGITLGM
DELYK|[…]

>pcDNA5/FRT-HaloTag7-SNAPf-tag- NLS-P2A-NLS-mTurquoise2

GG|MGSEIGTGFPFDPHYVEVLGERMHYVDVGPRDGTPVLFLHGNPTSSYVWRNIIPHVAPTHRCIAPDLIGMGKSDK
PDLGYFFDDHVRFMDAFIEALGLEEVVLVIH**D**WGSALGFHWAKRNPERVKGIAFMEFIRPIPTWDEWPEFARETFQAF
RTTDVGRKLIIDQNVFIEGTLPMGVVRPLTEVEMDHYREPFLNPVDREPLWRFPNELPIAGEPANIVALVEEYMDWLH
QSPVPKLLFWGTPGVLIPPAEAARLAKSLPNCKAVDIGPGLNLLQEDNPDLIGSEIARWLSTLEISG|SGRPPPPPPPPP
PPPPPPPPPPPPPPPPPPPPPGGRSRSLE|MDKDCEMKRTTLDSPLGKLELSGCEQGLH**R**IIFLGKGTSAADAVEVPA
PAAVLGGPEPLMQATAWLNAYFHQPEAIEEFPVPALHHPVFQQESFTRQVLWKLLKVVKFGEVISYSHLAALAGNPAA
TAAVKTALSGNPVPILIP**C**HRVVQGDLDVGGYEGGLAVKEWLLAHEGHRLGKPGLG|GAPDPKKKRKVDPKKKRKVD
PKKKRKELRASPQATNFSLLKQAGDVEENPGPSRMAPKKKRKVM|VSKGEELFTGVVPILVELDGDVNGHKFSVSGE
GEGDATYGKLTLKFICTTGKLPVPWPTLVTTLSWGVQCFARYPDHMKQHDFFKSAMPEGYVQERTIFFKDDGNYKTR
AEVKFEGDTLVNRIELKGIDFKEDGNILGHKLEYNYFSDNVYITADKQKNGIKANFKIRHNIEDGGVQLADHYQQNTPIG
DGPVLLPDNHYLSTQSKLSKDPNEKRDHMVLLEFVTAAGITLGMDELYK

>pcDNA5/FRT-HaloTag7-SNAP-tag2-NLS-P2A-NLS-mTurquoise2

GG|MGSEIGTGFPFDPHYVEVLGERMHYVDVGPRDGTPVLFLHGNPTSSYVWRNIIPHVAPTHRCIAPDLIGMGKSDK
PDLGYFFDDHVRFMDAFIEALGLEEVVLVIH**D**WGSALGFHWAKRNPERVKGIAFMEFIRPIPTWDEWPEFARETFQAF
RTTDVGRKLIIDQNVFIEGTLPMGVVRPLTEVEMDHYREPFLNPVDREPLWRFPNELPIAGEPANIVALVEEYMDWLH
QSPVPKLLFWGTPGVLIPPAEAARLAKSLPNCKAVDIGPGLNLLQEDNPDLIGSEIARWLSTLEISG|SGRPPPPPPPPP
PPPPPPPPPPPPPPPPPPPPPGGRSRSLE|MDKDCEMKRTT**Y**DSPLGKL**L**LSGCEQGLH**R**I**Y**F**V**G**NGQGEQGPP**GP
EPLMQATAWLNAYFHQPEAIEEFPVPALHHPVFQQESFTRQVLWKLLKVVKFGEVISYS**Q**LAALAGNPAATAAVKTAL
**R**GNPVPILIP**C**HRVVQGD**G**DVGGYEG**P**L**Y**VKEWLLAHEGHRLGKPGLG|GAPDPKKKRKVDPKKKRKVDPKKKRKEL
RASPQATNFSLLKQAGDVEENPGPSRMAPKKKRKVM|VSKGEELFTGVVPILVELDGDVNGHKFSVSGEGEGDATYG
KLTLKFICTTGKLPVPWPTLVTTLSWGVQCFARYPDHMKQHDFFKSAMPEGYVQERTIFFKDDGNYKTRAEVKFEGD
TLVNRIELKGIDFKEDGNILGHKLEYNYFSDNVYITADKQKNGIKANFKIRHNIEDGGVQLADHYQQNTPIGDGPVLLPD
NHYLSTQSKLSKDPNEKRDHMVLLEFVTAAGITLGMDELYK

>pcDNA5/FRT/TO-[Cox8a]x4-SNAPf-tag-T2A-EGFP

MSVLTPLLLRGLTGSARRLPVPRAKIHSLSVLTPLLLRGLTGSARRLPVPRAKIHSLSVLTPLLLRGLTGSARRLPVPRA
KIHSLSVLTPLLLRGLTGSARRLPVPRAKIHSLGGSGGS|DKDCEMKRTTLDSPLGKLELSGCEQGLH**R**IIFLGKGTSAA
DAVEVPAPAAVLGGPEPLMQATAWLNAYFHQPEAIEEFPVPALHHPVFQQESFTRQVLWKLLKVVKFGEVISYSHLAA
LAGNPAATAAVKTALSGNPVPILIP**C**HRVVQGDLDVGGYEGGLAVKEWLLAHEGHRLGKPGLG|GRGSGEGRGSLLT
CGDVEENPGP|VSKGEELFTGVVPILVELDGDVNGHKFSVSGEGEGDATYGKLTLKFICTTGKLPVPWPTLVTTLTYG
VQCFSRYPDHMKQHDFFKSAMPEGYVQERTIFFKDDGNYKTRAEVKFEGDTLVNRIELKGIDFKEDGNILGHKLEYNY
NSHNVYIMADKQKNGIKVNFKIRHNIEDGSVQLADHYQQNTPIGDGPVLLPDNHYLSTQSALSKDPNEKRDHMVLLEF
VTAAGITLGMDELYK

>pcDNA5/FRT/TO-[Cox8a]x4-SNAP-tag2-T2A-EGFP

MSVLTPLLLRGLTGSARRLPVPRAKIHSLSVLTPLLLRGLTGSARRLPVPRAKIHSLSVLTPLLLRGLTGSARRLPVPRA
KIHSLSVLTPLLLRGLTGSARRLPVPRAKIHSLGGSGGS|DKDCEMKRTT**Y**DSPLGKL**L**LSGCEQGLH**R**I**Y**F**V**G**NGQGE**
**QGPP**GPEPLMQATAWLNAYFHQPEAIEEFPVPALHHPVFQQESFTRQVLWKLLKVVKFGEVISYS**Q**LAALAGNPAAT
AAVKTAL**R**GNPVPILIP**C**HRVVQGD**G**DVGGYEG**P**L**Y**VKEWLLAHEGHRLGKPGLG|GRGSGEGRGSLLTCGDVEEN
PGP|VSKGEELFTGVVPILVELDGDVNGHKFSVSGEGEGDATYGKLTLKFICTTGKLPVPWPTLVTTLTYGVQCFSRYP

DHMKQHDFFKSAMPEGYVQERTIFFKDDGNYKTRAEVKFEGDTLVNRIELKGIDFKEDGNILGHKLEYNYNSHNVYIM
ADKQKNGIKVNFKIRHNIEDGSVQLADHYQQNTPIGDGPVLLPDNHYLSTQSALSKDPNEKRDHMVLLEFVTAAGITLG
MDELYK

>pcDNA5/FRT/TO-[Cox8a]x4-HaloTag7-T2A-EGFP

MSVLTPLLLRGLTGSARRLPVPRAKIHSLSVLTPLLLRGLTGSARRLPVPRAKIHSLSVLTPLLLRGLTGSARRLPVPRA
KIHSLSVLTPLLLRGLTGSARRLPVPRAKIHSLGGSGGS|GSEIGTGFPFDPHYVEVLGERMHYVDVGPRDGTPVLFLH
GNPTSSYVWRNIIPHVAPTHRCIAPDLIGMGKSDKPDLGYFFDDHVRFMDAFIEALGLEEVVLVIH**D**WGSALGFHWAK
RNPERVKGIAFMEFIRPIPTWDEWPEFARETFQAFRTTDVGRKLIIDQNVFIEGTLPMGVVRPLTEVEMDHYREPFLNP
VDREPLWRFPNELPIAGEPANIVALVEEYMDWLHQSPVPKLLFWGTPGVLIPPAEAARLAKSLPNCKAVDIGPGLNLL
QEDNPDLIGSEIARWLSTLEISG|GRGSGEGRGSLLTCGDVEENPGP|VSKGEELFTGVVPILVELDGDVNGHKFSVSG
EGEGDATYGKLTLKFICTTGKLPVPWPTLVTTLTYGVQCFSRYPDHMKQHDFFKSAMPEGYVQERTIFFKDDGNYKT
RAEVKFEGDTLVNRIELKGIDFKEDGNILGHKLEYNYNSHNVYIMADKQKNGIKVNFKIRHNIEDGSVQLADHYQQNTPI
GDGPVLLPDNHYLSTQSALSKDPNEKRDHMVLLEFVTAAGITLGMDELYK

>pcDNA5/FRT/TO-Vim-SNAP$_f$-tag

MSTRSVSSSSYRRMFGGPGTASRPSSSRSYVTTSTRTYSLGSALRPSTSRSLYASSPGGVYATRSSAVRLRSSVPG
VRLLQDSVDFSLADAINTEFKNTRTNEKVELQELNDRFANYIDKVRFLEQQNKILLAELEQLKGQGKSRLGDLYEEEMR
ELRRQVDQLTNDKARVEVERDNLAEDIMRLREKLQEEMLQREEAENTLQSFRQDVDNASLARLDLERKVESLQEEIA
FLKKLHEEEIQELQAQIQEQHVQIDVDVSKPDLTAALRDVRQQYESVAAKNLQEAEEWYKSKFADLSEAANRNNDALR
QAKQESTEYRRQVQSLTCEVDALKGTNESLERQMREMEENFAVEAANYQDTIGRLQDEIQNMKEEMARHLREYQDL
LNVKMALDIEIATYRKLLEGEESRISLPLPNFSSLNLRETNLDSLPLVDTHSKRTLLIKTVETRDGQVINETSQHHDDLEG
DPPVAT|DKDCEMKRTTLDSPLGKLELSGCEQGLHRIIFLGKGTSAADAVEVPAPAAVLGGPEPLMQATAWLNAYFHQ
PEAIEEFPVPALHHPVFQQESFTRQVLWKLLKVVKFGEVISYSHLAALAGNPAATAAVKTALSGNPVPILIP**C**HRVVQG
DLDVGGYEGGLAVKEWLLAHEGHRLGKPGLG|

>pcDNA5/FRT/TO-Vim-SNAP-tag2

MSTRSVSSSSYRRMFGGPGTASRPSSSRSYVTTSTRTYSLGSALRPSTSRSLYASSPGGVYATRSSAVRLRSSVPG
VRLLQDSVDFSLADAINTEFKNTRTNEKVELQELNDRFANYIDKVRFLEQQNKILLAELEQLKGQGKSRLGDLYEEEMR
ELRRQVDQLTNDKARVEVERDNLAEDIMRLREKLQEEMLQREEAENTLQSFRQDVDNASLARLDLERKVESLQEEIA
FLKKLHEEEIQELQAQIQEQHVQIDVDVSKPDLTAALRDVRQQYESVAAKNLQEAEEWYKSKFADLSEAANRNNDALR
QAKQESTEYRRQVQSLTCEVDALKGTNESLERQMREMEENFAVEAANYQDTIGRLQDEIQNMKEEMARHLREYQDL
LNVKMALDIEIATYRKLLEGEESRISLPLPNFSSLNLRETNLDSLPLVDTHSKRTLLIKTVETRDGQVINETSQHHDDLEG
DPPVAT|DKDCEMKRTT**Y**DSPLGKL**L**LSGCEQGLH**R**I**Y**F**V**G**NGQGEQGPP**GPEPLMQATAWLNAYFHQPEAIEEFPV
PALHHPVFQQESFTRQVLWKLLKVVKFGEVISYS**Q**LAALAGNPAATAAVKTAL**R**GNPVPILIP**C**HRVVQGD**G**DVGGYE
G**P**L**Y**VKEWLLAHEGHRLGKPGLG|

>pcDNA5/FRT/TO-CalR-SNAP-tag2-KDEL (Sec61b)

MLLSVPLLLGLLGLAVAGGSGGSEFGS|DKDCEMKRTT**Y**DSPLGKL**L**LSGCEQGLH**R**I**Y**F**V**G**NGQGEQGPP**GPEPLM
QATAWLNAYFHQPEAIEEFPVPALHHPVFQQESFTRQVLWKLLKVVKFGEVISYS**Q**LAALAGNPAATAAVKTAL**R**GNP
VPILIP**C**HRVVQGD**G**DVGGYEG**P**L**Y**VKEWLLAHEGHRLGKPGLG|GAPGFSSISAKDEL

>pcDNA5/FRT/TO-TOMM20-SNAP-tag2-T2A-EGFP

MVGRNSAIAAGVCGALFIGYCIYFDRKRRSDPNFKNRLRERRKKQKLAKERAGLSKLPDLKDAEAVQKFFLEEIQLGE
ELLAQGEYEKGVDHLTNAIAVCGQPQQLLQVLQQTLPPPVFQMLLTKLPTISQRIVSAQSLAEDDVEGGSGDPPVGG|
DKDCEMKRTT**Y**DSPLGKL**L**LSGCEQGLH**R**I**Y**F**V**G**NGQGEQGPP**GPEPLMQATAWLNAYFHQPEAIEEFPVPALHHP
VFQQESFTRQVLWKLLKVVKFGEVISYS**Q**LAALAGNPAATAAVKTAL**R**GNPVPILIP**C**HRVVQGD**G**DVGGYEG**P**L**Y**VK
EWLLAHEGHRLGKPGLG|GRGSGEGRGSLLTCGDVEENPGPVSKGEELFTGVVPILVELDGDVNGHKFSVSGEGEG
DATYGKLTLKFICTTGKLPVPWPTLVTTLTYGVQCFSRYPDHMKQHDFFKSAMPEGYVQERTIFFKDDGNYKTRAEV
KFEGDTLVNRIELKGIDFKEDGNILGHKLEYNYNSHNVYIMADKQKNGIKVNFKIRHNIEDGSVQLADHYQQNTPIGDG
PVLLPDNHYLSTQSALSKDPNEKRDHMVLLEFVTAAGITLGMDELYK

#### Expression on yeast surface

General color code: Gal1 promoter – Aga2P –affinity tags –TEV cleavage site -Factor Xa site - |Protein sequence| – **Introduced mutation – Catalytic residue –** linkers – eUnaG2

>pCTcon2-SNAP-tag1.1

MQKLHNHFTNTFNIFGLYYFLFKCNKSINKKLLIYLYTLTSRRKNPGSNSLLHTFSIKMQLLRCFSIFSVIASVLAQELTTI
CEQIPSPTLESTPYSLSTTTILANGKAMQGVFEYYKSVTFVSNCGSHPSTTSKGSPINTQYVFKDNSSTIEGRYPYDVP
DYALQENLYFQGLQASGGGGSGGGGSGGGGS|MDKDCEMKRTTLDSPLGKLELSGCEQGLH**R**IIFLGKGTSAADAV
EVPAPAAVLGGPEPLMQATAWLNAYFHQPEAIEEFPVPALHHPVFQQESFTRQVLWKLLKVVKFGEVISYSHLAALAG
NPAATAAVKTAL**R**GNPVPILIP**C**HRVVQGD**G**DVGGYEGGLAVKEWLLAHEGHRLGKPGLG|GSGGSEQKLISEEDL

>pJYDNg-SNAP-tag1.5

MYYFLFKCNKSINKKLLIYLYTLTSRRKNPGSNSLLHTFSIKMRFPSIFTAVVFAASSALAAPANGT|MDKDCEMKRTTL
DSPLGKLELSGCEQGLH**R**I**Y**F**V**G**N**GTSAADAVEVPAP**GHPEPQ**PEPLMQATAWLNAYFHQPEAIEEFPVPALHHPVF
QQESFTRQVLWKLLKVVKFGEVISYSHLAALAGNPAATAAVKTAL**R**GNPVPILIP**C**HRVVQGD**G**DVGGYEG**P**LAVKE
WLLAHEGHRLGKPGLG|AAAFSQKLDINLLDNVVNSSYHGEGVSGGSAQELTTICEQIPSPTLESTPYSLSTTTILANGK
AMQGVFEYYKSVTFVSNCGSHPSTTSKGSPINTQYVFKDNSSTIEGRYPYDVPDYALQASGGGGSGGGGSGGGGS
ASHEQKLISEEDL|MLEKFVGTWKIESSENFGEYLKAIGAPKELADAGDATTPVLYISQKDGDKMTVKIENGPPTFLDTQ
VSFKLGEEFDEFPSDRRKGVKSVVNLSGEKLVYVQKWDGKETTYVREIKDGKLVVTLTMGDVVAVRSYRRASE

#### Scripts

##### Rosetta scripts for unstructured loop redesign

Mutagenesis and minimization script

```
<ROSETTASCRIPTS>

<SCOREFXNS>

	<ScoreFunction name="sfxn_stand" weights="ref2015">
		<Reweight scoretype="atom_pair_constraint" weight="2"/>
		<Reweight scoretype="angle_constraint" weight="2"/>
		<Reweight scoretype="dihedral_constraint" weight="2"/>
		<Reweight scoretype="coordinate_constraint" weight="1"/>
		<Reweight scoretype="metalbinding_constraint" weight="2"/>
	</ScoreFunction>

</SCOREFXNS>

<TASKOPERATIONS>

	<ReadResfile name="resfile" filename="in/resfile"/>
	<InitializeFromCommandline name="extra_rot" />

</TASKOPERATIONS>

<FILTERS>

	<ScoreType name="score_filter" score_type="total_score" threshold="-200"
scorefxn="sfxn_stand" />

</FILTERS>

<MOVERS>

	<AddConstraints name="bond_cst">
		<FileConstraintGenerator name="bond_cst"
			filename="in/chemical_bond.cst"/>
	</AddConstraints>

	<SetupMetalsMover name="metal_cst"/>

	<DeclareBond name="ligand_bond" res1="124" res2="160"
		atom1="SG" atom2="C30"/>

	<AtomTree name="set_fold_tree" fold_tree_file="in/fold_tree"/>

	<FastRelax name="relax" scorefxn="sfxn_stand"
		disable_design="true"
		task_operations="extra_rot,resfile"
		repeats="5" cartesian="false"
		ramp_down_constraints="false"
		min_type="lbfgs_armijo_nonmonotone">
		<MoveMap name="move_map_1" bb="false" chi="true"
```

```
                    jump="true"/>
      </FastRelax>

      <FastDesign name="design" scorefxn="sfxn_stand"
             disable_design="false"
             task_operations="extra_rot,resfile"
             repeats="5" cartesian="false"
             ramp_down_constraints="false"
             min_type="lbfgs_armijo_nonmonotone">
             <MoveMap name="move_map_1" bb="true" chi="true"
                    jump="true"/>
      </FastDesign>

</MOVERS>

<PROTOCOLS>

      <Add mover="bond_cst"/>
      <Add mover="metal_cst"/>
      <Add mover="ligand_bond"/>
      <Add mover="set_fold_tree"/>
      <Add mover="relax"/>
      <Add mover="design"/>
      <Add filter="score_filter"/>

</PROTOCOLS>

</ROSETTASCRIPTS>
```

Resfile:

```
NATAA
start

27  A PIKAA R
29  A PIKAA Y
31  A PIKAA V
33  A PIKAA N
114 A PIKAA R
132 A PIKAA G
140 A PIKAA P
```

Constraint file:

```
AtomPair SG  124 C30 160                 HARMONIC 1.786     0.005
Angle    CB  124 SG  124 C30 160         HARMONIC 1.9457146 0.0628319
Angle    SG  124 C30 160 C32 160         HARMONIC 2.1478830 0.0628319
Dihedral CB  124 SG  124 C30 160 C32 160 HARMONIC 3.1102080 0.0628319
```

Fold tree:

```
FOLD_TREE  EDGE 124 34 -1  EDGE 124 158 -1 EDGE 64 159 1  EDGE 124 160 2  EDGE 42 7 3
EDGE 7 1 -1  EDGE 7 33 -1
```

RosettaScripts was run with the following flags:

```
-nstruct 10000
-parser:protocol in/relax.xml
-in:file:extra_res_path in/par
-ex1
-ex2
-ex3
-use_input_sc
-flip_HNQ
-no_optH false
-packing:extrachi_cutoff 8
-packing:ex1aro
-packing:ex2aro
-in:file:fullatom
-in:file:s in/SNAP.pdb
--out:path:all out
```

#### NGS analysis of libraries

Demultiplex command:

```
je demultiplex F1=[path/to/fwd_reads] \
       F2=[path/to/rev_reads] O=out_split_reads \
       BF=[path/to/barcodes_file] GZ=false UF1=unassigned_1 \
       UF2=unassigned_2 M=jemultiplexer_out_stats
```

Combining paired reads command:

```
cat [path/to/fwd_reads] [path/to/rev_reads] >> output
```

Trimming command:

```
cutadapt -g [primer_sequences_file] \
       -a [adapter_sequences_file] --times 3 --overlap 6 \
       -e 0.2 -m 16 -j 8 -o output [input_reads] > \
       trim.log
```

Align command:

```
bowtie2 -p 8 -N 1 -L 6 --n-ceil L,150,1 --mp 6,2 --np 0 \
       --rdg 50,6 --rfg 50,6 --score-min L,-1,-0.5 -x \
       [template_sequence_file] -U [input.reads] -S output 2> \
       align.log"
```

Conversion to bam file command:

```
samtools view -b [input_file] > output
```

### Example DynaFit[1] scripts

#### SNAP-tag variant kinetics fitted to model 1

```
[task]
   data  = progress
   task  = fit
   confidence = monte-carlo

[mechanism]
    P + S ----> P.S : kapp

[constants]; units: nM, sec
    kapp = 0.001?

[concentrations], units: nM
    S = 20?

[responses]
    P.S = 2?

[data]
   delay 0
   offset 134
   directory  path/to/data
   sheet      data.csv

    column 2 | conc P = 10.4 | label 10.4
   column 3 | conc P = 10.4 | label 10.4
   column 4 | conc P = 10.4 | label 10.4

   column 5 | conc P = 118.5 | label 118.5
   column 6 | conc P = 118.5 | label 118.5
   column 7 | conc P = 118.5 | label 118.5

   column 8  | conc P = 15.6 | label 15.6
   column 9  | conc P = 15.6 | label 15.6
   column 10 | conc P = 15.6 | label 15.6

   column 11 | conc P = 177.8 | label 177.8
   column 12 | conc P = 177.8 | label 177.8
   column 13 | conc P = 177.8 | label 177.8

   column 14 | conc P = 23.4 | label 23.4
   column 15 | conc P = 23.4 | label 23.4
   column 16 | conc P = 23.4 | label 23.4

   column 17 | conc P = 266.7 | label 266.7
   column 18 | conc P = 266.7 | label 266.7
   column 19 | conc P = 266.7 | label 266.7

   column 20 | conc P = 35.1 | label 35.1
   column 21 | conc P = 35.1 | label 35.1
   column 22 | conc P = 35.1 | label 35.1

   column 23 | conc P = 4.6 | label 4.6
   column 24 | conc P = 4.6 | label 4.6
   column 25 | conc P = 4.6 | label 4.6

   column 26 | conc P = 400 | label 400
   column 27 | conc P = 400 | label 400
   column 28 | conc P = 400 | label 400
```

```
   column 29 | conc P = 52.7 | label 52.7
   column 30 | conc P = 52.7 | label 52.7
   column 31 | conc P = 52.7 | label 52.7

   column 32 | conc P = 6.9 | label 6.9
   column 33 | conc P = 6.9 | label 6.9
   column 34 | conc P = 6.9 | label 6.9

   column 35 | conc P = 600 | label 600
   column 36 | conc P = 600 | label 600
   column 37 | conc P = 600 | label 600

   column 38 | conc P = 79 | label 79
   column 39 | conc P = 79 | label 79
   column 40 | conc P = 79 | label 79

   column 41 | conc P = 900 | label 900
   column 42 | conc P = 900 | label 900
   column 43 | conc P = 900 | label 900


[output]
  directory path/to/output/folder

[settings]
{ConfidenceIntervals}
   LevelPercent = 95
{Output}
   XAxisLabel = time [s]
   YAxisLabel = anisotropy

[end]
```

#### SNAP-tag2 kinetics fitted to model 2

```
[task]
   data  = progress
   task  = fit
   confidence = monte-carlo
[mechanism]
   P + S <===> P.S     :     k1    k_m1
   P.S ----> Z           :     k2
[constants] ; units: uM, sec
   k1 = 15?
   k_m1 = 0.01?
   k2 = 3?
[concentrations] ; units: uM
  S = 0.5?
[responses]
intensive
   S = 0.071
   P.S = 0.2?
   Z   = 1.0 * P.S

[data]
   directory  path/to/data
   sheet      data.csv

   column   2 | conc P = 2.5 | label 2.5

   column   3 | conc P = 2.0 | label 2.0

   column   4 | conc P = 1.5 | label 1.5

   column   5 | conc P = 1.0 | label 1.0

   column   6 | conc P = 0.5 | label 0.5

   column   7 | conc P = 0.417 | label 0.417


[output]
  directory path/to/output/folder

[settings]
{ConfidenceIntervals}
   LevelPercent = 95

{Constraints}
   Constants = 1000000000000000

{Output}
   XAxisLabel = time [s]
   YAxisLabel = anisotropy

{Marquardt}
   EqualizeDatasets = y

[end]
```

#### Chemical synthesis of SNAP-tag2 substrates

**General remarks.** All reagents were purchased from commercial suppliers (Acros Chemicals, Alfa Aesar, TCI Chemicals GmBH, ABCR, Sigma-Aldrich, Activate Scientific, Carl Roth GmBH + Co.KG, Merck KGaA, VWR International). All solvents used in reactions were anhydrous. Solvents were degassed, when necessary, by employing 3-times freeze-pump-thaw cycles or by purging $N_2$ for a minimum of 15 min. Anhydrous solvents were handled under argon atmosphere and reactions were performed in oven-dried glassware. Deionized water was used for all experiments. Reaction progress was followed by thin layer chromatography (TLC) or liquid chromatography-mass spectrometry (LCMS-2020 connected to a Nexera X-2 UHPLC system from Shimazu equipped with a C18 column (80 Å, 1.9 µm pore size, 2.1 x 50 mm by Supelco)). A solvent mixture gradient 10-95% MeCN/$H_2O$ with constant 0.1% v/v formic acid (1 mL/min) over a period of 6 min. was used. TLC was carried out on pre-coated silica gel plates (60G F254 on glass plates, Merck KGaA) in appropriate solvents followed by visualization of reaction spots using UV illumination (254 nm or 366 nm) and/or; by using the following staining agents (dip, dry & heat development). Staining solution: $KMnO_4$ (1 g), $K_2CO_3$ (6.6 g), 5% NaOH (1.7 mL) in $H_2O$ (90 mL). All synthetic products were either purified *via* normal phase flash column chromatography (FCC) using an automated system (Biotage Isolera One) with pre-packed silica gel columns (ultrapure silica gel 12 g or 25 g, SiliCycle Inc.) or *via* reverse-phase high-performance liquid chromatography (RP-HPLC) on a Thermo Fisher Scientific UltiMate 3000 system with a Supelco column (21.1 × 250 mm, 5 µm pore size, 8 mL/min flow rate) using a solvent gradient of 10-95% MeCN/$H_2O$ with constant 0.1% v/v trifluoroacetic acid additive. A standard RP-HPLC purification run took 45 min. Fractions containing the product were combined and concentrated *in vacuo* (rotary evaporator, Hei-Vap value, heidolph) with heating in a water bath (40 °C) and/or dried on a lyophilizer (Christ) connected to a vacuum pump (Vacuubrand). Final fluorophore substrates were stored as DMSO stocks at -20 °C.

Samples for nuclear magnetic resonance (NMR) spectroscopy were dissolved in deuterated solvents and NMR spectra were recorded at 298 K on a BRUKER Advance III HD 400 NMR spectrometer equipped with a CryoProbe™ ($^1$H: 400 MHz, $^{13}$C: 101 MHz). NMR spectra were analyzed using MestReNova 14.1.0 (MestreLab Research). Multiplicities are reported as s = singlet, d = doublet, t = triplet, q = quartet, m = multiplet and chemical shifts (δ) are calibrated to the residual chemical shifts of the solvents ($CDCl_3$, MeOD-$d_4$, MeCN-$d_3$, DMSO-$d_6$, DMF-$d_7$)[2]. Coupling constants J are reported in Hz. NMR spectra are reported as obtained. High-resolution mass spectrometry (HRMS) was acquired on a maXis II™ ETD-HRMS system (Bruker) using electron spray ionization (ESI) in positive mode conducted by the MS Core Facility.

#### Building blocks – nucleobases

##### *tert*-butyl (4,6-dichloropyrimidin-2-yl)carbamate (45):

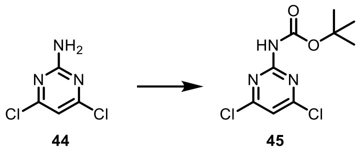

2-amino-4,6-dichloro pyrimidine **44** (4.2 g, 25.6 mmol, 1.0 equiv.) was dissolved in DCM (50 mL) and oxalyl chloride (9 mL) was added followed by the addition of 3 drops of DMF, at 25 °C. The reaction mixture was refluxed for 16 h. After removing volatiles, the residue was dissolved in DCM and an excess (10 mL) of *t*-BuOH was added. The reaction was refluxed for an additional 16 h. Volatiles were removed, and the residue dissolved in DCM and purified by FCC to give colorless tick oily product **45** (4.5 g, 66.5% yield).

**TLC:** *Rf* = 0.75 (EtOAc/*n*-Hexane = 2:8).

**$^1$H NMR** (400 MHz, DMSO-$d_6$) δ = 10.74 (s, 1H), 7.53 (s, 1H), 1.46 (s, 9H) ppm.

**$^{13}$C{$^1$H} NMR** (101 MHz, DMSO-$d_6$) *δ* = 161.2, 157.6, 150.0, 114.7, 80.5, 27.8 ppm.

**HRMS** (ESI$^+$) m/z: [M + Na]$^+$, calculated for $C_9H_{11}Cl_2N_3NaO_2^+$ 286.0121, found 286.0122.

##### *tert*-butyl (4-chloro-6-(methylthio)pyrimidin-2-yl)carbamate (46):

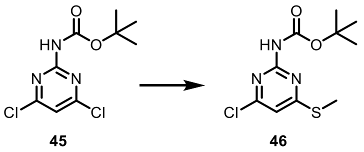

Carbamate **45** (4.5 g, 17 mmol) was dissolved in DMF (~100 mL) and solid sodium methanethiolate (1.2 g, 17 mmol, 1.0 equiv.), was added in one portion at 25 °C. The reaction mixture was stirred for 16 h after which DMF was removed under reduced pressure. Solid residue was dissolved in DCM (~200 mL) and washed with water (3 x ~50 mL). The organic fraction was dried with solid $Na_2SO_4$ and purified by FCC.

**TLC:** *Rf* = 0.38 (EtOAc/*n*-Hexane = 1:1).

**$^1$H NMR** (400 MHz, CDCl$_3$) *δ* = 7.36 (s, 1H), 6.86 (s, 1H), 2.59 (s, 3H), 1.55 (s, 9H) ppm.

**$^{13}$C{$^1$H} NMR** (101 MHz, CDCl$_3$) *δ* = 173.7, 159.6, 156.5, 149.7, 111.9, 81.9, 28.2, 12.7 ppm.

**HRMS** (ESI$^+$) m/z: [M + H]$^+$, calculated for $C_{10}H_{15}N_3O_2ClS^+$ 276.0568, found 276.0569.

***tert*-butyl (4-chloro-6-(methylsulfonyl)pyrimidin-2-yl)carbamate (47):**

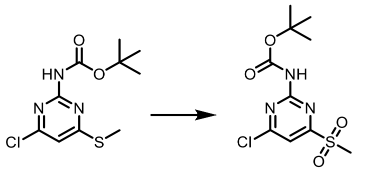

**45** **47**

Methylthiopyrimidine **46** (3.7 g, 13.6 mmol) was dissolved in DCM (~100 mL) and solid 3-chloroperoxybenzoic acid (16.8 g, 68.3 mmol, 5 equiv., 70% purity), was added in few small portions at 0 °C. Reaction mixture was stirred for 16 h letting it reach 25 °C. The reaction mixture was cooled to 0 °C and quenched by the addition of aqueous solution of $Na_2SO_3$. The organic fraction was separated and aqueous fraction was extracted with additional DCM (3 x ~150 mL). Combined organic fractions were dried with solid $Na_2SO_4$ and purified by FCC.

**$^1$H NMR** (400 MHz, DMSO-*d*$_6$) $\delta$ = 10.98 (s, 1H), 7.72 (s, 1H), 3.34 (s, 3H), 1.47 (s, 9H) ppm.

**$^{13}$C{$^1$H} NMR** (101 MHz, DMSO-*d*$_6$) $\delta$ = 167.2, 163.1, 158.1, 150.1, 110.7, 80.7, 27.8 ppm.

**HRMS** (ESI$^+$) m/z: [M + Na]$^+$, calculated for $C_{10}H_{14}ClN_3NaO_4S^+$ 330.0286, found 330.0288.

**4-chloro-6-(methylsulfonyl)pyrimidin-2-amine (49):**

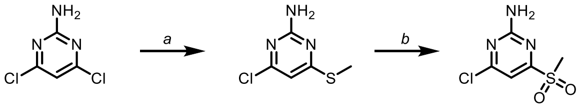

**44** **48** **49**

2-amino-4,6-dichloro pyrimidine **44** (2.0 g, 12.2 mmol, 1.5 equiv.) was dissolved in DMF (~100 mL) and solid sodium methanethiolate (0.57 g, 8.13 mmol, 1.0 equiv.), was added in one portion at 25 °C. The reaction mixture was stirred for 16 h after which DMF was removed under reduced pressure. Solid residue was dissolved in DCM (~200 mL) and washed with water (3 x ~50 mL). The organic fraction was dried with solid $Na_2SO_4$ and used in the next step without further purification.

Crude methylthiopyrimidine **48** was suspended in DCM/H2O (1:1, 50 mL) at 25 °C followed by addition of $Na_2WO_4$ (0.031 g, 0.096 mmol, 0.05 equiv.), $H_2O_2$ (0.9 mL, 7.7 mmol, 4 equiv., 30% in $H_2O$) and one drop of acetic acid. The reaction mixture was heated at 50 °C (oil bath) for 4 h. After cooling the reaction mixture to 0 °C, the remaining hydrogen peroxide was quenched by the addition of an aqueous solution of $Na_2SO_3$. The organic fraction was separated and aqueous fraction was extracted with additional DCM (3 x ~100 mL). Combined organic fractions were dried with solid $Na_2SO_4$ and purified by FCC.

**TLC:** *Rf* = 0.55 (EtOAc/*n*-Hexane = 1:1).

**$^1$H NMR** (400 MHz, DMSO-*d*$_6$) $\delta$ = 7.87 (s, 2H), 7.11 (s, 1H), 3.26 (s, 3H) ppm.

$^{13}$**C{$^1$H} NMR** (101 MHz, DMSO-*d$_6$*) *δ* = 167.5, 163.3, 162.7, 104.0, 39.3 ppm.

**HRMS** (ESI$^+$) m/z: [M + H]$^+$, calculated for $C_5H_7N_3O_2ClS^+$ 207.9942, found 207.9943.

**4-(methylsulfonyl)-6-(trifluoromethyl)pyrimidin-2-amine (52):**

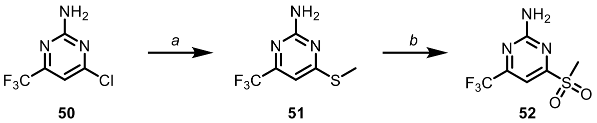


50 51 52

4-chloro-6-(trifluoromethyl)pyrimidin-2-amine **50** (2.0 g, 10.12 mmol, 1.0 equiv.) was dissolved in DMF (~100 mL) and solid sodium methanethiolate (0.85 g, 12.149 mmol, 1.2 equiv.), was added in one portion at 25 °C. The reaction mixture was stirred for 16 h after which DMF was removed under reduced pressure. Solid residue was dissolved in DCM (~200 mL) and washed with water (3 x ~50 mL). The organic fraction was dried with solid $Na_2SO_4$ and used in the next step without further purification after confirming conversion by LC/MS to methylthiopyrimidine **51**.

Crude methylthiopyrimidine **51** was suspended in DCM/H2O (1:1, 50 mL) at 25 °C followed by addition of $Na_2WO_4$ (0.031 g, 0.096 mmol, 0.05 equiv.), $H_2O_2$ (0.9 mL, 7.7 mmol, 4 equiv., 30% in $H_2O$) and one drop of acetic acid. The reaction mixture was heated at 50 °C (oil bath) for 4 h. After cooling the reaction mixture to 0 °C, the remaining hydrogen peroxide was quenched by the addition of aqueous solution of $Na_2SO_3$. The organic fraction was separated and aqueous fraction was extracted with additional DCM (3 x ~100 mL). Combined organic fractions were dried with solid $Na_2SO_4$ and purified by using a 25 g prepacked column (gradient 20-80% EtOAc/*n*-Hexane, over 30 column volumes) to give as white amorphous solid methylsulfon **52** (805 mg) in yield of 32% over 2 steps.

**TLC:** *Rf* = 0.65 (EtOAc/*n*-Hexane = 1:1).

$^1$**H NMR** $^1$H NMR (400 MHz, DMSO-*d$_6$*) *δ* = 8.10 (s, 2H), 7.34 (s, 1H), 3.32 (s, 3H) ppm.

$^{13}$**C{$^1$H} NMR** (101 MHz, DMSO-*d$_6$*) *δ* = 169.4, 164.1, 158.9, 158.5, 121.8, 119.1, 100.0, 39.6 ppm.

**19F{$^1$H}** (376 MHz, $CDCl_3$) *δ* = -69.4 ppm.

**HRMS** (ESI$^+$) m/z: [M + H]$^+$, calculated for $C_6H_6F_3N_3O_2S^+$ 242.0206 found 242.0205.

**4,6-dichloro-*N*[5]-methylpyrimidine-2,5-diamine (54):**

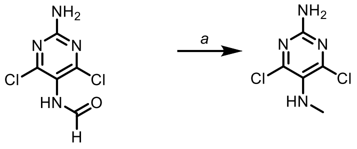


**53** **54**

For synthesis of **54** reduction of formamide, similar to previously described, was used.[3]

Formamide **53** (500 mg, 2.4 mmol, 1.0 equiv.) was dissolved in anhydrous THF (50 mL) at cooled to 0 °C. Borane-dimethyl sulfide (3 mL, 6 mmol, 2.5 equiv.) was added and the reaction mixture was left to reach 25 °C over 3 h. After the full conversion was confirmed by TLC and LC-MS, MeOH (~10 mL) was carefully added to quench the remaining borane-dimethyl sulfide complex and stirred for an additional 30 min. Volatiles were removed under reduced pressure, concentrated reaction mixture was dissolved in DCM (~100 mL) and washed with water (2x ~25 mL). Combined organic fractions were dried with solid $Na_2SO_4$ and purified by FCC to obtain white solid product **54** (264 mg, 56.6% yield).

**TLC:** *Rf* = 0.70 (EtOAc/*n*-Hexane = 3:7).

$^{1}$**H NMR** (400 MHz, $CDCl_3$) $\delta$ = 5.25 (s, 2H), 2.84 (s, 3H) ppm.

$^{13}$**C{$^{1}$H} NMR** (101 MHz, $CDCl_3$) $\delta$ = 156.8, 155.1, 129.8, 34.9 ppm.

**HRMS** (ESI$^+$) m/z: [M + H]$^+$, calculated for $C_5H_7N_4Cl_2$ 193.0042, found 193.0045.

**4,6-dichloropyrimidine-2,5-diamine (55):**

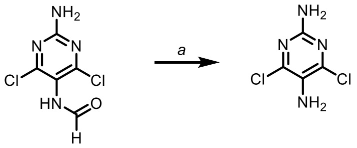


**53** **55**

Compound **55** was synthesized like previously described.[4]

**TLC:** *Rf* = 0.65 (EtOAc/*n*-Hexane = 3:7).

#### Building blocks – benzyl alcohol

***tert*-butyl (2-fluoro-4-(hydroxymethyl)benzyl)carbamate (58):**

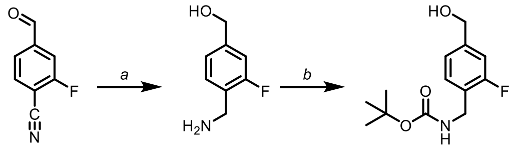


**56** **57** **58**

Nitrile **56** (3.0 g, 20.1 mmol, 1.0 equiv.) was dissolved and added to $LiAlH_4$ (3.8 g, 100.6 mmol, 5 equiv.), suspended in THF (~200 mL) at 0 °C. After stirring for 20 min at 0 °C, the solution was let to heat to 25 °C and then it was refluxed for 16 h. The reaction mixture was cooled to 0 °C, and the remaining $LiAlH_4$ was quenched by careful addition, under nitrogen atmosphere, of EtOAc (~50 mL), water, after which all volatiles were removed under reduced pressure to obtain crude amino-alcohol (**57**), which was used in the next step without further purification.

Crude amine **57** was dissolved in 1,4-Dioxane/Water (1:1, ~150 mL) and cooled down to 0 °C, to which $Et_3N$ (4.1 g, 40 mmol, 2 equiv.) and $Boc_2O$ (6.6 g, 30 mmol, 1.5 mmol) were added. The reaction mixture was let to reach 25 °C while stirring. Volatiles were removed under reduced pressure and the aqueous residue was extracted with EtOAc (2x ~250 mL), dried with solid $Na_2SO_4$ and purified by FCC to obtain a brownish solid product **58** (3.2 g, 62.3% yield).

**TLC:** *Rf* = 0.33 (EtOAc/*n*-Hexane = 3:7).

**$^1$H NMR** (400 MHz, $CDCl_3$) $\delta$ = 7.17 (s, 2H), 6.98 (s, 2H), 5.24 (s, 1H), 4.55 (s, 2H), 4.24 (d, *J*=6.1, 2H), 3.76 (s, 1H), 1.41 (s, 9H) ppm.

**$^{13}$C{$^1$H} NMR** (101 MHz, $CDCl_3$) $\delta$ = 162.0, 159.6, 156.2, 156.1, 143.1, 143.0, 140.1, 138.0, 129.5, 129.5, 127.4, 127.1, 124.6, 124.5, 122.2, 122.2, 122.1, 113.6, 113.4, 79.7, 64.5, 63.8, 63.7, 44.3, 38.3, 38.3, 28.3, 28.2 ppm.

**$^{19}$F NMR** (376 MHz, $CDCl_3$) δ = -119.3 ppm.

**HRMS** ($ESI^+$) m/z: $[M + Na]^+$, calculated for $C_{13}H_{18}NO_3FNa$ 278.1163, found 278.1161.

***tert*-butyl (3-fluoro-4-(hydroxymethyl)benzyl)carbamate (61):**

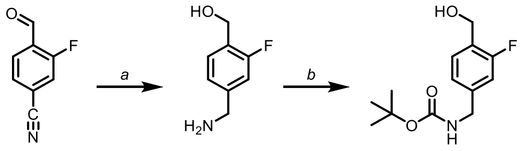


**59** **60** **61**

Carbamate (**61**) was synthesized as described above for **58**. A colorless oily liquid (0.4 g, 94% yield) was obtained, which solidified upon storage at 4 °C.

**TLC:** *Rf* = 0.70 (EtOAc/*n*-Hexane = 1:1).

Spectral data corresponds to data in previously published literature.[5]

***tert*-butyl (3-chloro-4-(hydroxymethyl)benzyl)carbamate (64):**

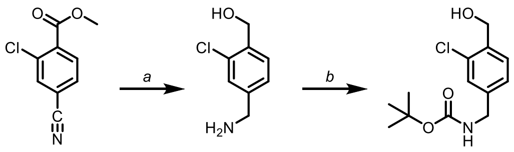


62 63 64

Carbamate (**64**) was synthesized as described above for **58**. Solid white product (202 mg, 58.2% yield) was obtained.

**TLC:** *Rf* = 0.77 (EtOAc/*n*-Hexane = 1:1).

Spectral data corresponds to data in previously published literature.[6]

***tert*-butyl (2-bromo-4-(hydroxymethyl)benzyl)carbamate (68):**

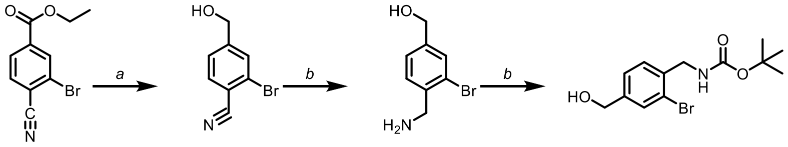


65 66 67 68

$LiBH_4$ (100 mg, 5.0 mmol, 4 equiv.) was suspended in diethyl ether (50 mL) followed by the addition of ester **65** (0.32 g, 1.26 mmol, 1.0 equiv.) and the mixture was cooled to 0 °C. MeOH (0.2 mL, 5.0 mmol, 4 equiv.) was added and the reaction mixture was stirred for 2 h, letting it reach room temperature. Unreacted $LiBH_4$ was quenched with water, diluted with EtOAc (~100 mL) and washed with water (2x ~20 mL). The organic fraction was dried with solid $Na_2SO_4$, volatiles were removed under reduced pressure and alcohol (**66**) product was used in the next step without further purification.

Alcohol (**66**) was dissolved in THF (~50 mL) and cooled to 0 °C. $BH_3 \cdot Me_2S$ (2.5 mL, 2M, 4 equiv.) was added dropwise and the reaction mixture was let to reach 25 °C over 16 h. The reaction mixture was cooled to 0 °C and unreacted borane was quenched by slow addition of MeOH (~5 mL) and water (~10 mL). The reaction mixture was diluted with DMC (~200 mL) and washed with water (3x ~50 mL). The organic fraction was dried with solid $Na_2SO_4$, volatiles were removed under reduced pressure and amino alcohol (**67**) was used in the next step without further purification.

Amino alcohol (**67**) was dissolved in a mixture of 1,4-dioxane and water (1:1, ~50 mL) and triethyl amine was added (0.254 g, 2.51 mmol, 2 equiv.), followed by the addition of $Boc_2O$ (0.41 g, 1.88 mmol, 1.5 equiv.). The reaction mixture was stirred over 16 h, diluted with EtOAc

(~150 mL) and washed with a saturated solution of $NaHCO_3$ (2x ~50 mL) and brine (2x ~50 mL). The organic fraction was dried with solid $Na_2SO_4$, volatiles was removed under reduced pressure and the product was purified by FCC to yield a yellowish thick liquid product **68** in yield of 57% over 3 consecutive steps (226 mg). Amorphous resin solidified during storage at 4 °C.

**TLC:** *Rf* = 0.73 (EtOAc/*n*-Hexane = 1:1).

**$^1$H NMR** (400 MHz, $CDCl_3$) $\delta$ = 7.43 (d, *J*=1.7, 1H), 7.33 – 6.92 (m, 2H), 5.14 (t, *J*=6.2, 1H), 4.50 (s, 2H), 4.22 (d, *J*=6.2, 2H), 1.35 (s, 9H) ppm.

**$^{13}$C NMR** (101 MHz, $CDCl_3$) $\delta$ = 156.0, 142.4, 136.7, 130.9, 129.5, 125.9, 123.4, 79.8, 63.7, 44.6, 28.4 ppm.

**HRMS** ($ESI^+$) m/z: $[M + Na]^+$, calculated for $C_{13}H_{18}NO_3BrNa^+$ 338.0362, found 338.0363.

#### Compound precursors

***tert*-butyl (4-(((4-amino-5-fluoropyrimidin-2-yl)oxy)methyl)benzyl)carbamate (71):**

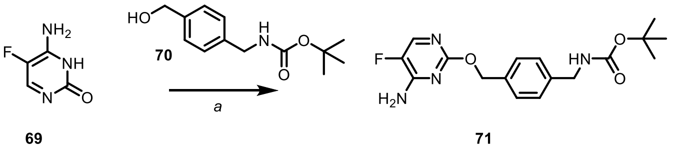

Benzyl alcohol **70** (200 mg, 0.84 mmol, 1.3 equiv.), $Ph_3P$ (170 mg, 0.65 mmol, 1.0 equiv.), and **69** (83 mg, 0.65 mmol, 1.0 equiv.) were dissolved in THF (~25 mL) and cooled down to 0 °C. DIAD (131 mg, 0.65 mmol, 1.0 equiv.) was added in one portion and the reaction mixture was let to reach 25 °C while stirring over 8 h. The Reaction mixture was diluted with EtOAc (~50 mL) and washed with a saturated solution of $NaHCO_3$ (~20 mL) and water (~20 mL). The organic fraction was dried with solid $Na_2SO_4$ and purified by FCC to obtain targeted *O*-alkyl product **71** (24 mg, 10.6% yield), after separation from the main side product (*N*-alkyl).

**TLC:** *Rf* = 0.48 (EtOAc/*n*-Hexane = 3:7).

**$^1$H NMR** (400 MHz, $CDCl_3$) $\delta$ = 7.93 (d, J=2.6, 1H), 7.49 – 7.36 (m, 2H), 7.29 (d, J=3.5, 3H), 5.41 – 5.25 (m, 2H), 4.33 (d, J=5.8, 2H), 1.48 (s, 9H) ppm.

**$^{13}$C{$^1$H} NMR** (101 MHz, $CDCl_3$) $\delta$ = 160.1, 155.9, 154.6, 154.4, 143.7, 141.3, 140.8, 140.6, 138.6, 135.8, 128.1, 127.5, 79.6, 68.9, 44.5, 28.4 ppm.

**$^{19}$F NMR** (376 MHz, $CDCl_3$) $\delta$ = -165.68 ppm.

**HRMS** ($ESI^+$) m/z: $[M + H]^+$, calculated for $C_{17}H_{22}FN_4O_3^+$ 349.1670, found 349.1674.

***tert*-butyl (4-(((2-amino-6-(trifluoromethyl)pyrimidin-4-yl)oxy)-methyl)-benzyl)carbamate (75):**

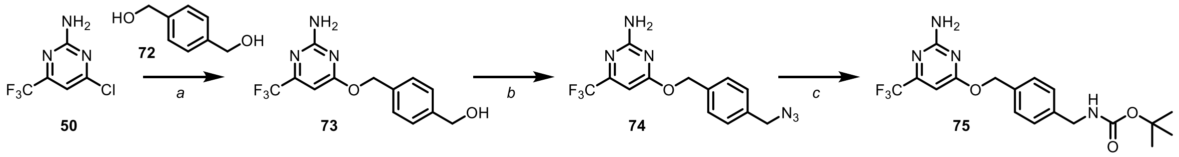

a) Bis-alcohol (**72**, 1.5 g, 11.1 mmol, 1.1 equiv.) was dissolved in a round bottom flask in a mixture of 1,4-dioxane and THF (4:1, 50 mL), cooled to 0 °C and solid NaH (0.48 g, 60%, 1.2 equiv.) was added. After stirring for 30 min at the same temperature under $N_2$ atmosphere, chloride (**50**) was added and the reaction mixture was heated to reflux (~100 °C, oil bath) and stirred for 16 h. After cooling down to 25 °C, the reaction was quenched with solid NH4Cl (~100 mg) and volatiles were removed under reduced pressure. The crude residue was dissolved in DCM (~200 mL) and washed with water (2 x ~50 mL) and the organic fraction was dried with solid anhydrous $Na_2SO_4$. After removing the volatiles under reduced pressure, alcohol (**75**) was used in the next step without further purification.

b) Alcohol (**75**) was dissolved in a mixture of toluene and THF (4:1, 50 mL) at room temperature (~25 °C) in the round bottom flask. To the reaction mixture, DPPA (5.5 g, 20.2 mmol, 2 equiv.) was added, followed by DBU (3.0 g, 20.2 mmol, 2 equiv.) and it was stirred under $N_2$ atmosphere for 16 h. The reaction mixture was diluted with EtOAc (~250 mL) and washed with a saturated solution of $NaHCO_3$ (3 x ~50 mL), the organic fraction was dried with solid anhydrous $Na_2SO_4$, volatiles removed under reduced pressure and azide (**74**) was used in the next step without further purification.

c) Crude azide (**74**) was dissolved in THF (50 mL) and tributylphosphine (2.7 g, 13.4 mmol, 1.33 equiv.) was added in one portion at room temperature while keeping the round bottom flask open to reduce the chance of overpressure in the flask caused by released gas. The reaction mixture was stirred for 2 h and water (3 mL) was added in one portion. After stirring for 16 h, NaOH (1M, 15 mL) was added at 0 °C followed by $Boc_2O$ (4.52 g, 20.7 mmol, 2.0 equiv.) and stirring was continued for another 12 h, letting the reaction mixture reach room temperature. The reaction mixture was extracted with EtOAc (3 x 100 mL), and organic fractions were combined and dried with solid anhydrous $Na_2SO_4$. After removing the volatiles under reduced pressure, automated FCC using 40 g prepacked column (gradient 10-50% EtOAc/*n*-Hexane, over 30 column volumes) provided 1.7 g of carbamate (**75**) as white amorphous solid (42% yield over 3 steps).

**$^1$H NMR** (400 MHz, $CDCl_3$) $\delta$ = 7.43 – 7.36 (m, 2H), 7.32 (d, *J*=7.9, 2H), 6.46 (s, 1H), 5.37 (s, 2H), 4.35 (d, *J*=6.0, 2H), 1.48 (s, 9H) ppm.

**$^{13}$C{$^1$H} NMR** (101 MHz, $CDCl_3$) $\delta$ = 171.0, 163.1, 157.2, 156.8, 155.9, 139.3, 134.8, 128.5, 127.7, 121.9, 119.2, 95.7, 95.6, 95.6, 95.6, 79.6, 68.2, 44.4, 28.4 ppm.

**$^{19}$F NMR** (376 MHz, $CDCl_3$) $\delta$ = -70.9 ppm.

**HRMS** (ESI$^+$) m/z: [M + H]$^+$, calculated for $C_{18}H_{22}F_3N_4O_3^+$ 399.1639, found 399.1637.

***tert*-butyl (4-(((2-amino-6-chloropyrimidin-4-yl)oxy)methyl)benzyl)carbamate (79):**

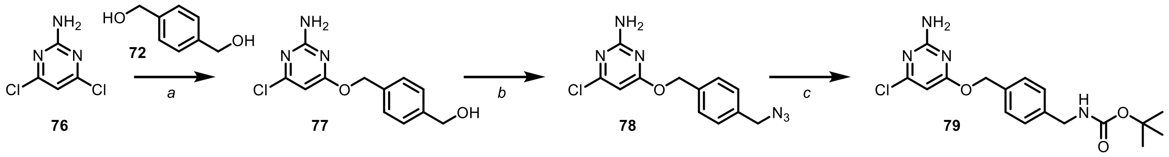


Carbamate (**79**) was synthesized following the procedure described for **75**, and a white amorphous solid was obtained as product (2.1 g, 45%, over 3 steps). For FCC, 40 g prepacked column was used, gradient 20% EtOAc in *n*-Hexane to 50% EtOAc, over 30 column volumes.

**$^1$H NMR** (400 MHz, $CDCl_3$) $\delta$ = 7.48 – 7.30 (m, 4H), 6.20 (s, 1H), 5.35 (s, 2H), 4.35 (d, *J*=5.6, 2H), 1.48 (s, 9H) ppm.

**$^{13}C\{^1H\}$ NMR** (101 MHz, $CDCl_3$) $\delta$ = 170.9, 162.2, 160.9, 155.9, 139.2, 135.1, 128.4, 127.7, 97.3, 79.6, 68.0, 44.4, 28.4 ppm.

Spectral data corresponds to data in previously published literature.[7]

***tert*-butyl (4-(((6-aminopyrazin-2-yl)oxy)methyl)benzyl)carbamate (83):**

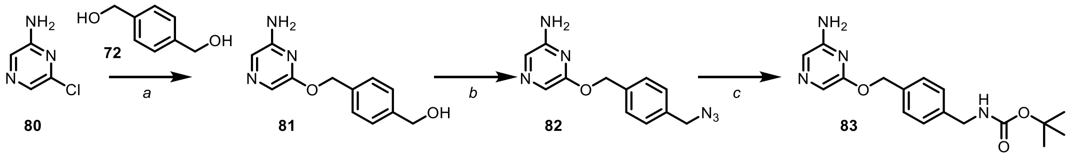


Carbamate (**83**) was synthesized following the procedure described for **75**, and a white amorphous solid was obtained as product (303 mg, 27%, over 3 steps). For FCC, 20 g prepacked column was used, gradient 20% EtOAc in *n*-Hexane to 100% EtOAc, over 20 column volumes.

**$^1H$ NMR** (400 MHz, $CDCl_3$) $\delta$ = 7.63 (s, 1H), 7.57 (s, 1H), 7.41 (d, *J*=8.2, 2H), 7.31 (d, *J*=7.9, 2H), 5.30 (s, 2H), 4.34 (d, *J*=5.7, 2H), 1.48 (s, 9H) ppm.

**$^{13}C\{^1H\}$ NMR** (101 MHz, $CDCl_3$) $\delta$ = 158.9, 155.9, 152.5, 138.9, 135.8, 128.3, 127.6, 122.8, 122.2, 79.6, 67.3, 44.4, 28.4 ppm.

**HRMS** ($ESI^+$) m/z: $[M + H]^+$, calculated for $C_{17}H_{23}N_4O_3^+$ 331.1765, found 331.1764.

***tert*-butyl (4-(((2-amino-6-fluoropyrimidin-4-yl)oxy)methyl)benzyl)carbamate (87):**

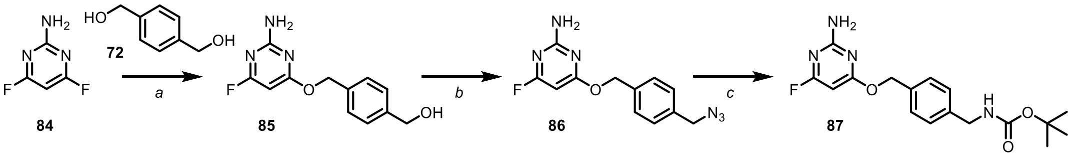


Carbamate (**87**) was synthesized following the procedure described for **75** and a white amorphous solid was obtained as product (25 mg, 22%, over 3 steps). For FCC, 20 g prepacked column was used, gradient 20% EtOAc in *n*-Hexane to 100% EtOAc, over 20 column volumes.

**TLC:** *Rf* = 0.45 (EtOAc/*n*-Hexane = 3:7).

**$^1H$ NMR** $^1H$ NMR (400 MHz, $CDCl_3$) $\delta$ = 7.28 (s, 2H), 7.23 (s, 2H), 5.59 (s, 1H), 5.24 (s, 2H), 4.24 (d, *J*=5.9, 2H), 1.39 (s, 9H) ppm.

**$^{13}C\{^1H\}$ NMR** (101 MHz, $CDCl_3$) $\delta$ = 173.2, 173.1, 173.0, 170.8, 162.4, 162.2, 155.9, 139.1, 135.2, 128.4, 127.7, 81.2, 80.9, 79.6, 68.3, 44.4, 28.4 ppm.

**$^{19}F$ NMR** (376 MHz, $CDCl_3$) $\delta$ = -64.2 ppm.

**HRMS** (ESI⁺) m/z: [M + H]⁺, calculated for $C_{17}H_{22}FN_4O_3^+$ 349.1670, found 349.1669.

***tert*-butyl (4-(((2-aminopyrimidin-4-yl)oxy)methyl)benzyl)carbamate (91):**

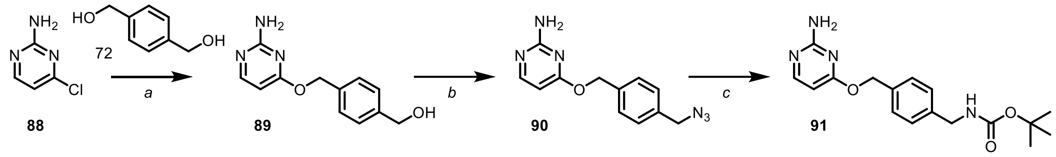


Carbamate (**91**) was synthesized following the procedure described for **75** and a white amorphous solid was obtained as product (65 mg, 20%, over 3 steps). For FCC, 20 g prepacked column was used, gradient 20% EtOAc in *n*-Hexane to 100% EtOAc, over 20 column volumes.

**TLC:** *Rf* = 0.52 (EtOAc/*n*-Hexane = 7:3).

**$^1$H NMR** (400 MHz, DMSO-$d_6$) $\delta$ = 8.15 (d, *J*=6.8, 1H), 7.44 (d, *J*=7.8, 3H), 7.27 (d, *J*=7.9, 2H), 6.41 (d, *J*=6.8, 1H), 5.41 (s, 2H), 4.13 (d, *J*=5.7, 2H), 1.39 (s, 9H) ppm.

**$^{13}$C{$^1$H} NMR** (101 MHz, DMSO-$d_6$) $\delta$ = 171.4, 159.4, 159.1, 157.9, 156.3, 149.1, 141.2, 133.9, 129.4, 129.3, 127.5, 99.1, 78.3, 69.1, 43.6, 43.5, 28.7 ppm.

**HRMS** (ESI⁺) m/z: [M + H]⁺, calculated for $C_{17}H_{23}N_3O_3^+$ 331.1765, found 331.1766.

***tert*-butyl (4-(((2-amino-6-bromo-pyrimidin-4-yl)oxy)methyl)benzyl)carbamate (95):**

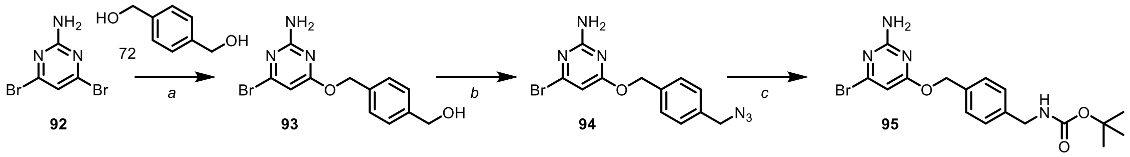


Carbamate (**95**) was synthesized following the procedure described for **75** and a yellowish amorphous solid was obtained as product (36 mg, 9.5%, over 3 steps). For FCC, 20 g prepacked column was used, gradient 20% EtOAc in *n*-Hexane to 100% EtOAc, over 20 column volumes.

**TLC:** *Rf* = 0.38 (EtOAc/*n*-Hexane = 3:7).

**$^1$H NMR** (400 MHz, $CDCl_3$) $\delta$ = 7.40 – 7.28 (m, 4H), 6.35 (s, 1H), 5.31 (s, 2H), 4.34 (d, *J*=5.9, 2H), 1.48 (s, 9H) ppm.

**$^{13}$C{$^1$H} NMR** (101 MHz, $CDCl_3$) $\delta$ = 170.3, 161.8, 155.9, 151.6, 139.2, 135.0, 128.4, 127.7, 101.4, 79.6, 68.0, 44.4, 28.4 ppm.

**HRMS** (ESI⁺) m/z: [M + H]⁺, calculated for $C_{17}H_{22}BrN_4O_3^+$ 409.0870, found 409.0868.

**tert-butyl (4-(((2-amino-6-methylpyrimidin-4-yl)oxy)methyl)benzyl)carbamate (99):**

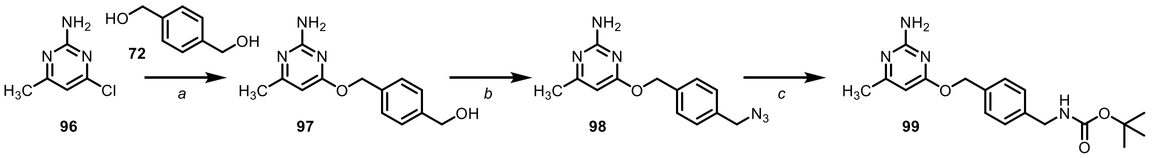


Carbamate (**99**) was synthesized following the procedure described for **75** and a yellowish amorphous solid was obtained as product (32 mg, 9.5%, over 3 steps). For FCC, 20 g prepacked column was used, gradient 20% EtOAc in *n*-Hexane to 100% EtOAc, over 20 column volumes.

**TLC:** *Rf* = 0.54 (EtOAc/*n*-Hexane = 7:3).

**$^{1}$H NMR** (400 MHz, $CDCl_3$:MeOD-*d*$_4$ = 4:1) δ = 7.32 – 7.12 (m, 5H), 5.94 (s, 1H), 5.28 (s, 2H), 4.16 (s, 2H), 2.28 (s, 3H), 1.32 (s, 9H) ppm.

**$^{13}$C{$^{1}$H} NMR** (101 MHz, $CDCl_3$:MeOD-*d*$_4$ = 4:1) $\delta$ = 172.2, 157.8, 156.8, 140.1, 133.7, 129.0, 127.7, 98.5, 69.8, 44.1, 28.4, 18.5 ppm.

**HRMS** ($ESI^+$) m/z: $[M + H]^+$, calculated for $C_{18}H_{25}N_4O_3^+$ 345.1921, found 345.1922.

**tert-butyl (4-(((6-aminopyrimidin-4-yl)oxy)methyl)benzyl)carbamate (103):**

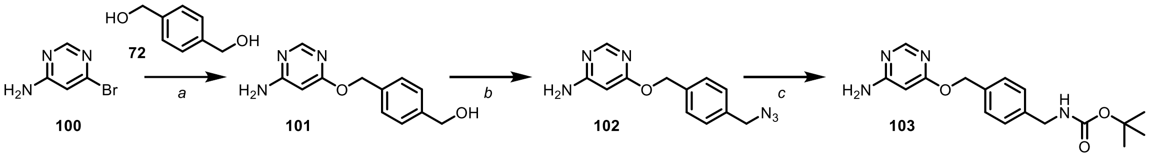


Carbamate (**103**) was synthesized following the procedure described for **75** and a yellowish amorphous solid was obtained as product (51 mg, 15.5%, over 3 steps). For FCC, 20 g prepacked column was used, gradient 20% EtOAc in *n*-Hexane to 100% EtOAc, over 20 column volumes.

**TLC:** *Rf* = 0.48 (EtOAc/*n*-Hexane = 7:3).

**$^{1}$H NMR** (400 MHz, $CDCl_3$:MeOD-*d*$_4$ = 4:1) $\delta$ = 8.18 (s, 1H), 7.21 (dd, *J*=29.1, 7.6, 4H), 5.91 (s, 1H), 5.29 (s, 2H), 4.17 (s, 2H), 1.33 (s, 9H) ppm.

**$^{13}$C{$^{1}$H} NMR** (101 MHz, $CDCl_3$:MeOD-*d*$_4$ = 4:1) $\delta$ = 169.9, 158.8, 156.4, 150.6, 139.6, 134.0, 128.6, 128.5, 127.5, 88.0, 79.7, 69.5, 44.0, 28.2 ppm.

**HRMS** ($ESI^+$) m/z: $[M + H]^+$, calculated for $C_{17}H_{23}N_4O_3^+$ 331.1765, found 331.1761.

**tert-butyl (4-(((2,5-diamino-6-chloropyrimidin-4-yl)oxy)methyl)benzyl)carbamate (106):**

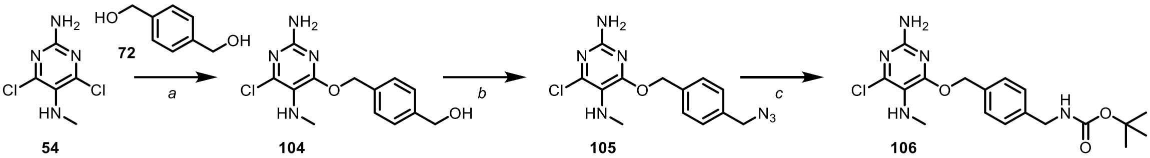


**54** **104** **105** **106**

Carbamate (**106**) was synthesized following the procedure described for **75** and a white amorphous solid was obtained as product (43 mg, 7.5%, over 3 steps). For FCC, 20 g prepacked column was used, gradient 20% EtOAc in *n*-Hexane to 100% EtOAc, over 20 column volumes.

**TLC:** *Rf* = 0.45 (EtOAc/*n*-Hexane = 3:7).

**$^1$H NMR** (400 MHz, $CDCl_3$) $\delta$ = 7.34 – 7.25 (m, 2H), 7.25 – 7.17 (m, 2H), 5.28 (s, 2H), 4.25 (d, *J*=6.0, 2H), 2.69 (s, 3H), 1.39 (s, 9H) ppm.

**$^{13}$C{$^1$H} NMR** (101 MHz, $CDCl_3$) $\delta$ = 163.4, 156.0, 155.9, 149.9, 139.2, 135.2, 128.5, 127.6, 120.3, 79.6, 68.5, 44.4, 34.8, 28.4 ppm.

**HRMS** (ESI$^+$) m/z: [M + H]$^+$, calculated for $C_{18}H_{25}N_5O_3Cl^+$ 394.1640, found 394.1647.

**tert-butyl (4-(((2-amino-6-chloro-5-(methylamino)pyrimidin-4-yl)oxy)methyl)benzyl)carbamate (109):**

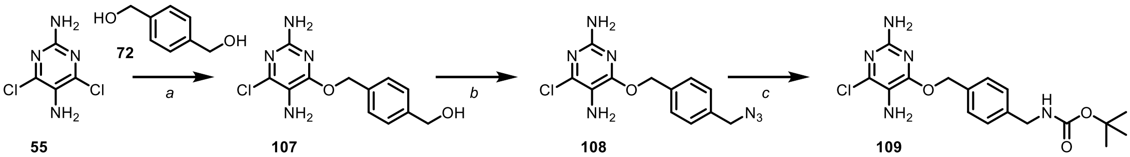


**55** **107** **108** **109**

Carbamate (**109**) was synthesized following the procedure described for **75** and a white amorphous solid was obtained as product (110 mg, 20%, over 3 steps). For FCC, 20 g prepacked column was used, gradient 20% EtOAc in *n*-Hexane to 100% EtOAc, over 20 column volumes.

**TLC:** *Rf* = 0.42 (EtOAc/*n*-Hexane = 3:7).

**$^1$H NMR** (400 MHz, DMF-*d*$_7$) $\delta$ = 7.68 (d, *J*=7.8, 2H), 7.52 (d, *J*=8.0, 3H), 5.57 (s, 2H), 4.46 (d, *J*=6.3, 2H), 1.61 (s, 9H) ppm.

**$^{13}$C{$^1$H} NMR** (101 MHz, DMF-*d*$_7$) $\delta$ = 159.9, 156.3, 155.3, 142.8, 140.6, 135.2, 128.5, 127.2, 116.1, 77.9, 67.9, 43.6, 27.9 ppm.

**HRMS** (ESI$^+$) m/z: [M + H]$^+$, calculated for $C_{17}H_{23}ClN_5O_3^+$ 380.1484, found 380.1482.

**_tert_-butyl (4-(((2-chloropyrrolo[2,1-f][1,2,4]triazin-4-yl)oxy)methyl)benzyl)carbamate (111):**

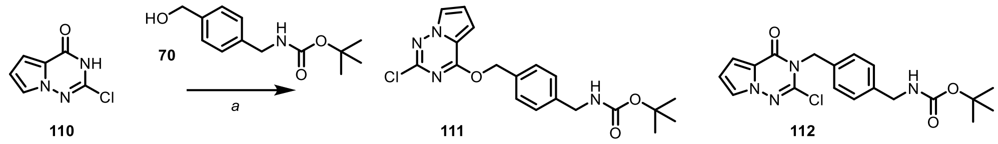


Benzyl alcohol **70** (703 mg, 2.96 mmol, 1.0 equiv.), $Ph_3P$ (778 mg, 2.96 mmol, 1.0 equiv.), and **110** (500 mg, 2.96 mmol, 1.0 equiv.) were dissolved in THF (~50 mL) and cooled down to 0 °C. DIAD (516 mg, 2.96 mmol, 1.0 equiv.) was added in one portion and the reaction mixture was let to reach 25 °C while stirring over 8 h. The reaction mixture was diluted with EtOAc (~150 mL) and washed with saturated solution of $NaHCO_3$ (~30 mL) and water (~30 mL). The organic fraction was dried with solid $Na_2SO_4$ and purified by FCC to obtain two products (*N*- and *O*-alkylated), both as while waxy resins, main (*O*-alkylated) product **111** (272 mg, 23.5% yield) and 301 mg of side (*N*-alkylated) product **112** in 26% yield.

**TLC:** *Rf* = 0.85 (EtOAc/*n*-Hexane = 3:7).

**$^1$H NMR** (400 MHz, $CDCl_3$) $\delta$ = 7.67 (dd, *J*=2.6, 1.5, 1H), 7.48 (d, *J*=8.1, 2H), 7.34 (d, *J*=8.1, 2H), 6.87 (dd, *J*=4.5, 1.5, 1H), 6.74 (dd, *J*=4.5, 2.6, 1H), 5.59 (s, 2H), 4.45 – 4.21 (m, 2H), 1.48 (s, 9H) ppm.

**$^{13}$C{$^1$H} NMR** (101 MHz, $CDCl_3$) $\delta$ = 161.6, 155.9, 151.0, 139.7, 134.0, 129.0, 127.7, 120.4, 113.4, 112.7, 103.5, 79.6, 69.0, 44.4, 28.4 ppm.

**HRMS** ($ESI^+$) m/z: $[M + H]^+$, calculated for $C_{19}H_{21}N_4O_3ClNa^+$ 411.1194, found 411.1195.

**_tert_-butyl (4-(((2-((tert-butoxycarbonyl)amino)pyrrolo[2,1-f][1,2,4]triazin-4-yl)oxy)-methyl)benzyl)-carbamate (113):**

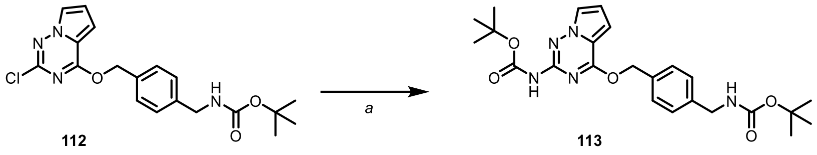


Chloride **112** (53 mg, 0.137 mmol, 1.0 equiv.) was dissolved in anhydrous 1,4-dioxane (~50 mL) followed by the addition of $Pd(OAc)_2$ (4.6 mg, 0.02 mmol, 0.15 equiv.), *tert*-Butyl carbamate (96 mg, 0.82 mmol, 6 equiv.), XPhos (29 mg, 0.062 mmol, 0.45 equiv.) and $CsCO_3$ (178 mg, 0.55 mmol, 4.0 equiv.). The reaction mixture was refluxed under an inert nitrogen atmosphere for 6 h, let to cool down to room temperature and diluted with EtOAc (~50 mL) and washed with a saturated solution of $NaHCO_3$, ammonium chloride and water (each ~10 mL). The organic fraction was dried with solid $Na_2SO_4$ and purified by FCC to provide 54 mg of carbamate **113** in 84.3% yield, as a white solid.

**TLC:** *Rf* = 0.43 (EtOAc/*n*-Hexane = 3:7).

**$^1$H NMR** (400 MHz, $CDCl_3$) $\delta$ = 7.61 (dd, *J*=2.6, 1.5, 1H), 7.34 (d, *J*=8.2, 2H), 7.23 (d, *J*=7.9, 2H), 6.89 (s, 1H), 6.68 (dd, *J*=4.5, 1.5, 1H), 6.53 (dd, *J*=4.4, 2.5, 1H), 5.41 (s, 2H), 4.85 (s, 1H), 4.25 (d, *J*=5.9, 2H), 1.47 (s, 9H), 1.38 (s, 9H) ppm.

**$^{13}$C{$^1$H} NMR** (101 MHz, $CDCl_3$) $\delta$ = 161.8, 156.0, 150.4, 149.2, 139.4, 134.6, 128.6, 127.7, 120.7, 113.0, 111.4, 102.7, 81.6, 79.6, 68.1, 44.4, 28.4, 28.2 ppm.

**HRMS** (ESI$^+$) m/z: [M + H]$^+$, calculated for $C_{24}H_{32}N_5O_5^+$ 470.2398, found 470.2396.

***tert*-butyl (4-(((2-amino-6-methoxypyrimidin-4-yl)oxy)methyl)benzyl)carbamate (114):**

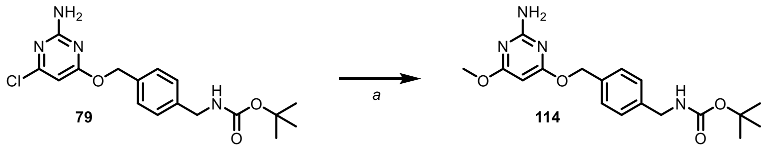


Chloride **79** (50 mg, 0.137 mmol, 1.0 equiv.) was dissolved in 1,4-dioxane (10 mL) and sodium methoxide was added (44 mg, 0.05 mL, 0.2 mmol, 1.5 equiv., 25% in MeOH) at 25 °C. The reaction was heated at reflux temperature for 16 h. The remaining unreacted methoxide was quenched with solid $NH_4Ac$ (~20 mg) and volatiles were removed under reduced pressure. The mixture was purified by automated FCC to provide 39 mg of methyl ether **114** in 79% yield, as a white solid.**TLC:** *Rf* = 0.22 (EtOAc/*n*-Hexane = 3:7).

**$^1$H NMR** $^1$H NMR (400 MHz, DMF-*d*$_7$) $\delta$ = 7.43 (d, *J*=7.9, 2H), 7.33 (d, *J*=7.8, 3H), 6.74 (s, 2H), 5.43 (s, 1H), 5.30 (s, 2H), 4.27 (d, *J*=6.3, 2H), 3.81 (s, 3H), 1.42 (s, 9H) ppm.

**$^{13}$C{$^1$H} NMR** $^{13}$C NMR (101 MHz, DMF-*d*$_7$) $\delta$ = 172.2, 171.6, 163.2, 156.3, 140.4, 135.9, 128.4, 127.2, 78.6, 77.9, 67.0, 53.0, 43.6, 27.9 ppm.

**HRMS** (ESI$^+$) m/z: [M + H]$^+$, calculated for $C_{24}H_{25}N_4O_4^+$ 361.1870, found 361.1870.

***tert*-butyl (4-(((2-amino-6-(dimethylamino)pyrimidin-4-yl)oxy)methyl)benzyl)carbamate (115):**

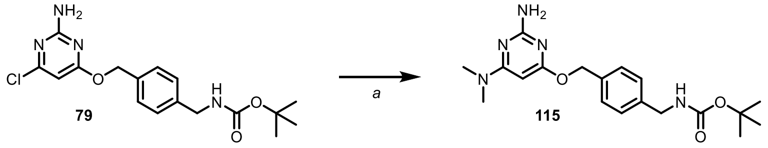


Chloride **79** (70 mg, 0.191 mmol, 1.0 equiv.) was dissolved in anhydrous 1,4-dioxane (~15 mL) followed by addition of $Pd(OAc)_2$ (6.4 mg, 0.028 mmol, 0.15 equiv.), dimethylamine hydrochloride (62 mg, 0.77 mmol, 4.0 equiv.), XPhos (41 mg, 0.086 mmol, 0.45 equiv.) and $CsCO_3$ (500 mg, 1.53 mmol, 8 equiv.). The reaction mixture was Heated at 110 °C under an inert nitrogen atmosphere for 4 h in a sealed vial, let to cool down to room temperature and diluted with EtOAc (~70 mL) and washed with a saturated solution of $NaHCO_3$, ammonium chloride and water (each ~15 mL). The organic fraction was dried with solid $Na_2SO_4$ and purified by FCC to provide 45 mg of dimethylamine-pyrimidine **115** in 84.3% yield, as a brownish solid.

**TLC:** *Rf* = 0.82 (EtOAc/*n*-Hexane = 7:3).

**$^1$H NMR** (400 MHz, $CDCl_3$) $\delta$ = 7.37 (s, 2H), 7.28 (s, 2H), 5.29 (s, 1H), 5.27 (s, 2H), 4.30 (s, 2H), 3.00 (s, 6H), 1.47 (s, 9H) ppm.

**$^{13}$C{$^1$H} NMR** (101 MHz, $CDCl_3$) $\delta$ = 170.8, 165.3, 162.1, 155.9, 138.5, 136.4, 128.2, 127.6, 79.5, 67.0, 44.4, 37.2, 28.4 ppm.

**HRMS** ($ESI^+$) m/z: $[M + H]^+$, calculated for $C_{19}H_{28}N_5O_3^+$ 374.2187, found 374.2183.

***tert*-butyl (4-(((2-amino-6-((tert-butoxycarbonyl)amino)pyrimidin-4-yl)oxy)methyl)benzyl)carbamate (116):**

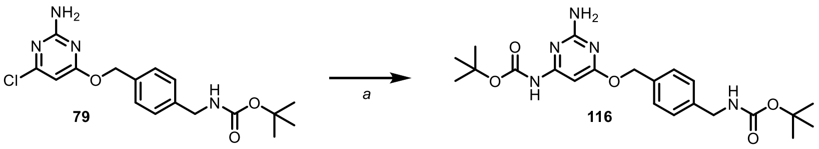


Chloride **79** (70 mg, 0.191 mmol, 1.0 equiv.) was dissolved in anhydrous 1,4-dioxane (~15 mL) followed by the addition of $Pd(OAc)_2$ (6.4 mg, 0.028 mmol, 0.15 equiv.), *tert*-butyl carbamate (134 mg, 1.15 mmol, 6 equiv.), XPhos (41 mg, 0.086 mmol, 0.45 equiv.) and $CsCO_3$ (150 mg, 0.767 mmol, 4 equiv.). The reaction mixture was Heated at 110 °C under an inert nitrogen atmosphere for 4 h in a sealed vial, let to cool down to room temperature and diluted with EtOAc (~70 mL) and washed with a saturated solution of $NaHCO_3$, ammonium chloride and water (each ~15 mL). The organic fraction was dried with solid $Na_2SO_4$ and purified by FCC to provide 60 mg of carbamate **116** in 80.7% yield, as a brownish solid.

**TLC:** *Rf* = 0.18 (EtOAc/*n*-Hexane = 3:7).

**$^1$H NMR** (400 MHz, $CDCl_3$) $\delta$ = 7.91 (d, *J*=40.7, 1H), 7.37 (s, 2H), 7.28 (s, 2H), 6.73 (s, 1H), 5.30 (s, 2H), 4.33 (d, *J*=5.9, 2H), 1.48 (s, 9H) ppm.

**$^{13}$C{$^1$H} NMR** (101 MHz, $CDCl_3$) $\delta$ = 171.7, 162.0, 159.6, 155.9, 152.0, 138.6, 135.9, 128.1, 127.6, 84.4, 81.8, 67.4, 44.4, 28.4, 28.2 ppm.

**HRMS** ($ESI^+$) m/z: $[M + H]^+$, calculated for $C_{22}H_{32}N_5O_5^+$ 466.2398, found 446.2395.

***tert*-butyl (4-(((4-amino-6-chloro-1,3,5-triazin-2-yl)oxy)methyl)benzyl)carbamate (119):**

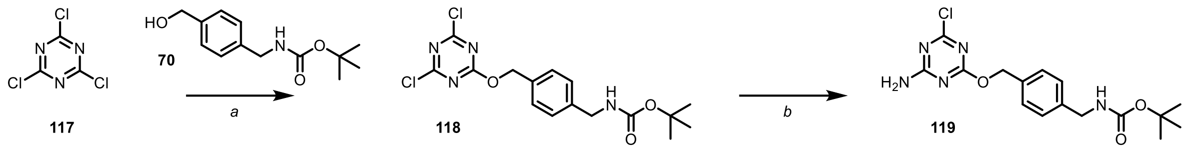

Triazine (**119**) was synthesized using the procedure described before for similar compounds.[8] Benzyl alcohol **70** (200 mg, 0.84 mmol, 1.0 equiv.) was dissolved in $CH_2Cl_2$ (~50 mL) followed by the addition of 2,6-lutidine (361 mg, 0.39 mL, 3.37 mmol, 4 equiv.) and cyanuric chloride **117** (155 mg, 0.84 mmol, 1.0 equiv.). The reaction mixture was stirred at reflux temperature under an inert $N_2$ atmosphere for 16 h, then diluted with additional DCM (~100 mL) and washed with water and brine (each ~30 mL). The organic fraction was dried with solid $Na_2SO_4$ and volatiles were removed under reduced pressure to give a crude intermediate (**118**) which was used in the next step without purification.

Crude bis-chloride triazine (**118**) was dissolved in DCM (~50 mL) and aqueous ammonium hydroxide (30%) was added (~15 mL). The biphasic reaction mixture was stirred for 3 h, diluted with additional DCM (~100 mL) and washed with water (2x ~50 mL). The organic fraction was dried with solid $Na_2SO_4$ and volatiles were removed under reduced pressure. Automated FCC gave white amino triazine (**119**) as a white solid (258 mg, 83.6% yield).

**TLC:** *Rf* = 0.18 (EtOAc/*n*-Hexane = 3:7).

**$^1$H NMR** (400 MHz, $CDCl_3$) $\delta$ = 7.15 (dd, *J*=8.3, 2.1, 2H), 7.09 – 6.97 (m, 2H), 5.14 (s, 2H), 4.03 (s, 2H), 1.22 (s, 9H) ppm.

**$^{13}$C{$^1$H} NMR** (101 MHz, $CDCl_3$) $\delta$ = 171.3, 170.9, 168.6, 157.2, 139.9, 134.5, 128.9, 127.7, 80.0, 69.9, 44.3, 28.6 ppm.

**HRMS** (ESI$^+$) m/z: [M + H]$^+$, calculated for $C_{16}H_{21}N_5O_3Cl^+$ 366.1327, found 366.1327.

***tert*-butyl (4-(((4-amino-1,3,5-triazin-2-yl)oxy)methyl)benzyl)carbamate (122):**

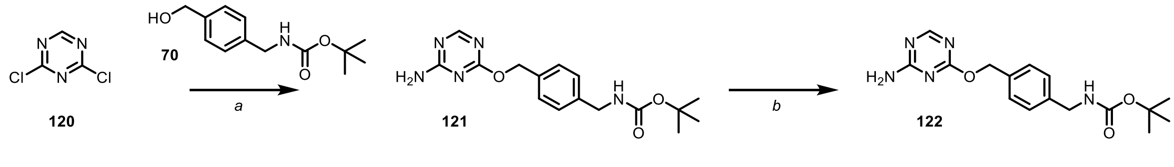

Triazine (**122**) was synthesized using the procedure described before for similar compounds.[8] Benzyl alcohol **70** (316 mg, 1.33 mmol, 1.0 equiv.) was dissolved in $CH_2Cl_2$ (~50 mL) followed by the addition of 2,6-lutidine (571 mg, 0.62 mL, 5.33 mmol, 4 equiv.) and triazine **120** (200 mg, 1.33 mmol, 1.0 equiv.). The reaction mixture was stirred at reflux temperature under an inert $N_2$ atmosphere for 16 h, then diluted with additional DCM (~100 mL) and washed with water and brine (each ~30 mL). The organic fraction was dried with solid $Na_2SO_4$ and volatiles were removed under reduced pressure to give a crude intermediate (**121**) which was used in the next step without purification.

Crude bis-chloride triazine (**118**) was dissolved in DCM (~50 mL) and aqueous ammonium hydroxide (30%) was added (~15 mL). The biphasic reaction mixture was stirred for 3 h, diluted with additional DCM (~100 mL) and washed with water (2x ~50 mL). The organic fraction was dried with solid $Na_2SO_4$ and volatiles were removed under reduced pressure. Automated FCC gave white amino triazine (**122**) as a white solid (70 mg, 15.8% yield).

**TLC:** *Rf* = 0.75 (EtOAc).

**$^1$H NMR** (400 MHz, $CDCl_3$:MeOD-$d_4$ = 4:1) $\delta$ = 8.06 (d, *J*=2.0, 1H), 7.24 – 7.13 (m, 2H), 7.07 (d, *J*=8.4, 2H), 5.16 (s, 2H), 4.06 (s, 2H), 1.24 (s, 9H) ppm.

**$^{13}$C{$^1$H} NMR** (101 MHz, $CDCl_3$: MeOD-$d_4$ = 4:1) $\delta$ = 169.7, 167.5, 167.2, 156.5, 139.1, 134.3, 128.1, 127.1, 79.4, 68.5, 43.7, 28.0 ppm.

**HRMS** (ESI$^+$) m/z: [M + H]$^+$, calculated for $C_{16}H_{22}N_5O_3^+$ 332.1717, found 332.1707.

***tert*-butyl (4-(((2-amino-6-(2,2,2-trifluoroethoxy)pyrimidin-4-yl)oxy)methyl)benzyl)carbamate (123):**

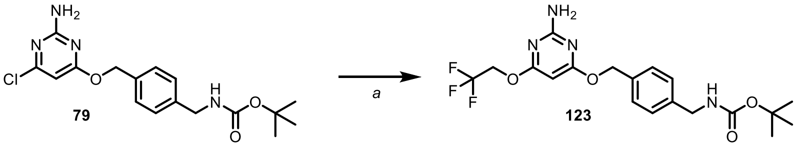


Trifluoroethanol (36 mg, 25 µL, 0.36 mmol, 1.3 equiv.) was added to anhydrous 1,4-dioxane (~10 mL) at 25 °C. Solid NaH (15 mg, 0.356 mmol, 1.3 equiv., 60%) was added and stirred at the same temperature for 30 min. Chloride **79** (100 mg, 0.274 mmol, 1.0 equiv.) was added and the reaction mixture was heated at 110 °C in a sealed vial over 16 h. After letting to cool to room temperature, unreacted alkoxide was quenched by the addition of solid $NH_4$Ac (~10 mg), volatiles were removed under reduced pressure and purified by automated FCC to give 110 mg of white solid product **123** (94% yield).

**$^1$H NMR** (400 MHz, $CDCl_3$) $\delta$ = 7.33 – 7.08 (m, 4H), 5.53 (s, 1H), 5.19 (d, *J*=11.6, 2H), 4.58 (d, *J*=8.6, 2H), 4.22 (d, *J*=5.8, 2H), 1.37 (s, 9H) ppm.

**$^{13}$C{$^1$H} NMR** (101 MHz, $CDCl_3$) $\delta$ = 175.0, 172.0, 172.0, 170.9, 170.1, 162.3, 161.8, 160.8, 156.0, 139.2, 138.9, 135.6, 135.0, 128.4, 128.3, 128.2, 127.7, 127.6, 127.5, 124.8, 122.0, 119.3, 97.2, 81.3, 79.6, 68.0, 67.8, 62.1, 61.8, 61.4, 44.4, 28.4 ppm.

**$^{19}$F NMR** (376 MHz, $CDCl_3$) $\delta$ = -73.9 ppm.

**HRMS** (ESI$^+$) m/z: [M + H]$^+$, calculated for $C_{19}H_{24}F_3N_4O_4^+$ 429.1744, found 429.1743.

**_tert_-butyl (4-(((2-amino-6-(methylthio)pyrimidin-4-yl)oxy)methyl)benzyl)carbamate (124):**

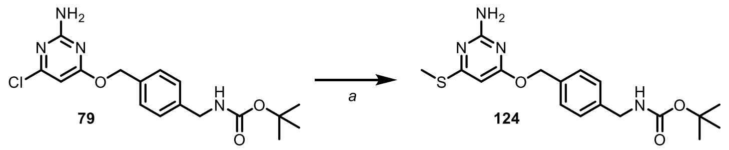


Chloride **79** (200 mg, 0.548 mmol, 1.0 equiv.) was dissolved in DMF (~25 mL) and sodium thiomethoxide (38 mg, 0.548 mmol, 1.0 equiv.) was added. The reaction mixture was warmed up to 50 °C and stirred at a constant temperature for 16 h. Volatiles were removed under reduced pressure and solid rest was purified by automated FCC to give 90 mg of yellowish solid product **124** (38% yield).

**TLC:** *Rf* = 0.42 (EtOAc/*n*-Hexane = 3:7).

**$^{1}$H NMR** (400 MHz, $CDCl_3$) *δ* = 7.35 (d, *J*=7.9, 2H), 7.29 (d, *J*=1.4, 2H), 5.99 (s, 1H), 5.28 (s, 2H), 4.31 (d, *J*=6.0, 2H), 2.43 (s, 3H), 1.47 (s, 9H) ppm.

**$^{13}$C{$^{1}$H} NMR** (101 MHz, $CDCl_3$) *δ* = 171.7, 169.6, 161.9, 156.0, 138.9, 135.6, 128.3, 127.6, 93.1, 79.5, 67.3, 44.4, 28.4, 12.9 ppm.

**HRMS** ($ESI^+$) m/z: $[M + H]^+$, calculated for $C_{18}H_{25}N_4O_3S^+$ 377.1642, found 377.1632.

**_tert_-butyl (4-(((2-((tert-butoxycarbonyl)amino)-6-chloropyrimidin-4-yl)oxy)methyl)-3-fluorobenzyl)-carbamate (125):**

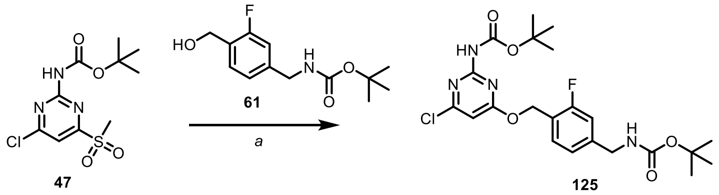


Benzyl alcohol **61** (24 mg, 0.09 mmol, 1.0 equiv.) was dissolved in THF (5 mL) and cooled down to 0 °C. Sodium hydride (4.3 mg, 0.126 mmol, 1.3 equiv.) was added and the suspension was stirred at the constant temperature for 30 min. Methyl sulfone **47** (30 mg, 0.09 mmol, 1.0 equiv.) was added in one portion and the reaction mixture was let to reach 25 °C over 16 h. The remaining unreacted hydride and alkoxide were quenched by the addition of solid $NH_4Ac$ (~10 mg), volatiles were removed under reduced pressure and the mixture was purified by automated FCC, 20 g prepacked column was used, gradient 10% EtOAc in *n*-Hexane to 50% EtOAc, over 20 column volumes. A solid white product was obtained (36 mg, 76% yield).

**TLC:** *Rf* = 0.68 (EtOAc/*n*-Hexane = 3:7).

**$^{1}$H NMR** (400 MHz, $CDCl_3$) *δ* = 7.49 (d, *J*=7.7, 1H), 7.40 (s, 1H), 7.11 – 6.96 (m, 2H), 6.45 (s, 1H), 5.47 (s, 2H), 4.33 (d, *J*=6.0, 2H), 1.56 (s, 9H), 1.48 (s, 9H) ppm.

**$^{13}C\{^1H\}$ NMR** (101 MHz, $CDCl_3$) $\delta$ = 170.6, 162.5, 161.2, 160.1, 156.7, 155.9, 149.8, 142.4, 142.4, 131.5, 131.4, 122.9, 121.7, 121.6, 114.4, 114.2, 101.8, 81.9, 79.9, 62.8, 62.8, 44.0, 28.4, 28.2 ppm.

**$^{19}F$ NMR** (376 MHz, $CDCl_3$) δ = -117.37 – -117.52 (m).

**HRMS** ($ESI^+$) m/z: $[M + H]^+$, calculated for $C_{22}H_{29}ClFN_4O_5^+$ 483.1805, found 483.1810.

***tert*-butyl (4-(((2-((tert-butoxycarbonyl)amino)-6-chloropyrimidin-4-yl)oxy)methyl)-3-chlorobenzyl)-carbamate (126):**

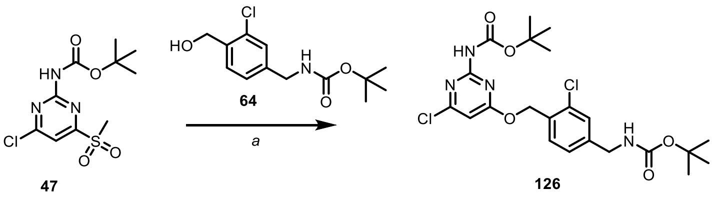


Carbamate **126** was synthesized following the procedure described above for **125** from alcohol **64** and methylsulfone **47**. White solid product (32 mg, 65% yield) was obtained after purification by automated FCC.

**TLC:** *Rf* = 0.64 (EtOAc/*n*-Hexane = 3:7).

**$^1H$ NMR** (400 MHz, $CDCl_3$) $\delta$ = 7.52 (d, *J*=7.9, 1H), 7.43 (s, 1H), 7.35 (d, *J*=1.7, 1H), 7.20 (d, *J*=1.8, 1H), 6.48 (s, 1H), 5.51 (s, 2H), 4.31 (s, 2H), 1.55 (s, 9H), 1.48 (s, 9H) ppm.

**$^{13}C\{^1H\}$ NMR** (101 MHz, $CDCl_3$) $\delta$ = 170.5, 161.3, 156.7, 155.9, 149.8, 141.3, 134.2, 132.2, 130.8, 128.3, 125.8, 101.8, 81.9, 79.9, 66.1, 43.8, 28.4, 28.2, 28.2 ppm.

**HRMS** ($ESI^+$) m/z: $[M + H]^+$, calculated for $C_{22}H_{29}Cl_2N_4O_5^+$ 499.1510, found 499.1505.

***tert*-butyl (3,5-dichloro-4-(hydroxymethyl)benzyl)carbamate (129):**

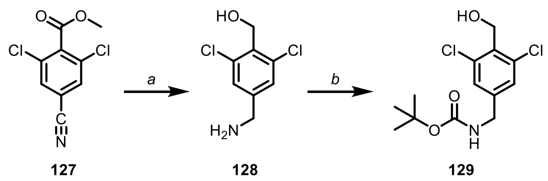


Carbamate (**129**) was synthesized as described above for **58**. Solid white product (178 mg, 53.5% yield) was obtained as an inseparable mixture of the carbamate **129** and a side product of hydrodechlorination. A mixture was used in the next step.

**TLC:** *Rf* = 0.85 (EtOAc/*n*-Hexane = 1:1).

**_tert_-butyl (4-(((2-((tert-butoxycarbonyl)amino)-6-chloropyrimidin-4-yl)oxy)methyl)-3,5-dichlorobenzyl)carbamate (130):**

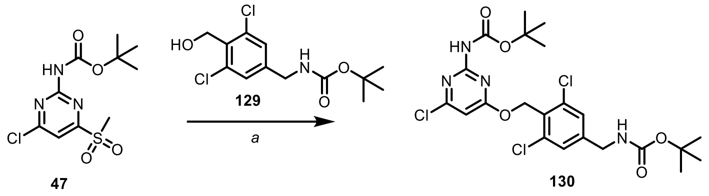

Carbamate **130** was synthesized following the procedure described above for **125** from alcohol **129** and methylsulfone **47**. White solid product (36 mg, 69% yield) was obtained after purification by automated FCC when the monochloro analogue, originating from the previous step mixture, was separated.

**TLC:** *Rf* = 0.75 (EtOAc/*n*-Hexane = 3:7).

**$^1$H NMR** (400 MHz, $CDCl_3$) $\delta$ = 7.39 (s, 1H), 7.30 (s, 2H), 6.43 (s, 1H), 5.63 (s, 2H), 5.00 (t, *J*=6.4, 1H), 4.31 (d, *J*=6.2, 2H), 1.56 (s, 9H), 1.49 (s, 9H) ppm.

**$^{13}$C{$^1$H} NMR** (101 MHz, $CDCl_3$) $\delta$ = 170.5, 161.2, 156.7, 155.8, 149.7, 142.9, 137.2, 129.8, 127.1, 101.7, 81.9, 80.2, 63.9, 43.5, 28.4, 28.2 ppm.

**HRMS** ($ESI^+$) m/z: $[M + H]^+$, calculated for $C_{22}H_{28}Cl_3N_4O_5^+$ 533.1120, found 533.1114.

**_tert_-butyl (3-chloro-5-fluoro-4-(hydroxymethyl)benzyl)carbamate (133):**

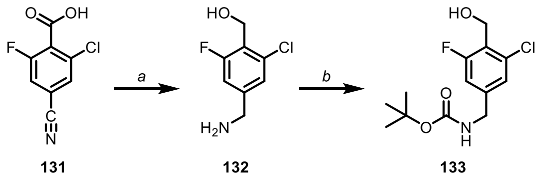

Carbamate (**133**) was synthesized as described above for **58**. Solid white product (46 mg, 35.2% yield) was obtained as a mixture of the title compounds and a minor product of hydrodechlorination, which was used in the next step without further purification.

**TLC:** *Rf* = 0.45 (EtOAc/*n*-Hexane = 3:7).

**HRMS** ($ESI^+$) m/z: $[M + Na]^+$, calculated for $C_{13}H_{17}NO_3ClFNa$ 312.0773, found 312.0772.

**tert-butyl (4-(((2-((tert-butoxycarbonyl)amino)-6-chloropyrimidin-4-yl)oxy)methyl)-3-chloro-5-fluorobenzyl)carbamate (134):**

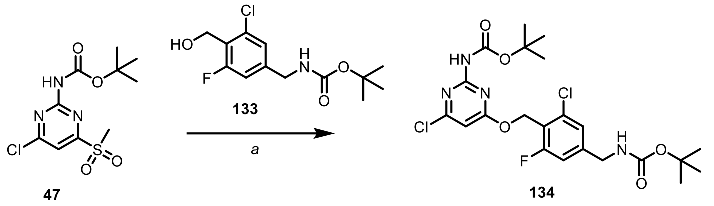


Carbamate **134** was synthesized following the procedure described above for **125** from alcohol **133** and methylsulfone **47**. White solid product (17 mg, 33% yield) was obtained after purification by automated FCC.

**TLC:** *Rf* = 0.79 (EtOAc/*n*-Hexane = 3:7).

**$^{1}$H NMR** (400 MHz, $CDCl_3$) *δ* = 7.31 (s, 1H), 7.17 (s, 1H), 6.98 (d, *J*=9.7, 1H), 6.41 (s, 1H), 5.51 (d, *J*=1.8, 2H), 4.30 (d, *J*=6.2, 2H), 1.54 (s, 9H), 1.47 (s, 9H) ppm.

**$^{13}$C{$^{1}$H} NMR** (101 MHz, $CDCl_3$) *δ* = 170.5, 163.4, 161.2, 156.7, 155.8, 149.7, 143.6, 143.5, 136.8, 136.7, 124.0, 120.2, 120.0, 113.1, 112.9, 101.7, 81.9, 80.2, 60.0, 60.0, 43.7, 28.4, 28.2 ppm.

**$^{19}$F NMR** (376 MHz, $CDCl_3$) δ = -112.34 (d, *J*=9.6) ppm.

**HRMS** ($ESI^+$) m/z: $[M + H]^+$, calculated for $C_{22}H_{28}Cl_2FN_4O_5^+$ 517.1415, found 517.1404.

**tert-butyl (4-(((2-((tert-butoxycarbonyl)amino)-6-chloropyrimidin-4-yl)oxy)methyl)-2-fluorobenzyl)carbamate (135):**

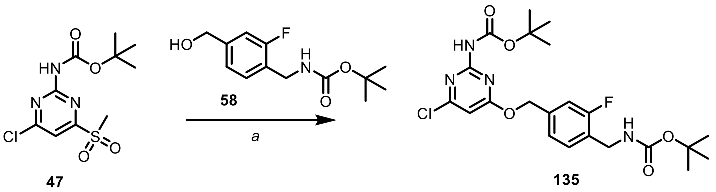


Carbamate **135** was synthesized following the procedure described above for **125** from alcohol **58** and methylsulfone **47**. White solid product (33 mg, 70% yield) was obtained after purification by automated FCC.

**TLC:** *Rf* = 0.82 (EtOAc/*n*-Hexane = 3:7).

**$^{1}$H NMR** (400 MHz, $CDCl_3$) *δ* = 7.38 – 7.21 (m, 3H), 7.13 (s, 2H), 6.37 (s, 1H), 5.33 (s, 2H), 4.28 (d, *J*=6.1, 2H), 1.47 (s, 9H), 1.37 (s, 9H) ppm.

$^{13}$C NMR (101 MHz, $CDCl_3$) *δ* = 170.5, 161.2, 156.7, 155.8, 149.7, 137.2, 137.1, 129.9, 129.0, 124.2, 115.7, 115.5, 101.8, 101.8, 81.9, 81.8, 79.7, 67.9, 67.9, 38.5, 28.4, 28.2 ppm.

**$^{19}$F NMR** (376 MHz, $CDCl_3$) *δ* = -118.8 ppm.

**HRMS** ($ESI^+$) m/z: $[M + Na]^+$, calculated for $C_{22}H_{28}ClFN_4NaO_5^+$ 505.1624, found 505.1613.

***tert*-butyl (4-(((2-((tert-butoxycarbonyl)amino)-6-chloropyrimidin-4-yl)oxy)methyl)-2-chlorobenzyl)carbamate (140):**

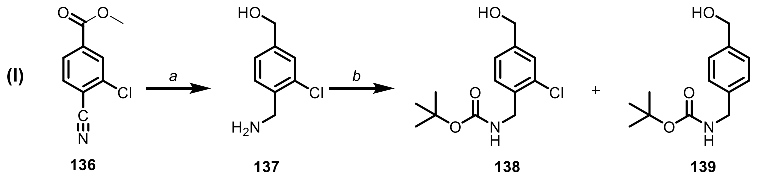


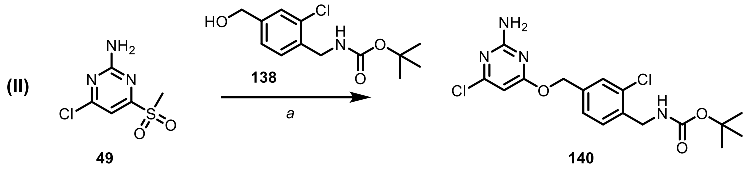


Alcohol (**138**) was synthesized as described above for **58**. Solid white product (241 mg) was obtained as a mixture of the target product and dehalogenated product, which were inseparable by FCC. A mixture was used in the next step.

**HRMS** ($ESI^+$) m/z: $[M + H]^+$, calculated for $C_{13}H_{18}ClNO_3Na^+$ 312.0773, found 312.0772.

Carbamate **140** was synthesized following the procedure described above for **135** from alcohol **138** (mixture with **139**) and methylsulfone **49**. A slightly yellowish solid product (19 mg, 43% yield) was obtained after purification by automated FCC.

**TLC:** *Rf* = 0.39 (EtOAc/*n*-Hexane = 3:7).

**$^1$H NMR** (400 MHz, $CDCl_3$) $\delta$ = 7.42 (d, *J*=1.7, 2H), 7.28 (s, 3H), 6.20 (s, 1H), 5.31 (s, 2H), 4.42 (d, *J*=6.2, 2H), 1.47 (s, 9H) ppm.

**$^{13}$C{$^1$H} NMR** (101 MHz, $CDCl_3$) $\delta$ = 170.7, 162.0, 161.0, 155.8, 137.0, 136.3, 133.6, 129.7, 128.9, 126.5, 97.4, 79.8, 67.1, 42.3, 28.4 ppm.

**HRMS** ($ESI^+$) m/z: $[M + H]^+$, calculated for $C_{17}H_{21}N_4O_3Cl_2^+$ 399.0985, found 399.0987.

***tert*-butyl (2,6-difluoro-4-(hydroxymethyl)benzyl)carbamate (143):**

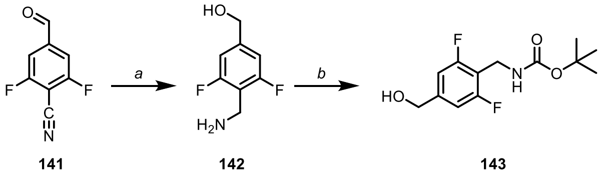


Carbamate (**143**) was synthesized as described above for **58**. A solid white product (89 mg, 28% yield) was obtained.

**TLC:** *Rf* = 0.6 (EtOAc/*n*-Hexane = 1:1).

**HRMS** (ESI$^+$) m/z: [M + Na]$^+$, calculated for $C_{13}H_{17}NO_3Na^+$ 296.1069, found 296.1069.

***tert*-butyl (4-(((2-((tert-butoxycarbonyl)amino)-6-chloropyrimidin-4-yl)oxy)methyl)-2,6-difluorobenzyl)carbamate (144):**

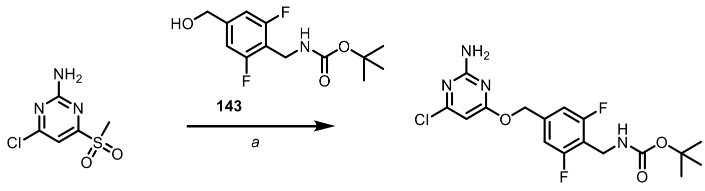


49 144

Carbamate **144** was synthesized following the procedure described above for **125** from alcohol **143** and methylsulfone **49**. White solid product (21 mg, 48% yield) was obtained after purification by automated FCC.

**TLC:** *Rf* = 0.38 (EtOAc/*n*-Hexane = 3:7).

$^1$**H NMR** (400 MHz, $CDCl_3$) $\delta$ = 7.27 (s, 1H), 7.10 – 6.77 (m, 2H), 6.12 (d, *J*=6.6, 1H), 5.23 (d, *J*=2.0, 2H), 4.29 (d, *J*=6.0, 2H), 1.38 (s, 9H) ppm.

$^{13}$**C{$^1$H} NMR** (101 MHz, $CDCl_3$) $\delta$ = 170.8, 170.5, 162.1, 161.8, 160.4, 155.8, 137.5, 137.4, 126.1, 125.9, 123.6, 114.9, 114.7, 110.4, 97.5, 97.4, 79.8, 67.4, 67.4, 66.7, 38.5, 29.7, 28.4 ppm.

$^{19}$**F NMR** (376 MHz, $CDCl_3$) $\delta$ = -114.26 (d, *J*=7.7), -118.67 (t, *J*=9.2) ppm.

**HRMS** (ESI$^+$) m/z: [M + H]$^+$, calculated for $C_{17}H_{20}N_4O_3ClF_2^+$ 401.1187, found 401.1191.

***tert*-butyl (2-bromo-4-(((2-((tert-butoxycarbonyl)amino)-6-chloropyrimidin-4-yl)oxy)-methyl)benzyl)-carbamate (145):**

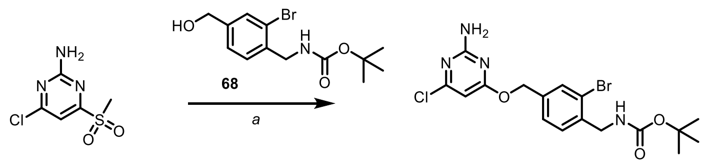


48 145

Carbamate **145** was synthesized following the procedure described above for **125** from alcohol **68** and methylsulfone **49**. White solid product (23 mg, 55% yield) was obtained after purification by automated FCC.

**TLC:** *Rf* = 0.38 (EtOAc/*n*-Hexane = 3:7).

$^1$**H NMR** (400 MHz, $CDCl_3$) $\delta$ = 7.44 – 7.29 (m, 4H), 6.18 (s, 1H), 5.32 (d, *J*=1.7, 2H), 4.34 (d, *J*=5.9, 2H), 1.48 (s, 10H) ppm.

$^{13}C\{^1H\}$ **NMR** (101 MHz, $CDCl_3$) $\delta$ = 170.9, 162.1, 160.8, 155.9, 139.2, 135.0, 128.4, 127.7, 97.4, 79.6, 68.1, 28.4 ppm.

**HRMS** ($ESI^+$) m/z: $[M + H]^+$, calculated for $C_{17}H_{21}N_4O_3BrCl^+$ 443.0480, found 443.0478.

***tert*-butyl (4-(((2-amino-6-(trifluoromethyl)pyrimidin-4-yl)oxy)methyl)-2-fluorobenzyl)carbamate (146):**

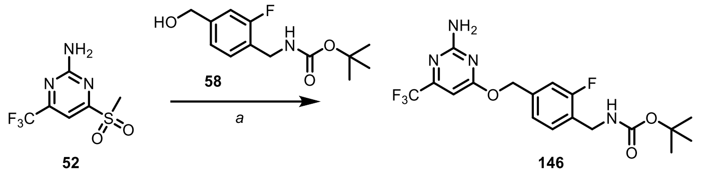


Carbamate **146** was synthesized following the procedure described above for **125** from alcohol **58** and methylsulfone **52**. White solid product (175 mg, 68% yield) was obtained after purification by automated FCC and additional purification by HPLC.

**TLC:** *Rf* = 0.48 (EtOAc/*n*-Hexane = 3:7).

$^1$**H NMR** (400 MHz, $CDCl_3$) δ = 7.36 (s, 1H), 7.21 – 7.00 (m, 2H), 6.44 (s, 1H), 5.35 (s, 2H), 4.37 (d, *J*=5.9, 2H), 1.46 (s, 9H) ppm.

$^{13}C\{^1H\}$ **NMR** (101 MHz, $CDCl_3$) δ = 170.7, 163.3, 163.2, 162.0, 159.6, 157.2, 156.8, 155.9, 137.4, 137.3, 130.0, 130.0, 128.5, 126.1, 126.0, 123.6, 123.6, 121.9, 119.2, 114.9, 114.7, 95.3, 95.3, 95.2, 95.2, 79.7, 68.1, 67.3, 67.3, 38.4, 28.4, 28.2 ppm.

$^{19}$**F NMR** (376 MHz, $CDCl_3$) $\delta$ = -76.55 (TFA), -124.42 – -124.48 (m) ppm.

**HRMS** ($ESI^+$) m/z: $[M + H]^+$, calculated for $C_{18}H_{21}N_4O_3F_4^+$ 417.1544, found 417.1542.

**methyl (tert-butoxycarbonyl)glycyl-*L*-serinate (147):**

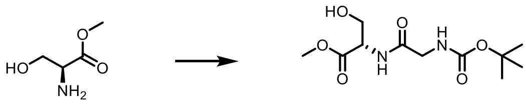


To a solution of *L*-serine methyl ester (1.47 g, 9 mmol, 1.0 equiv.) and (*tert*-butoxycarbonyl)glycine (1.89 g, 10.8 mmol, 1.2 equiv.) in anhydrous DCM/DMF (50:1) (50 mL) cooled down to 0 °C with ice-water bath were added 1-hydroxybenzotriazole hydrate (1.51 g, 9.9 mmol, 1.1 equiv.), 1-(3-dimethylaminopropyl)-3-ethylcarbodiimide hydrochloride (1.89 g, 9.9 mmol, 1.1 equiv.) and triethylamine (2.5 mL, 18 mmol, 2.0 equiv.). The reaction mixture was stirred 16 h. The mixture was poured into a phosphate buffer (50 mL) and diluted with DCM (150 mL). Fractions were separated and an aqueous layer was extracted 5 times with 250 mL of DCM. Organic fractions were combined, dried with $Na_2SO_4$ and concentrated under reduced pressure. FCC (10-30% EtOAc/n-Hexane, linear gradient) provided 1.57 g of the product in 63% yield.

Spectral data corresponds to data in previously published literature.[9]

**methyl (*S*)-2-(((*tert*-butoxycarbonyl)amino)methyl)-4,5-dihydrooxazole-4-carboxylate (148):**

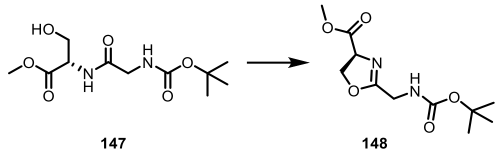

To a solution of **147** (1.56 g, 3.38 mmol, 1.0 equiv.) in anhydrous DCM at -78 °C was added dropwise diethylaminosulfur trifluoride (1.2 g, 7.45 mmol, 2.2 equiv.). The reaction mixture was slowly warmed up to 25 °C and stirred for a total of 3 hours. Then the mixture was cooled to -40°C and solid potassium carbonate (1.4 g, 10.163 mmol, 3.0 equiv.) was added. After stirring the reaction mixture at -40°C for 30 minutes, the mixture was allowed to warm up to 25 °C and it was stirred for an additional 15 minutes. The mixture was treated with saturated $NaHCO_3$ and diluted with DCM. Layers were separated and the aqueous layer was extracted 3 times with 150 mL of DCM. Organic fractions were combined, dried with $Na_2SO_4$ and concentrated under reduced pressure. The crude oxazoline was used directly in the next step.

**methyl 2-(((tert-butoxycarbonyl)amino)methyl)oxazole-4-carboxylate (149):**

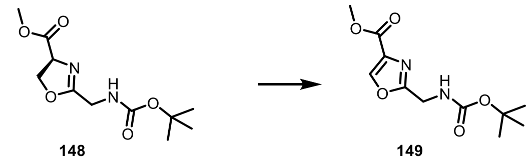

To a solution of **148** (875 mg, 3.38 mmol, 1.0 equiv.) in anhydrous DCM (30 mL) at -55°C was added dropwise solution of DBU (1.36 mL, 9.14 mmol, 2.7 equiv.) in anhydrous DCM (4 mL). The reaction mixture was stirred for 40 minutes at -55°C. Then solution of bromotrichloromethane (538 µL, 5.42 mmol, 1.6 equiv.) in anhydrous DCM (3 mL) was added. The reaction mixture was allowed to warm up to 25 °C and it was stirred for a total of 5 hours. The reaction mixture was cooled down to 0°C and was treated with phosphate buffer and it was diluted in DCM. Layers were separated and the aqueous layer was extracted 3 times with 150 mL of DCM. Organic fractions were combined, dried with $Na_2SO_4$ and concentrated under reduced pressure. Silica gel flash liquid chromatography (1-7% DCM/methanol, linear gradient) provided 835 mg of the product in 96% yield.

Spectral data corresponds to data in previously published literature.[10]

***tert*-butyl ((4-(hydroxymethyl)oxazol-2-yl)methyl)carbamate (150):**

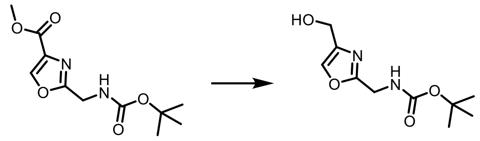


To a solution of **149** (835 mg, 3.26 mmol, 1.0 equiv.) in $Et_2O$ (40 mL) were added lithium borohydride (284 mg, 13.03 mmol, 4 equiv.) and methanol (528 µL, 13.03 mmol, 4 equiv.). The mixture was left refluxing overnight. The mixture was diluted with water and EtOAc. Layers were separated and the aqueous layer was extracted 3 times with 100 mL of EtOAc. Organic fractions were combined, washed with brine, dried with $Na_2SO_4$ and concentrated under reduced pressure. Silica gel flash liquid chromatography (10-40% EtOAc/n-Hexane, linear gradient) provided 200 mg of the product in 67% yield.

Spectral data corresponds to data in previously published literature.[11]

**tert-butyl ((4-(((2-amino-6-chloropyrimidin-4-yl)oxy)methyl)oxazol-2-yl)methyl)carbamate (151):**

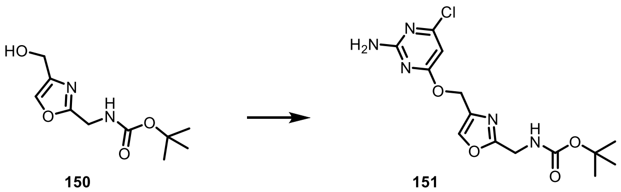


To a solution of **150** (220 mg, 0.964 mmol, 1.0 equiv.) in THF (20 mL) at 0°C was added sodium hydride (58 mg, 1.45 mmol, 1.5 equiv.). After stirring the reaction mixture for 30 minutes, **49** (231 mg, 0.964 mmol, 1.0 equiv.) was added. The reaction was stirred overnight. The mixture was diluted with water and EtOAc. Layers were separated and the aqueous layer was extracted 3 times with 150 mL of EtOAc. Organic fractions were combined, washed with brine, dried with $Na_2SO_4$ and concentrated under reduced pressure. Silica gel flash liquid chromatography (20-50% EtOAc/*n*-Hexane, linear gradient) provided 270 mg of the product in 79% yield.

**TLC:** *Rf* = 0.3 (EtOAc/*n*-Hexane = 1:1).

**$^1$H NMR** (400 MHz, DMSO-*d*$_6$) $\delta$ = 8.13 (s, 1H), 7.49 (s, 1H), 7.13 (s, 2H), 6.12 (s, 1H), 5.24 – 5.11 (m, 2H), 4.23 (d, *J*=6.0, 2H), 1.39 (s, 9H) ppm.

**$^{13}$C{$^1$H} NMR** (101 MHz, DMSO-*d*$_6$) $\delta$ = 170.5, 163.2, 162.4, 160.5, 156.0, 139.2, 135.7, 94.8, 78.8, 59.9, 37.8, 28.6 ppm.

**HRMS** (ESI$^+$) m/z: $[M + H]^+$, calculated for $C_{14}H_{19}ClN_5O_4^+$ 356.1120, found 356.1123.

**(4-bromothiophen-2-yl)methanol (152):**

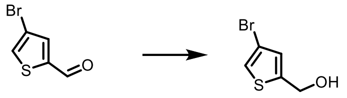

152

4-bromo-2-thiophenecarboxaldehyde (2.42 g, 12.67 mmol, 1.0 equiv.) was suspended in 100 mL of ethanol and cooled down to 0 °C. Sodium borohydride (958 mg, 25.33 mmol, 3.0 equiv.) was added to the mixture. The reaction was stirred for 2 hours. The consumption of the aldehyde was monitored by TLC. The reaction was quenched cautiously with 1.0 M aqueous HCl. Ethanol was removed under reduced pressure. The mixture was diluted with water and EtOAc. Layers were separated and the aqueous layer was extracted 2 times with 250 mL of EtOAc. Organic fractions were combined, dried with $Na_2SO_4$ and concentrated under reduced pressure. FCC (20-40% EtOAc/*n*-Hexane, linear gradient) provided 2.3 g of the product (94% yield).

Spectral data corresponds to data in previously published literature.[12]

**((4-bromothiophen-2-yl)methoxy)triisopropylsilane (153):**

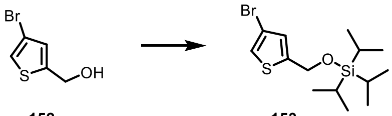

152 153

To a solution of **152** (2.3 g, 11.913 mmol, 1.0 equiv.) in DCM (70 mL) was added imidazole (2.43 g, 35.74 mmol, 3.0 equiv.) and chlorotriisopropylsilane (2.8 mL, 13.1 mmol, 1.1 equiv.). The reaction mixture was stirred overnight. The mixture was diluted in water and DCM. Layers were separated and the aqueous layer was extracted 3 times with 100 mL of DCM. Organic fractions were combined, dried with $Na_2SO_4$ and concentrated under reduced pressure. Silica gel flash liquid chromatography (1-5% EtOAc/n-Hexane, linear gradient) provided 3.32 g of the product in 80% yield.

Spectral data corresponds to data in previously published literature.[13]

**5-(((triisopropylsilyl)oxy)methyl)thiophene-3-carbaldehyde (154):**

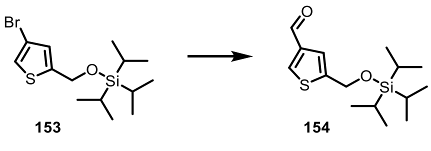

153 154

Alcohol **153** (3.32 g, 9.51 mmol, 1.0 equiv.) was dissolved in anhydrous THF (70 mL). Solution was cooled down to -78 °C and *n*-butyllithium (2.7 M, 4.22 mL, 1.2 equiv.) was added dropwise.

The mixture was stirred for 30 min. DMF (1.17 mL, 15.22 mmol, 1.6 equiv.) was added. The mixture was allowed to warm up to 25 °C and was stirred for 3 hours. The reaction mixture was quenched with a saturated solution of ammonia chloride and diluted with EtOAc. Layers were separated and the aqueous layer was extracted 3 times with 250 mL of EtOAc. Organic fractions were combined, dried with $Na_2SO_4$ and concentrated under reduced pressure. The crude aldehyde was used directly in the next step.

**(5-(((triisopropylsilyl)oxy)methyl)thiophen-3-yl)methanol (155):**

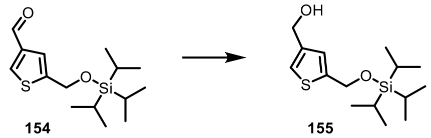


To a solution of **154** (2.93 g, 9.82 mmol, 1.0 equiv.) in ethanol (100 mL) at 0 °C was added sodium borohydride (1.11 g, 29.44 mmol, 3.0 equiv.). The reaction was stirred for 2 hours. The consumption of the aldehyde was monitored by TLC. The reaction was quenched cautiously with 1.0 M aqueous HCl. Ethanol was removed under reduced pressure. The mixture was diluted with water and EtOAc. Layers were separated and the aqueous layer was extracted 2 times with 250 mL of EtOAc. Organic fractions were combined, dried with $Na_2SO_4$ and concentrated under reduced pressure. The crude hydroxide was used crude in the next step.

**4-chloro-6-((5-(((triisopropylsilyl)oxy)methyl)thiophen-3-yl)methoxy)pyrimidin-2-amine (156):**

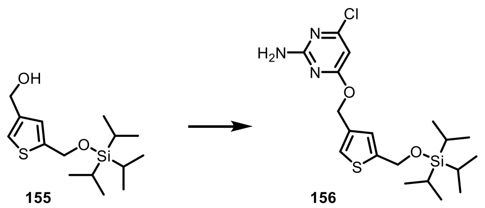


To a solution of **155** (500 mg, 1.66 mmol, 1.0 equiv.) in THF (20 mL) at 0°C was added sodium hydride (99.8 mg, 2.5 mmol, 1.5 equiv.). After stirring the reaction mixture for 30 minutes, methyl sulfone **49** (398 mg, 1.66 mmol, 1.0 equiv.) was added. The reaction was stirred overnight. The mixture was diluted with water and EtOAc. Layers were separated and aqueous fraction was extracted 3 times with 250 mL of EtOAc. Organic fractions were combined, washed with brine, dried with $Na_2SO_4$ and concentrated under reduced pressure. The crude mixture was used directly in the next step.

**(4-(((2-amino-6-chloropyrimidin-4-yl)oxy)methyl)thiophen-2-yl)methanol (157):**

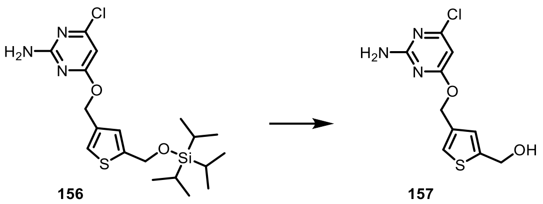


To a solution of **156** (710 mg, 1.66 mmol, 1.0 equiv.) in THF (30 mL) was added TBAF (1 M, 4.97 mL, 3.0 equiv.). The reaction mixture was stirred overnight. The mixture was diluted with water and EtOAc. Layers were separated and the aqueous layer was extracted 3 times with 250 mL of EtOAc. Organic fractions were combined, washed with brine, dried with $Na_2SO_4$ and concentrated under reduced pressure. Silica gel flash liquid chromatography (20-60% EtOAc/n-Hexane, linear gradient) provided 380 mg of the product in 84.3% yield.

Rf = 0.33 (EtOAc/*n*-Hexane = 1:1).

**$^1$H NMR** (400 MHz, DMSO-$d_6$) $\delta$ = 7.47 (d, *J*=1.4, 1H), 7.11 (s, 2H), 7.00 (d, *J*=1.3, 1H), 6.12 (s, 1H), 5.45 (s, 1H), 5.24 (s, 2H), 4.59 (d, *J*=3.2, 2H) ppm.

**$^{13}$C{$^1$H} NMR** (101 MHz, DMSO-$d_6$) δ = 170.7, 163.2, 160.4, 147.5, 136.8, 125.3, 124.6, 94.9, 63.3, 58.7 ppm.

**HRMS** (ESI$^+$) m/z: [M + H]$^+$, calculated for $C_{10}H_{11}ClN_3O_2S^+$ 272.0255, found 272.0260.

**4-((5-(azidomethyl)thiophen-3-yl)methoxy)-6-chloropyrimidin-2-amine (158):**

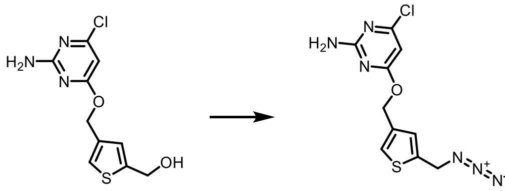


To a solution of **157** (380 mg, 1.40 mmol, 1.0 equiv.) in THF/toluene (1:1, 30 mL) at 0°C was added diphenylphosphoryl azide (453 µL, 2.1 mmol, 1.5 equiv.). After stirring the reaction mixture for 10 minutes, DBU (627 µL, 4.19 mmol, 3.0 equiv.) was added. The reaction was stirred overnight. The mixture was diluted with water and EtOAc. Layers were separated and the aqueous layer was extracted 3 times with 150 mL of EtOAc. Organic fractions were combined, washed with brine, dried with $Na_2SO_4$ and concentrated under reduced pressure. The crude mixture was used directly in the next step.

**_tert_-butyl ((4-(((2-amino-6-chloropyrimidin-4-yl)oxy)methyl)thiophen-2-yl)methyl)carbamate (159):**

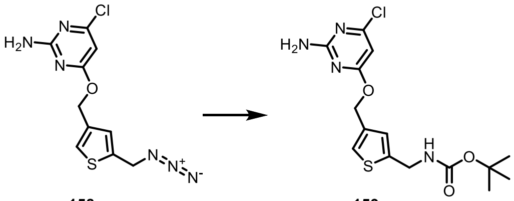

**158** **159**

To a solution of **158** (410 mg, 1.38 mmol, 1.0 equiv.) in THF (15 mL) was added tributylphosphine (682 µL, 2.76 mmol, 2.0 equiv.). After 4 hours water (10 mL) was added. The reaction mixture was stirred overnight. To the reaction mixture was added 1 M NaOH (20 mL) and the mixture was cooled down to 0°C. Di-tert-butyl decarbonate (603 mg, 2.76 mmol, 2.0 equiv.) was added. The reaction mixture was stirred overnight. The mixture was diluted with water and EtOAc. Layers were separated and the aqueous layer was extracted 3 times with 250 mL of EtOAc. Organic fractions were combined, washed with brine, dried with $Na_2SO_4$ and concentrated under reduced pressure. Silica gel flash liquid chromatography (20-60% EtOAc/n-Hexane, linear gradient) provided 380 mg of the product in 84.3% yield.

**TLC:** *Rf* = 0.35 (EtOAc/*n*-Hexane = 3:7).

**HRMS** ($ESI^+$) m/z: $[M + H]^+$, calculated for $C_{15}H_{20}ClN_4O_3S^+$ 371.0939, found 371.0940.

**(6-(bromomethyl)pyridin-2-yl)methanol (160):**

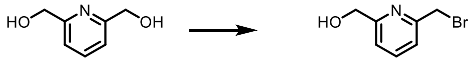

**160**

To a solution of 2,6-pyridinedimethanol (500 mg, 3.59 mmol, 1.0 equiv.) in dicholoromethane (30 mL) was added carbon tetrabromide (1.19 g, 3.59 mmol, 1.0 equiv.) and triphenylphosphine (942 mg, 3.59 mmol, 1.0 equiv.). The reaction was stirred at ambient temperature overnight. The reaction mixture was poured into a saturated solution of sodium bicarbonate and diluted with DCM. Layers were separated and an aqueous layer was extracted with 100 mL of DCM. Organic fractions were combined, dried with $Na_2SO_4$ and concentrated under reduced pressure. Silica gel flash liquid chromatography (20-60% EtOAc/n-Hexane, linear gradient) provided 300 mg of the product in 41.3% yield.

Spectral data corresponds to data in previously published literature.[14]

**2-((6-(hydroxymethyl)pyridin-2-yl)methyl)isoindoline-1,3-dione (161):**

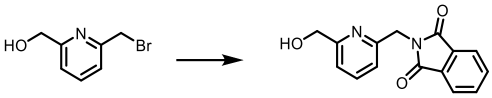


To a stirred solution of potassium phthalimide (268 mg, 1.45 mmol, 1.0 equiv.) at 25 °C under argon in DMF (20 mL) was added a DMF solution (10 mL) of **160** (146 mg, 723 µmol, 1.0 equiv.) over 5 min. The reaction mixture was warmed at 40°C and stirred overnight. The reaction mixture was cooled to 25 °C and diluted with DCM, washed with water 4 times. The organic layer was dried with $Na_2SO_4$ and concentrated under reduced pressure. The crude mixture was used directly in the next step.

Spectral data corresponds to data in previously published literature.[15]

**(6-(aminomethyl)pyridin-2-yl)methanol (162):**

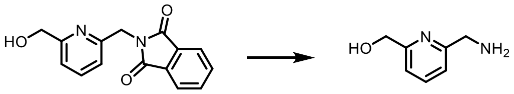


To a solution of **161** (236 mg, 880 µmol, 1.0 equiv.) in ethanol (30 mL) was added hydrazine hydrate (28.2 mg, 880 µmol, 1.0 equiv.). The reaction mixture was refluxed overnight. The mixture was cooled to 25 °C and filtered from precipitate. The filtrate was evaporated to obtain crude primary amine which was used in the next step.

Spectral data corresponds to data in previously published literature.[15]

**tert-butyl ((6-(hydroxymethyl)pyridin-2-yl)methyl)carbamate (163):**

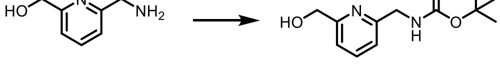


To a solution of **162** (120 mg, 0.868 mmol, 1.0 equiv.) in DCM (30 mL) at 0°C was added di-tert-butyl carbonate (228 mg, 1.04 mmol, 1.2 equiv.). The reaction was stirred overnight. The reaction mixture was concentrated under reduced pressure. Silica gel flash liquid chromatography (40-70% EtOAc/n-Hexane, linear gradient) provided 30 mg of the product in 14.5% yield.

Spectral data corresponds to data in previously published literature.[15]

**tert-butyl ((6-(((2-amino-6-chloropyrimidin-4-yl)oxy)methyl)pyridin-2-yl)methyl)carbamate (164):**

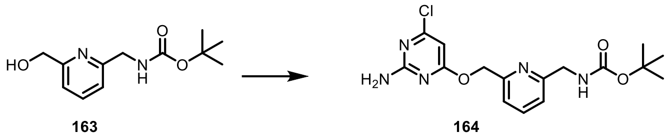

To a solution of **163** (30 mg, 0.126 mmol, 1.0 equiv.) in THF (10 mL) at 0°C was added sodium hydride (7.5 mg, 0.189 mmol, 1.5 equiv.). After stirring the reaction mixture for 30 minutes, **49** (30.2 mg, 0.126 mmol, 1.0 equiv.) was added. The reaction was stirred overnight. The mixture was diluted with water and EtOAc. Layers were separated and the aqueous layer was extracted 3 times with 50 mL of EtOAc. Organic fractions were combined, washed with brine, dried with $Na_2SO_4$ and concentrated under reduced pressure. Silica gel flash liquid chromatography (20-50% EtOAc/n-Hexane, linear gradient) provided 44 mg of the product in 95.5% yield.

**TLC:** *Rf* = 0.30 (EtOAc/*n*-Hexane = 1:1).

**$^1$H NMR** (400 MHz, DMSO-*d6*) $\delta$ = 7.90 (t, *J*=7.8, 1H), 7.55 (t, *J*=6.2, 1H), 7.38 (d, *J*=7.6, 1H), 7.30 (d, *J*=7.8, 1H), 7.21 (s, 2H), 6.32 (s, 1H), 5.47 (s, 2H), 4.31 (d, *J*=6.2, 2H), 1.50 (s, 9H) ppm.

**$^{13}$C{$^1$H} NMR** (101 MHz, DMSO-*d6*) $\delta$ = 170.6, 163.3, 160.6, 159.7, 156.4, 155.6, 138.1, 120.2, 120.0, 94.9, 78.5, 68.3, 45.8, 28.7 ppm.

**HRMS** (ESI$^+$) m/z: [M + H]$^+$, calculated for $C_{16}H_{21}ClN_5O_3^+$ 366.1327, found 366.1330.

**tert-butyl (2-amino-2-oxoethyl)carbamate (165):**

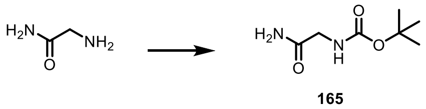

Glycinamide hydrochloride (1 g, 9.05 mmol, 1.0 equiv.) was dissolved in THF/water 4:1. Triethylamine (1.26 mL, 9.05 mmol, 1.0 equiv.) and di-tert-butyl dicarbonate (2.32 mL, 10.9 mmol, 1.2 equiv.) were added to the solution. The reaction was stirred overnight. The mixture was acidified with saturated $NaHSO_4$, and diluted with water and EtOAc. Layers were separated and the aqueous layer was extracted 3 times with 250 mL of EtOAc. Organic fractions were combined, dried with $Na_2SO_4$ and concentrated under reduced pressure. Silica gel flash liquid chromatography (40-80% EtOAc/n-Hexane, linear gradient) provided 1.57 g of the product in 99% yield.

Spectral data corresponds to data in previously published literature.[16]

**_tert_-butyl (2-amino-2-thioxoethyl)carbamate (166):**

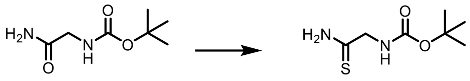

165 166

To a solution of **165** (1.5 g, 8.61 mmol, 1.0 equiv.) in THF (70 mL) was added Lawesson's reagent (6.96 g, 17.22 mmol, 2.0 equiv.). The reaction was stirred reflux overnight. The mixture was diluted with water and EtOAc. Layers were separated and the aqueous layer was extracted 3 times with 250 mL of EtOAc. Organic fractions were combined, dried with $Na_2SO_4$ and concentrated under reduced pressure. Silica gel flash liquid chromatography (10-30% EtOAc/*n*-Hexane, linear gradient) provided 1.08 g of the product in 66% yield.

Spectral data corresponds to data in previously published literature.[17]

**ethyl 2-(((_tert_-butoxycarbonyl)amino)methyl)thiazole-4-carboxylate (167):**

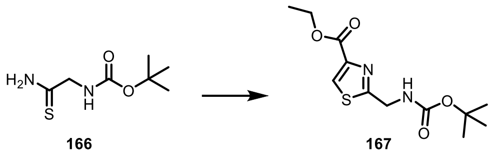

166 167

To a solution of **166** (700 mg, 3.68 mmol, 1.0 equiv.) in dry ethanol (50 mL) were added ethyl bromopyruvate (509 µL, 4.04 mmol, 1.1 equiv.) and calcium carbonate (220 mg, 2.2 mmol, 0.6 equiv.). The mixture was stirred at room temperature under nitrogen overnight. The reaction mixture was evaporated on a rotary evaporator to a viscous residue. The mixture was diluted with saturated $NaHCO_3$ and DCM. Layers were separated and the aqueous layer was extracted 3 times with 200 mL of DCM. Organic fractions were combined, dried with $Na_2SO_4$ and concentrated under reduced pressure. Silica gel flash liquid chromatography (10-30% EtOAc/n-Hexane, linear gradient) provided 605 mg of the product in 57% yield.

Spectral data corresponds to data in previously published literature.[18]

**_tert_-butyl ((4-(hydroxymethyl)thiazol-2-yl)methyl)carbamate (168):**

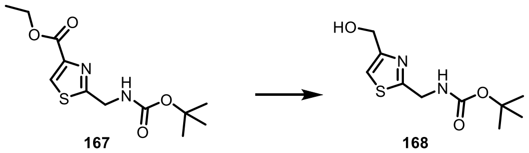

167 168

To a solution of **167** (605 mg, 2.11 mmol, 1.0 equiv.) in $Et_2O$ (40 mL) were added lithium borohydride (184 mg, 8.45 mmol, 4.0 equiv.) and methanol (342 µL, 8.45 mmol, 4.0 equiv.). The mixture was refluxed overnight. The mixture was diluted with water and EtOAc. Layers were separated and the aqueous layer was extracted 3 times with 100 mL of EtOAc. Organic fractions were combined, washed with brine, dried with $Na_2SO_4$ and concentrated under reduced pressure. Silica gel flash liquid chromatography (10-40% EtOAc/n-Hexane, linear gradient) provided 314 mg of the product in 61% yield.

Spectral data corresponds to data in previously published literature.[19]

***tert*-butyl ((4-(((2-amino-6-chloropyrimidin-4-yl)oxy)methyl)thiazol-2-yl)methyl)carbamate (169):**

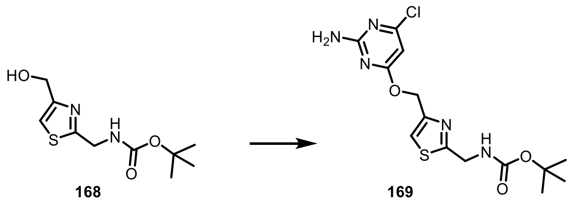


To a solution of **168** (105 mg, 0.43 mmol, 1.0 equiv.) in THF (15 mL) at 0°C was added sodium hydride (26 mg, 0.645 mmol, 1.5 equiv.). After stirring the reaction mixture for 30 minutes, **49** (103 mg, 0.43 mmol, 1.0 equiv.) was added. The reaction was stirred overnight. The mixture was diluted with water and EtOAc. Layers were separated and the aqueous layer was extracted 3 times with 100 mL of EtOAc. Organic fractions were combined, washed with brine, dried with $Na_2SO_4$ and concentrated under reduced pressure. Silica gel flash liquid chromatography (20-50% EtOAc/n-Hexane, linear gradient) provided 120 mg of the product in 75% yield.

**TLC:** *Rf* = 0.42 (EtOAc/*n*-Hexane = 1:1).

**$^1$H NMR** (400 MHz, DMSO-$d_6$) $\delta$ = 7.78 (s, 1H), 7.63 (s, 1H), 7.13 (s, 2H), 6.16 (s, 1H), 5.41 – 5.25 (m, 2H), 4.37 (d, *J*=6.1, 2H), 1.41 (s, 9H) ppm.

**$^{13}$C{$^1$H} NMR** (101 MHz, DMSO-$d_6$) $\delta$ = 172.0, 170.5, 163.2, 160.5, 156.2, 150.9, 119.4, 94.9, 79.0, 63.5, 42.4, 28.6 ppm.

**HRMS** (ESI$^+$) m/z: $[M + H]^+$, calculated for $C_{14}H_{19}ClN_5O_3S^+$ 372.0892, found 372.0892.

#### Fluorophore Compounds – TMR substrates

**Compound 3:**

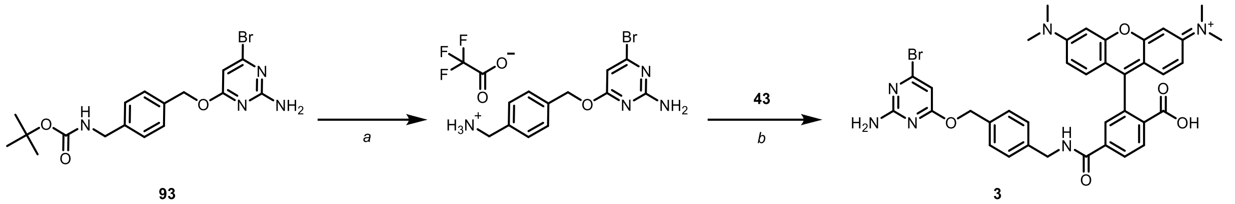


93 3

Active ester of TMR-6-COOH: TMR-6-COOH (**43**) (13 mg, 0.032 mmol, 1.2 equiv.) was dissolved in DMF (~3 mL) and DIPEA (13 mg, 0.1 mmol, 4.0 equiv.) was added followed by addition of PyAOP (17 mg, 0.32 mmol, 1.2 equiv.). The reaction mixture was stirred for 10 min.

Carbamate **95** (11 mg, 0.026 mmol, 1.0 equiv.) was dissolved in DCM (~5 mL) and cooled to 0 °C. TFA (2 mL) was added and the reaction mixture was stirred for 1 h at a constant temperature. After removing volatiles (at 25 °C), active ester of TMR was added at 25 °C to the crude ammonium residue in DMF (~ 3 mL) and reaction was stirred for 4 h. Unreacted components were quenched by addition of 0.5 mL of 10% AcOH in water, volatiles removed under reduced pressure and purified by preparative HPLC to provide 8 mg (41% yield) of fluorophore Compound **3** as a red solid.

**HRMS** (ESI$^+$) m/z: [M + 2H]$^{2+}$, calculated for $C_{37}H_{35}BrN_6O_5^{2+}$ 361.0921, found 361.0918.

**Compound 1:**

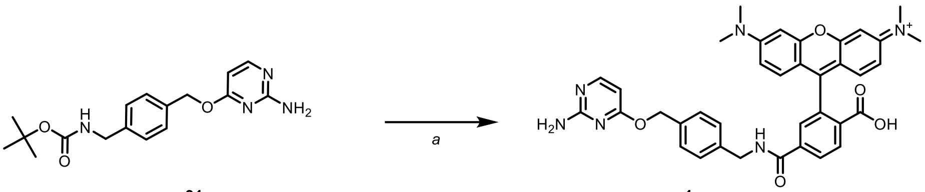


91 1

Compound (**1**) was synthesized as described above for **3**. Solid red product was isolated by preparative HPLC (9.8 mg, 45.7% yield).

**HRMS** (ESI$^+$) m/z: [M + 2H]$^{2+}$, calculated for $C_{37}H_{36}N_6O_5^{2+}$ 322.1368, found 322.1369.

**Compound 2:**

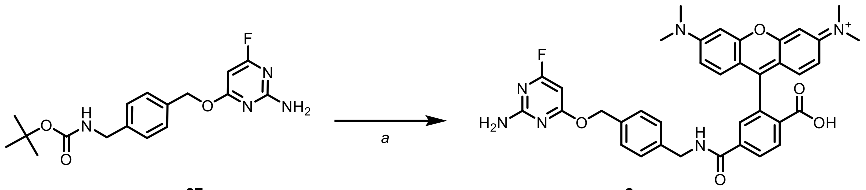


**87** **2**

Compound (**2**) was synthesized as described above for **3**. Solid red product was isolated by preparative HPLC (12 mg, 92% yield).

**HRMS** (ESI$^+$) m/z: $[M + H]^+$, calculated for $C_{37}H_{34}FN_6O_5^+$ 661.2569, found 661.2568.

**Compound 4:**

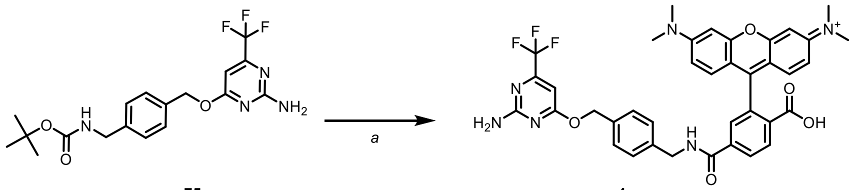


**75** **4**

Compound (**4**) was synthesized as described above for **3**. Solid red product was isolated by preparative HPLC (11 mg, 62% yield).

Spectral data corresponds to data in previously published literature.[20]

**Compound 5:**

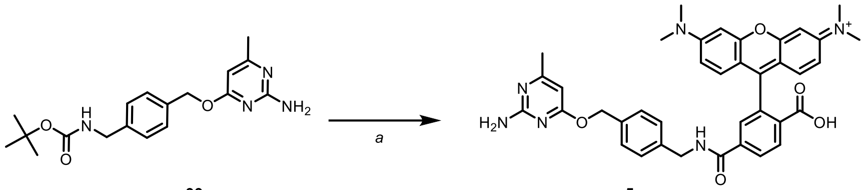


**99** **5**

Compound (**5**) was synthesized as described above for **3**. Solid red product was isolated by preparative HPLC (18 mg, 78% yield).

**HRMS** (ESI$^+$) m/z: $[M + 2H]^{2+}$, calculated for $C_{38}H_{38}N_6O_5^{2+}$ 329.1447, found 329.1446.

**Compound 6:**

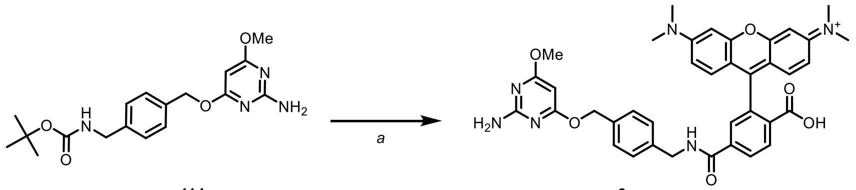

114 6

Compound (**6**) was synthesized as described above for **3**. Solid red product was isolated by preparative HPLC (3 mg, 77% yield).

**HRMS** (ESI$^+$) m/z: $[M + H]^+$, calculated for $C_{38}H_{37}N_6O_6^+$ 673.2769, found 673.2768.

**Compound 7:**

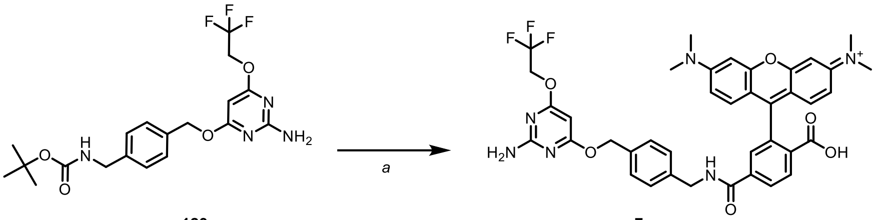

123 7

Compound (**7**) was synthesized as described above for **3**. Solid red product was isolated by preparative HPLC (16 mg, 93% yield).

**HRMS** (ESI$^+$) m/z: $[M + 2H]^{2+}$, calculated for $C_{39}H_{37}F_3N_6O_6^{2+}$ 371.1358, found 371.1360.

**Compound 8:**

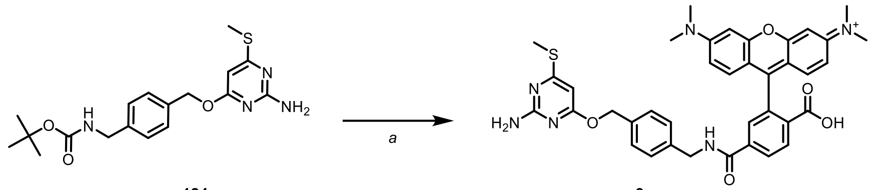

124 8

Compound (**8**) was synthesized as described above for **3**. Solid red product was isolated by preparative HPLC (11 mg, 60% yield).

**HRMS** (ESI$^+$) m/z: $[M]^+$, calculated for $C_{38}H_{37}N_6O_5S^+$ 689.2541, found 689.2541.

**Compound 9:**

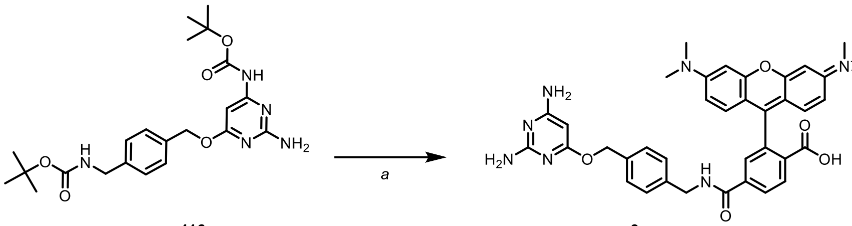

116 9

Compound (**9**) was synthesized as described above for **3**. Solid red product was isolated by preparative HPLC (15 mg, 74% yield).

**HRMS** (ESI$^+$) m/z: $[M + 2H]^{2+}$, calculated for $C_{37}H_{37}N_7O_5^{2+}$ 329.6423, found 329.6423.

**Compound 10:**

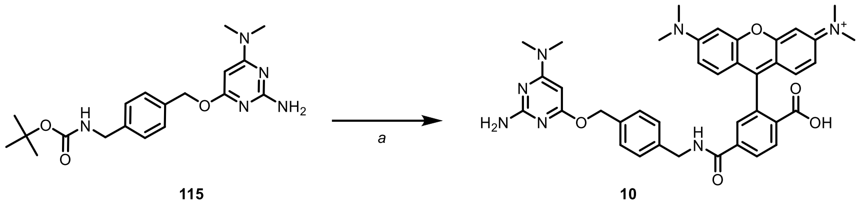

115 10

Compound (**10**) was synthesized as described above for **3**. Solid red product was isolated by preparative HPLC (15 mg, 74% yield).

**HRMS** (ESI$^+$) m/z: $[M + 2H]^{2+}$, calculated for $C_{39}H_{41}N_7O_5^{2+}$ 343.6579, found 343.6576.

**Compound 11:**

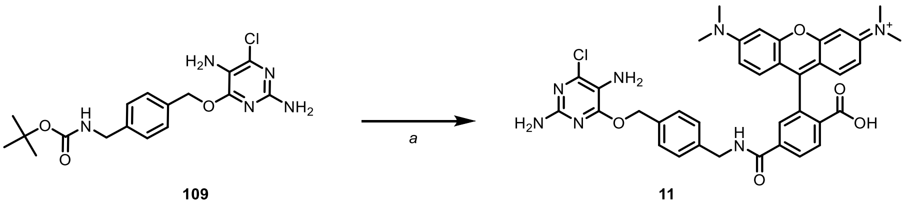

109 11

Compound (**11**) was synthesized as described above for **3**. Solid red product was isolated by preparative HPLC (6 mg, 38% yield).

**HRMS** (ESI$^+$) m/z: $[M + H]^+$, calculated for $C_{37}H_{35}ClN_7O_5^+$ 692.2383, found 692.2383.

**Compound 12:**

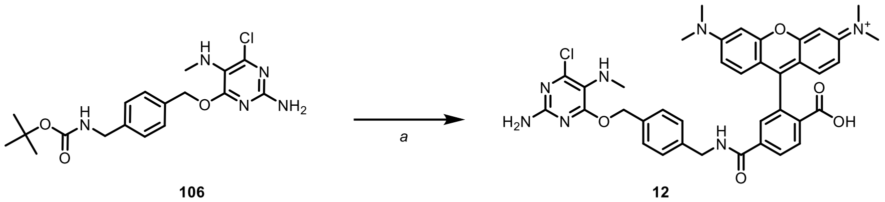


106 12

Compound (**12**) was synthesized as described above for **3**. Solid red product was isolated by preparative HPLC (6 mg, 38% yield).

**HRMS** ($ESI^+$) m/z: $[M + H]^+$, calculated for $C_{38}H_{37}ClN_7O_5^+$ 706.2539, found 706.2539.

**Compound 13:**

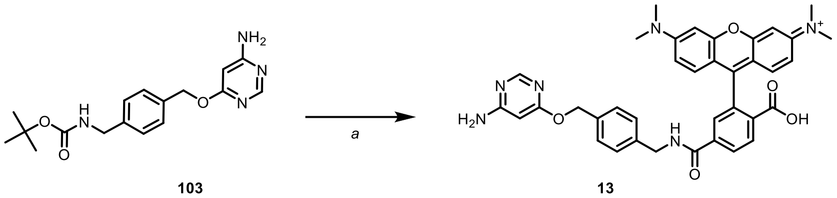


103 13

Compound (**13**) was synthesized as described above for **3**. Solid red product was isolated by preparative HPLC (14 mg, 59% yield).

**HRMS** ($ESI^+$) m/z: $[M + 2H]^{2+}$, calculated for $C_{37}H_{36}N_6O_5^{2+}$ 322.1368, found 322.1368.

**Compound 14:**

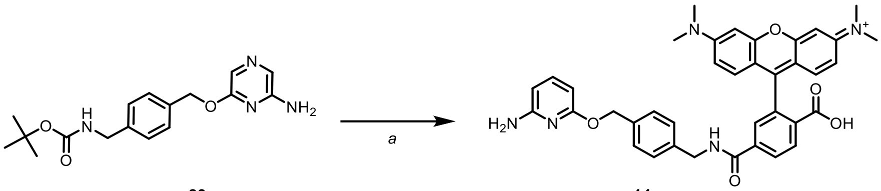


83 14

Compound (**14**) was synthesized as described above for **3**. Solid red product was isolated by preparative HPLC (12 mg, 62% yield).

**HRMS** ($ESI^+$) m/z: $[M + H]^+$, calculated for $C_{37}H_{35}N_6O_5^+$ 643.2663, found 643.2659.

**Compound 15:**

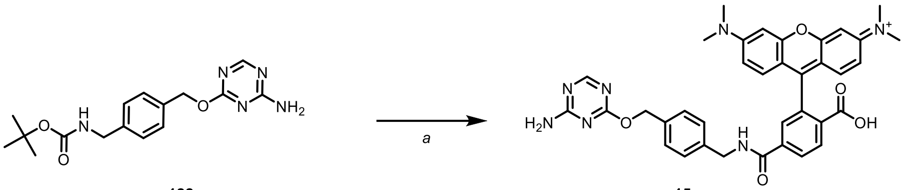

122 15

Compound (**15**) was synthesized as described above for **3**. Solid red product was isolated by preparative HPLC (7 mg, 41% yield).

**HRMS** (ESI$^+$) m/z: $[M + H]^+$, calculated for $C_{36}H_{34}N_7O_5^+$ 644.2616, found 644.2610.

**Compound 16:**

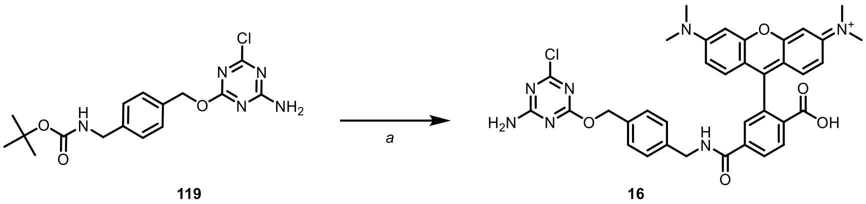

119 16

Compound (**16**) was synthesized as described above for **3**. Solid red product was isolated by preparative HPLC (11 mg, 59% yield).

HRMS (ESI$^+$) m/z: $[M + H]^+$, calculated for $C_{36}H_{33}ClN_7O_5^+$ 678.2226, found 678.2233.

**Compound 17:**

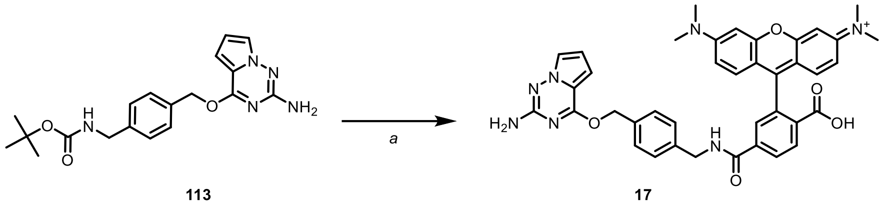

113 17

Compound (**17**) was synthesized as described above for **3**. Solid red product was isolated by preparative HPLC (4 mg, 27% yield).

**HRMS** (ESI$^+$) m/z: $[M + H]^+$, calculated for $C_{39}H_{36}N_7O_5^+$ 682.2772, found 682.2776.

**Compound 18:**

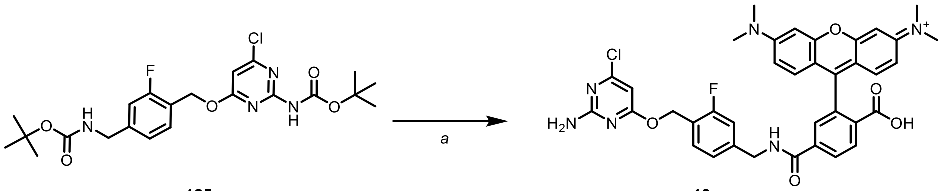

125 18

Compound (**18**) was synthesized as described above for **3**. Solid red product was isolated by preparative HPLC (6 mg, 59% yield).

**HRMS** (ESI$^+$) m/z: $[M + H]^+$, calculated for $C_{37}H_{34}ClFN_6O_5^+$ 695.2180, found 695.2179.

**Compound 19:**

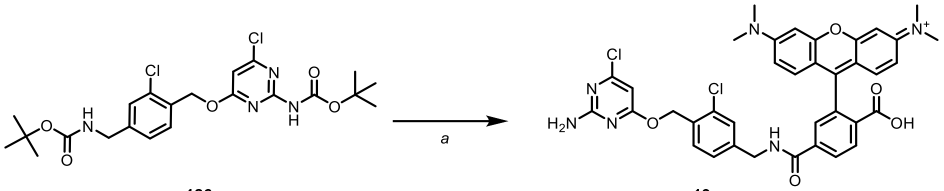

126 19

Compound (**19**) was synthesized as described above for **3**. Solid red product was isolated by preparative HPLC (9 mg, 63% yield).

**HRMS** (ESI$^+$) m/z: $[M + H]^+$, calculated for $C_{37}H_{33}Cl_2N_6O_5^+$ 711.1884, found 711.1885.

**Compound 20:**

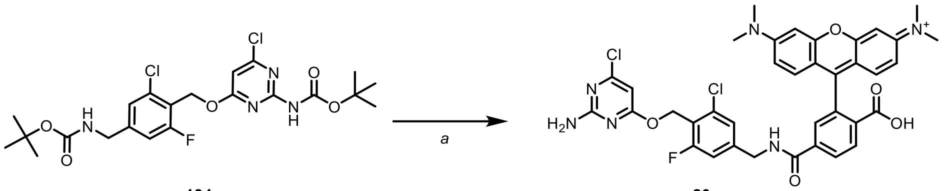

134 20

Compound (**20**) was synthesized as described above for **3**. Solid red product was isolated by preparative HPLC (9 mg, 64% yield).

**HRMS** (ESI$^+$) m/z: $[M + H]^+$, calculated for $C_{37}H_{32}N_6O_5Cl_2F^+$ 729.1790, found 729.1786.

**Compound 21:**

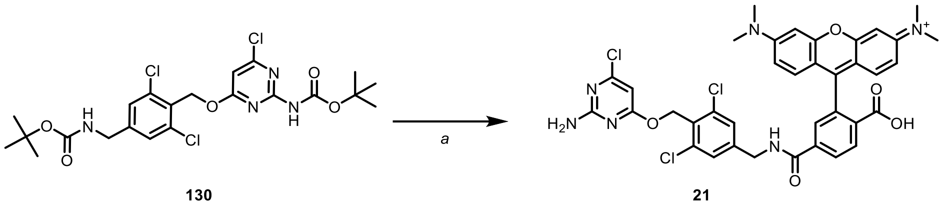


130 21

Compound (**21**) was synthesized as described above for **3**. Solid red product was isolated by preparative HPLC (4 mg, 29% yield).

**HRMS** (ESI$^+$) m/z: $[M + 2H]^{2+}$, calculated for $C_{37}H_{32}N_6O_5Cl_3H^{2+}$ 373.0784, found 373.0782.

**Compound 22: CF-TMR**

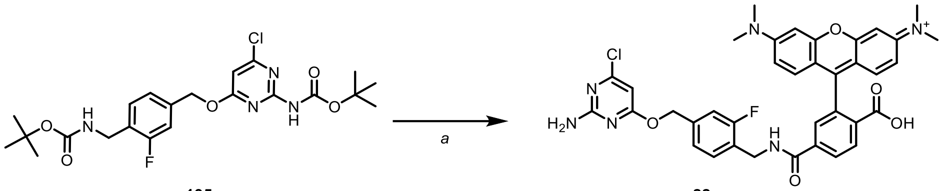


135 22

Compound (**22**) was synthesized as described above for **3**. Solid red product was isolated by preparative HPLC (7 mg, 48% yield).

**HRMS** (ESI$^+$) m/z: $[M + H]^+$, calculated for $C_{37}H_{34}ClFN_6O_5^+$ 695.2180, found 695.2182.

**Compound 23:**

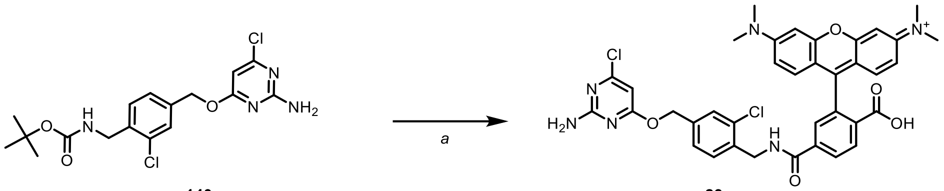


140 23

Compound (**23**) was synthesized as described above for **3**. Solid red product was isolated by preparative HPLC (6.5 mg, 40% yield).

**HRMS** (ESI$^+$) m/z: $[M + H]^+$, calculated for $C_{37}H_{33}Cl_2N_6O_5^+$ 711.1884, found 711.1891.

**Compound 24:**

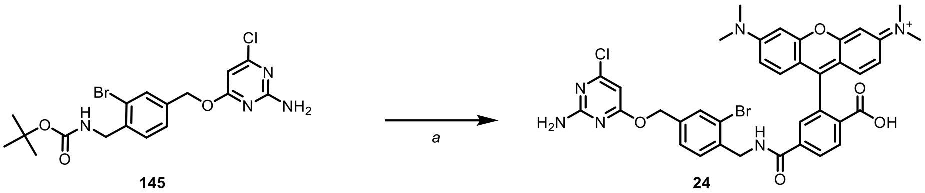

145 24

Compound (**24**) was synthesized as described above for **3**. Solid red product was isolated by preparative HPLC (4 mg, 21% yield).

**HRMS** (ESI$^+$) m/z: [M + H]$^+$, calculated $C_{37}H_{33}BrClN_6O_5^+$ 755.1379, found 755.1385.

**Compound 25:**

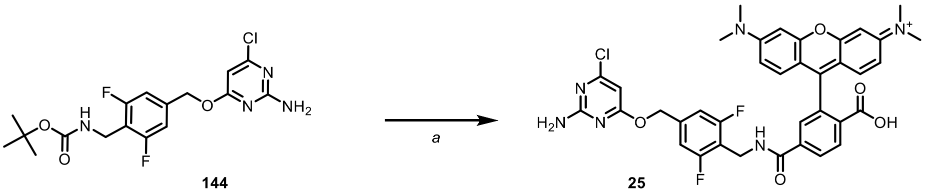

144 25

Compound (**25**) was synthesized as described above for **3**. Solid red product was isolated by preparative HPLC (2 mg, 30% yield).

**HRMS** (ESI$^+$) m/z: [M + H]$^+$, calculated $C_{37}H_{32}ClF_2N_6O_5^+$ 713.2085, found 713.2089.

**Compound 26:**

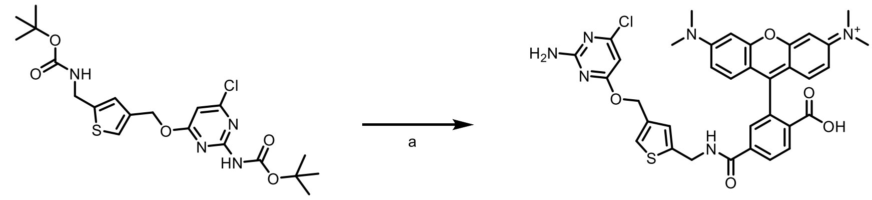

156 26

Compound (**26**) was synthesized as described above for **3**. Solid red product was isolated by preparative HPLC (4 mg, 27% yield).

**HRMS** (ESI$^+$) m/z: [M + H]$^+$, calculated for $C_{35}H_{32}ClN_6O_5S^+$ 683.1838, found 683.1841.

**Compound 27:**

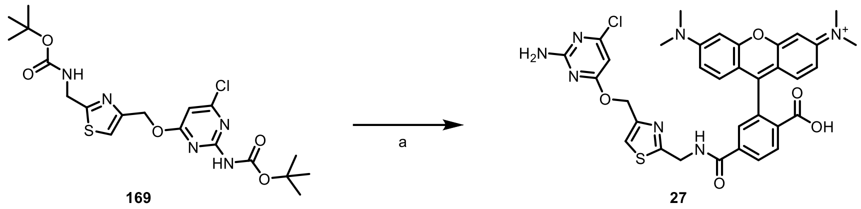


Compound (**27**) was synthesized as described above for **3**. Solid red product was isolated by preparative HPLC (12 mg, 89% yield).

**HRMS** ($ESI^+$) m/z: $[M + H]^+$, calculated for $C_{34}H_{31}ClN_7O_5S^+$ 684.1790, found 684.1797.

**Compound 28:**

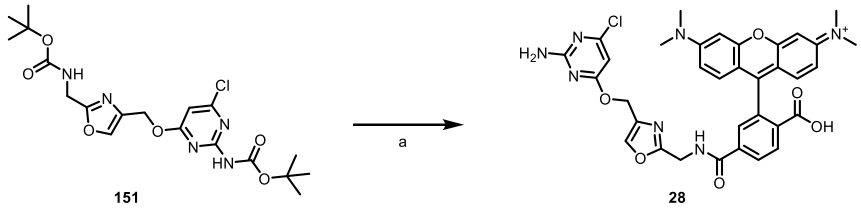


Compound (**28**) was synthesized as described above for **3**. Solid red product was isolated by preparative HPLC (13 mg, 87% yield).

**HRMS** ($ESI^+$) m/z: $[M + H]^+$, calculated for $C_{34}H_{31}ClN_7O_6^+$ 668.2019, found 683.1841.

**Compound 29:**

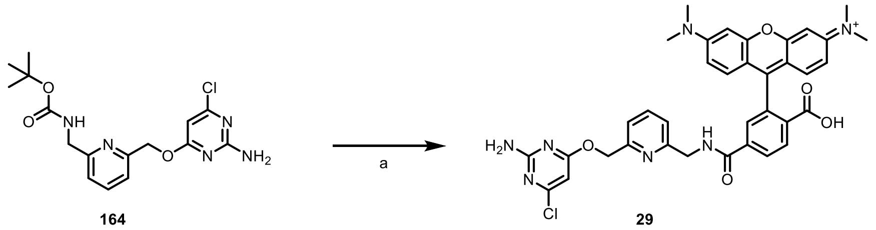


Compound (**28**) was synthesized as described above for **3**. Solid red product was isolated by preparative HPLC (20 mg, 54% yield).

**HRMS** ($ESI^+$) m/z: $[M + H]^+$, calculated for $C_{36}H_{33}ClN_7O_5^+$ 678.2226, found 678.2232.

**Compound 30: TF-TMR**

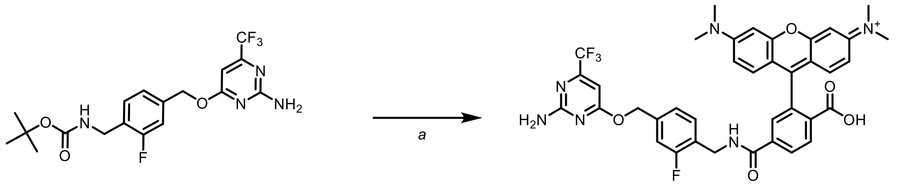



**146** **30**

Compound (**30**) was synthesized as described above for **3**. Solid red product was isolated by preparative HPLC (9 mg, 47% yield).

**HRMS** ($ESI^+$) m/z: $[M]^+$, calculated $C_{38}H_{33}F_4N_6O_5^+$ 729.2443, found 729.2441.

#### Fluorophore Compounds – Other fluorophores

##### Compound 31: TF-SiR

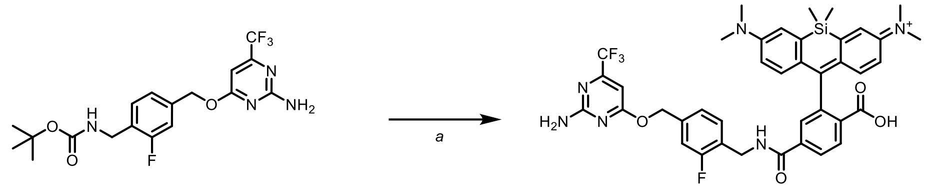

146 31

Compound (**31**) was synthesized as described above for **22**, using SiR-6-COOH as a fluorophore. Dark blue-green product was isolated by preparative HPLC (10 mg, 54% yield).

**HRMS** ($ESI^+$) m/z: $[M + H]^+$, calculated for $C_{40}H_{39}F_4N_6O_4Si^+$ 771.2733, found 771.2734.

##### Compound 32: TF-CPY

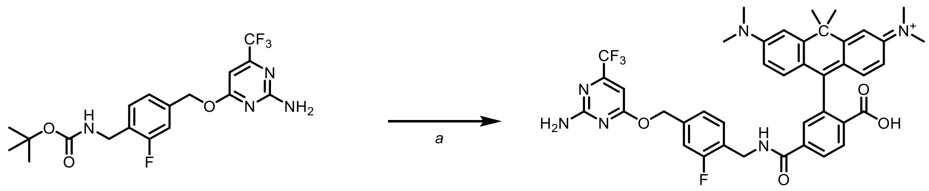

146 32

Compound (**32**) was synthesized as described above for **22**, using CPY-6-COOH as a fluorophore. Dark blue to violet product was isolated by preparative HPLC (7 mg, 77% yield).

**HRMS** ($ESI^+$) m/z: $[M]^+$, calculated for $C_{41}H_{39}F_4N_6O_4^+$ 755.2963, found 755.2963.

##### Compound 33: TF-MaP618

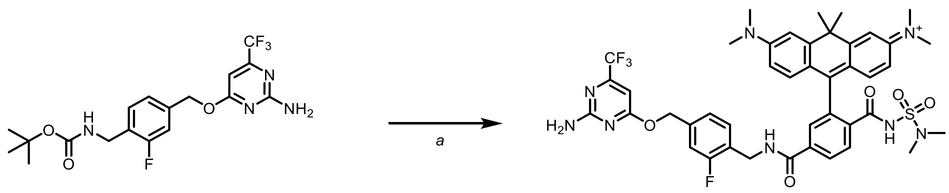

146 33

Compound (**33**) was synthesized as described above for **22**, using MaP618-6-COOH as a fluorophore. Dark blue to grey product was isolated by preparative HPLC (2 mg, 32% yield).

**HRMS** ($ESI^+$) m/z: $[M+1H]^+$, calculated for $C_{43}H_{45}F_4N_8O_5S^+$ 861.3164, found 861.3168.

**Compound 34: TF-MaP555**

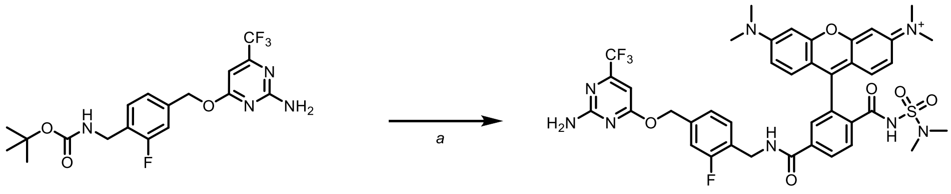

146 34

Compound (**34**) was synthesized as described above for **22**, using MaP555-6-COOH as a fluorophore. Pale red product was isolated by preparative HPLC (10 mg, 54% yield).

**HRMS** ($ESI^+$) m/z: $[M + H]^+$, calculated for $C_{40}H_{39}F_4N_8O_6S^+$ 835.2644, found 835.2647.

**Compound 35: $CF_3$P-MaP618**

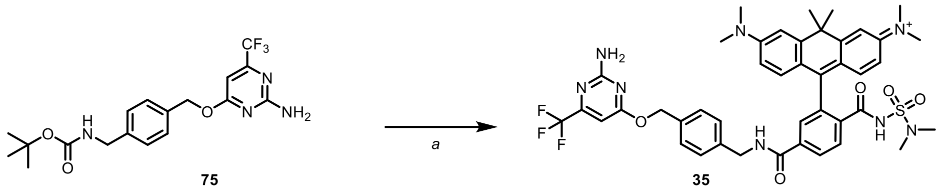

75 35

Compound (**35**) was synthesized as described above for **22**, using MaP618-6-COOH as a fluorophore. Dark blue to grey product was isolated by preparative HPLC (3 mg, 57% yield).

**HRMS** ($ESI^+$) m/z: $[M]^+$, calculated for $C_{43}H_{46}F_3N_8O_5S^+$ 843.3258, found 843.3256.

**Compound 36: CF-SiR**

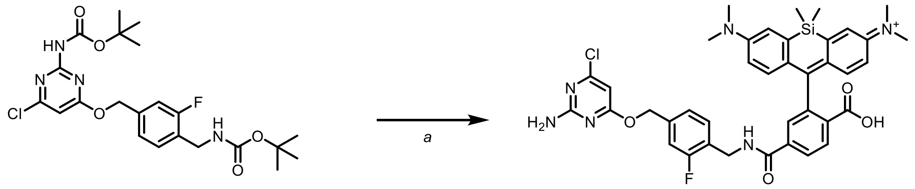

135 36

Compound (**36**) was synthesized as described above for **22**, using SiR-6-COOH as a fluorophore. Dark blue-green product was isolated by preparative HPLC (3 mg, 65% yield).

**HRMS** ($ESI^+$) m/z: $[M + H]^+$, calculated for $C_{39}H_{39}ClFN_6O_4Si^+$ 737.2469, found 737.2478.

**Compound 37: CF-CPY**

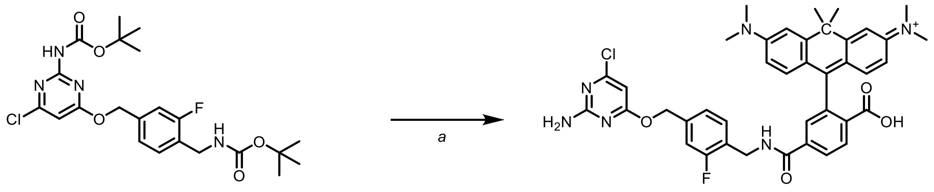

135 37

Compound (**37**) was synthesized as described above for **22**, using CPY-6-COOH as a fluorophore. Dark blue to violet product was isolated by preparative HPLC (2.5 mg, 55.7% yield).

**HRMS** ($ESI^+$) m/z: $[M]^+$, calculated for $C_{40}H_{39}ClFN_6O_4^+$ 721.2700, found 721.2706.

**Compound 38: CF-MaP555**

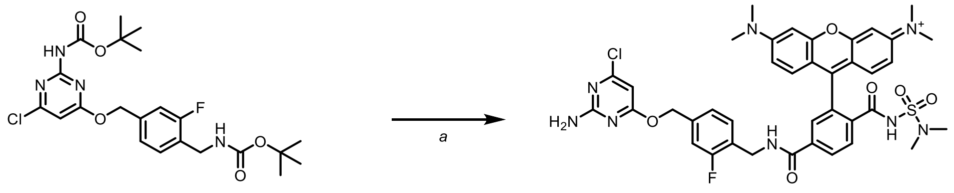

135 38

Compound (**38**) was synthesized as described above for **22**, using MaP555-6-COOH as a fluorophore. Pink to red solid product was isolated by preparative HPLC (4 mg, 24% yield).

**HRMS** ($ESI^+$) m/z: $[M+H]^+$, calculated for $C_{39}H_{39}ClFN_8O_6S^+$ 801.2380, found 801.2395.

#### Non-Fluorescent Compounds

##### Compound 39: TF-Ac

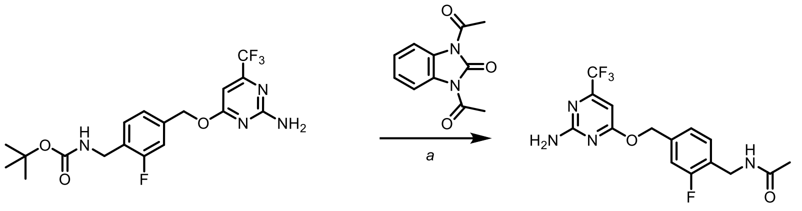

146 39

Carbamate **146** (15 mg, 0.036 mmol, 1.0 equiv.) was dissolved in DCM (~5 mL) and cooled to 0 °C. TFA (3 mL) was added and the reaction mixture was stirred at the same temperature for 2 h. Volatiles were removed in the flow of nitrogen and crude residue was used for coupling with acylating reagent (1,3-Dihydro-1,3-diacetyl-2H-benzimidazol-2-one) (9 mg, 0.043 mmol, 1.0 equiv.) like previously described[21] in DCM as solvent (~5 mL) and using triethylamine as a base (7.3 mg, 0.072 mmol, 2.0 equiv.). The reaction mixture was stirred for 8 h. White amorphous product was isolated by preparative HPLC (9 mg, 69% yield).

**HRMS** ($ESI^+$) m/z: $[M+H]^+$, calculated for $C_{15}H_{15}F_4N_4O_2^+$ 359.1126, found 359.1126.

##### Compound 40: TF-Nor

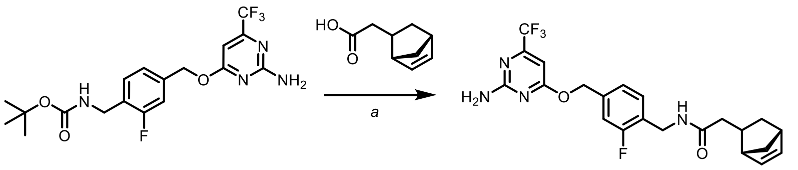

146 40

Carbamate **146** (10 mg, 0.024 mmol, 1.0 equiv.) was dissolved in DCM (~5 mL) and cooled to 0 °C. TFA (3 mL) was added and the reaction mixture was stirred at the same temperature for 2 h. Volatiles were removed in the flow of nitrogen and the crude residue was used for coupling with previously activated 5-Norbornene-2-acetic acid (5.9 mg, 0.024 mmol, 1.0 equiv.) with PyAOP (4.6 mg, 0.024 mmol, 1.0 equiv.) in DMF for 3 min in presence of DIPEA as a base (12 mg, 0.96 mmol, 4.0 equiv.). The reaction mixture was stirred for 8 h. The White amorphous product was isolated by preparative HPLC (3 mg, 27% yield).

**HRMS** ($ESI^+$) m/z: $[M+H]^+$, calculated for $C_{22}H_{23}F_4N_4O_2^+$ 451.1752, found 451.1750.

**Compound 41: TF-BCN**

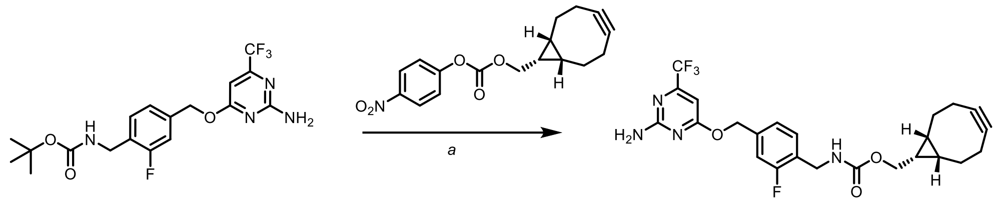


146 41

Carbamate **146** (10 mg, 0.024 mmol, 1.0 equiv.) was dissolved in DCM (~5 mL) and cooled to 0 °C. TFA (3 mL) was added and the reaction mixture was stirred at the same temperature for 2 h. Volatiles were removed in flow of nitrogen and crude residue was used for coupling with active ester rel-((1R,8S,9s)-Bicyclo[6.1.0]non-4-yn-9-yl)methyl (4-nitrophenyl) carbonate (7.5 mg, 0.024 mmol, 1.0 equiv.) in DMF in presence of DIPEA as a base (12 mg, 0.96 mmol, 4.0 equiv.). The reaction mixture was stirred for 8 h. The white amorphous product was isolated by preparative HPLC (3 mg, 25% yield).

**HRMS** ($ESI^+$) m/z: $[M+H]^+$, calculated for $C_{24}H_{25}F_4N_4O_3^+$ 493.1857, found 493.1861.

**Compound 42: TF-PhN$_3$:**

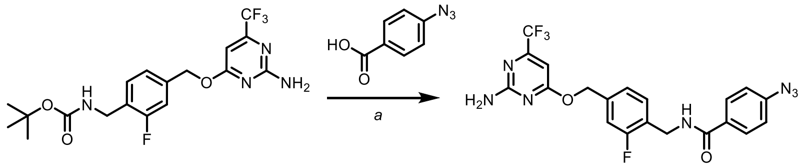


146 42

Carbamate **146** (10 mg, 0.024 mmol, 1.0 equiv.) was dissolved in DCM (~5 mL) and cooled to 0 °C. TFA (3 mL) was added and the reaction mixture was stirred at the same temperature for 2 h. Volatiles were removed in the flow of nitrogen and the crude residue was used for coupling with previously activated 5-Norbornene-2-acetic acid (3.9 mg, 0.024 mmol, 1.0 equiv.) with PyAOP (4.6 mg, 0.024 mmol, 1.0 equiv.) in DMF for 3 min in presence of DIPEA as a base (12 mg, 0.96 mmol, 4.0 equiv.). The reaction mixture was stirred for 8 h. The white amorphous product was isolated by preparative HPLC (6 mg, 54% yield).

#### NMR Spectra

##### 1H spectrum of compound 45:

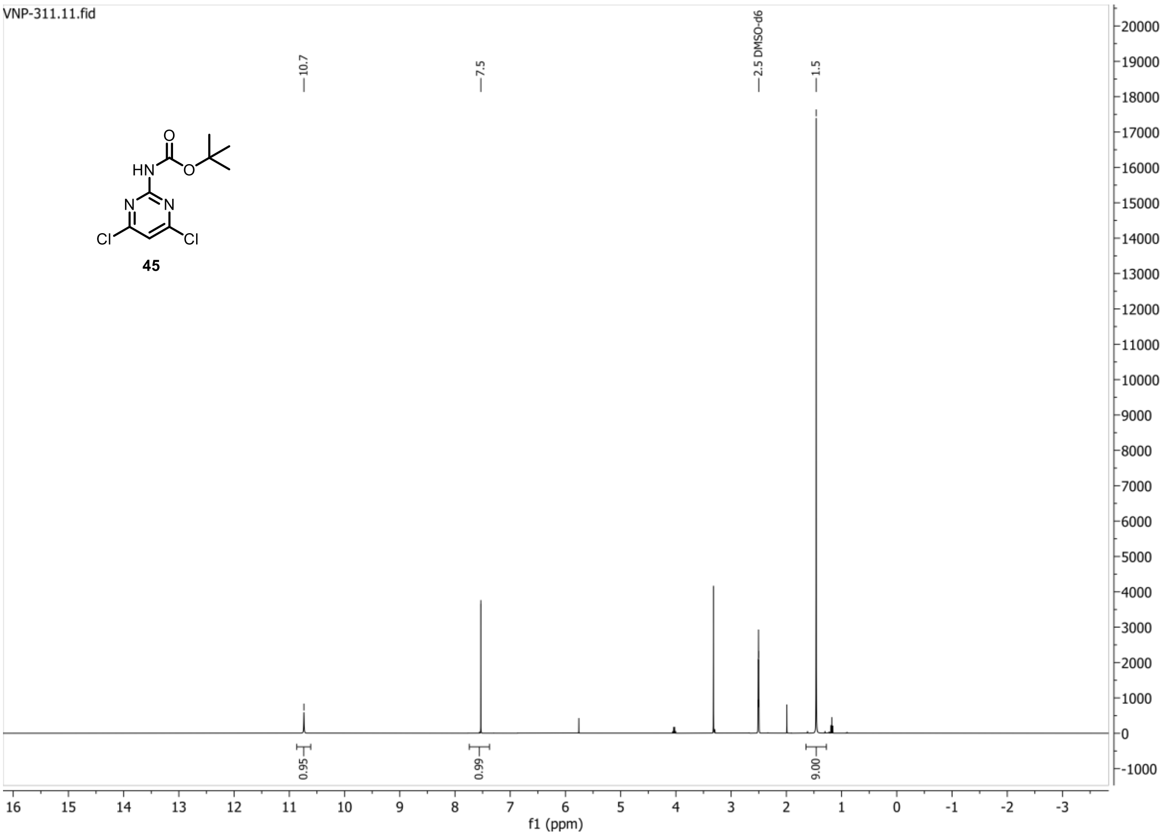


##### 13C{1H} spectrum of compound 45:

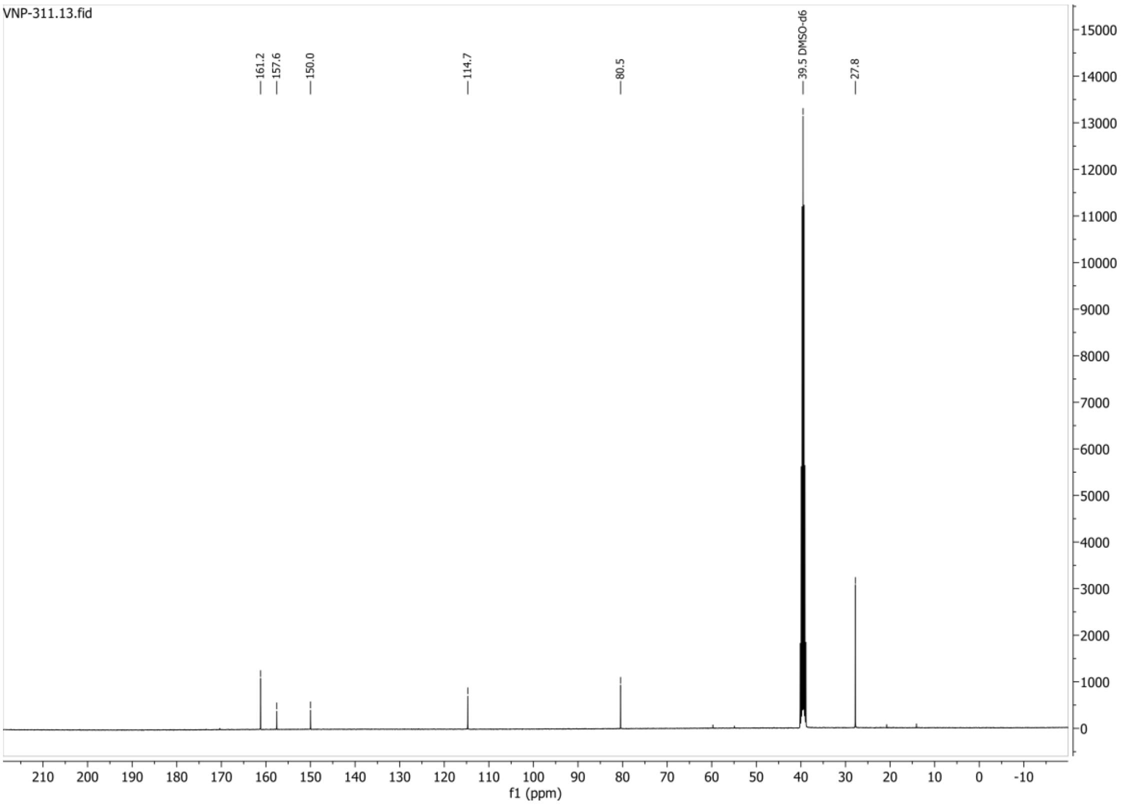

**HSQC-DEPT135 spectrum of compound 45:**

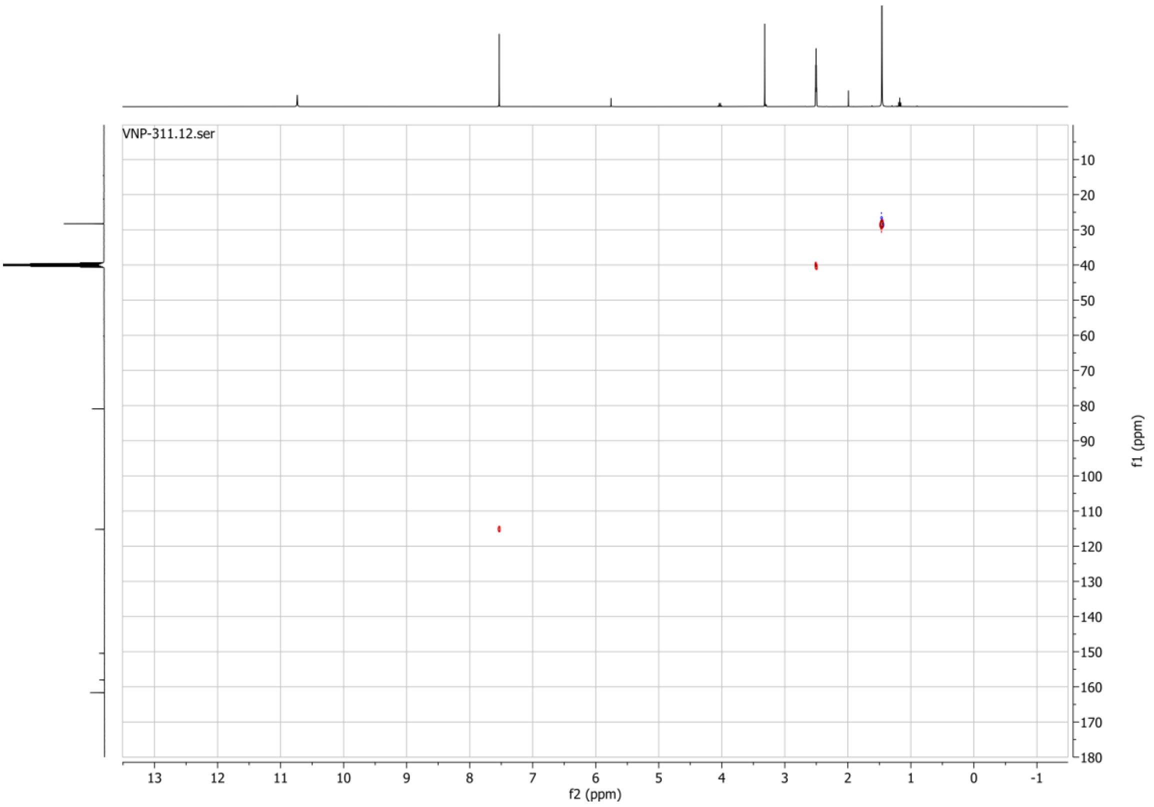

#### 1H spectrum of compound 46:

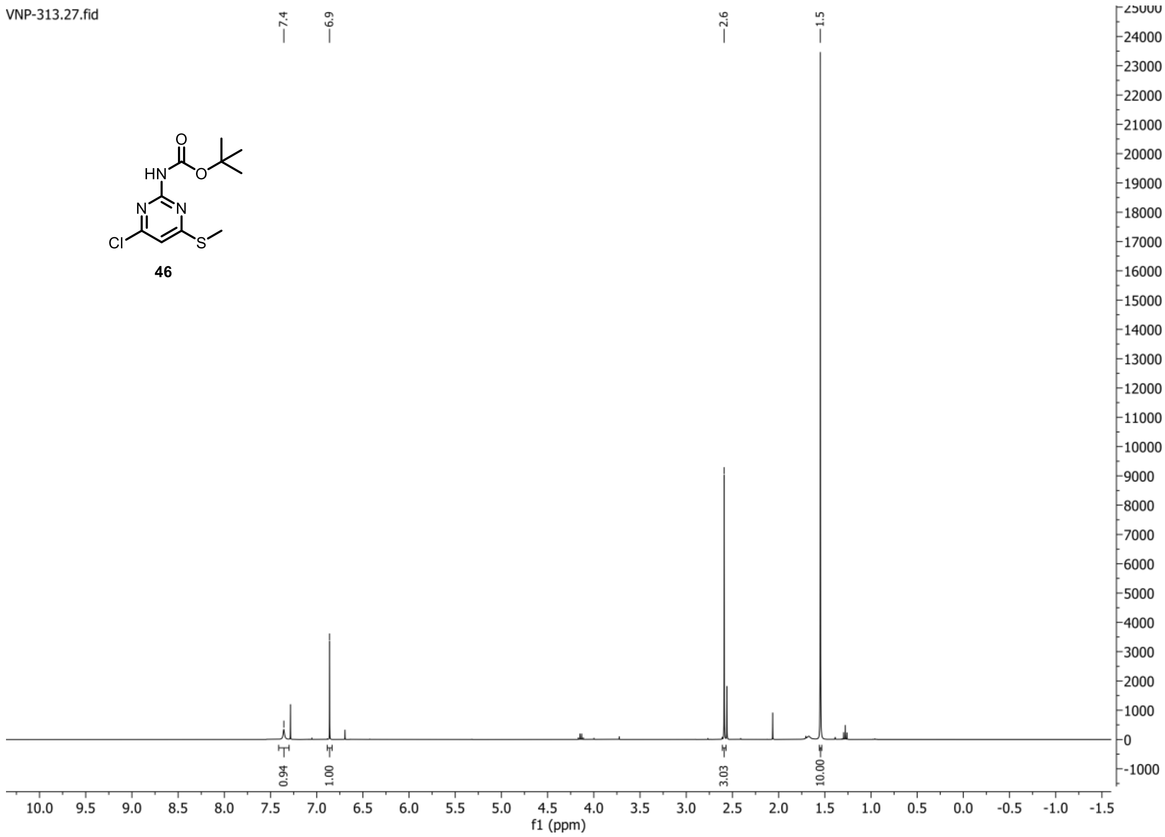


#### 13C{1H} spectrum of compound 46:

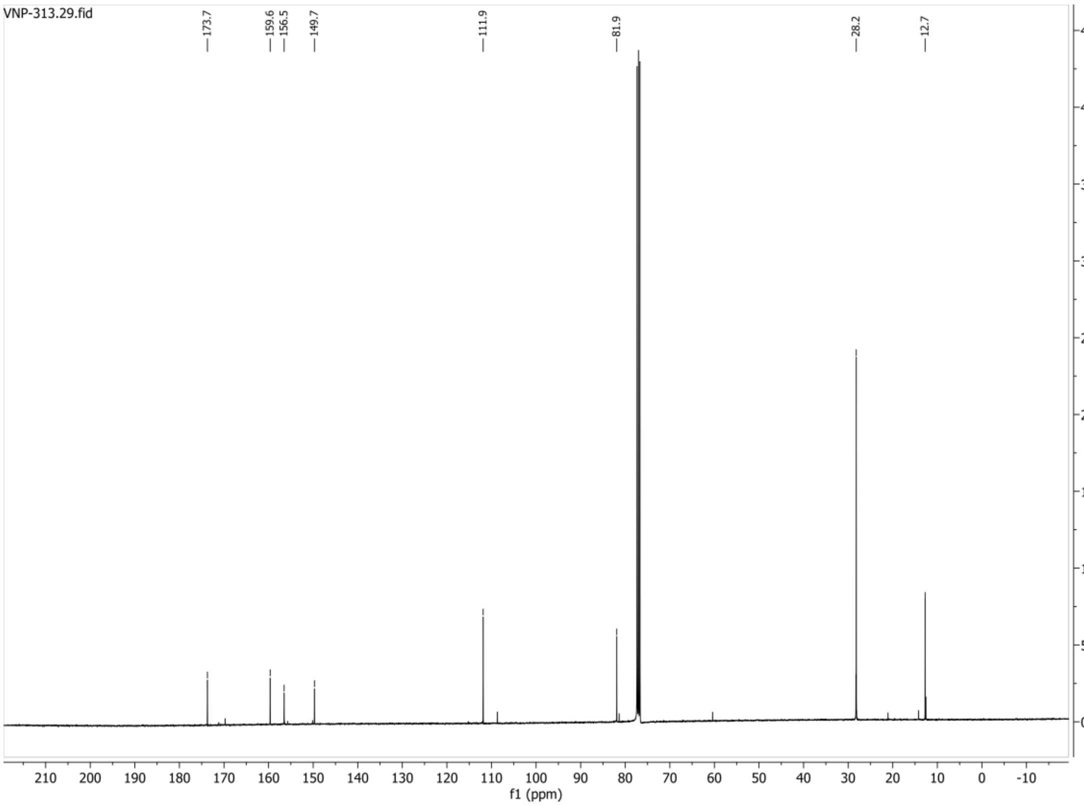

#### 1H spectrum of compound 47:

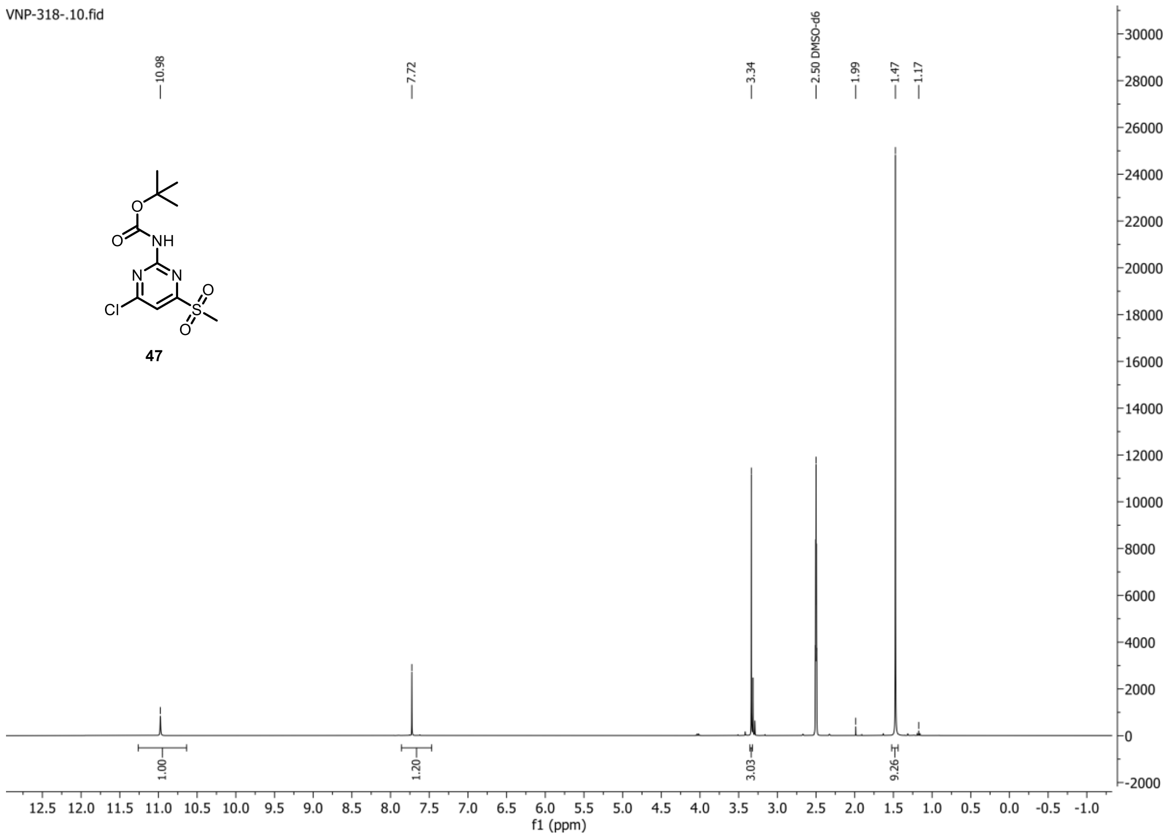


#### 13C{1H} spectrum of compound 47:

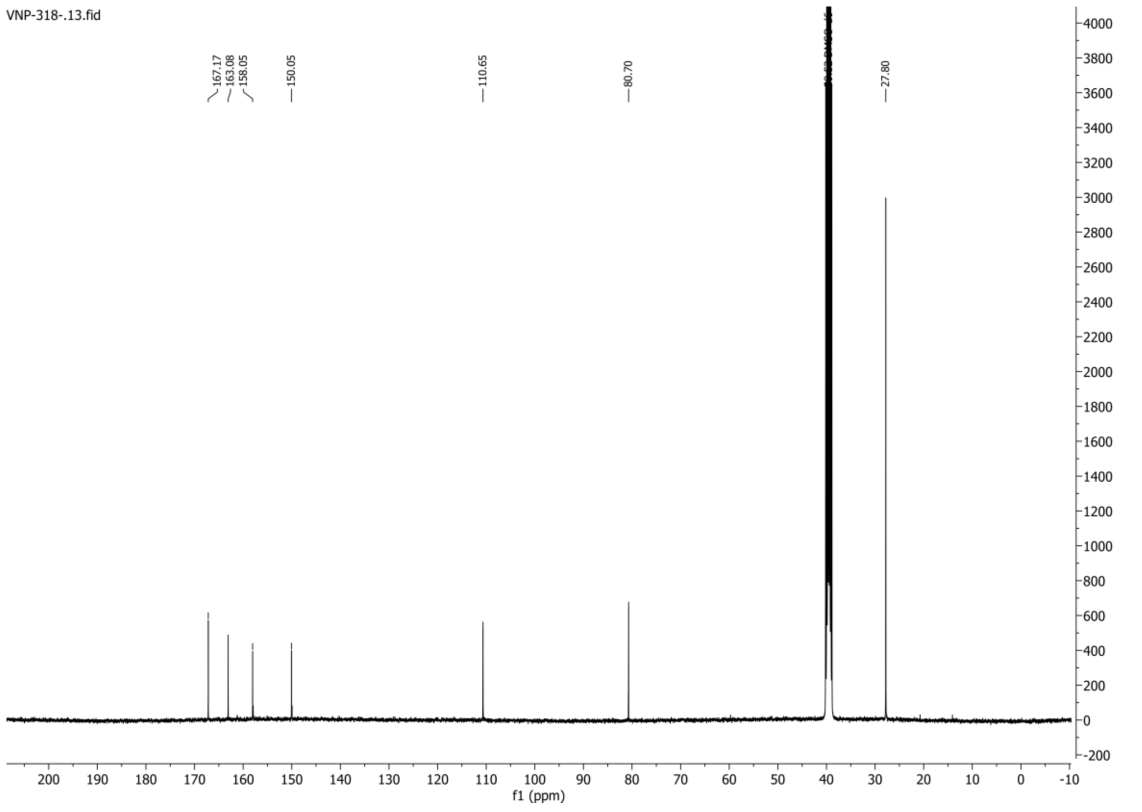

**1H spectrum of compound 49:**

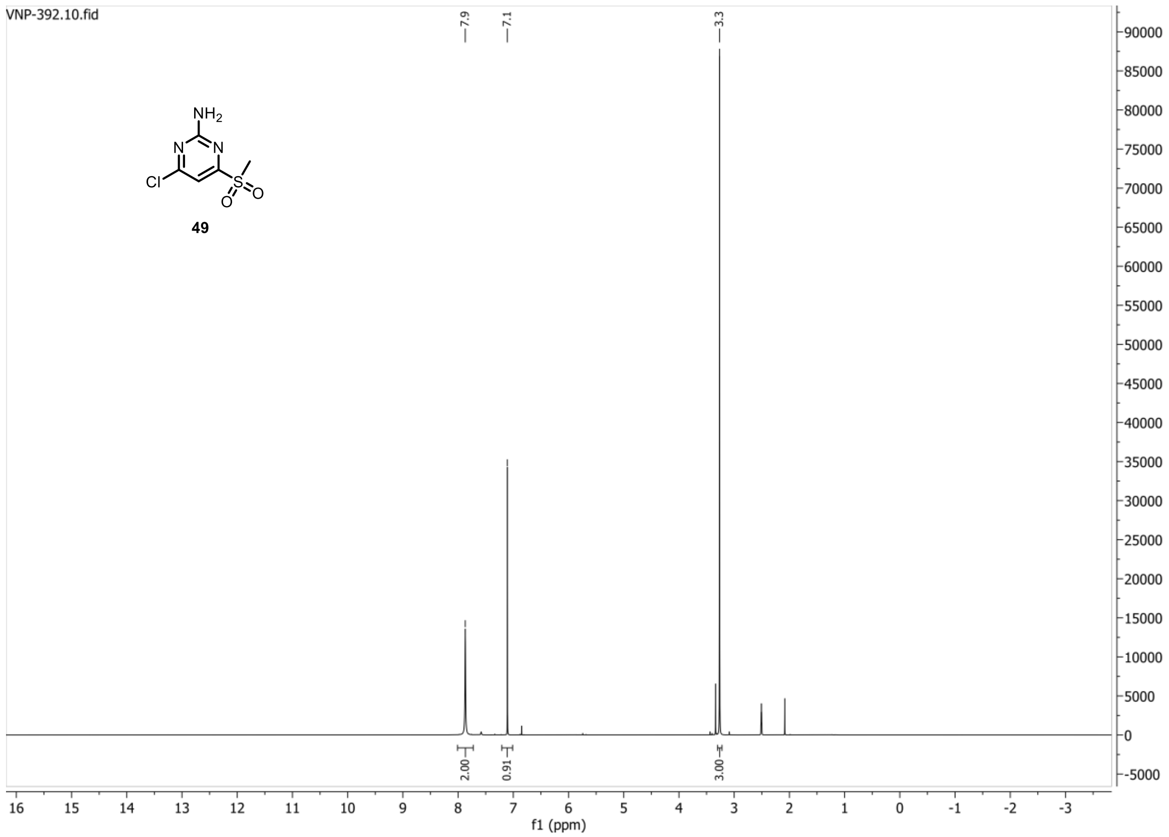


**13C{1H} spectrum of compound 49:**

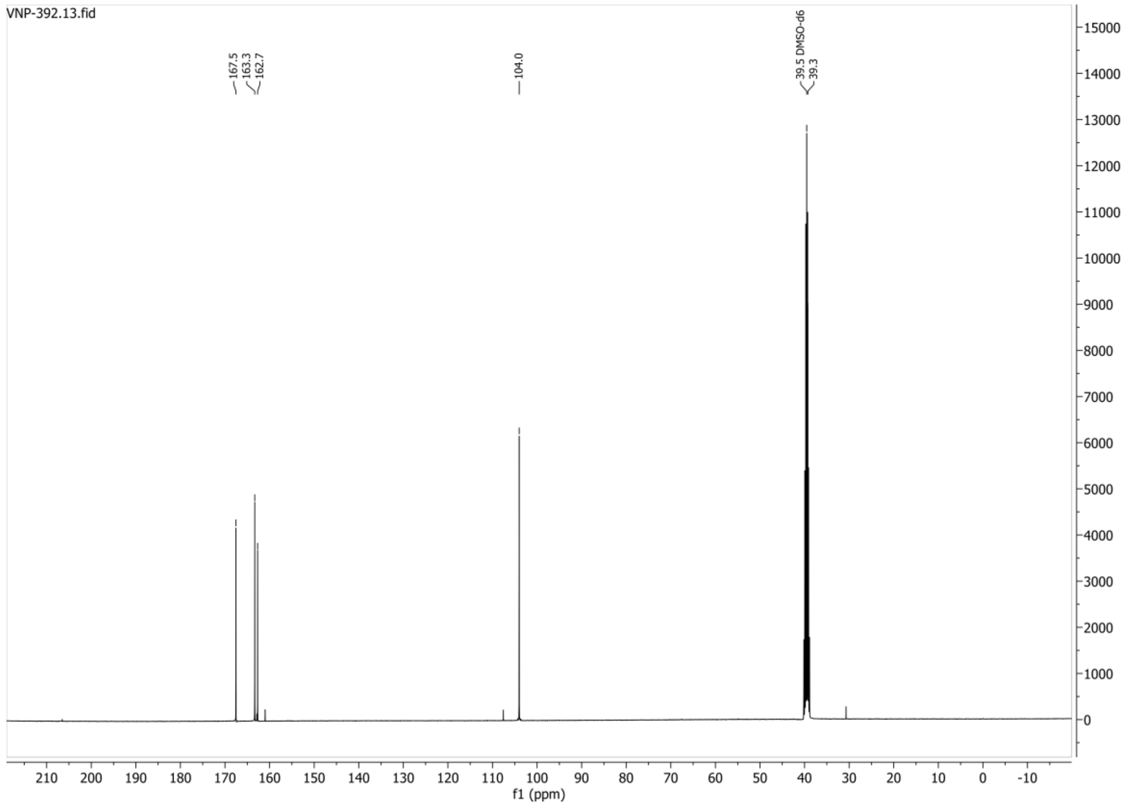

**HSQC-DEPT135 spectrum of compound 49:**

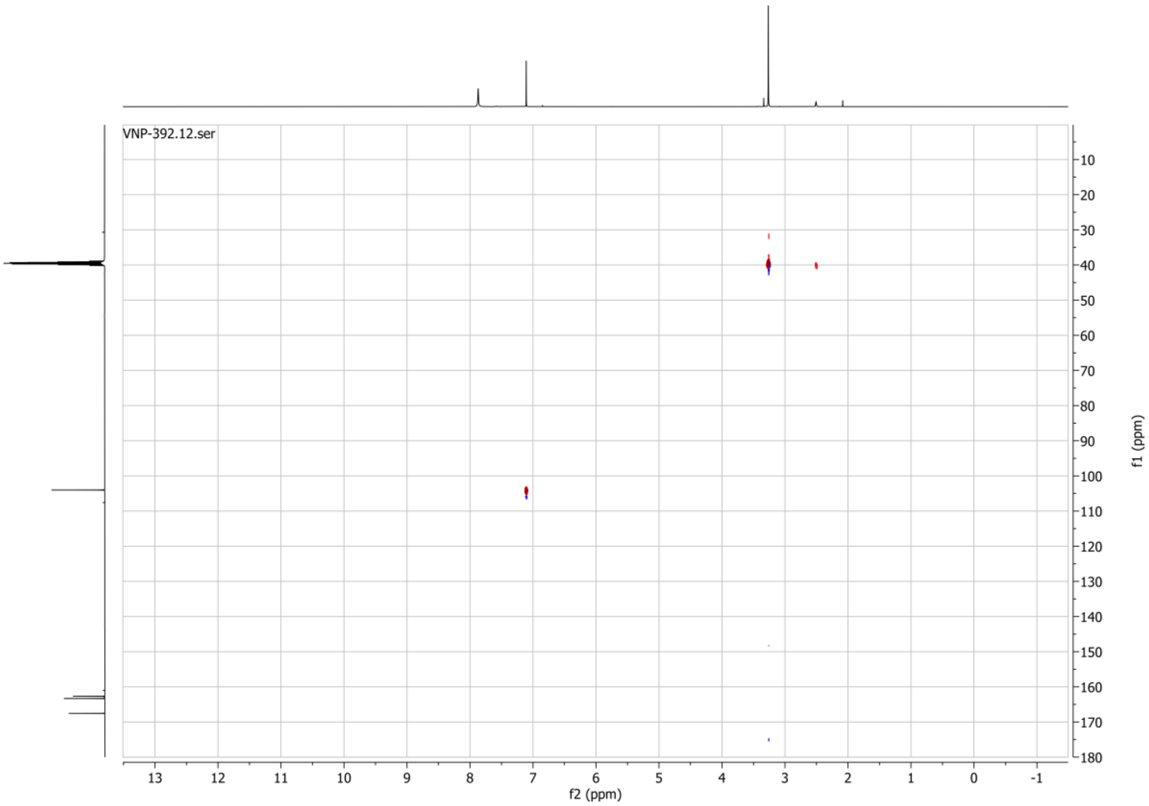

**1H spectrum of compound 52:**

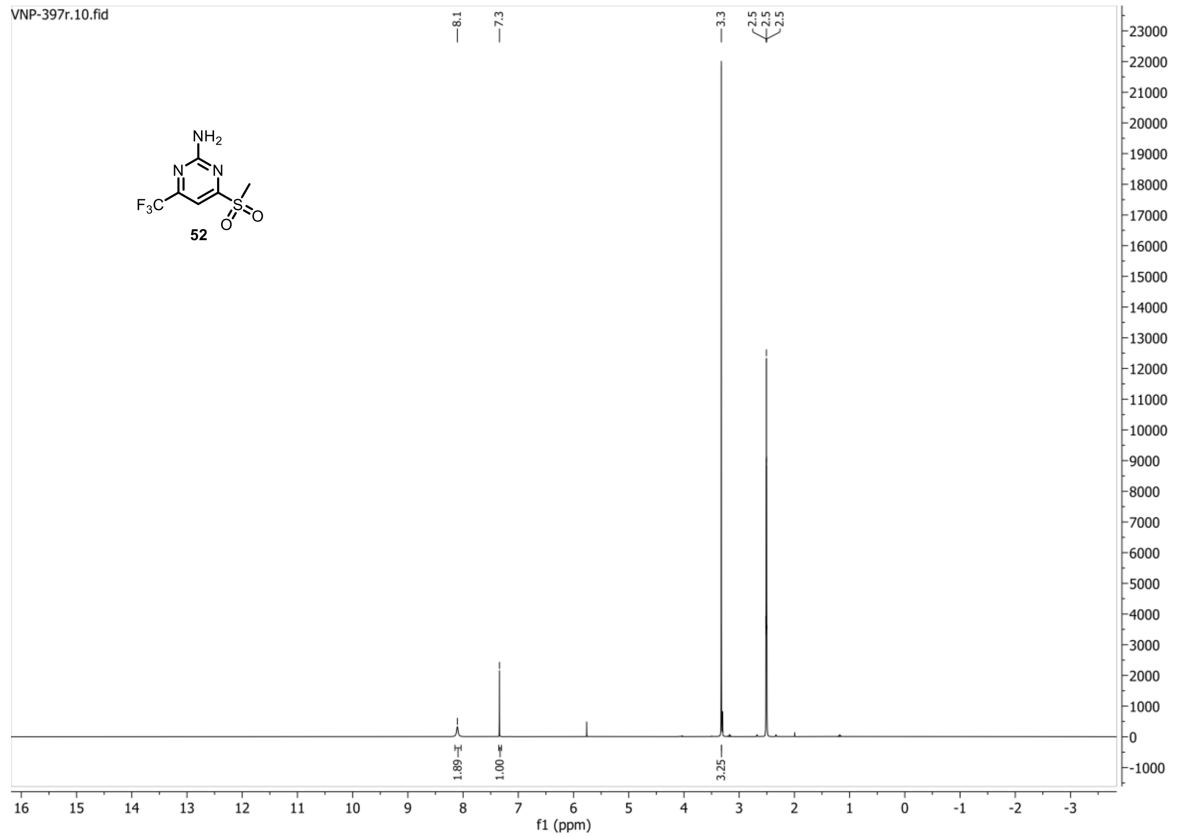


**13C{1H} spectrum of compound 52:**

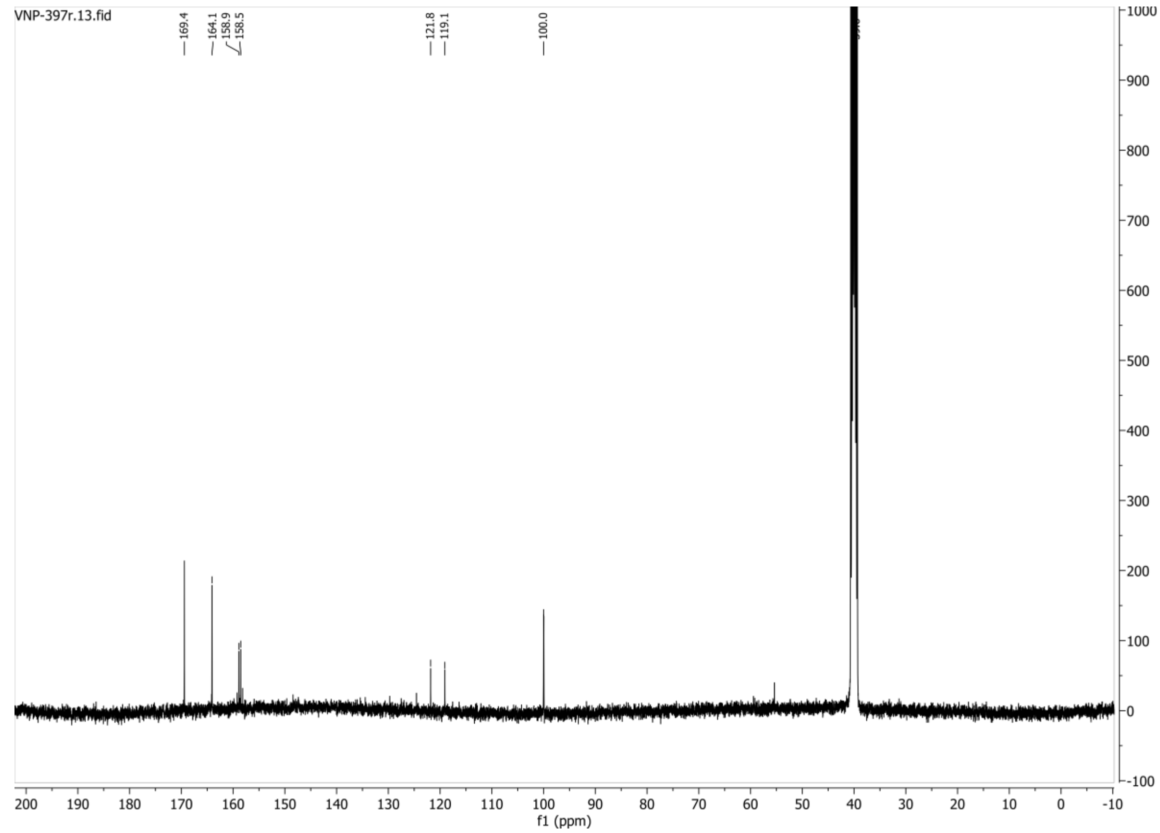

**HSQC-DEPT135 spectrum of compound 52:**

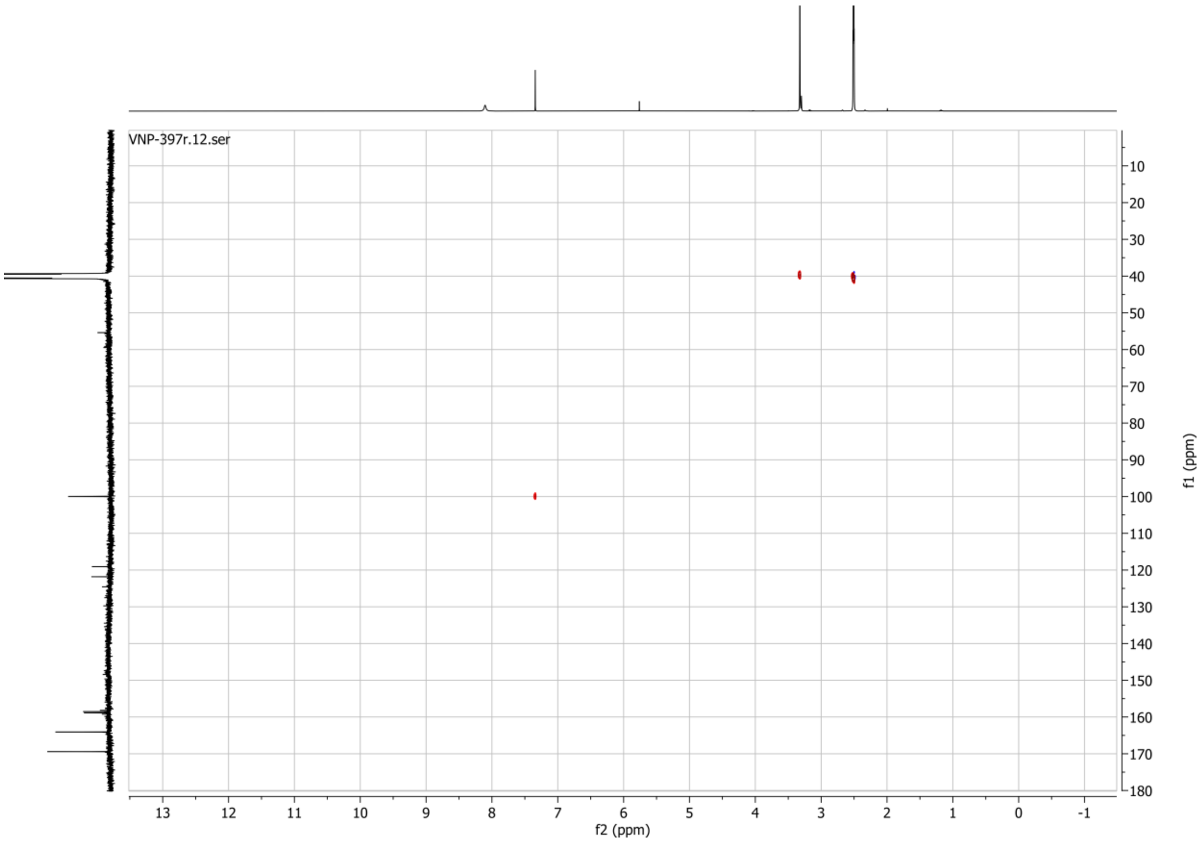


**19F spectrum of compound 52:**

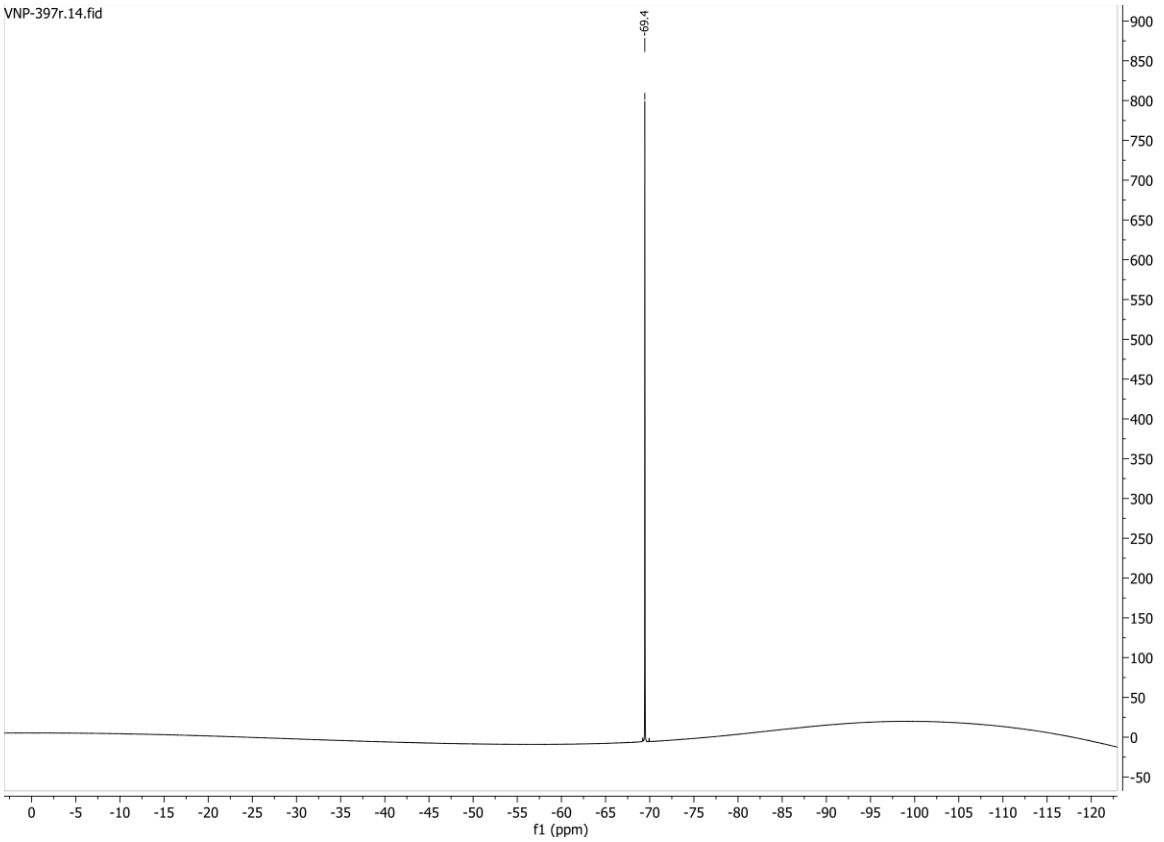

#### 1H spectrum of compound 54:

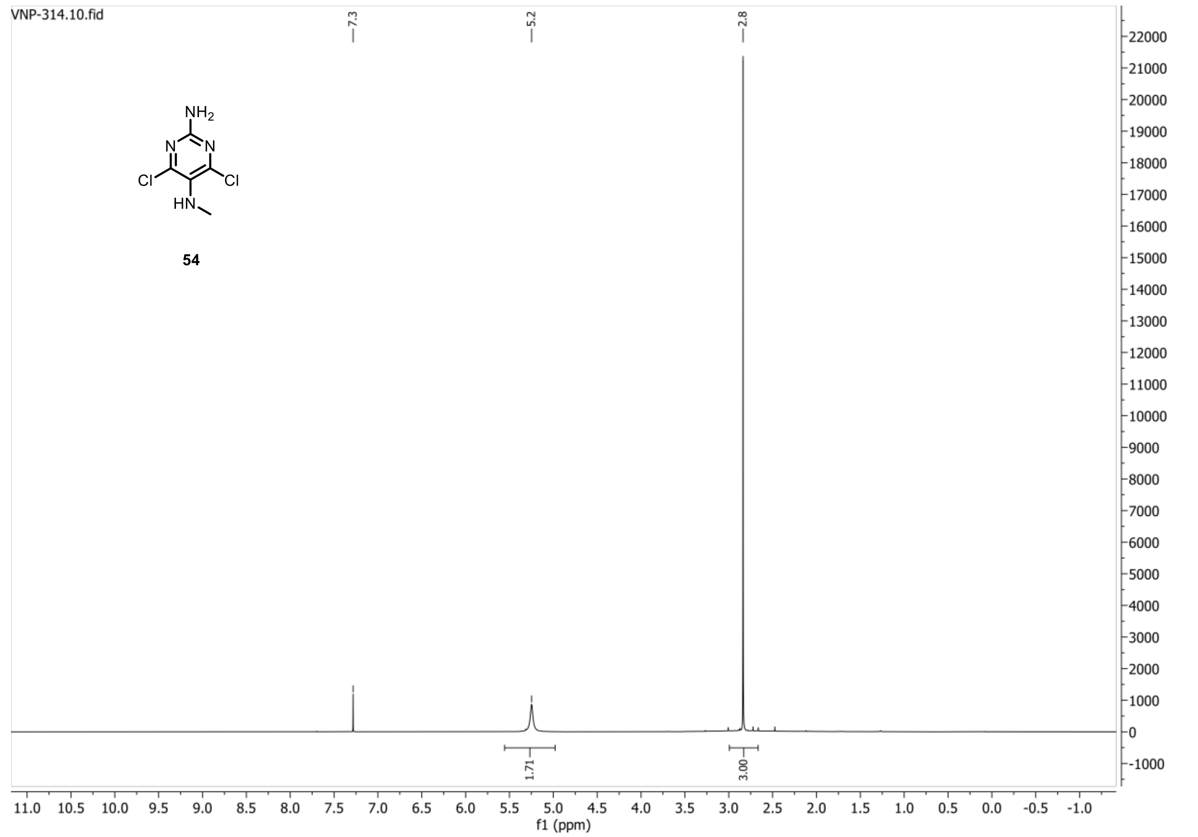


#### 13C{1H} spectrum of compound 54:

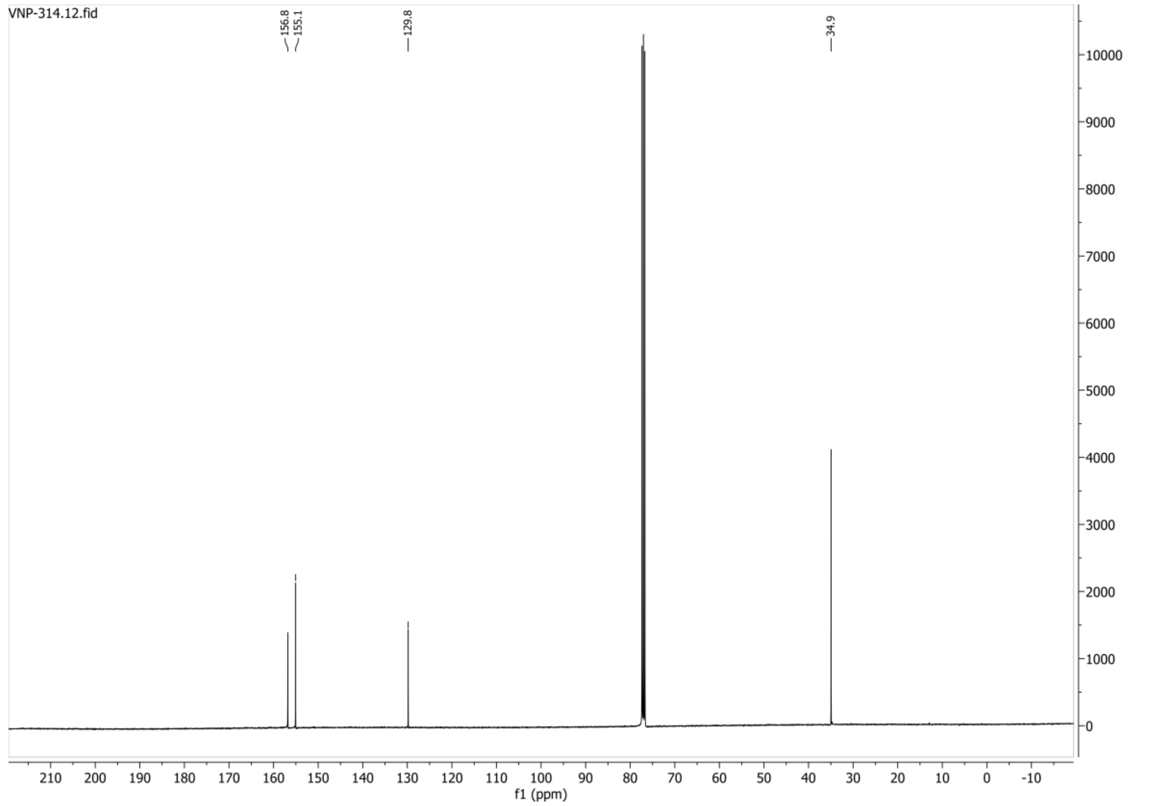

**HSQC-DEPT135 spectrum of compound 54:**

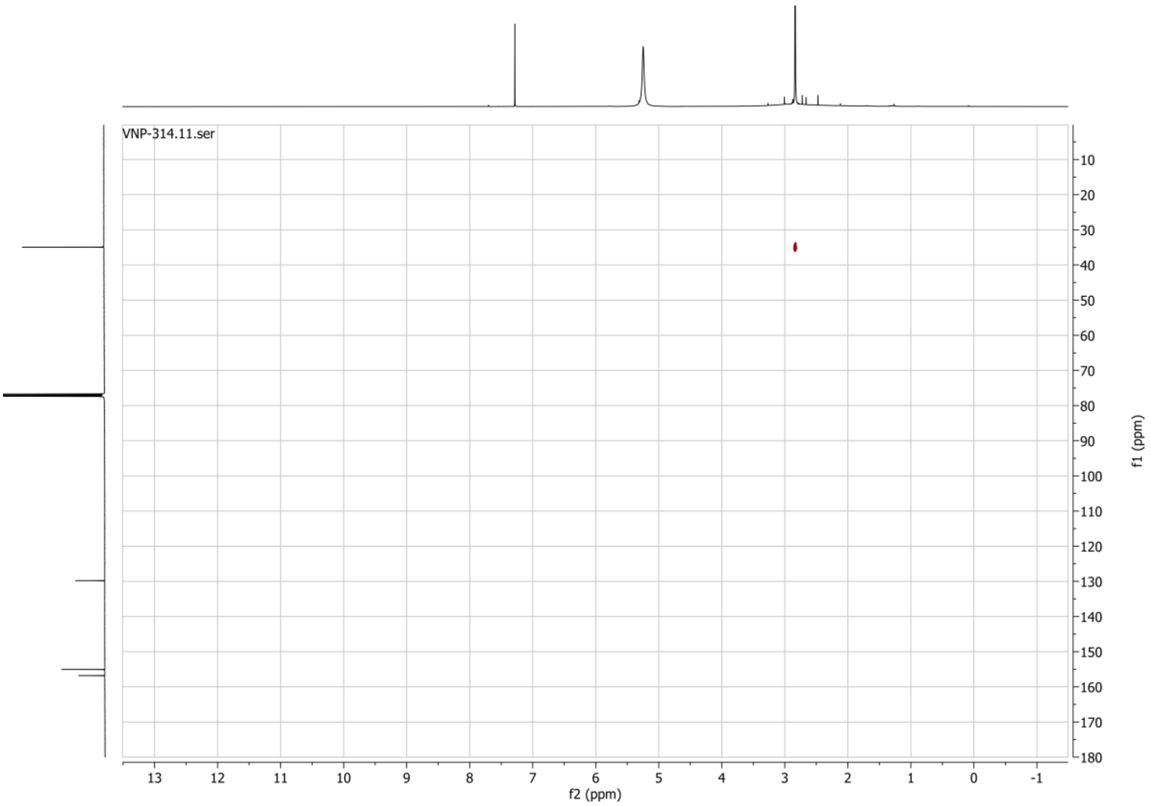

**1H spectrum of compound 58:**

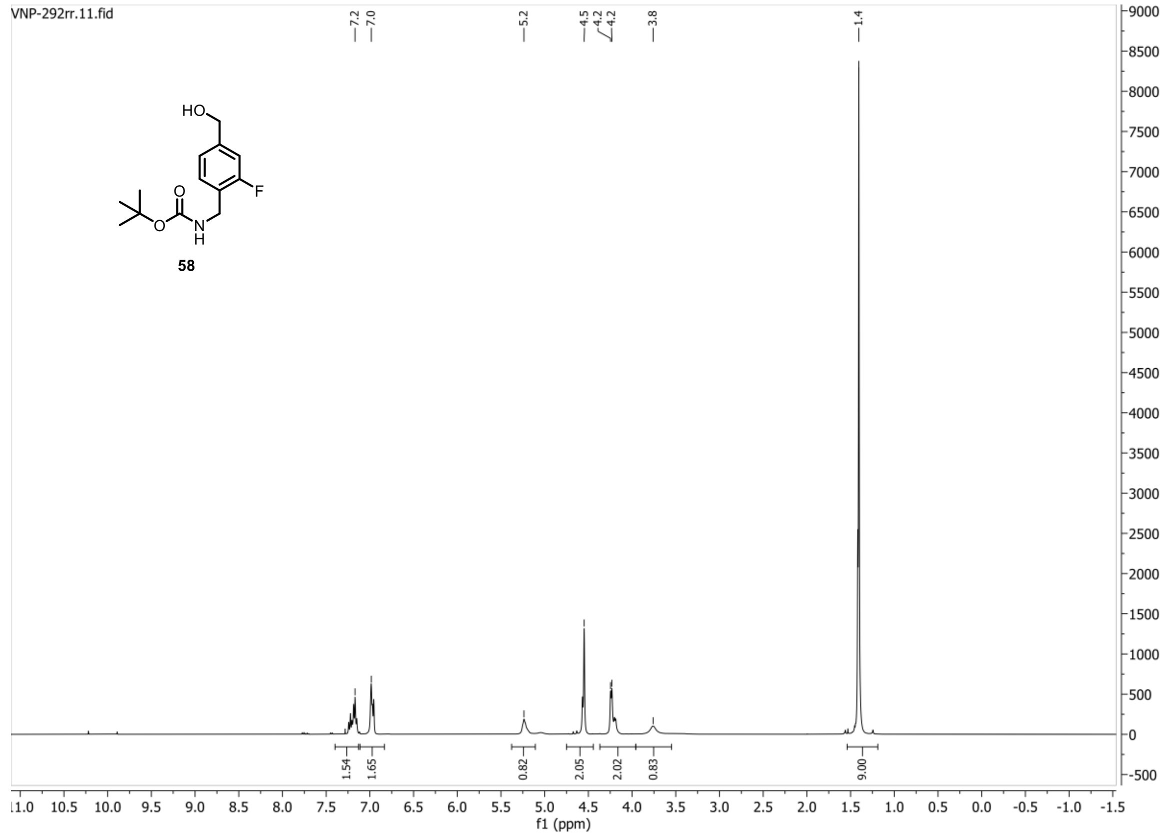


**13C{1H} spectrum of compound 58:**

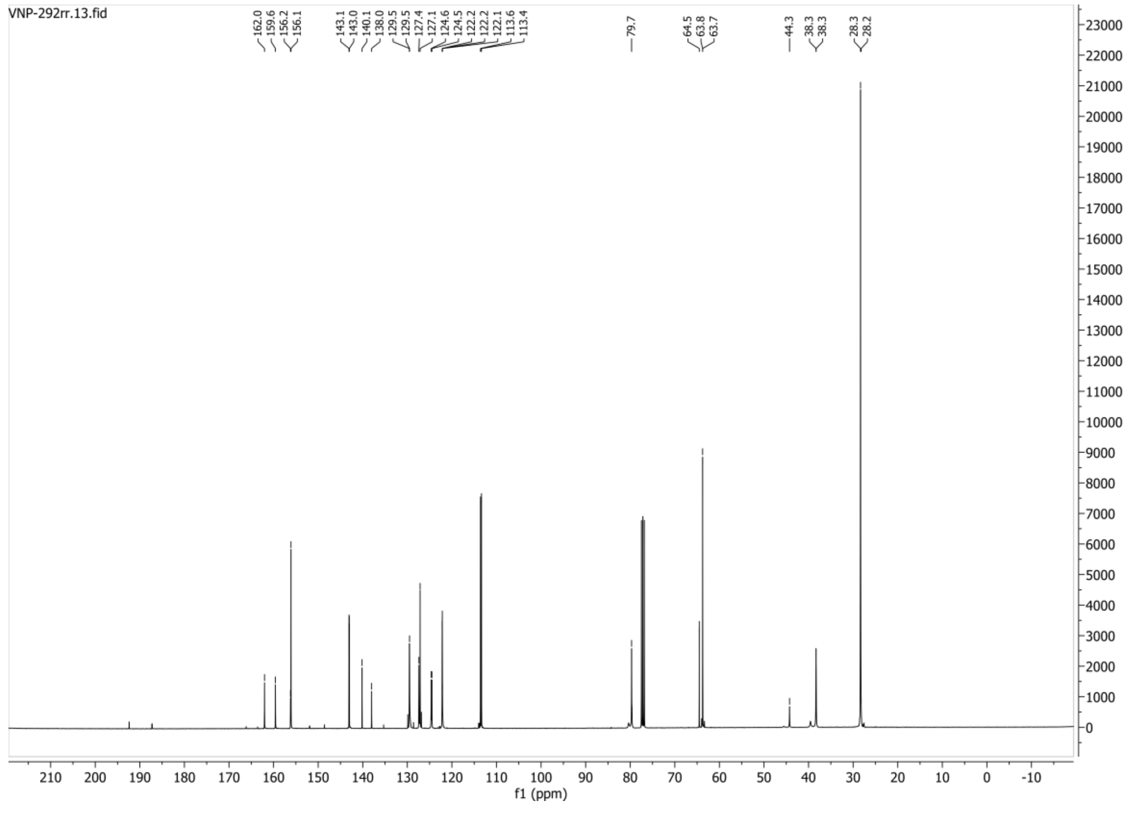

**19F spectrum of compound 58:**

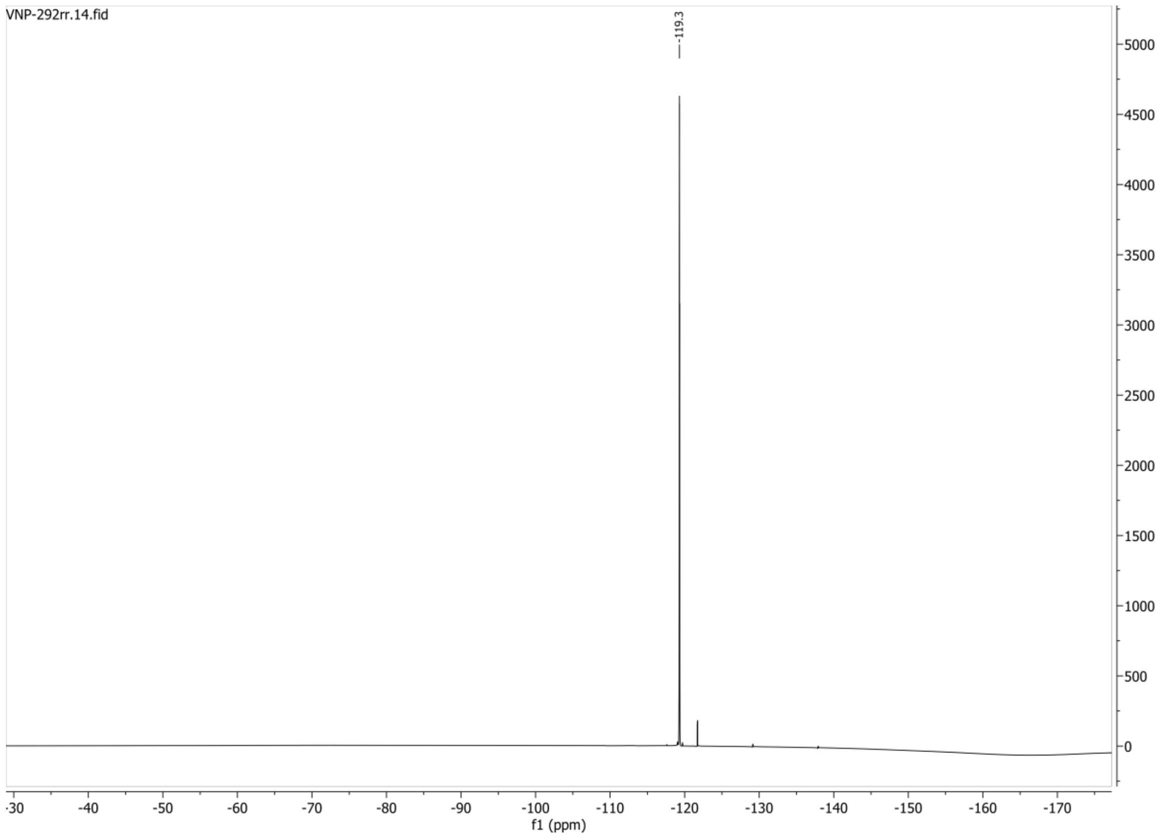


**HSQC-DEPT135 spectrum of compound 58:**

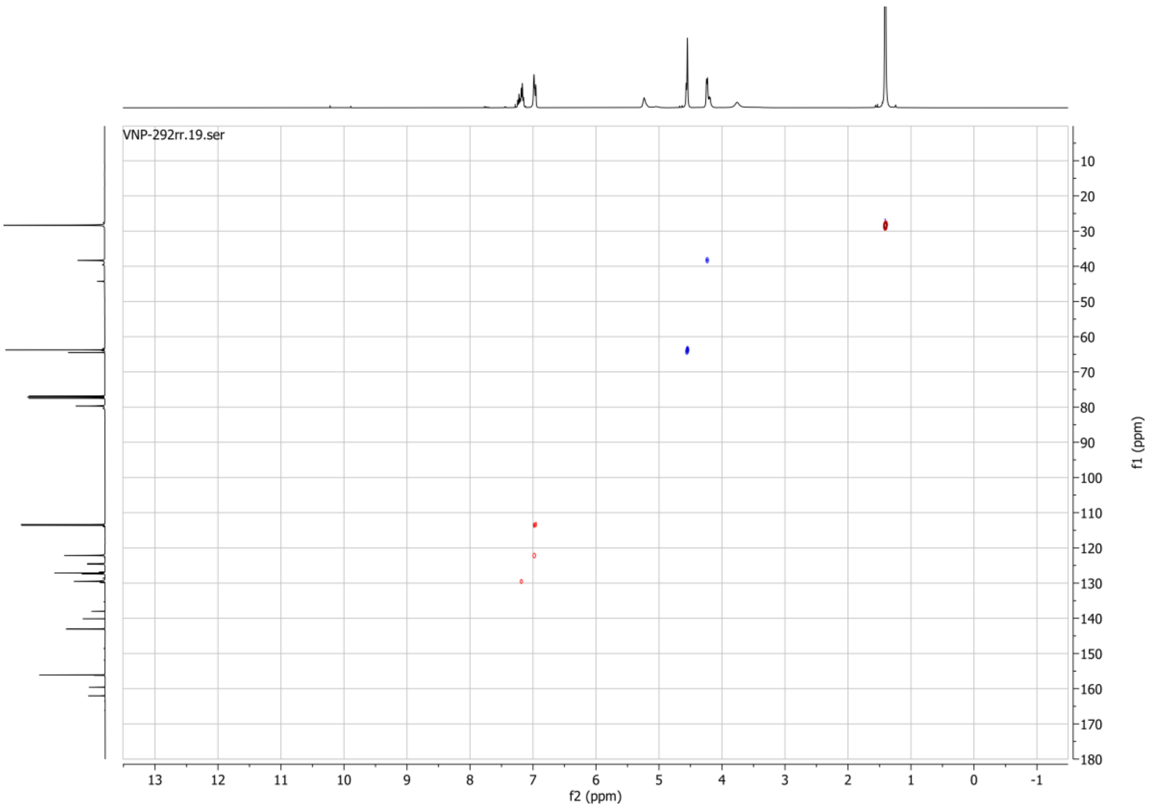

**1H spectrum of compound 68:**

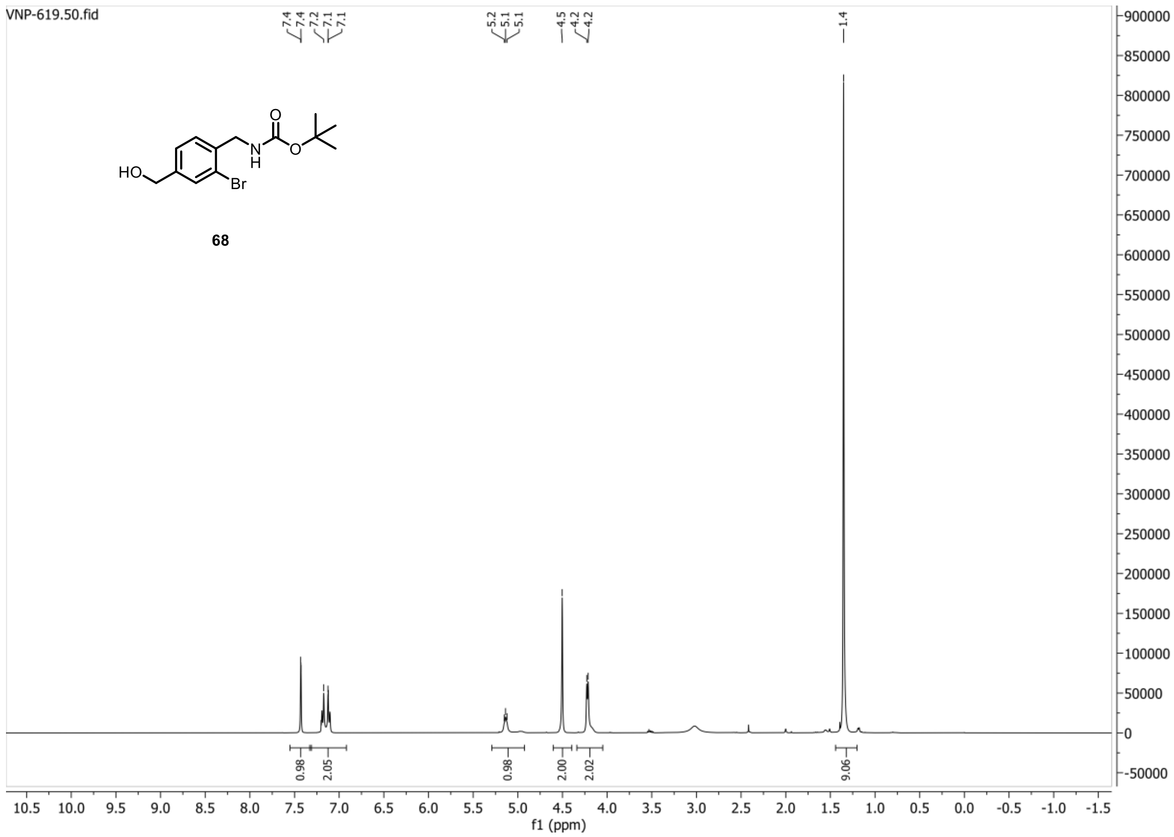


**13C{1H} spectrum of compound 68:**

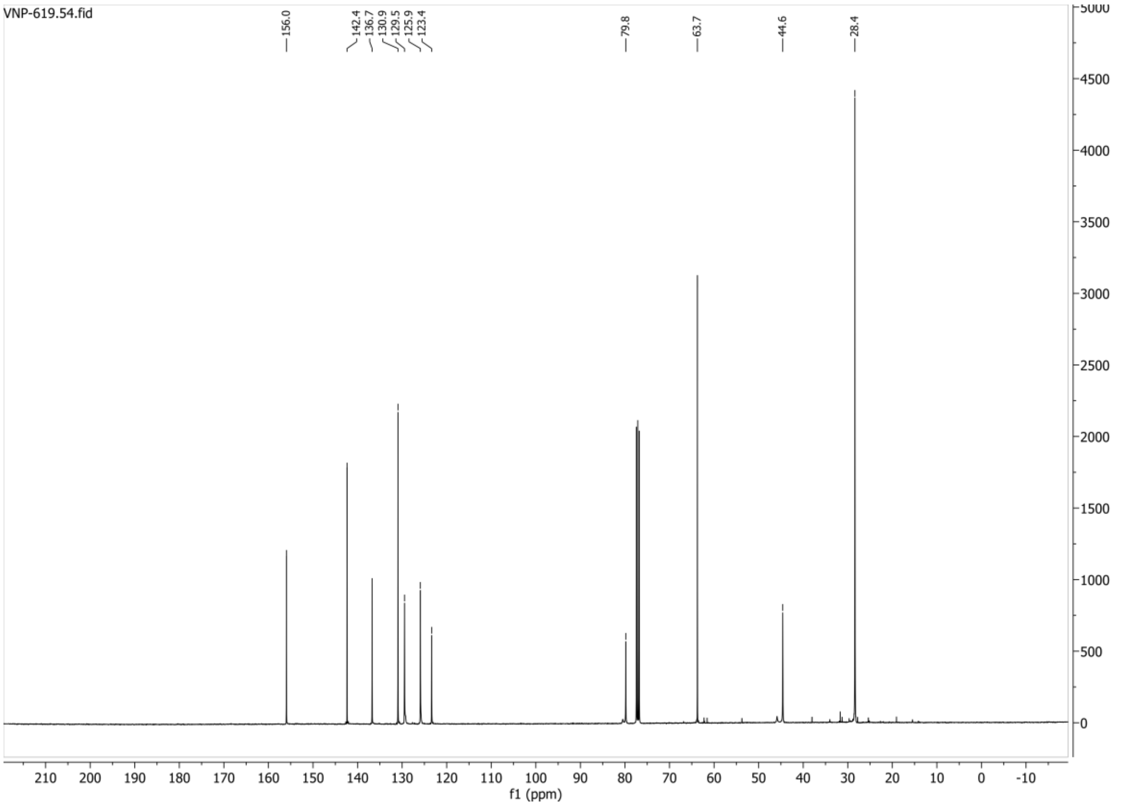

**HSQC-DEPT spectrum of compound 68:**

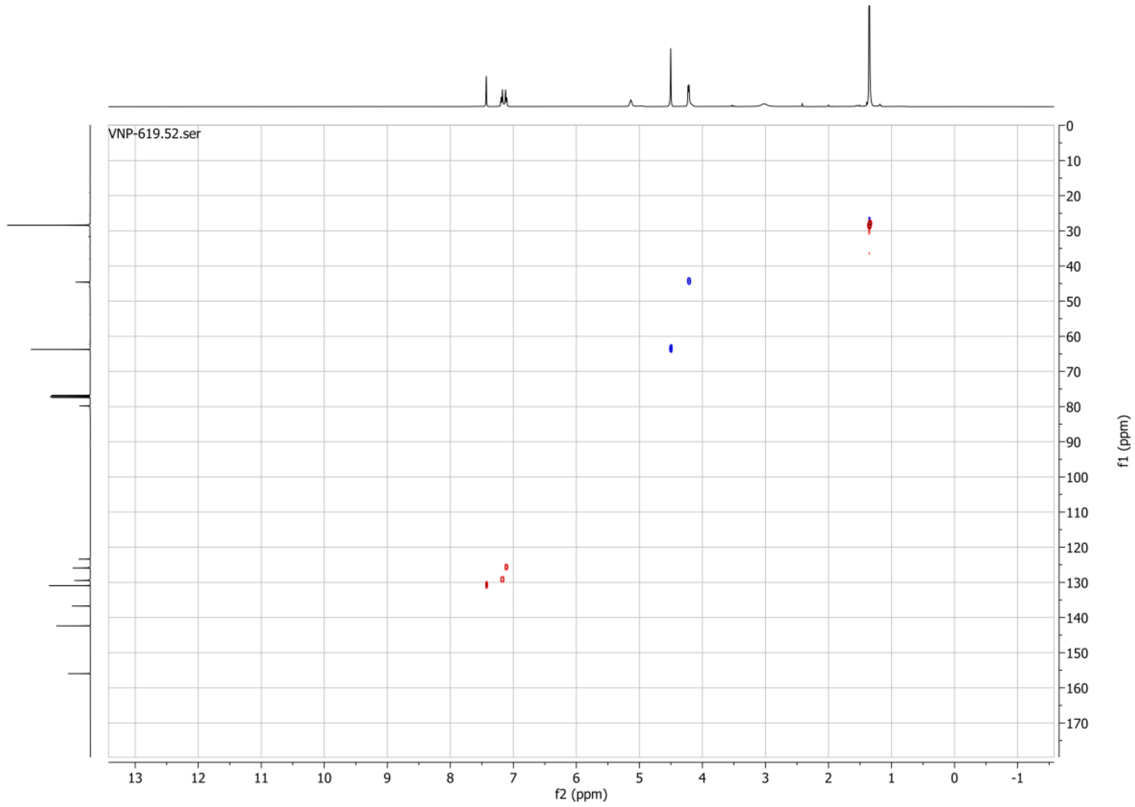

**1H spectrum of compound 71:**

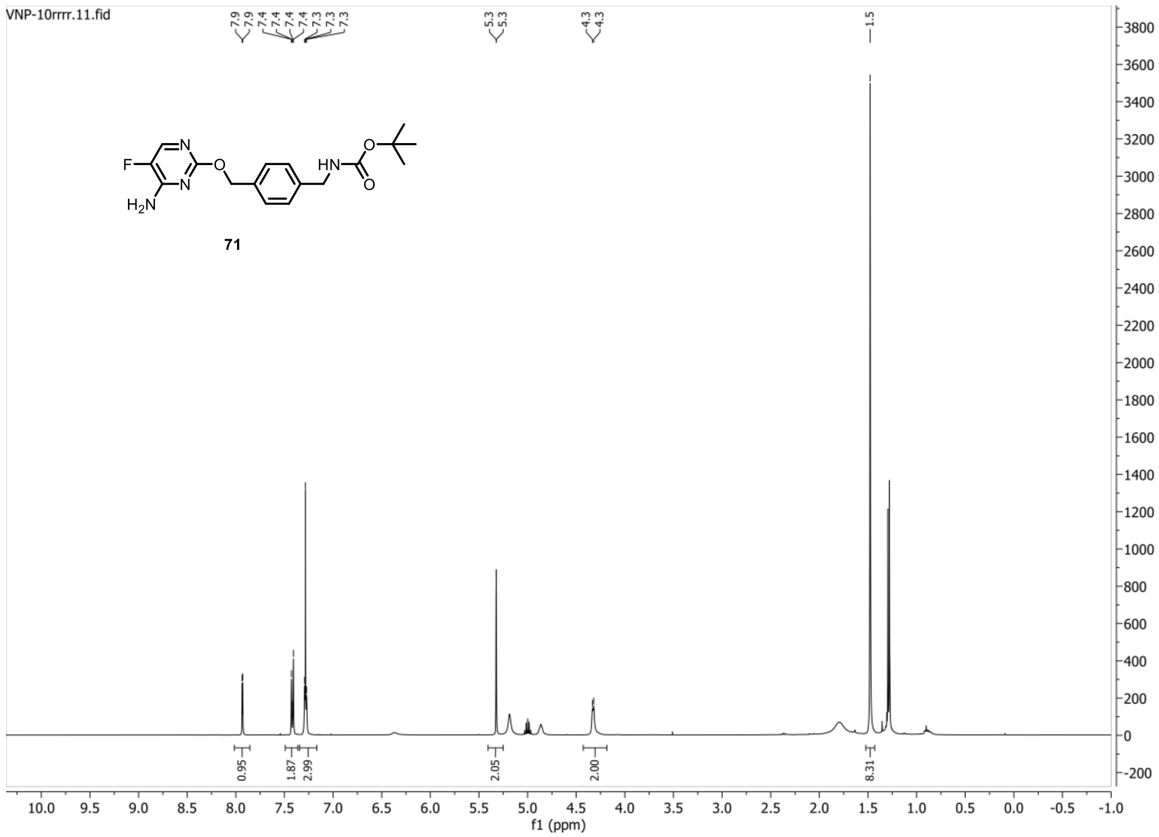


**13C{1H} spectrum of compound 71:**

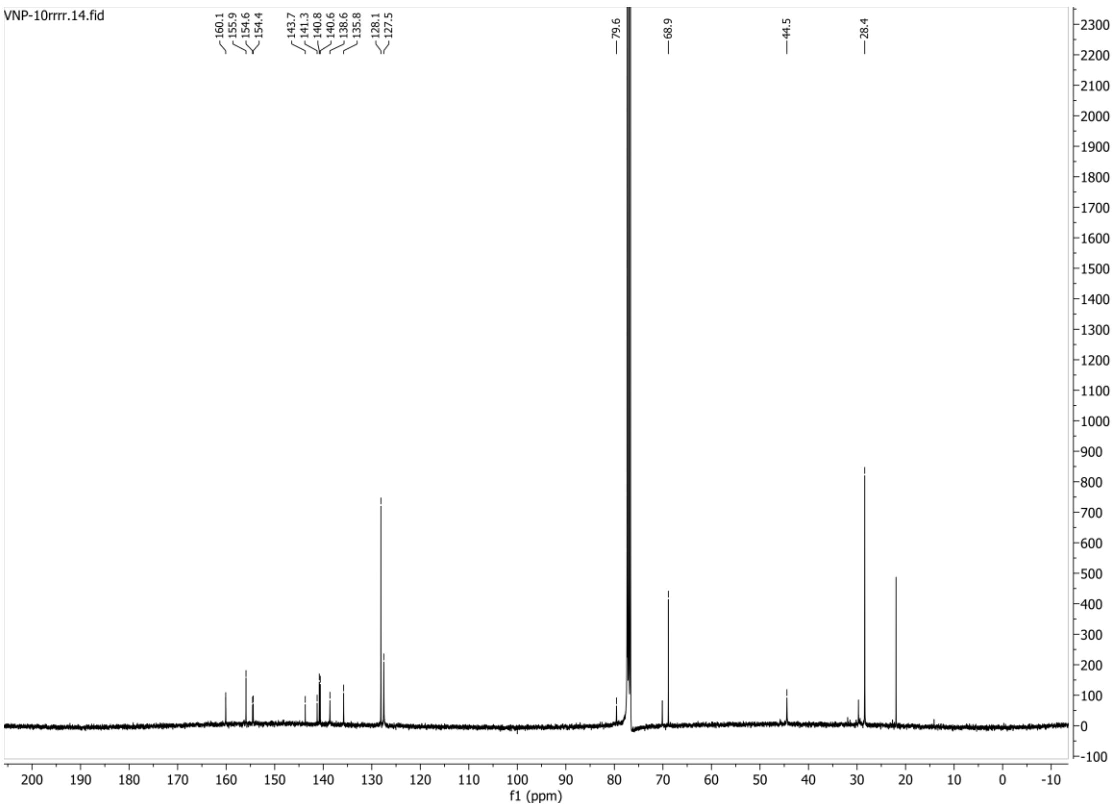

**HSQC-DEPT135 spectrum of compound 71:**

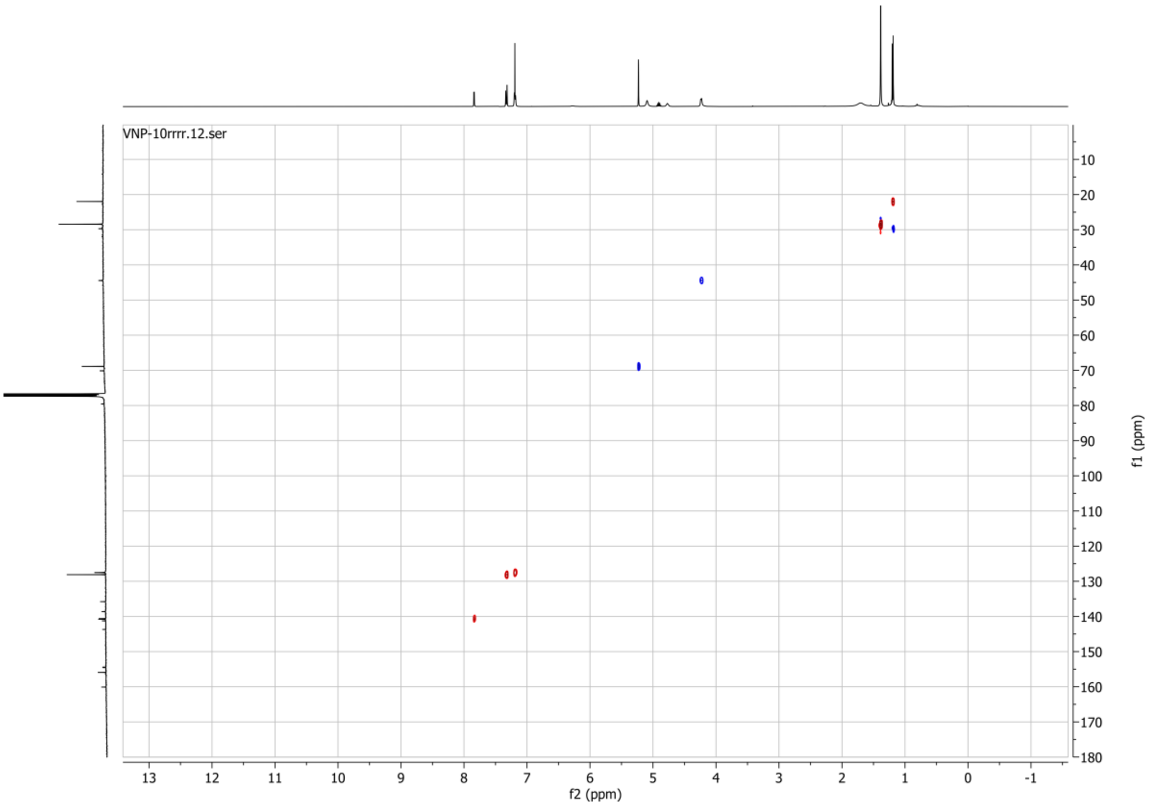


**19F spectrum of compound 71:**

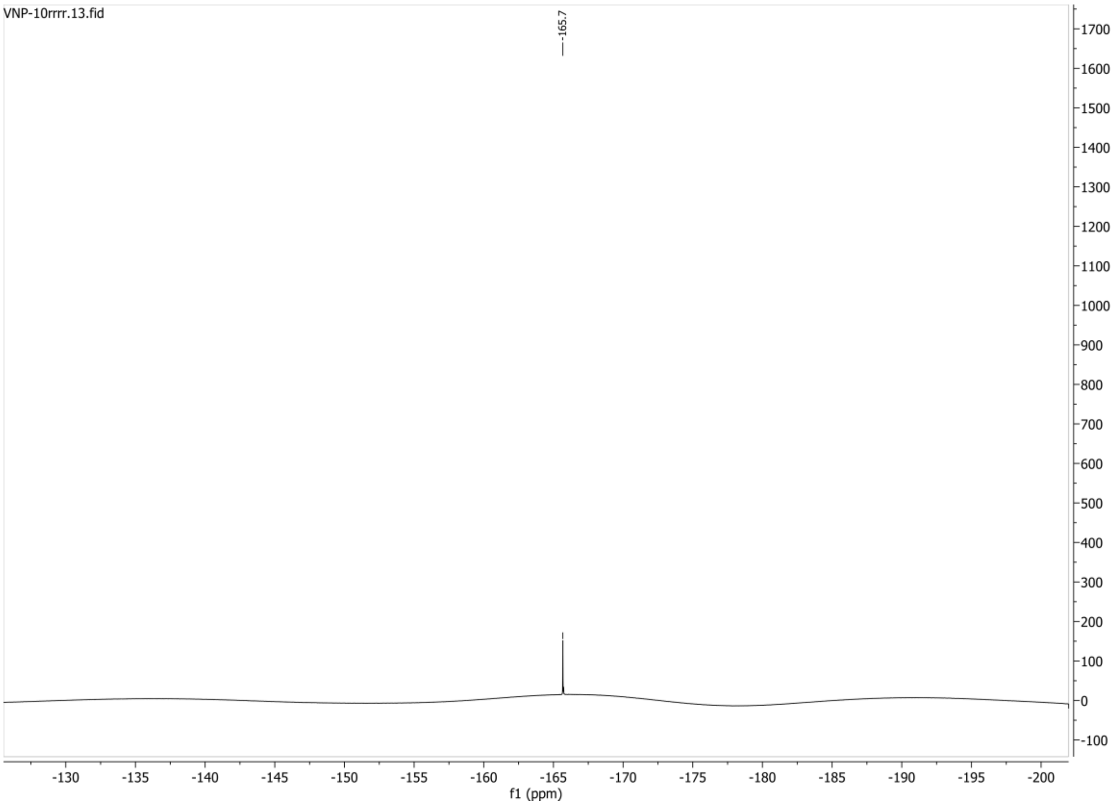

**1H spectrum of compound 75:**

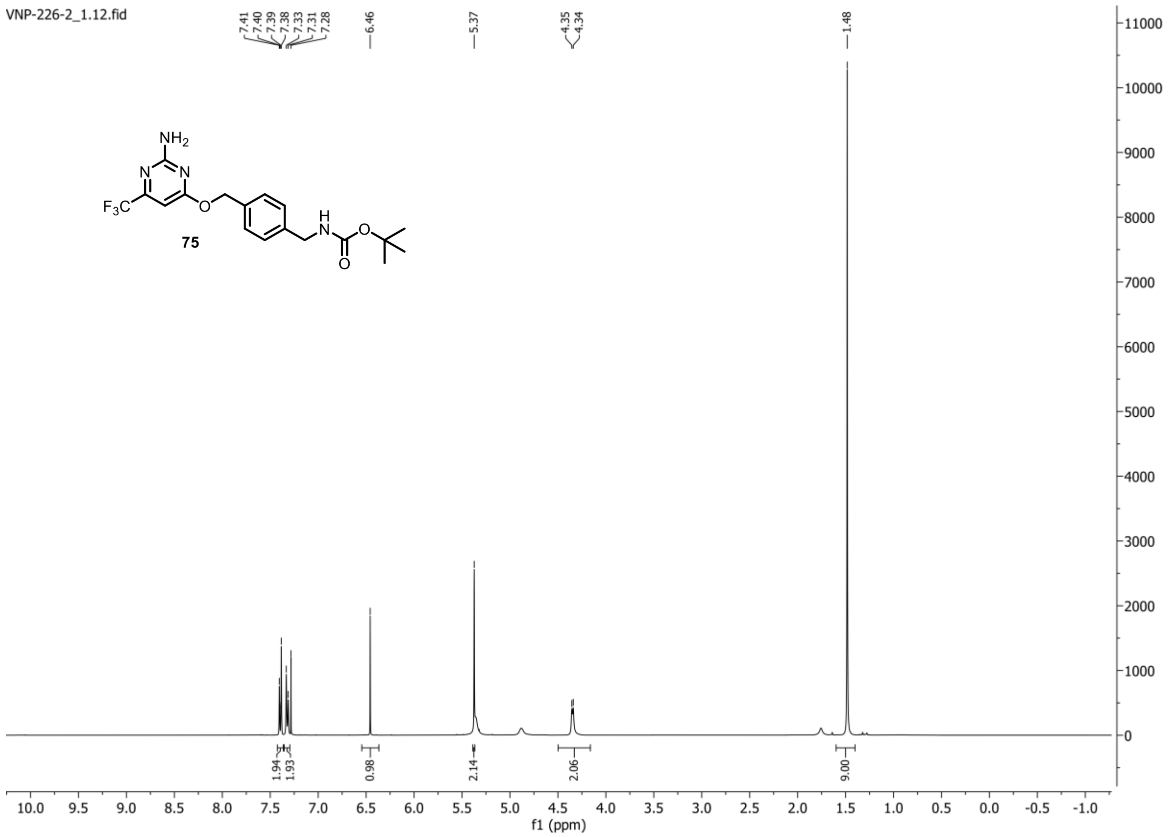


**13C{1H} spectrum of compound 75:**

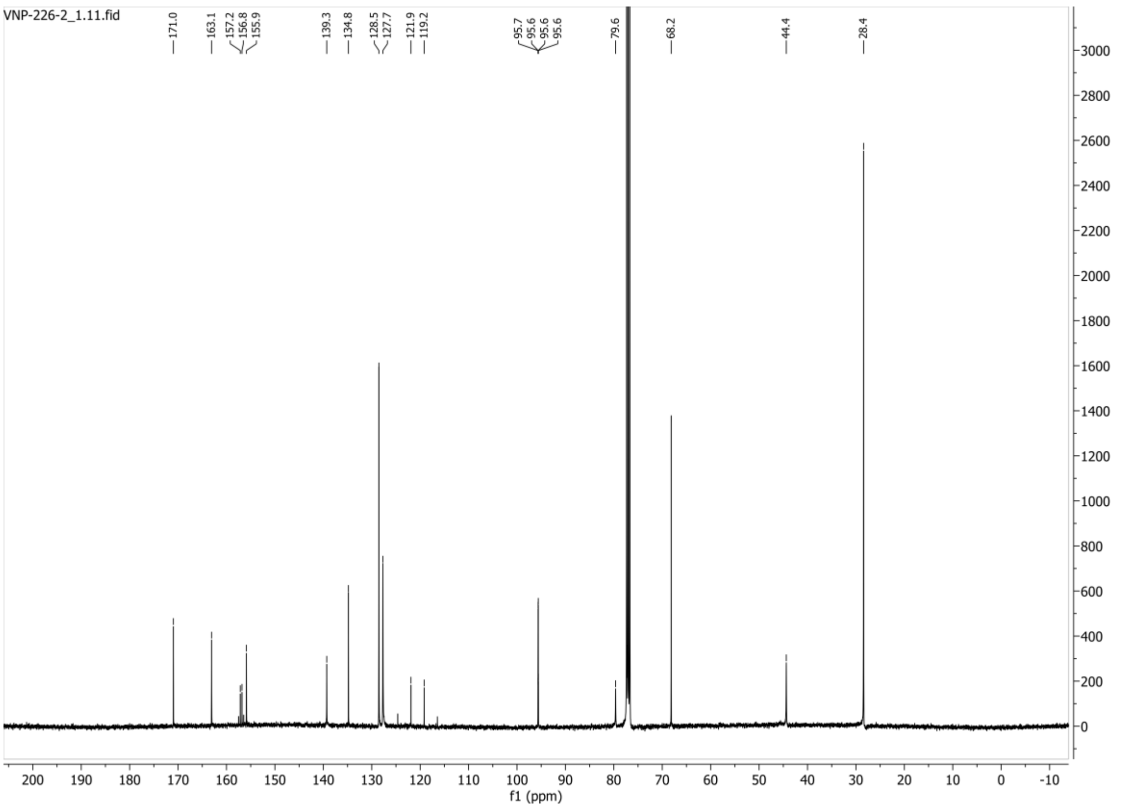

**DEPT-135 spectrum of compound 75:**

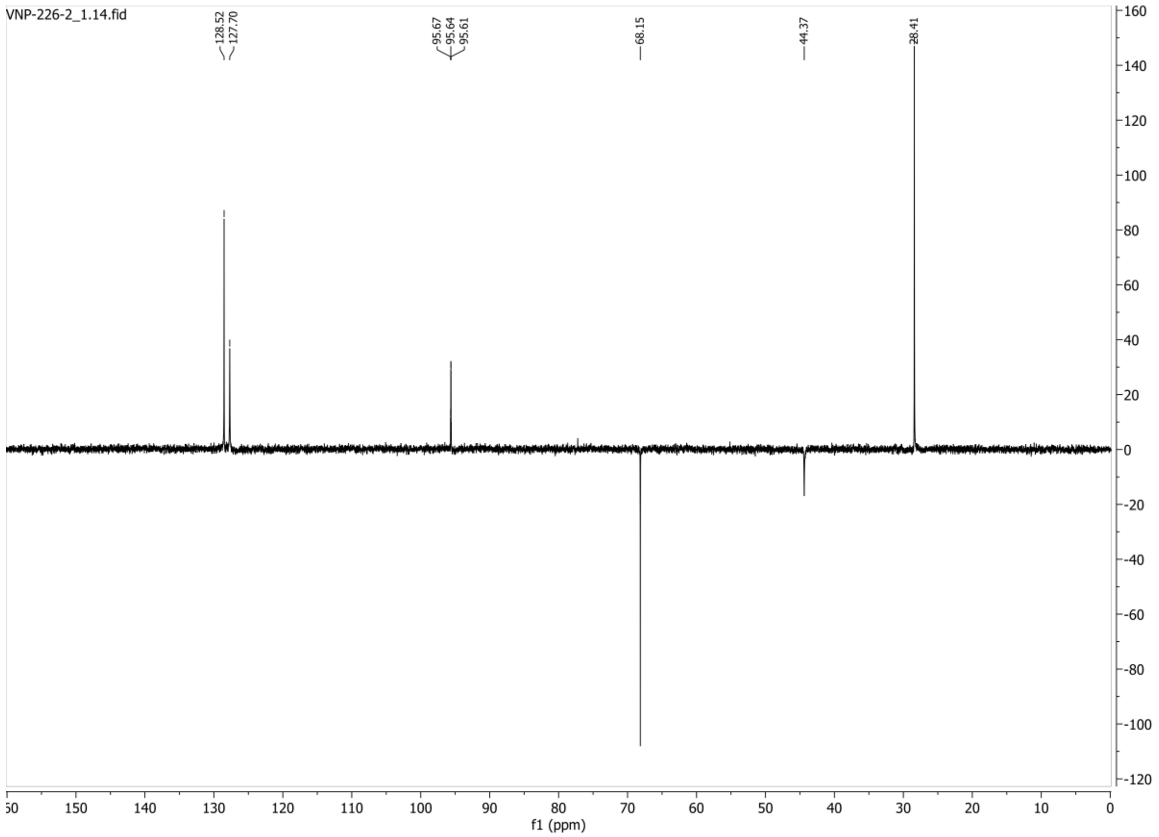


**19F spectrum of compound 75:**

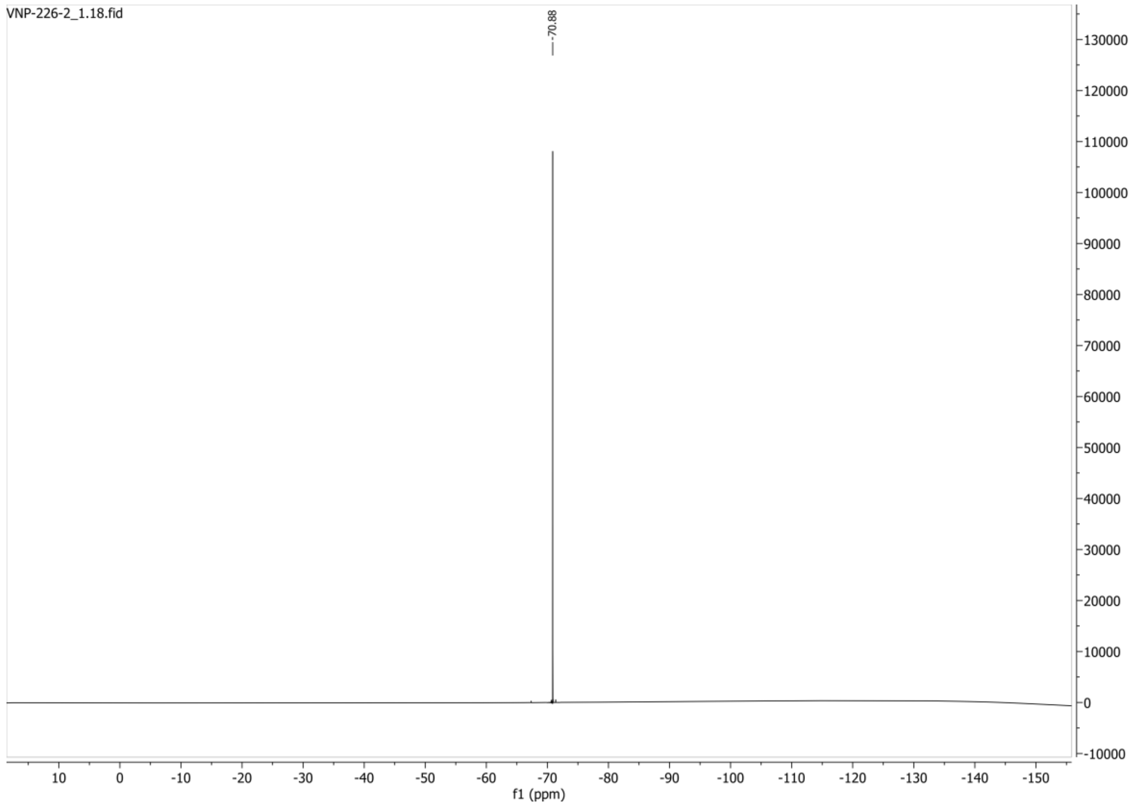

**1H spectrum of compound 79:**

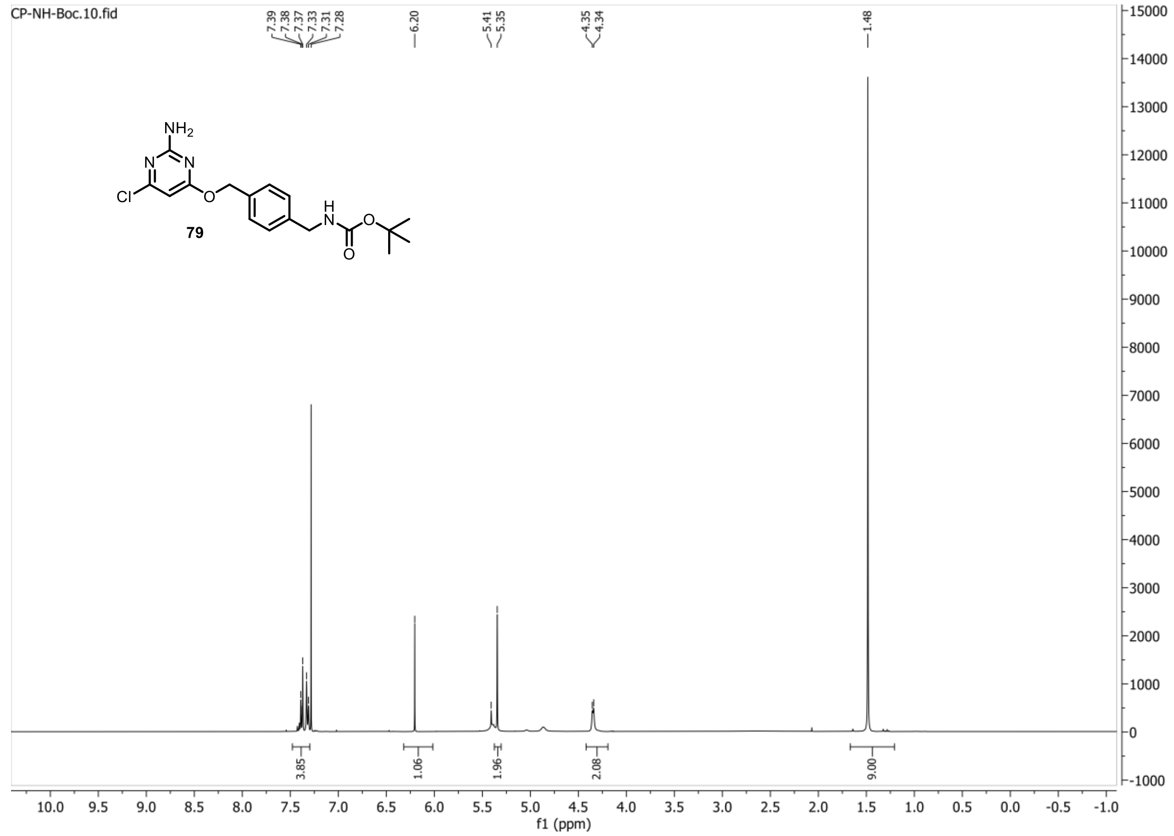


**13C{1H} spectrum of compound 79:**

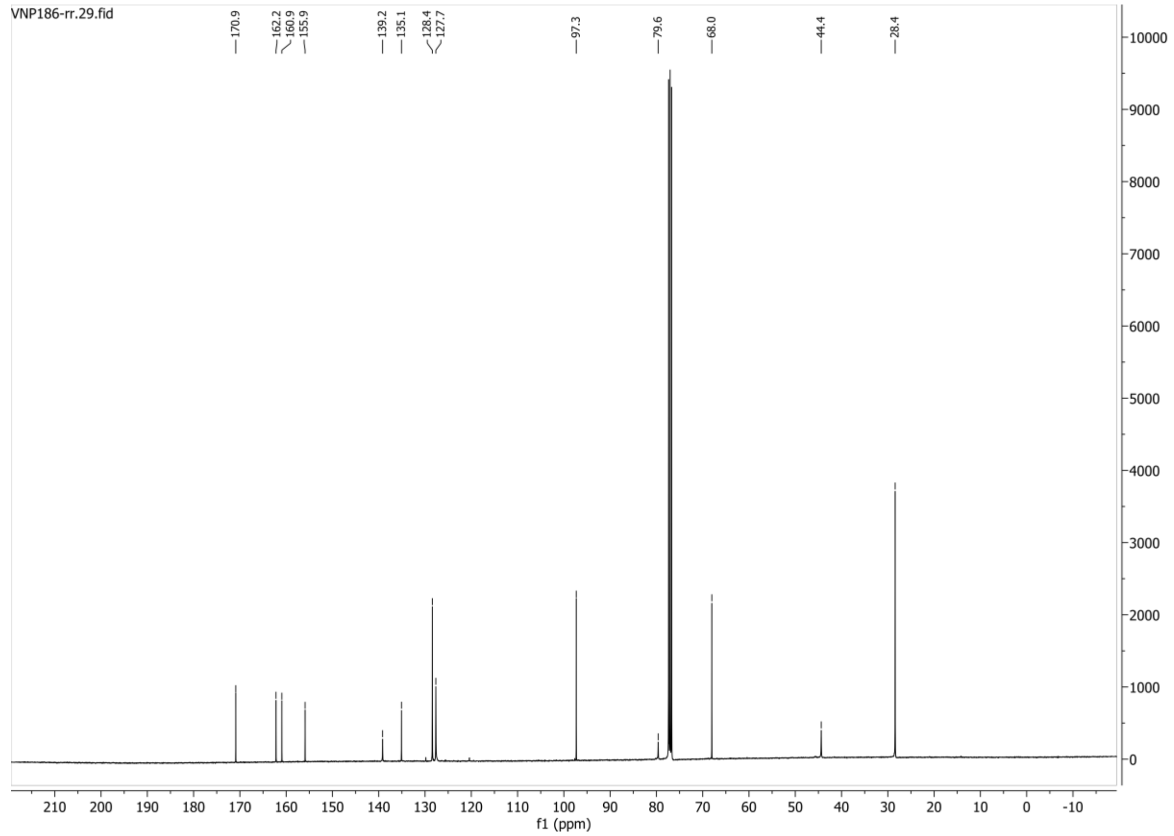

**HSQC-DEPT135 spectrum of compound 79:**

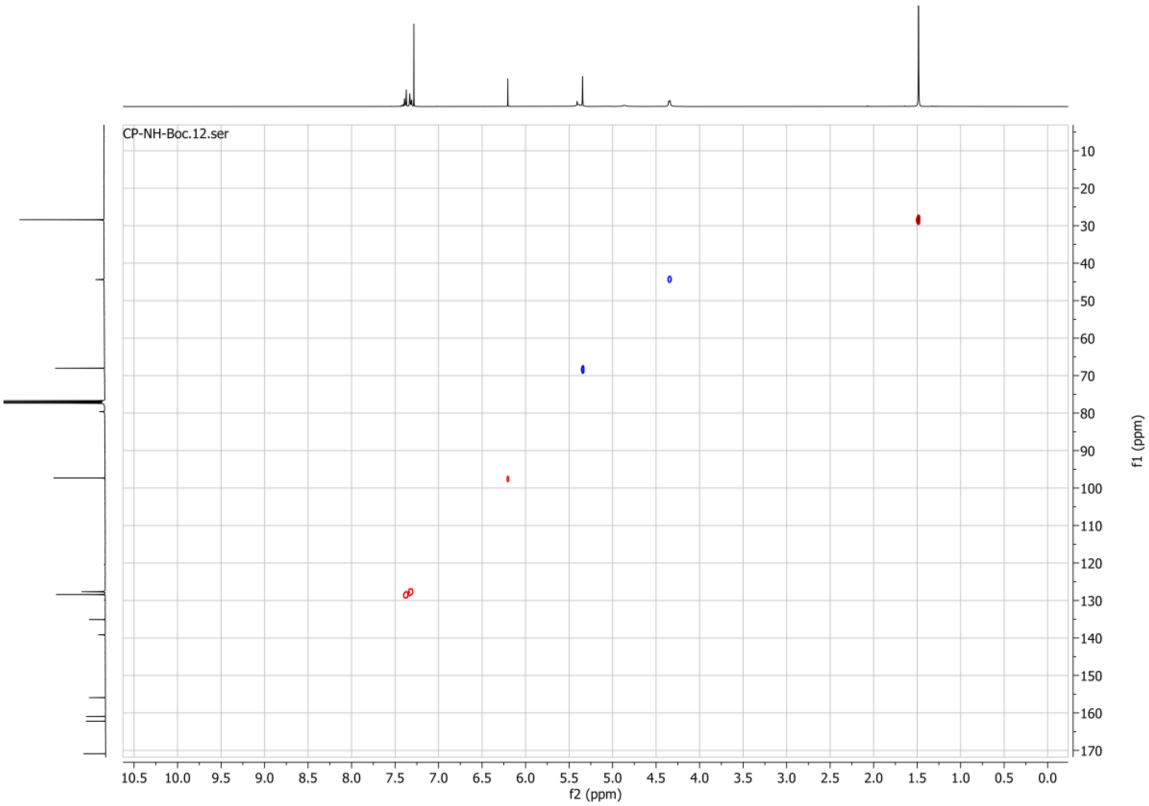

**1H spectrum of compound 83:**

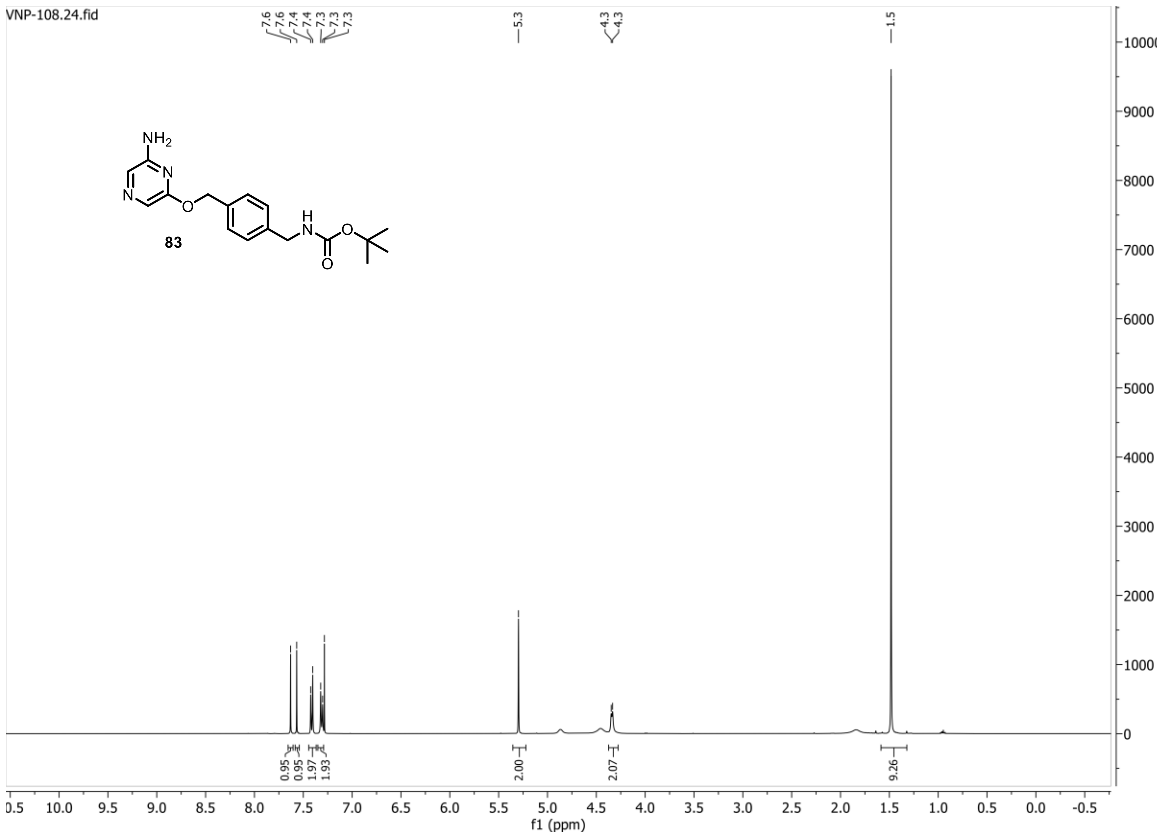


**13C{1H} spectrum of compound 83:**

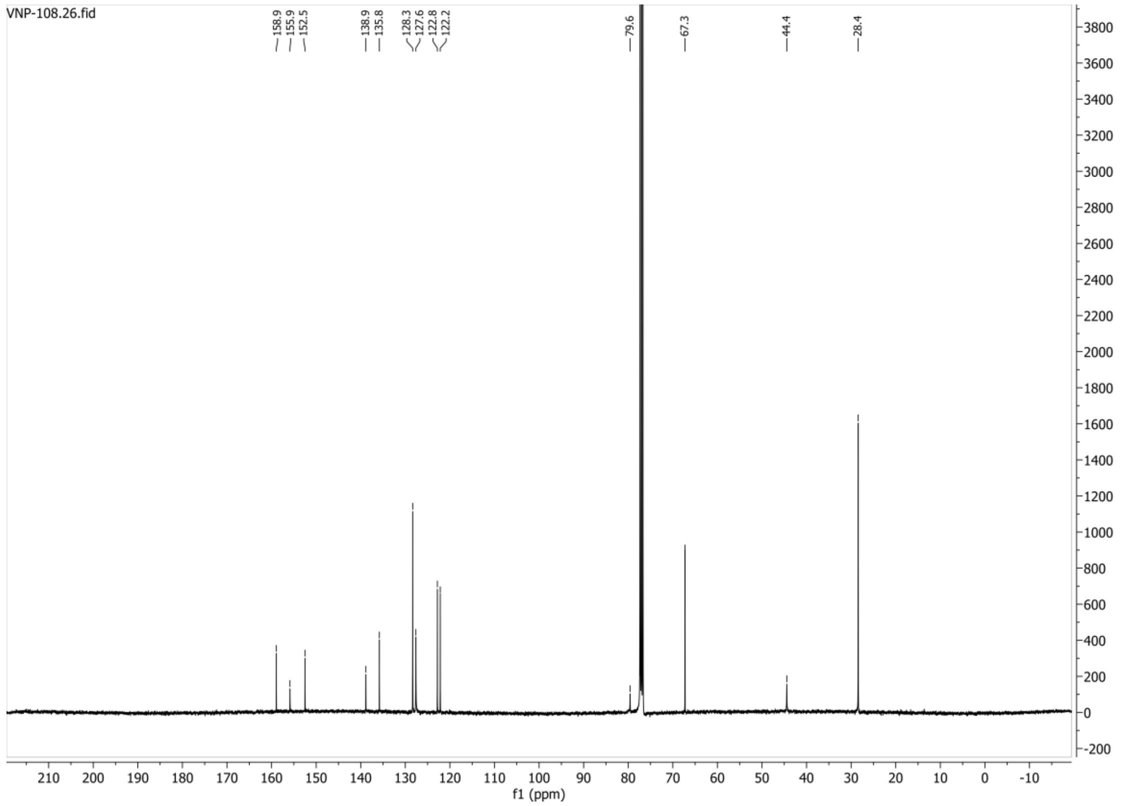

**HSQC-DEPT135 spectrum of compound 83:**

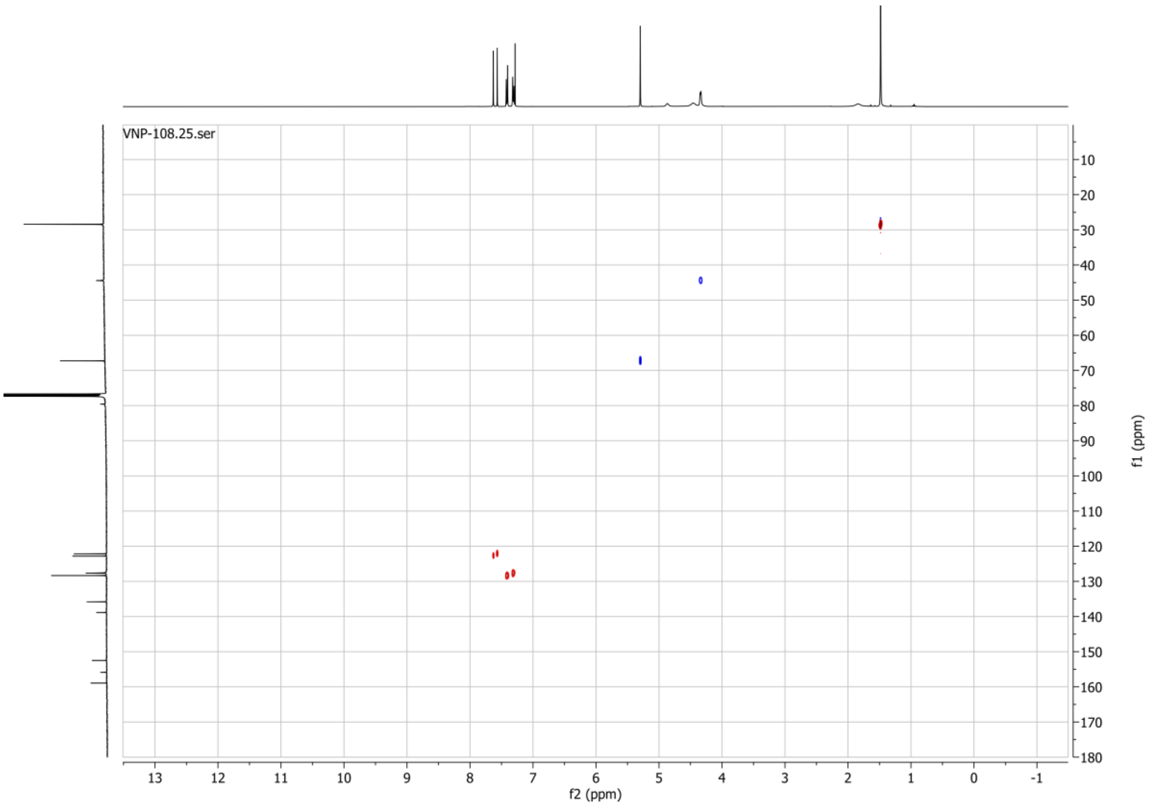

**1H spectrum of compound 87:**

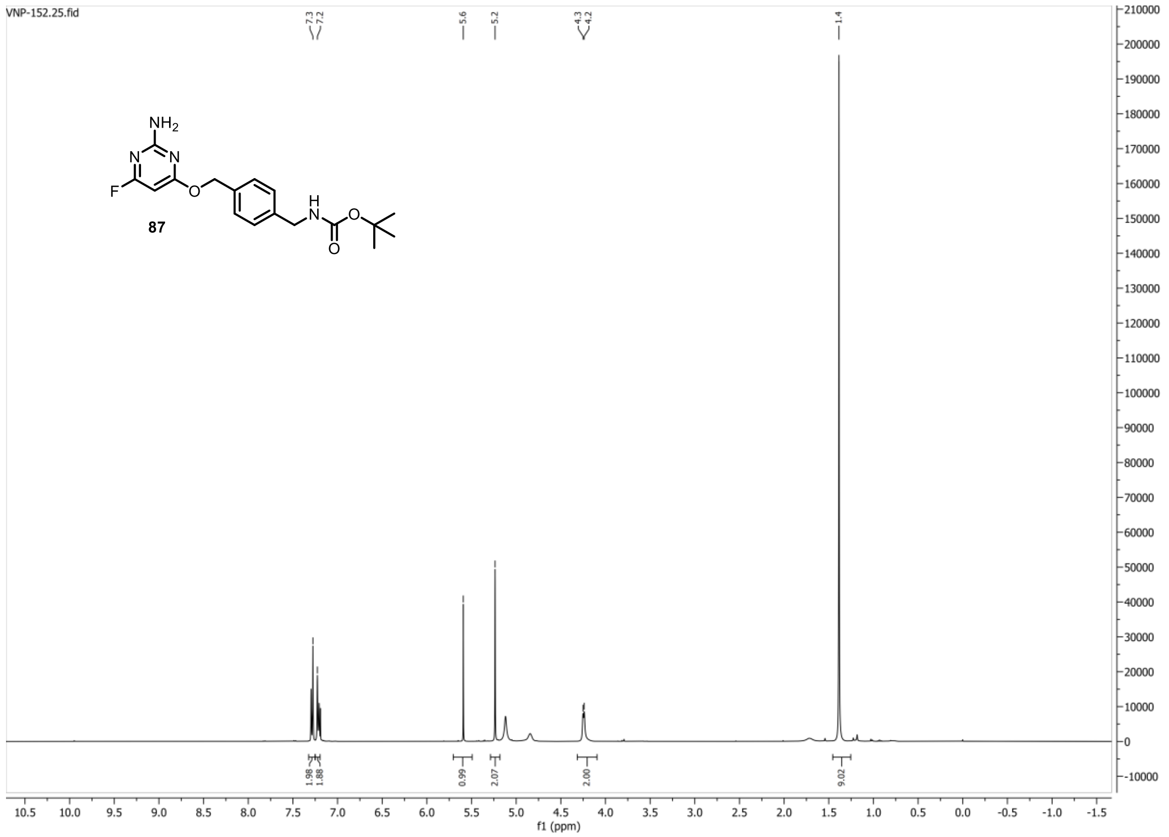


**13C{1H} spectrum of compound 87:**

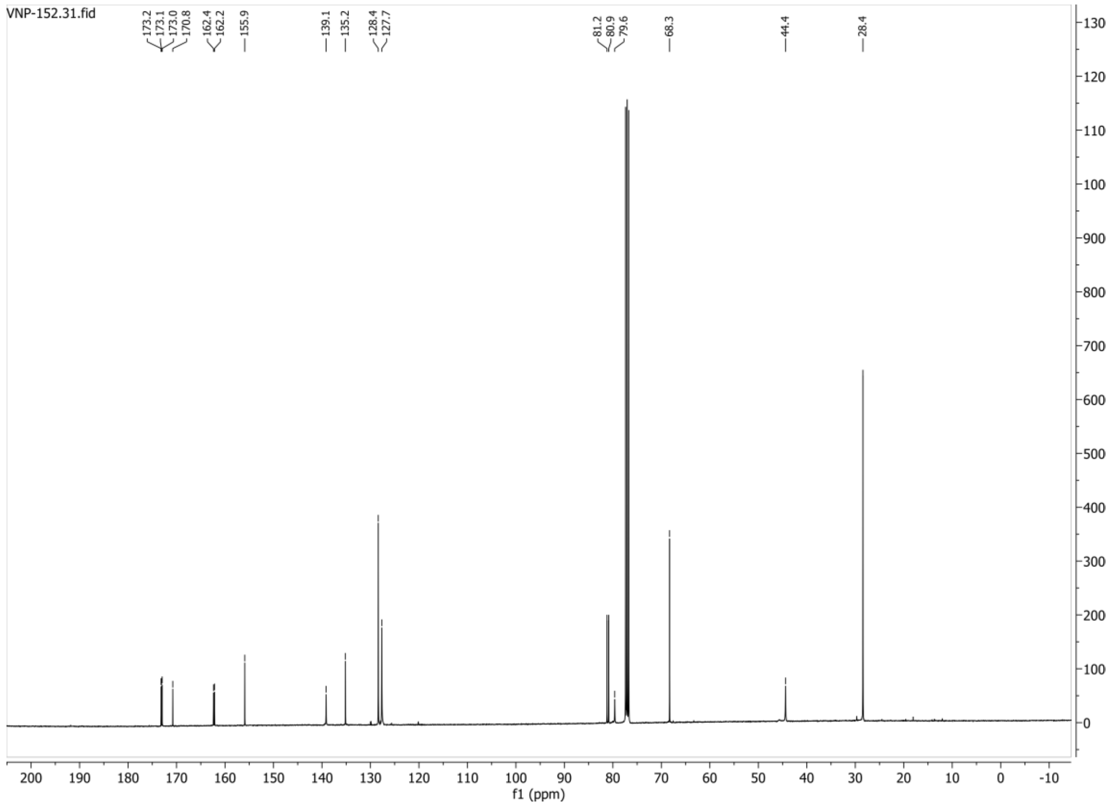

**HSQC-DEPT135 spectrum of compound 87:**

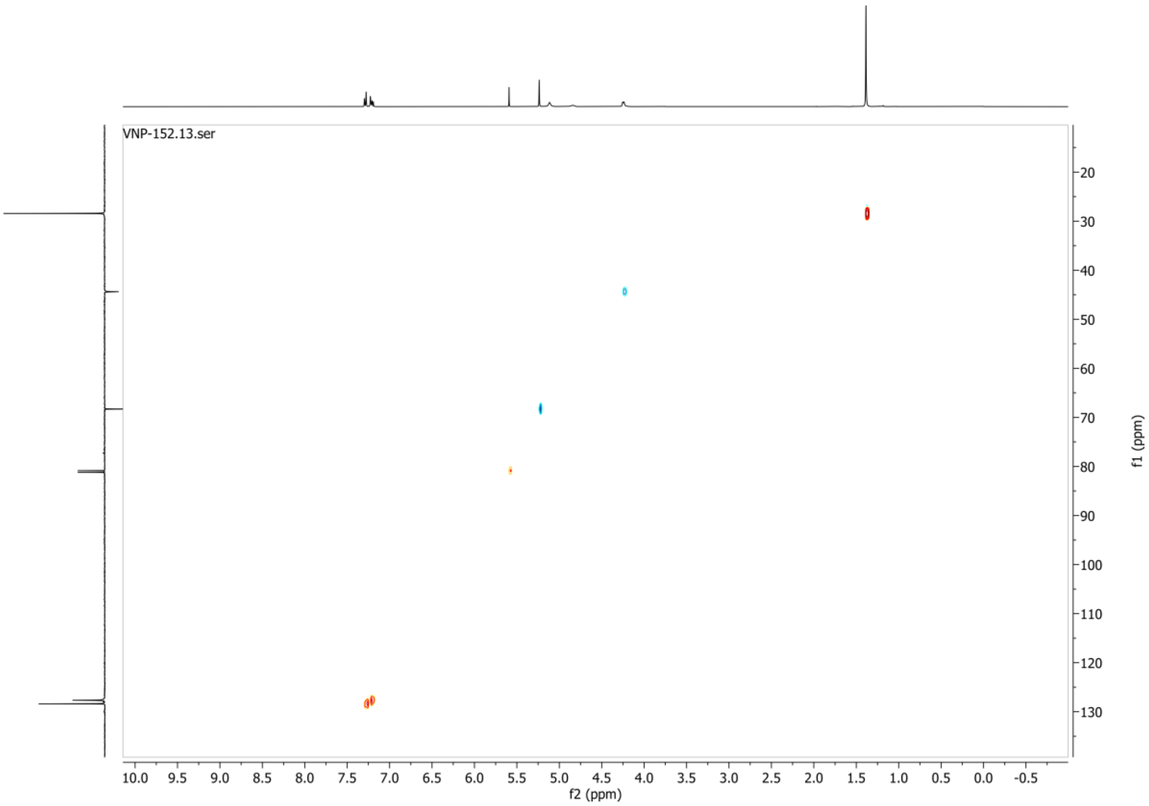


**19F spectrum of compound 87:**

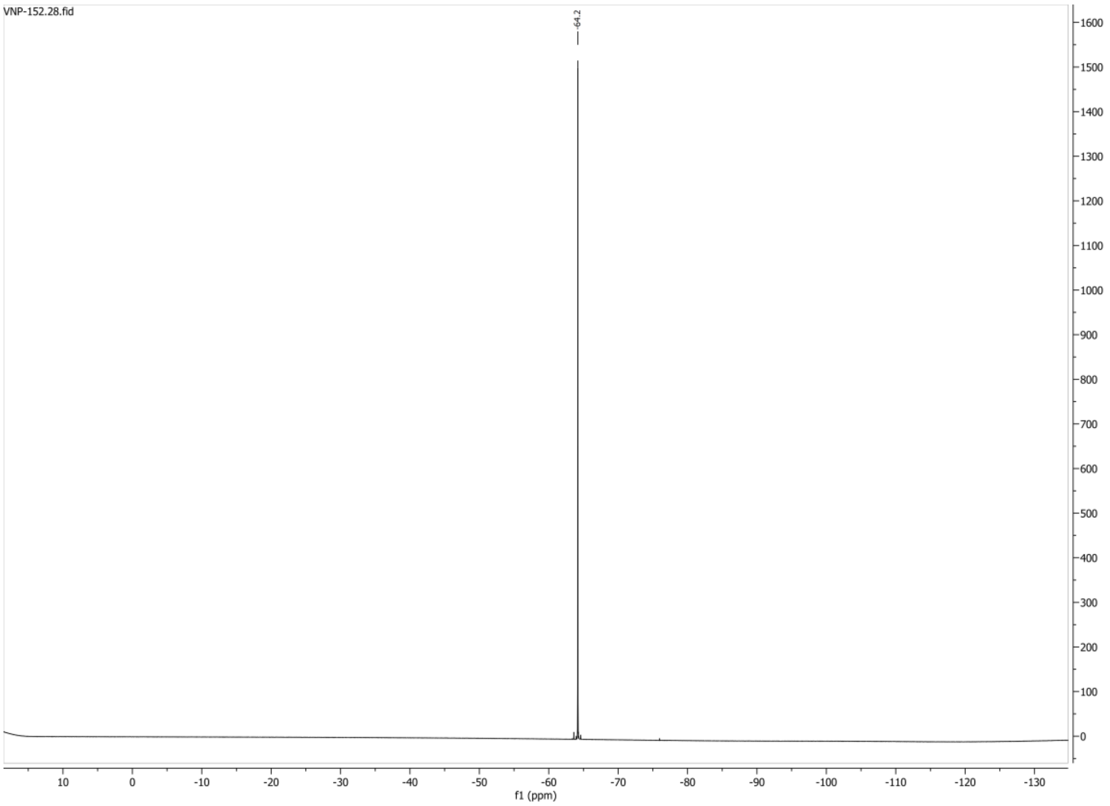

**1H spectrum of compound 91:**

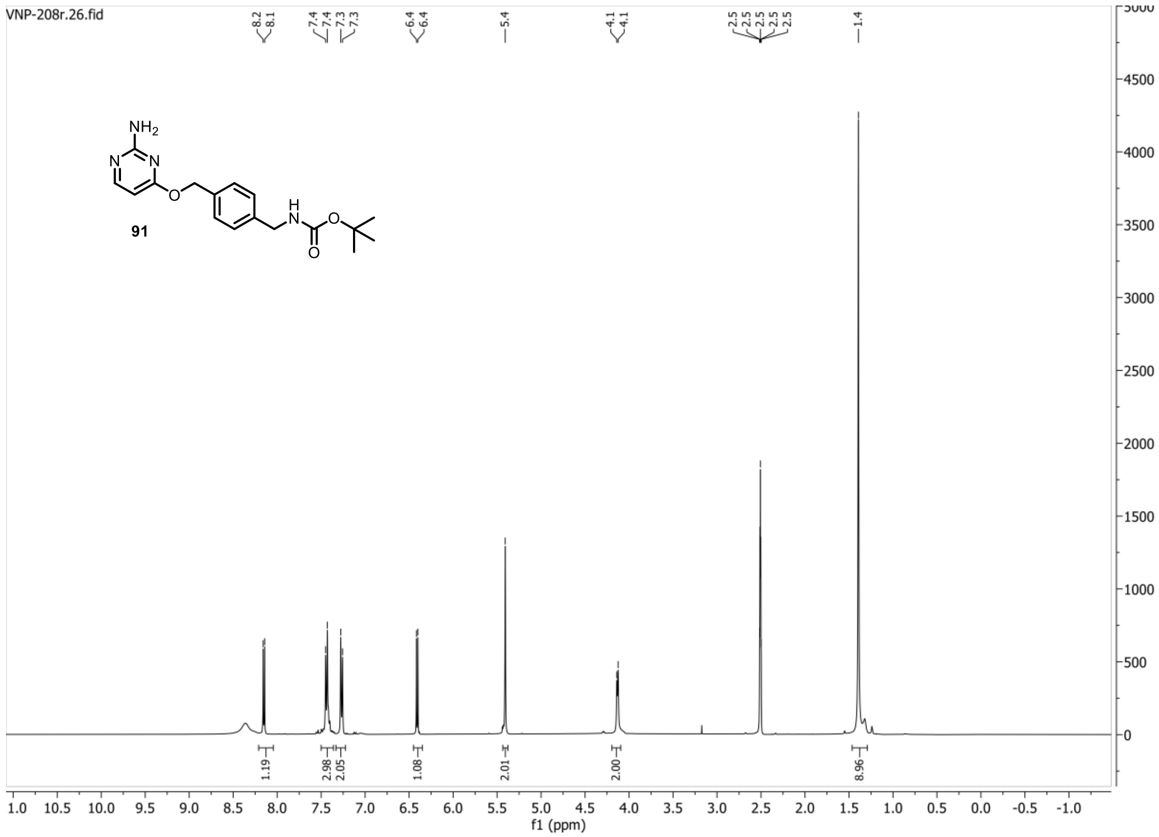


**13C{1H} spectrum of compound 91:**

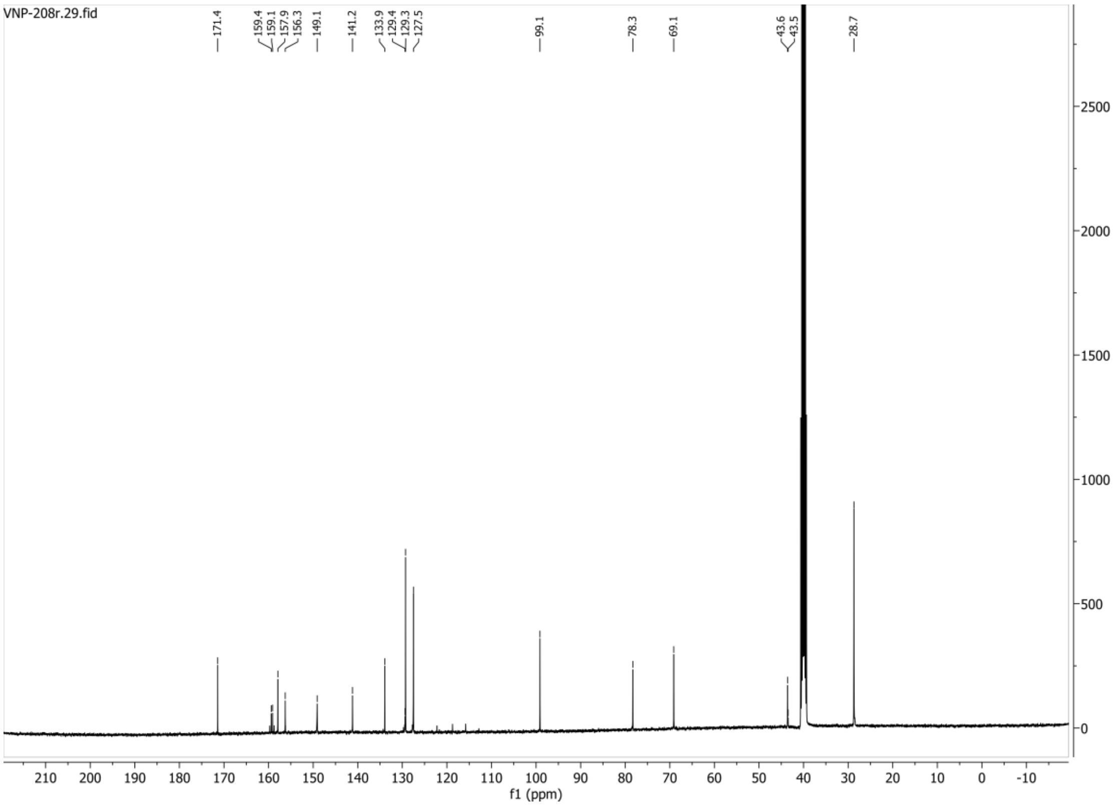

**HSQC-DEPT135 spectrum of compound 91:**

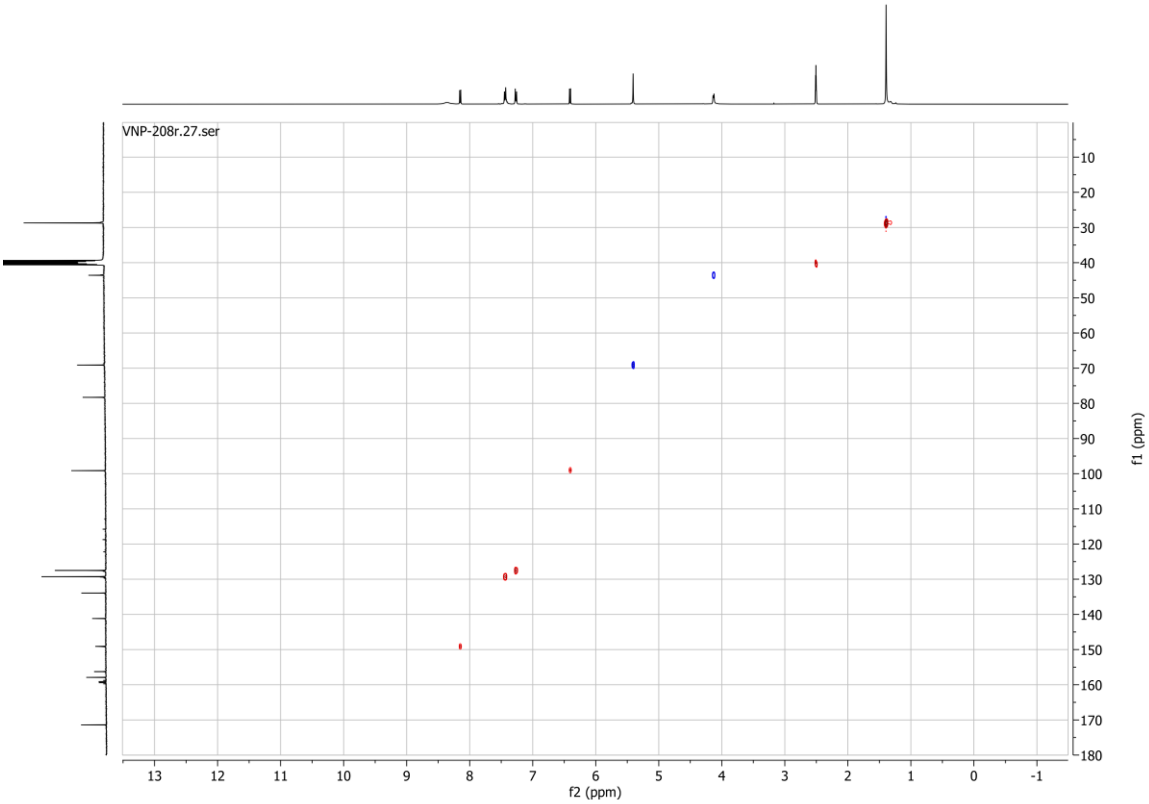

**1H spectrum of compound 95:**

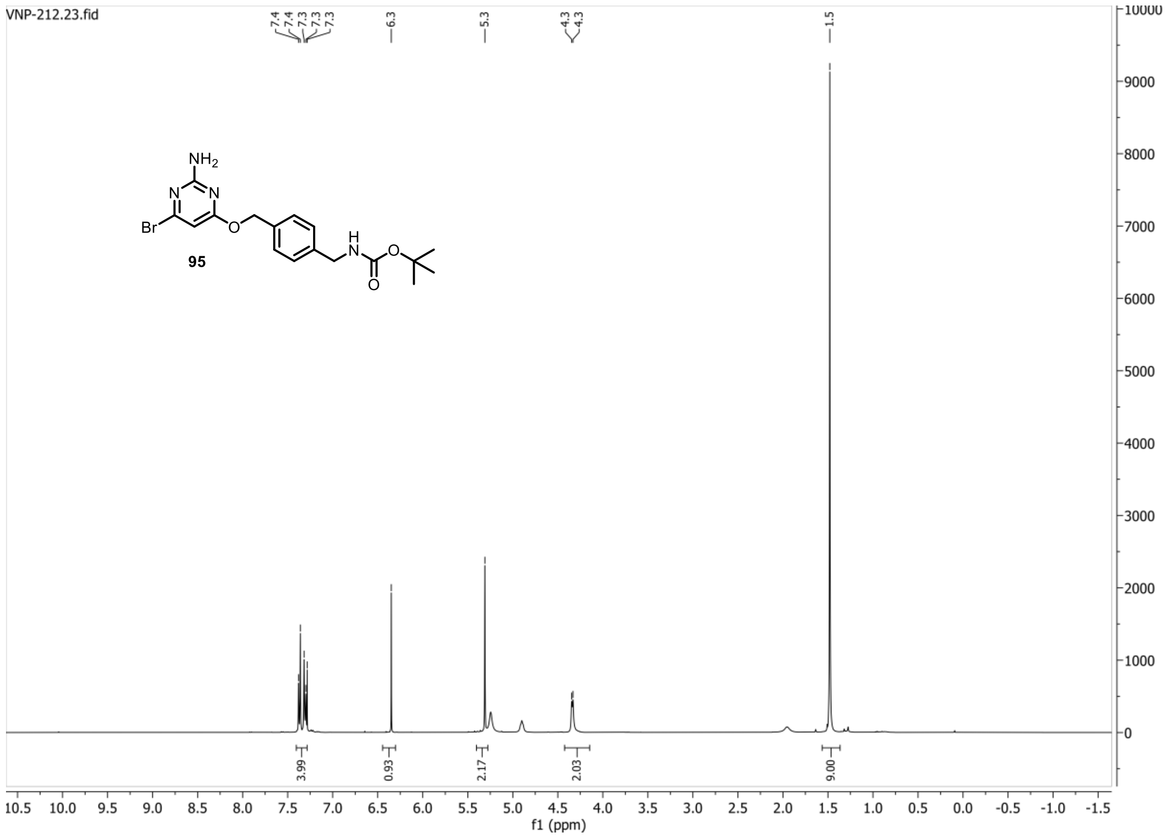


**13C{1H} spectrum of compound 95:**

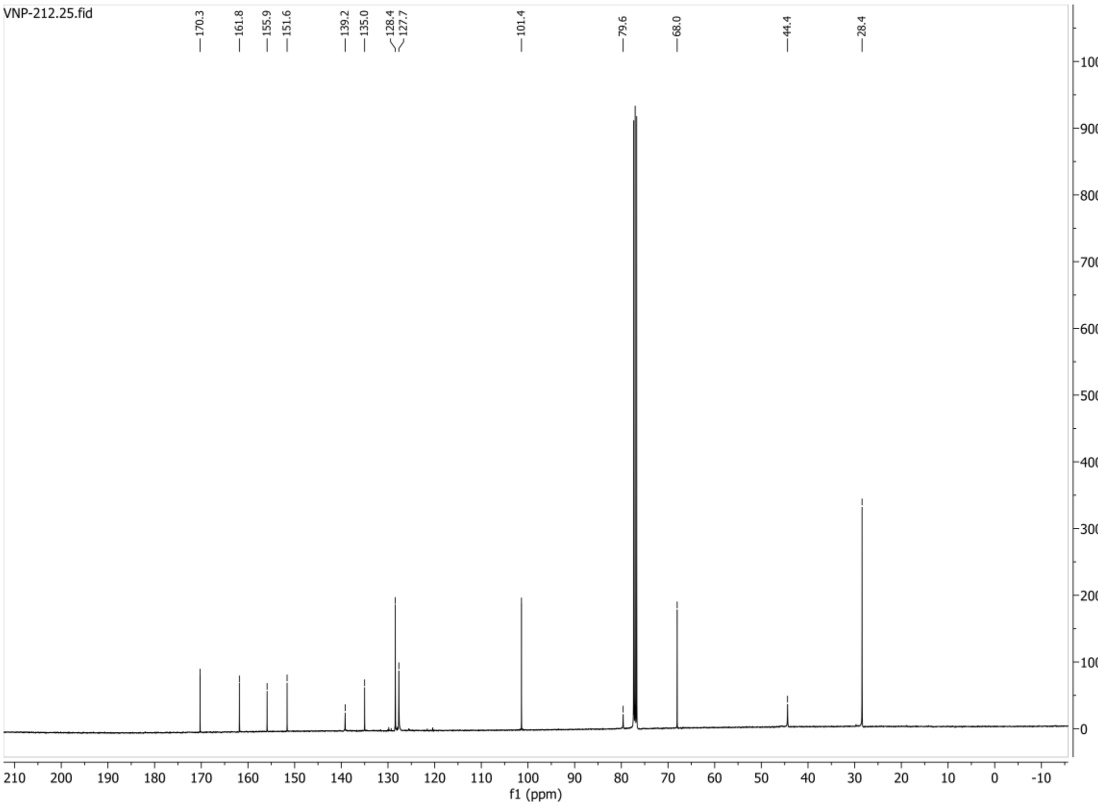

**HSQC-DEPT135 spectrum of compound 95:**

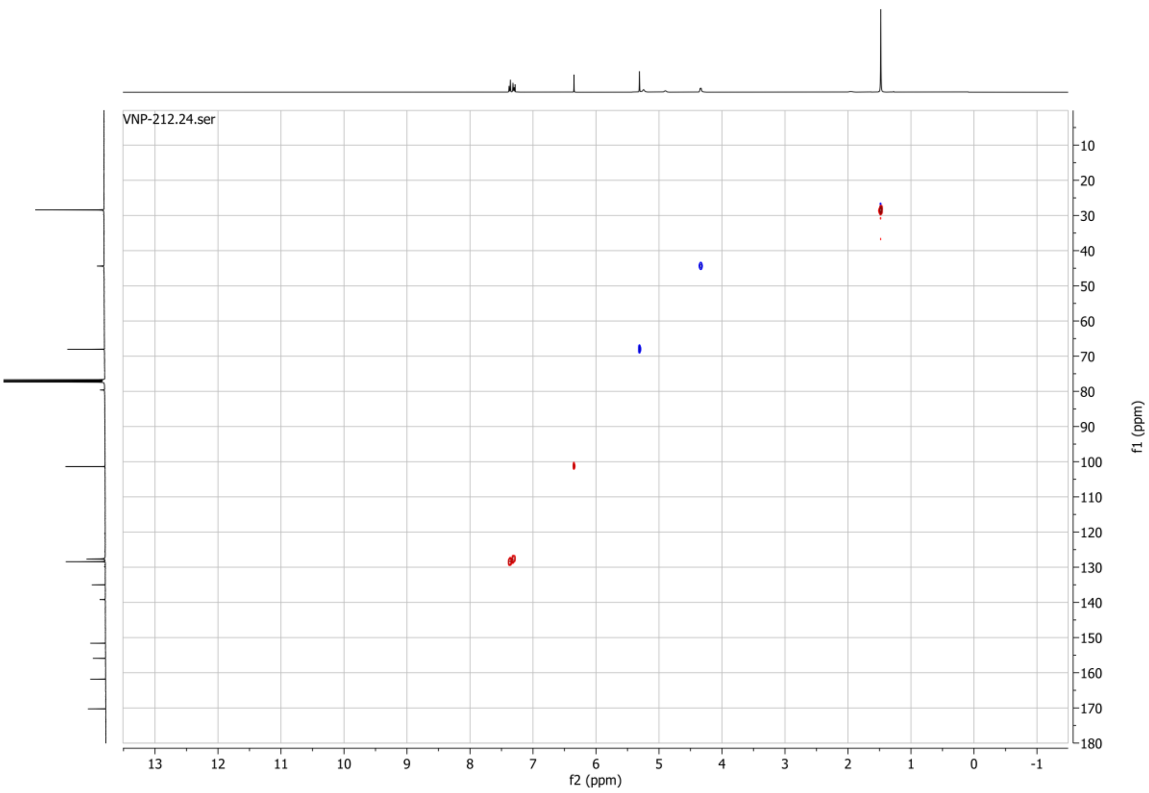

**1H spectrum of compound 99:**

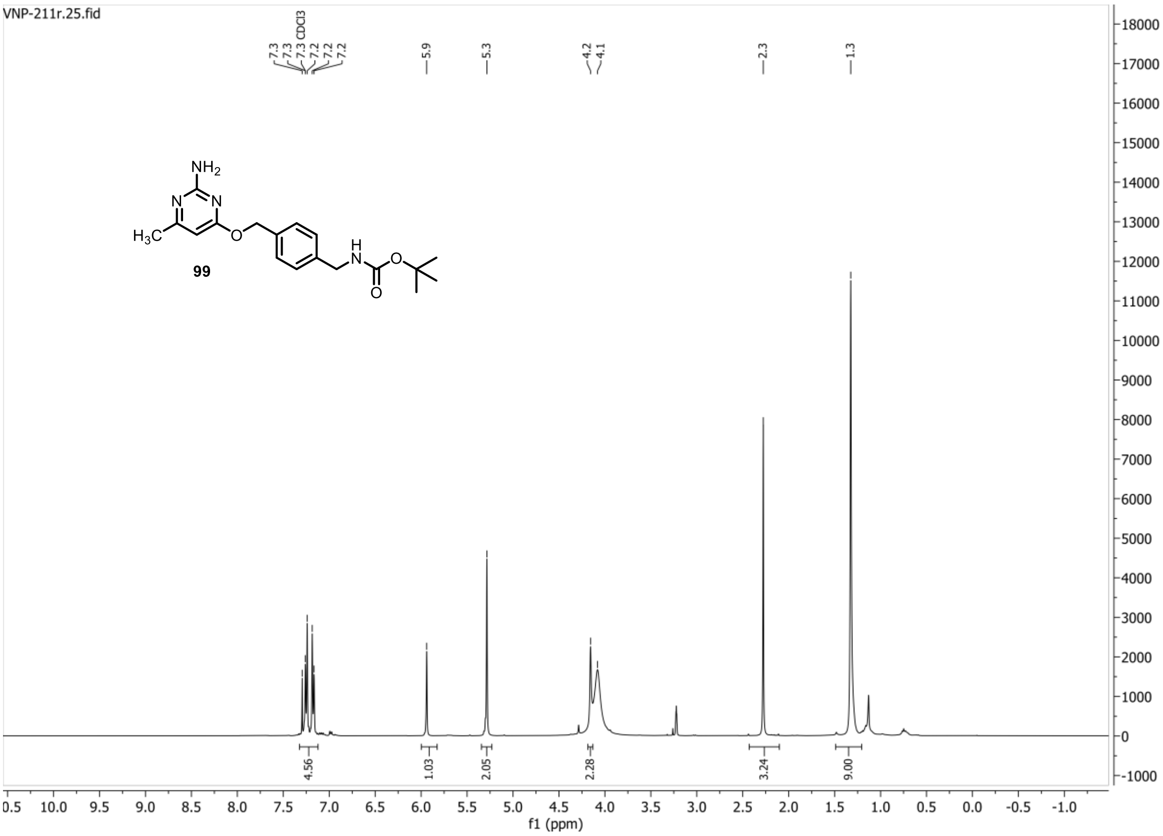


**13C{1H} spectrum of compound 99:**

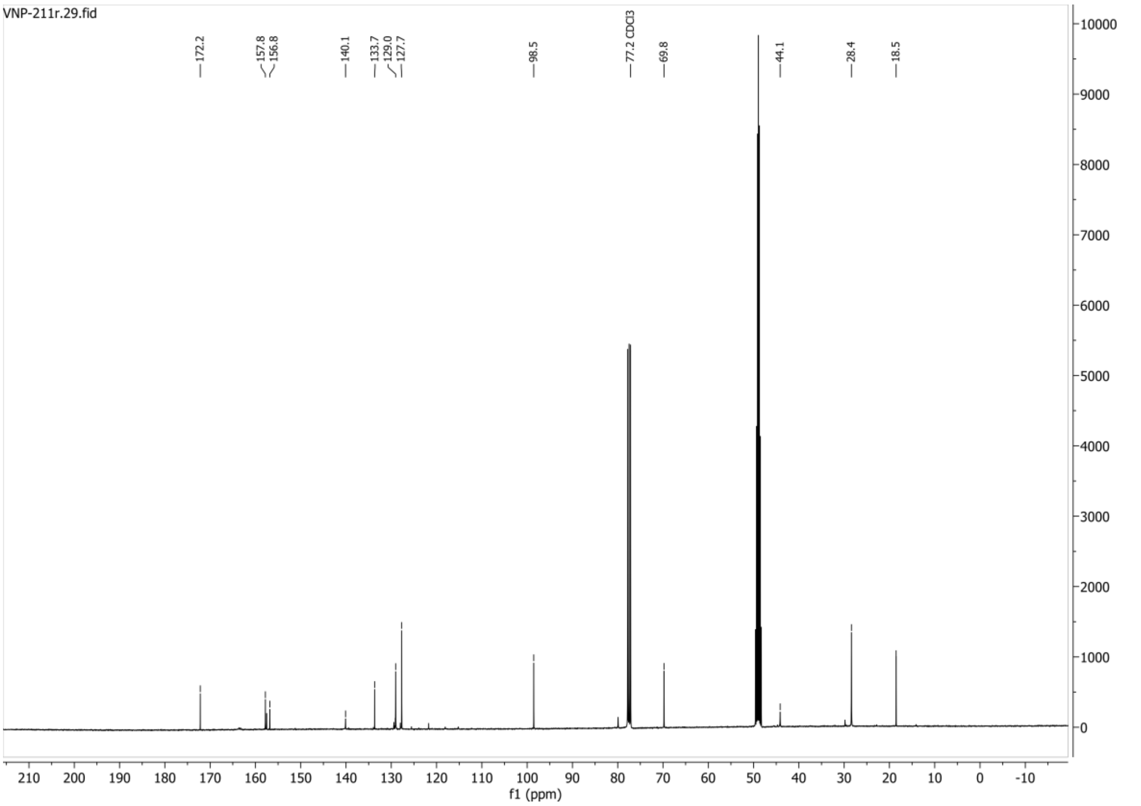

**HSQC-DEPT135 spectrum of compound 99:**

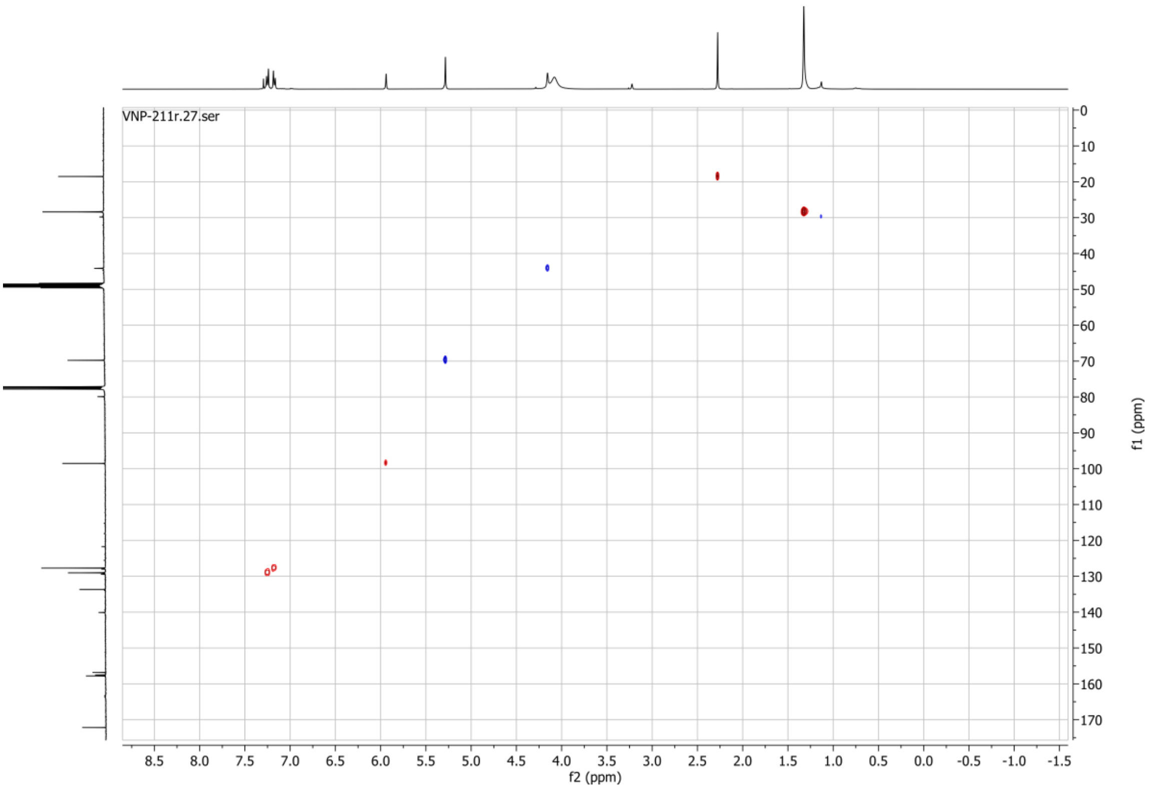

**1H spectrum of compound 103:**

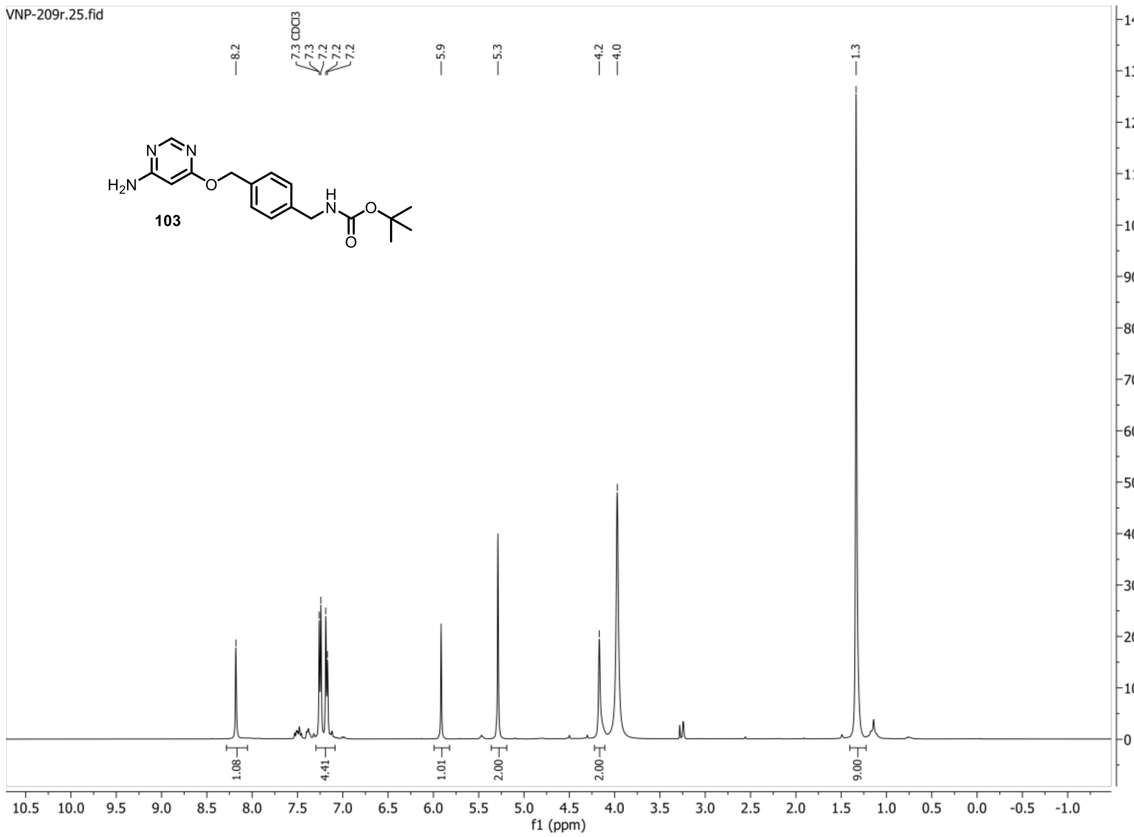


**13C{1H} spectrum of compound 103:**

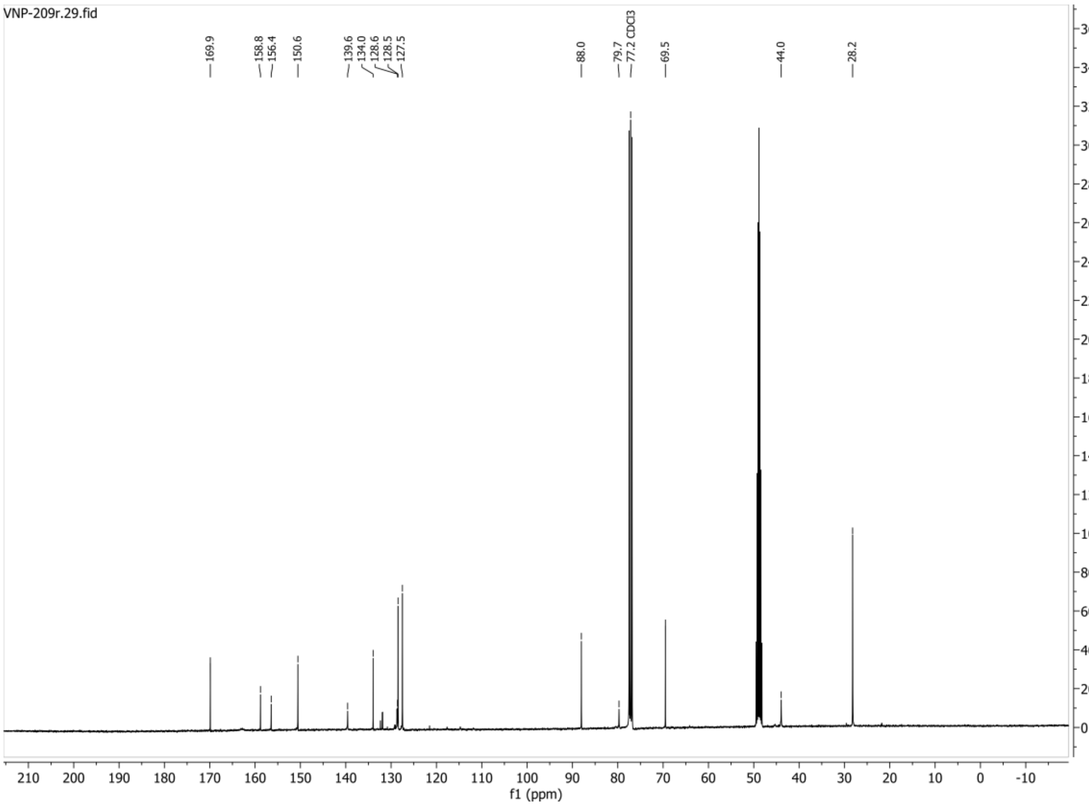

**HSQC-DEPT135 spectrum of compound 103:**

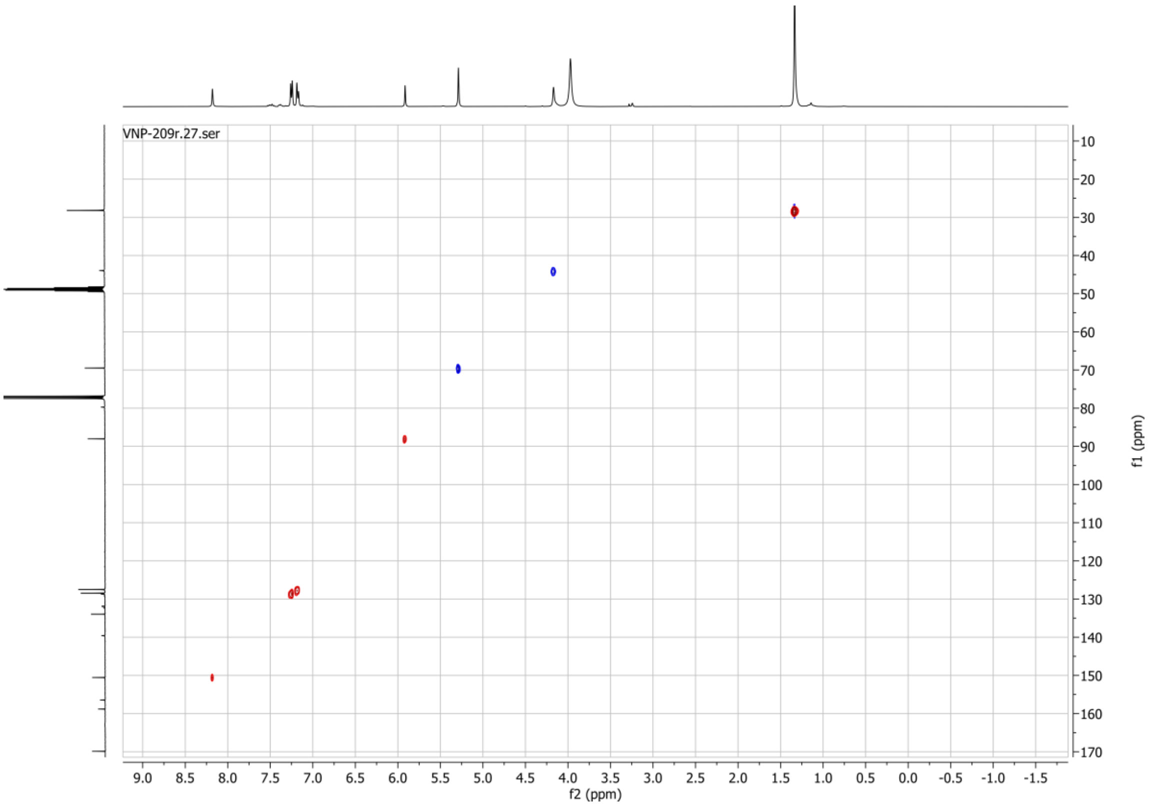

**1H spectrum of compound 106:**

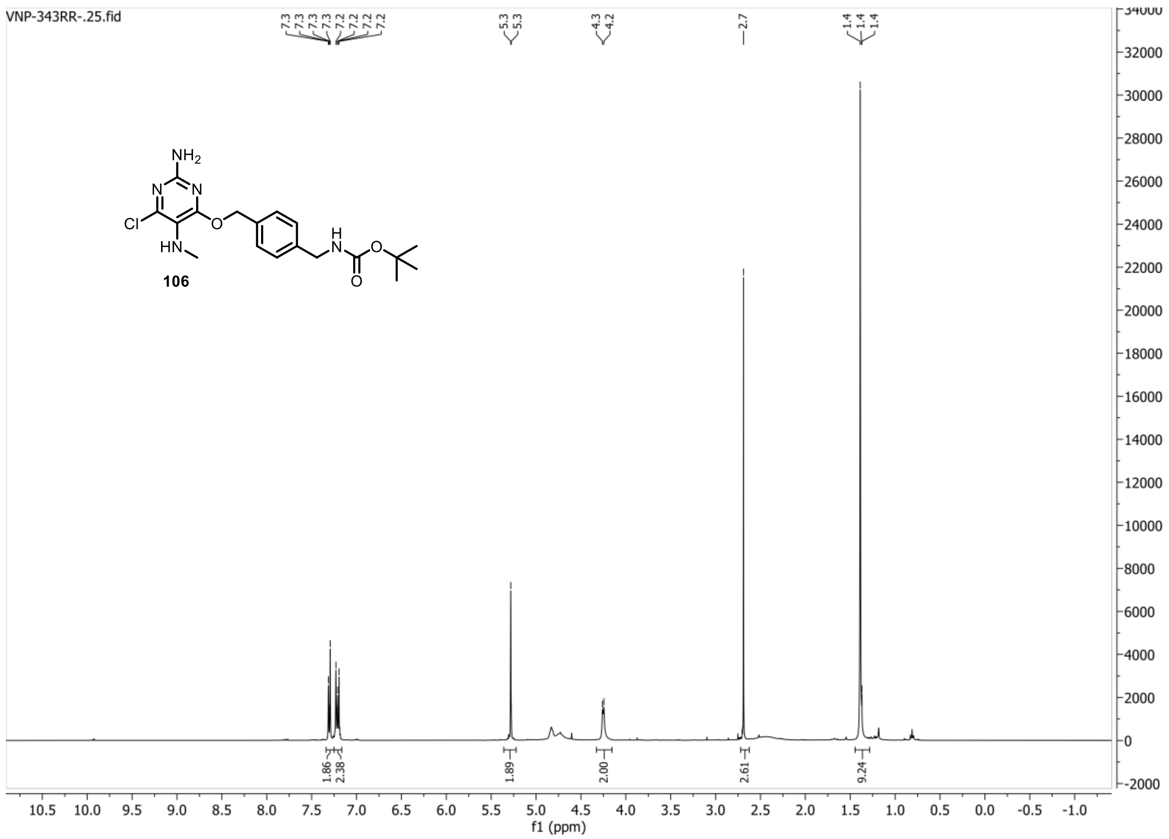


**13C{1H} spectrum of compound 106:**

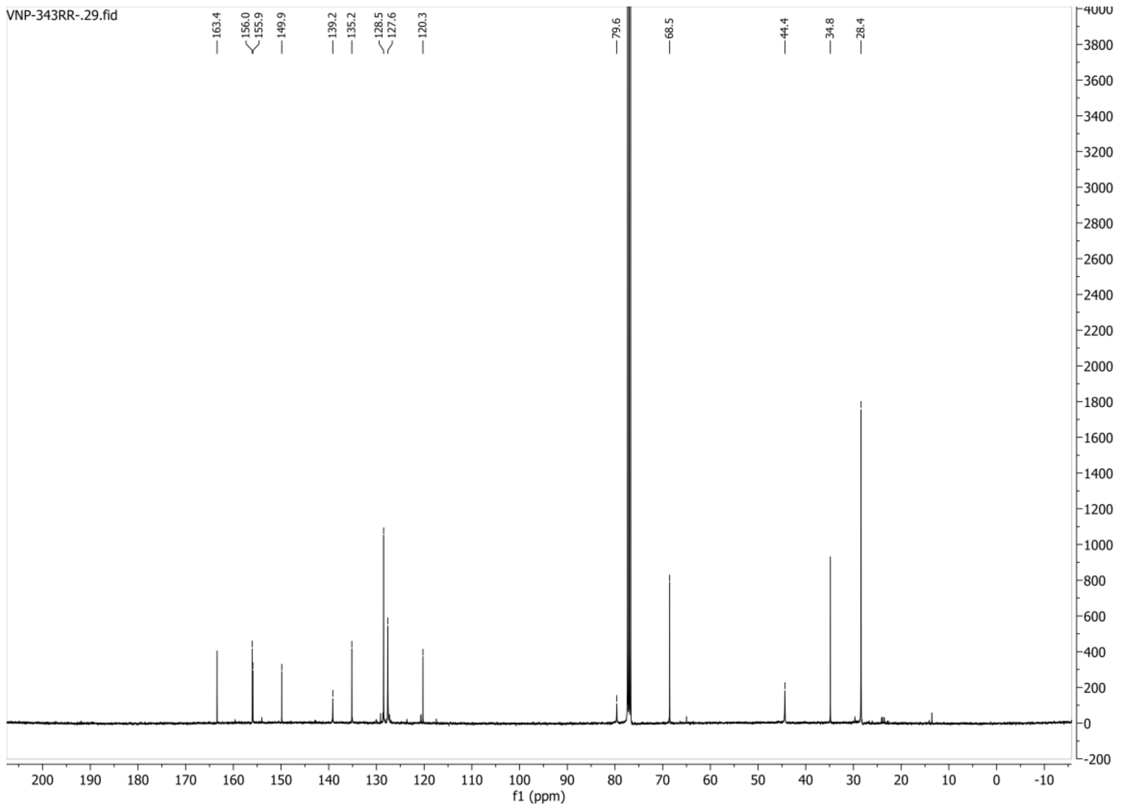

**HSQC-DEPT spectrum of compound 106:**

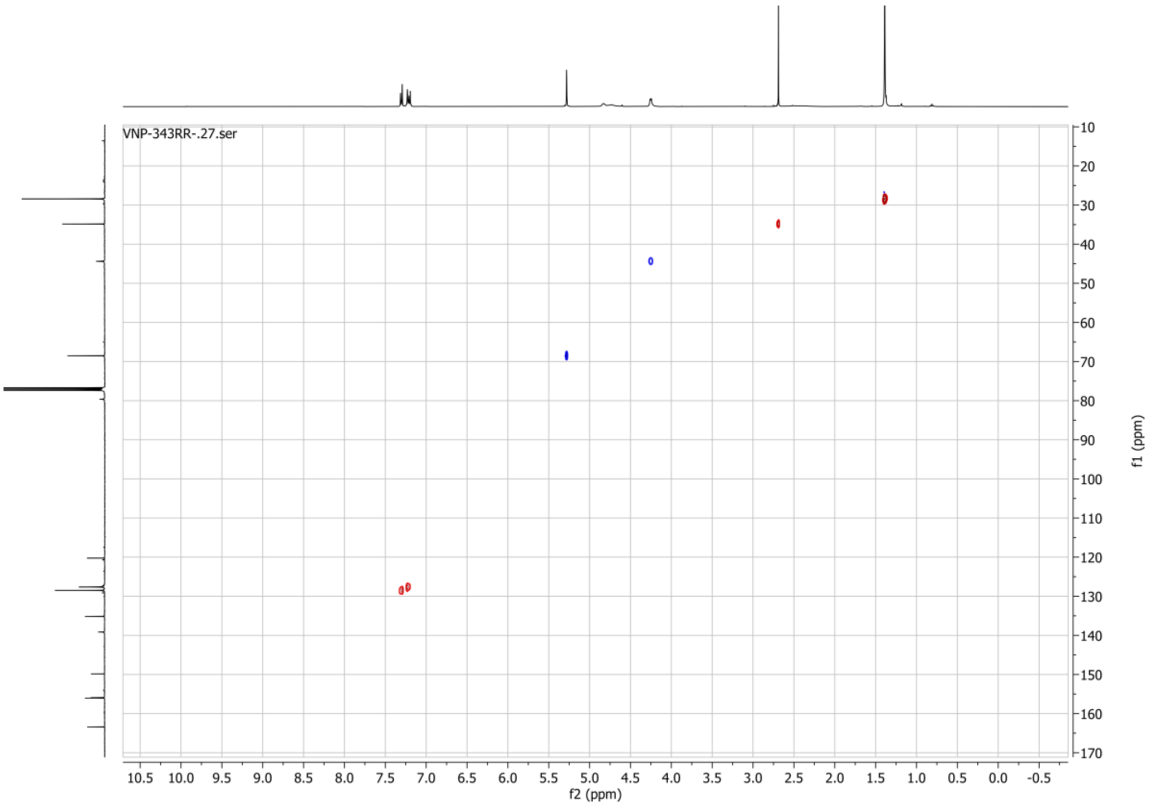

#### 1H spectrum of compound 109:

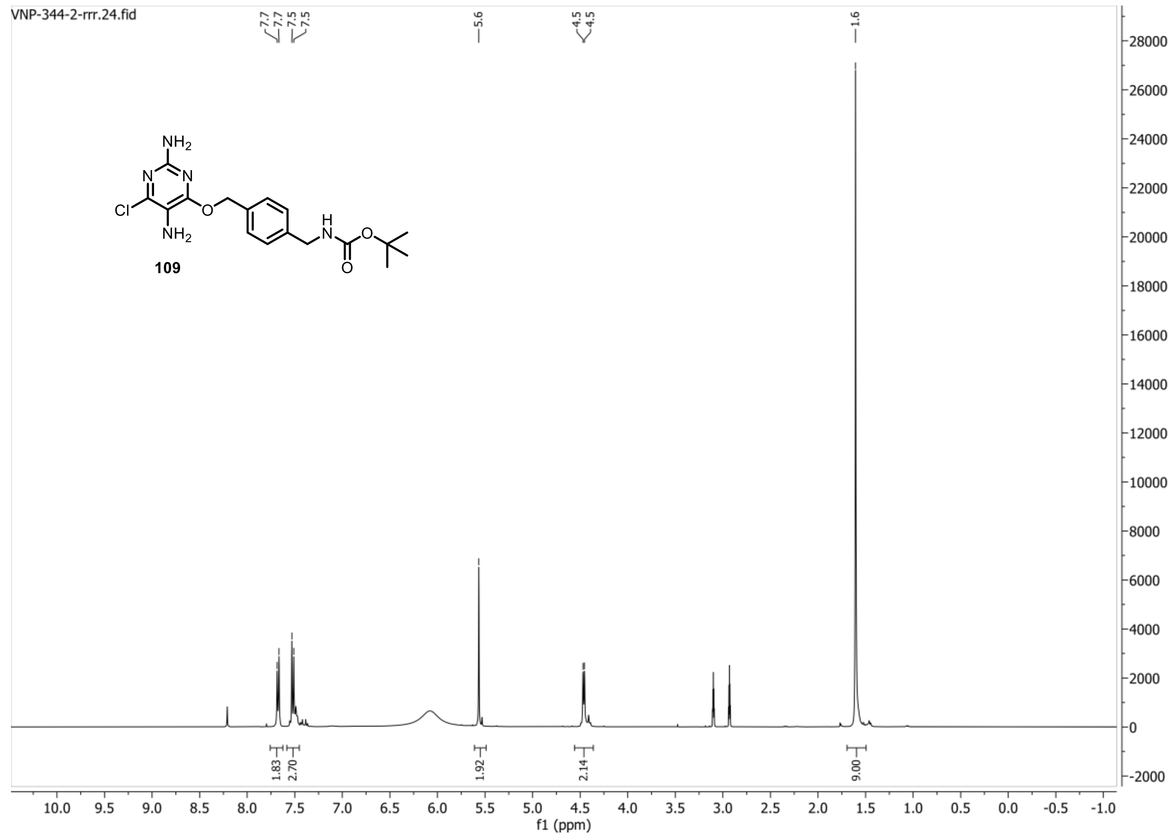


#### 13C{1H} spectrum of compound 109:

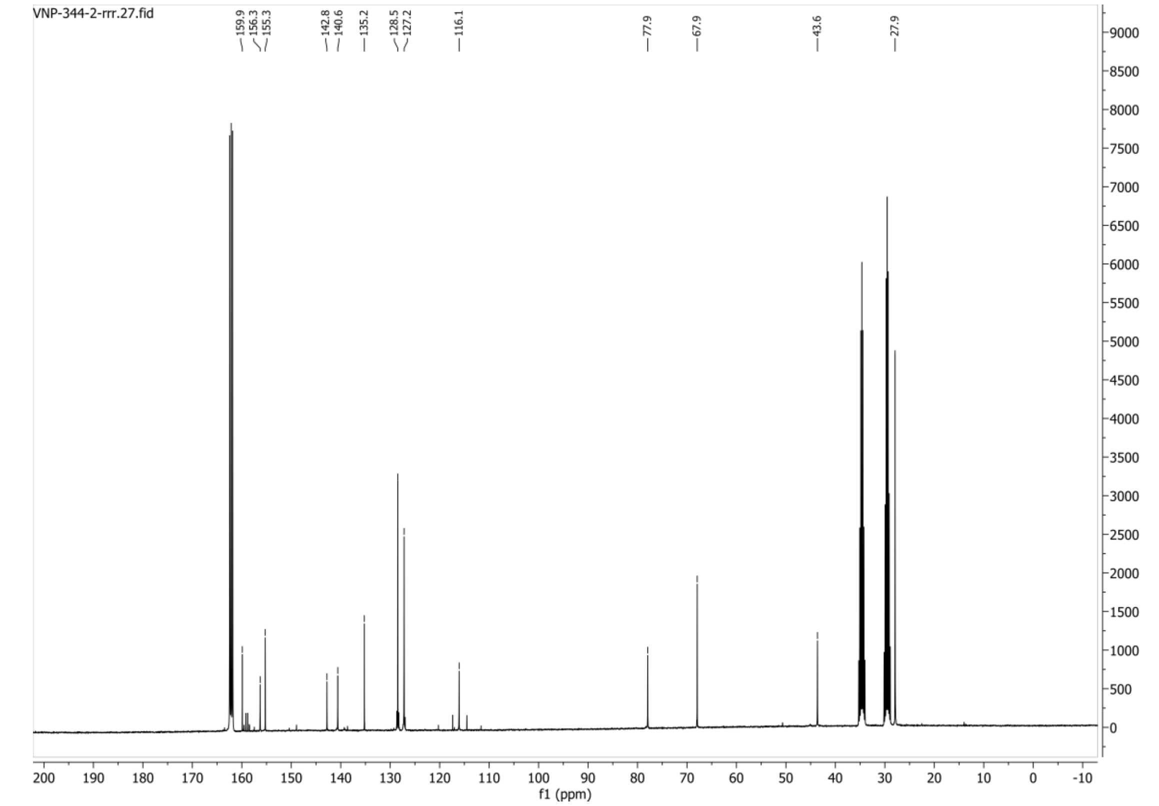

**HSQC-DEPT spectrum of compound 109:**

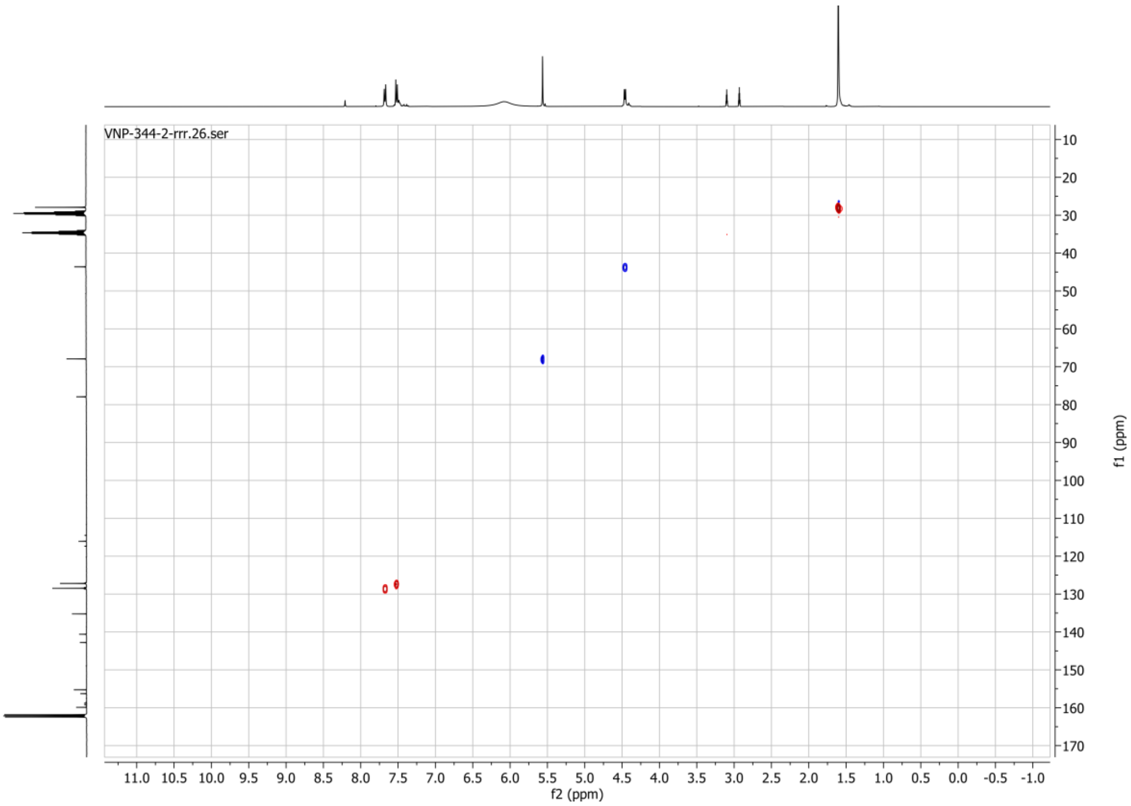

#### 1H spectrum of compound 111:

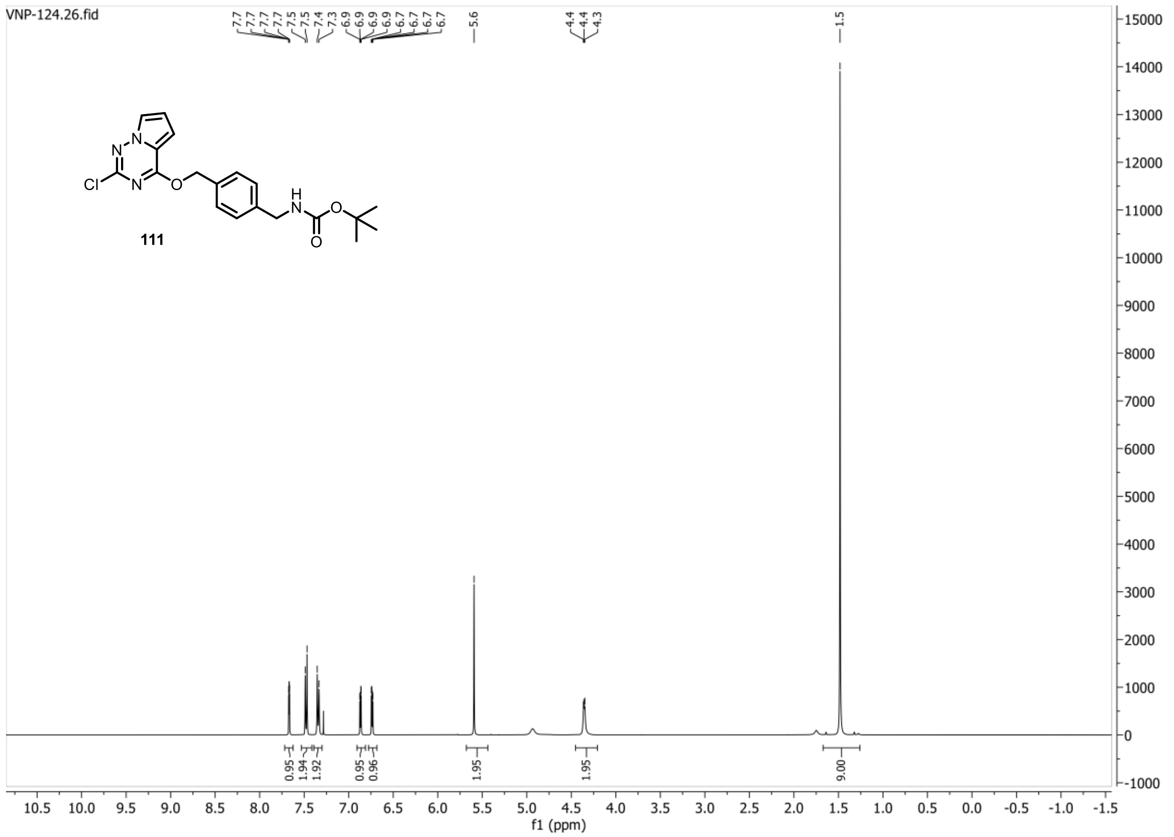


#### 13C{1H} spectrum of compound 111:

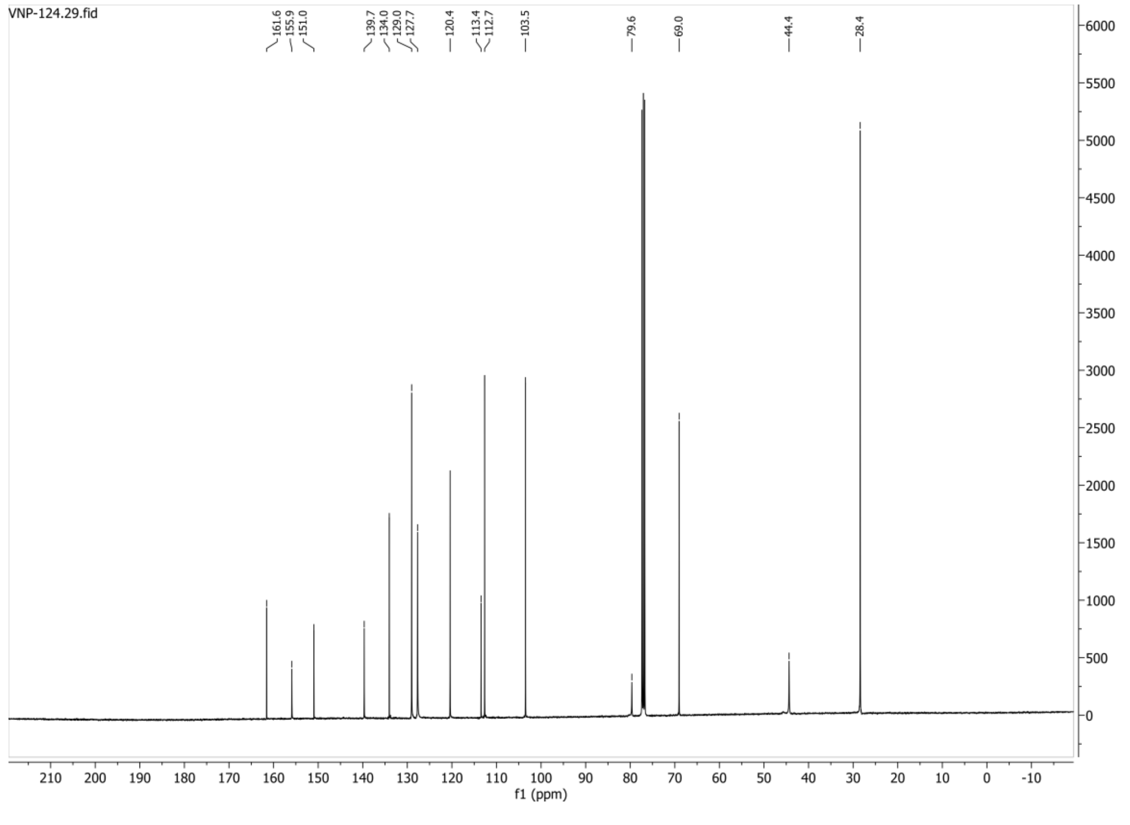

**HSQC-DEPT135 spectrum of compound 111:**

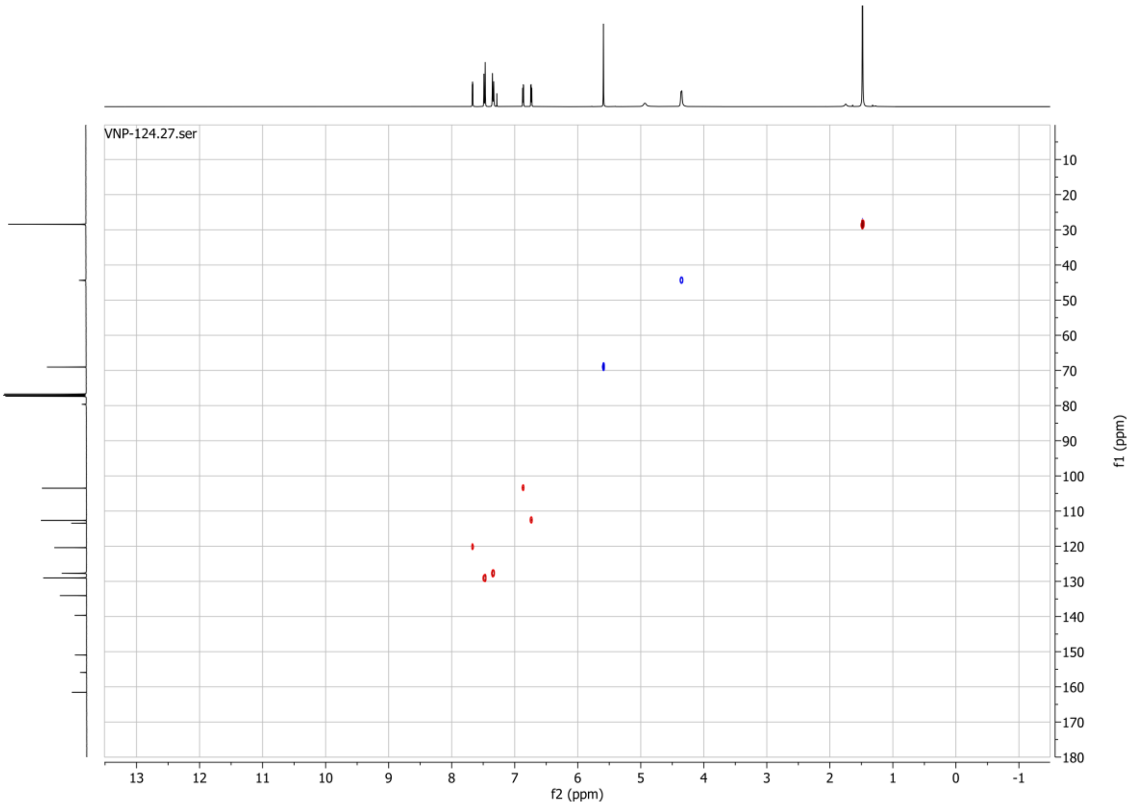

1H spectrum of compound 113:

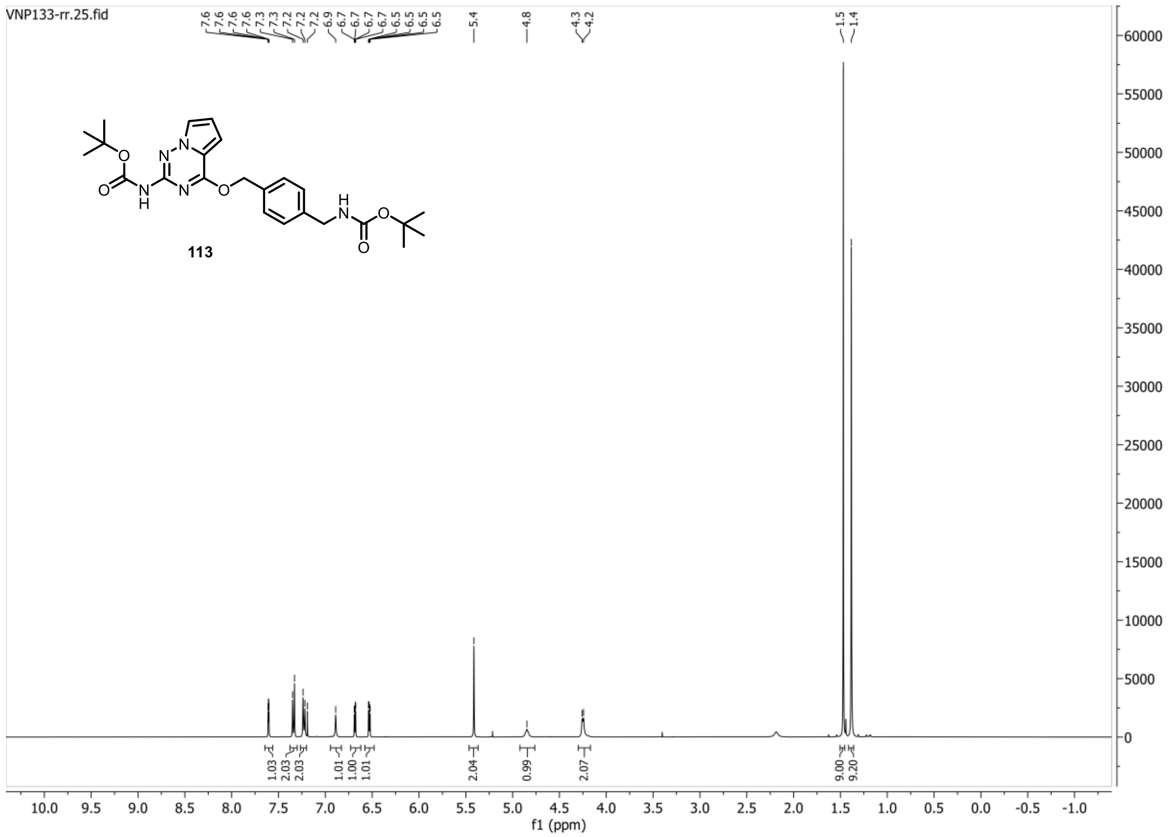


13C{1H} spectrum of compound 113:

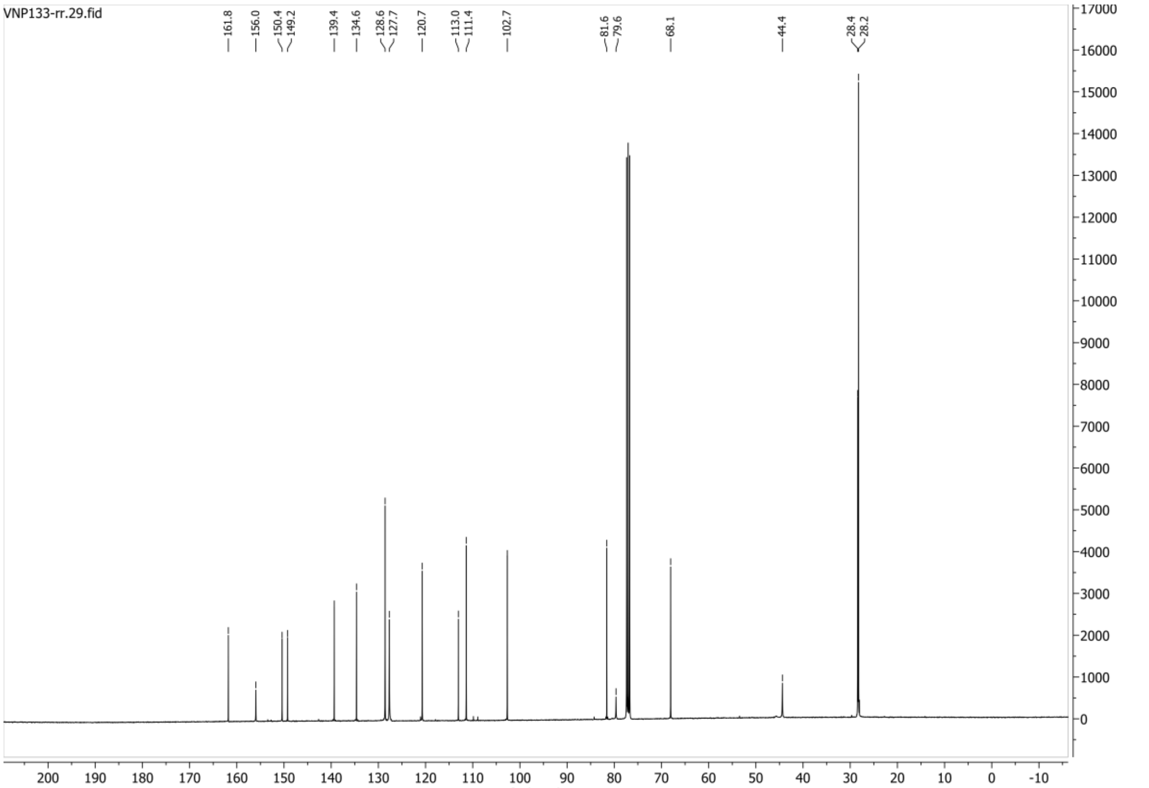

**HSQC-DEPT135 spectrum of compound 113:**

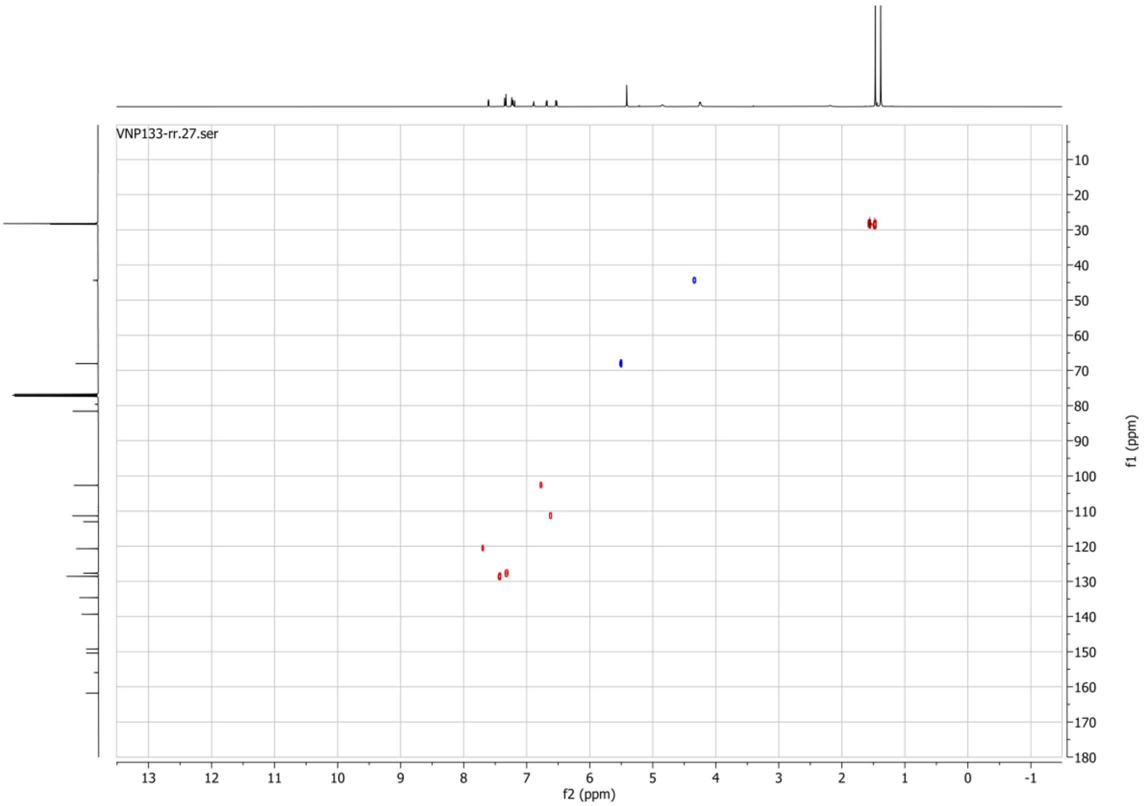

**1H spectrum of compound 114:**

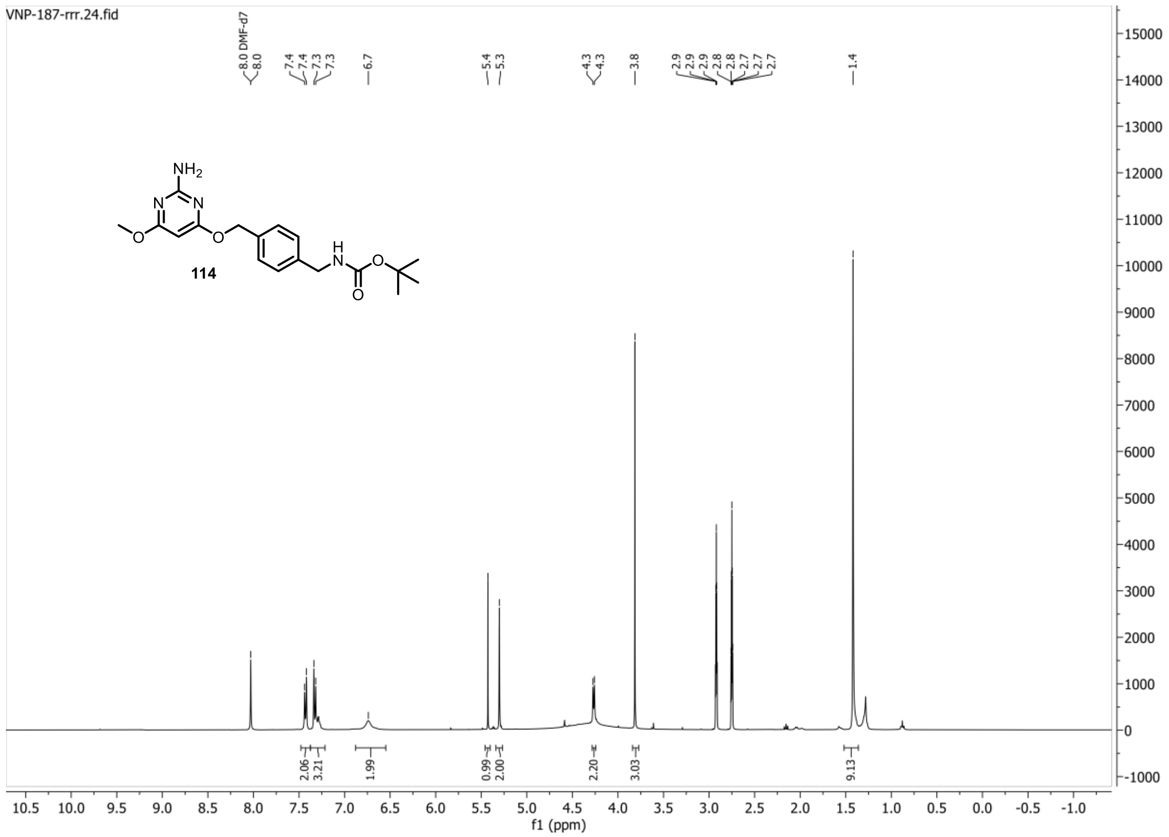


**13C{1H} spectrum of compound 114:**

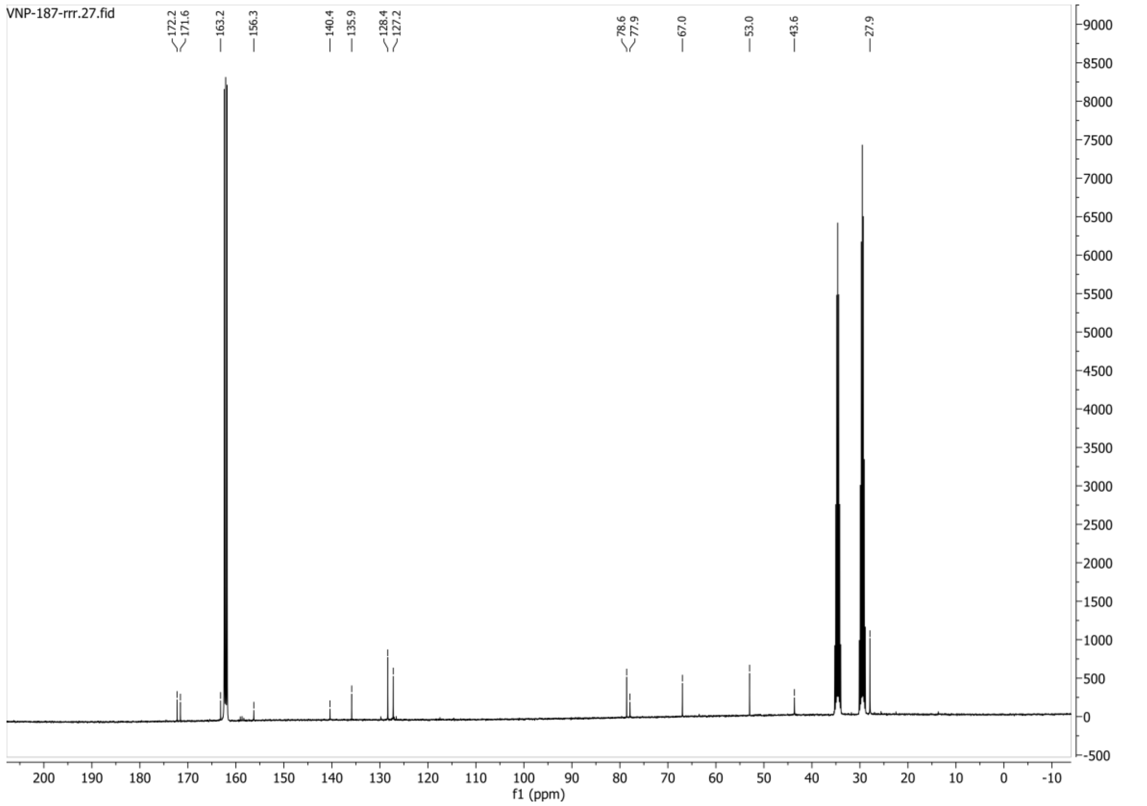

**1H spectrum of compound 115:**

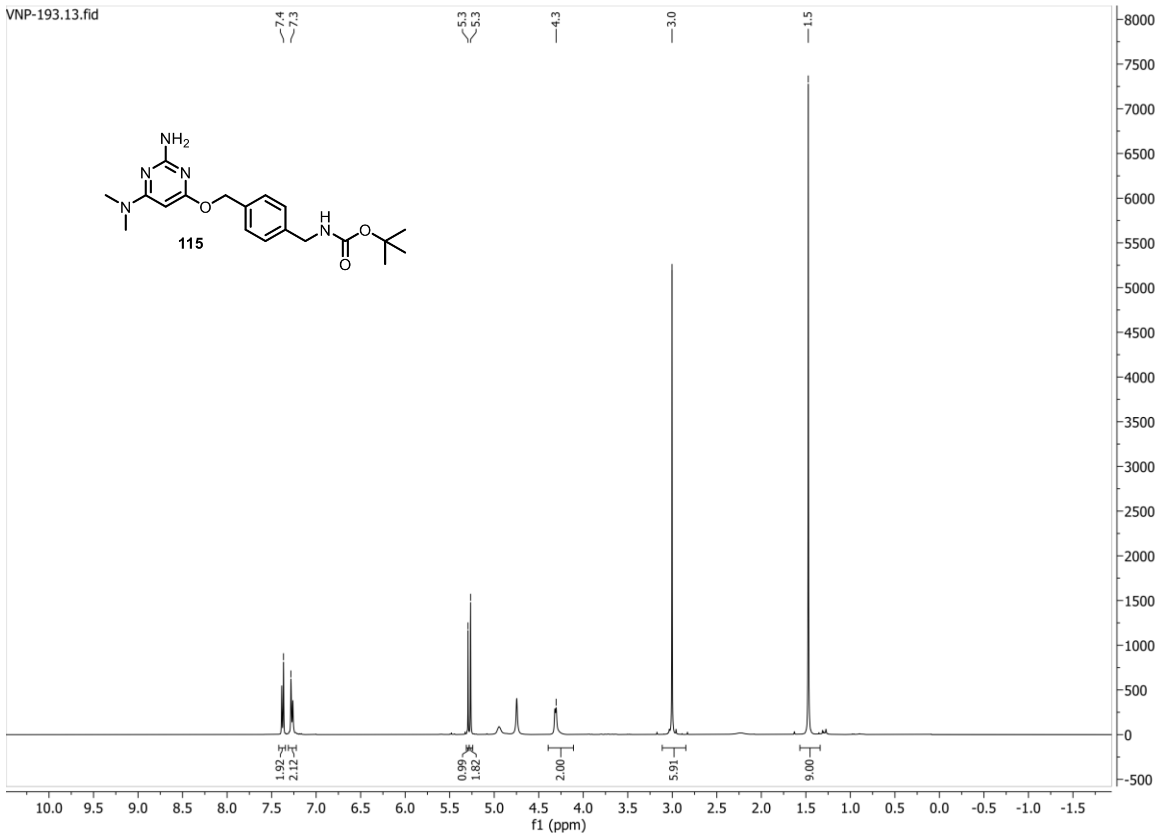


**13C{1H} spectrum of compound 115:**

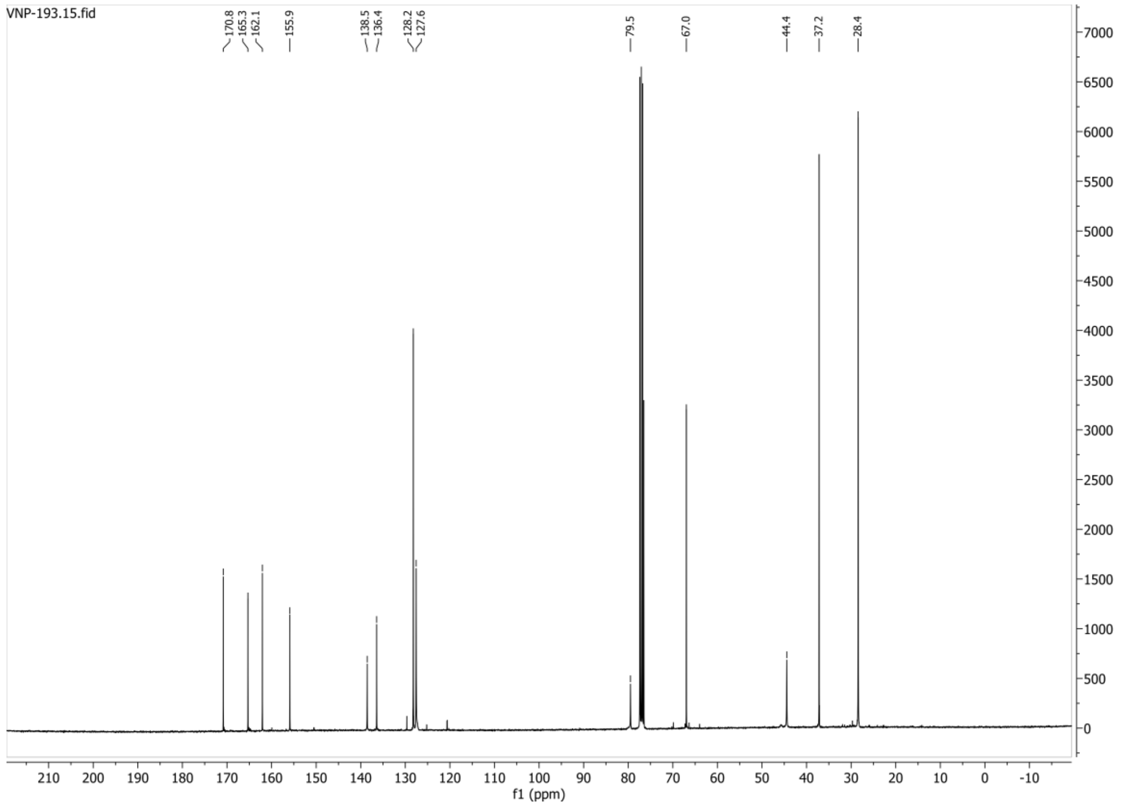

**HSQC-DEPT135 spectrum of compound 115:**

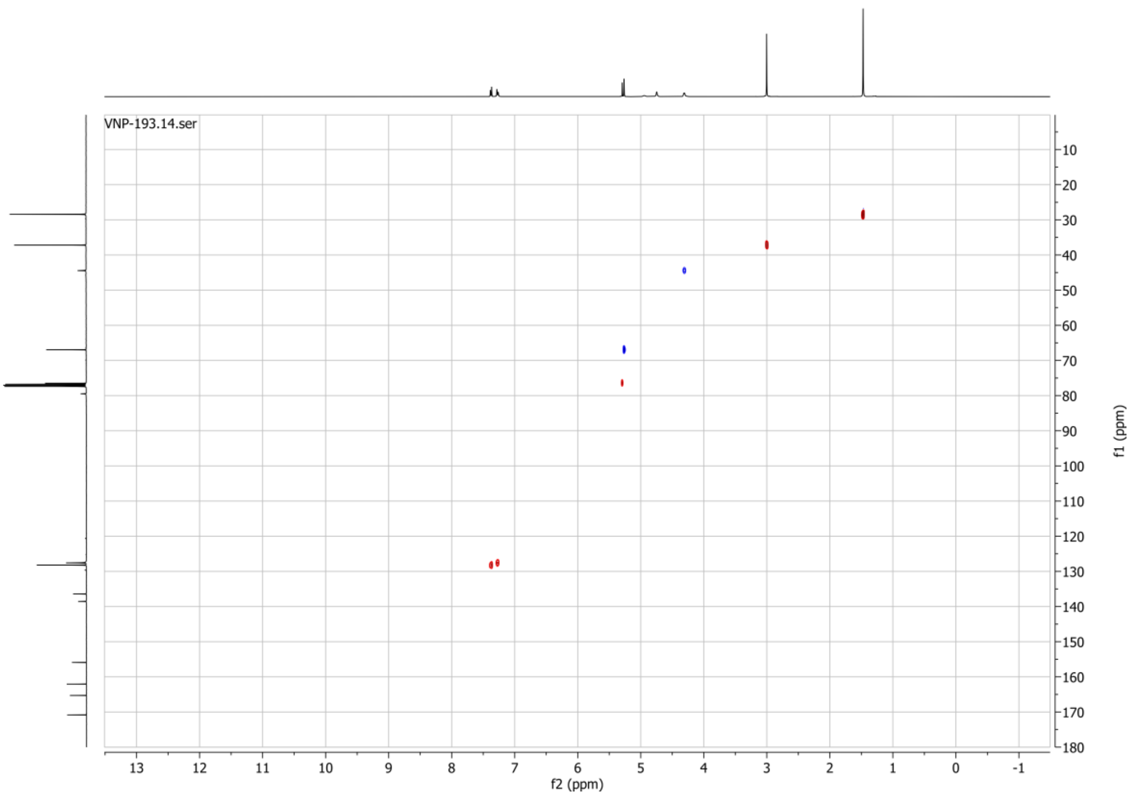

**1H spectrum of compound 116:**

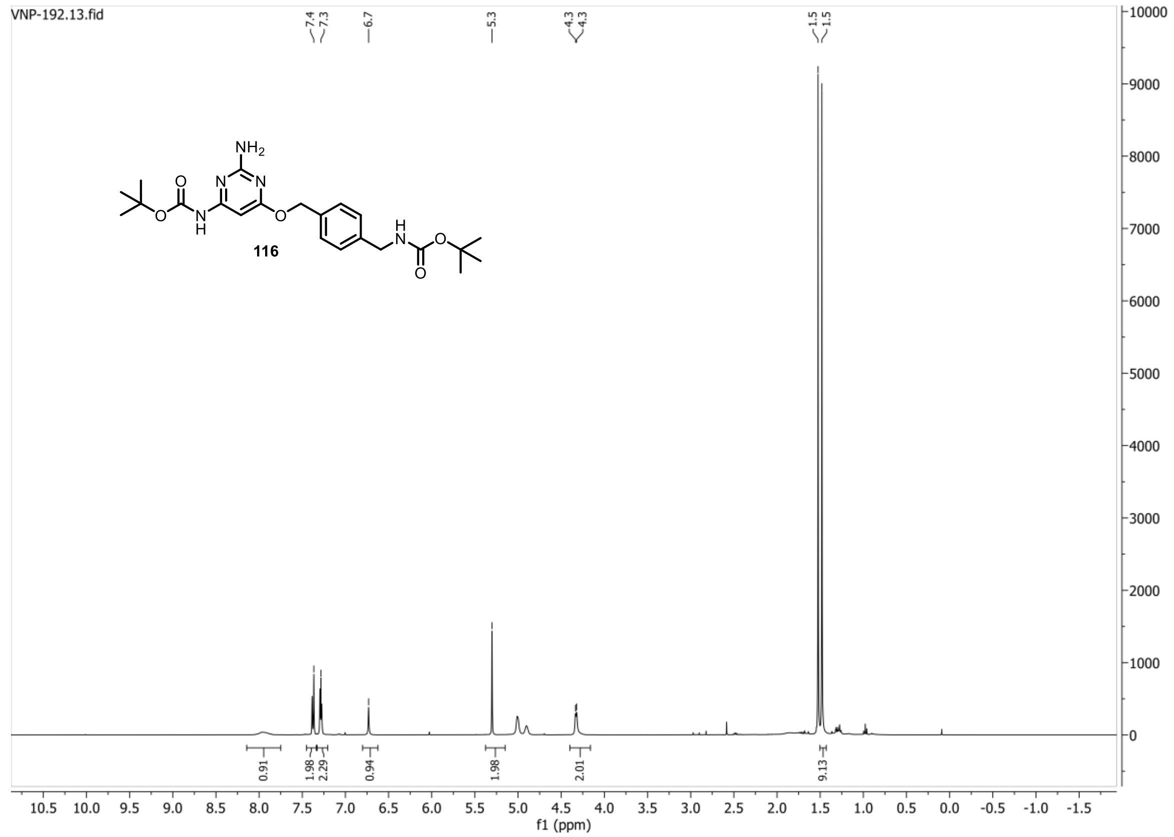


**13C{1H} spectrum of compound 116:**

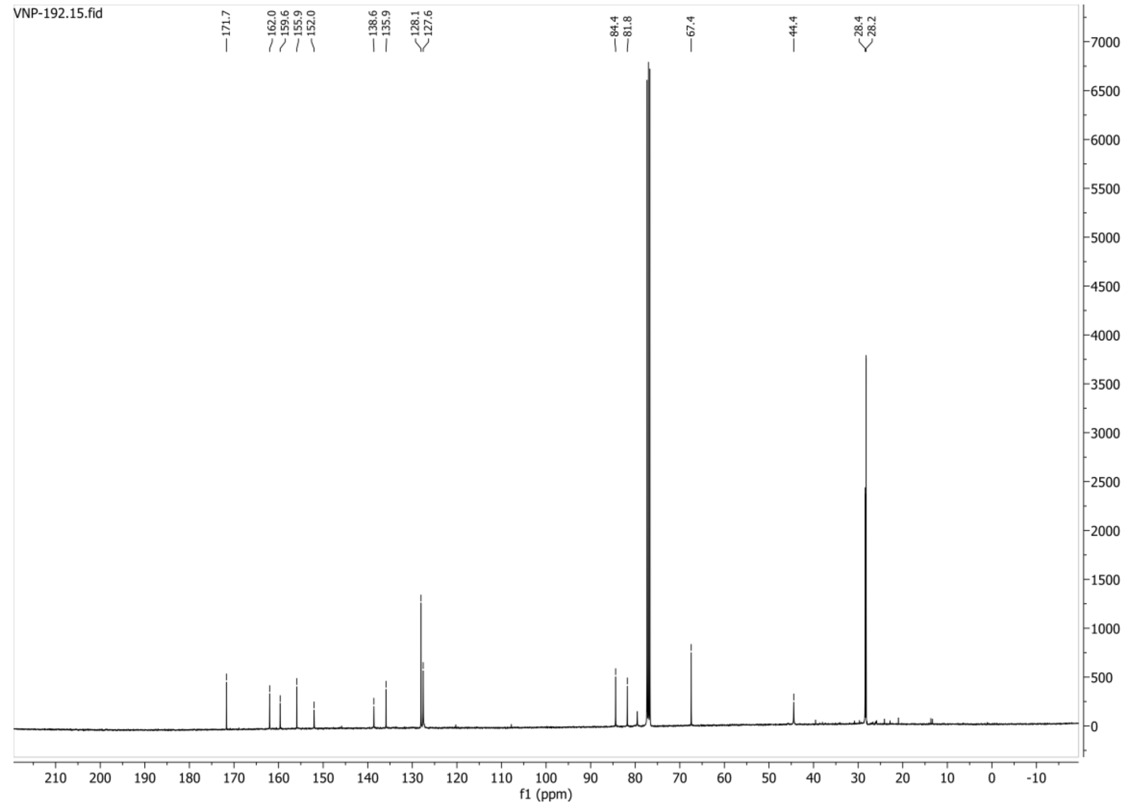

**HSQC-DEPT135 spectrum of compound 116:**

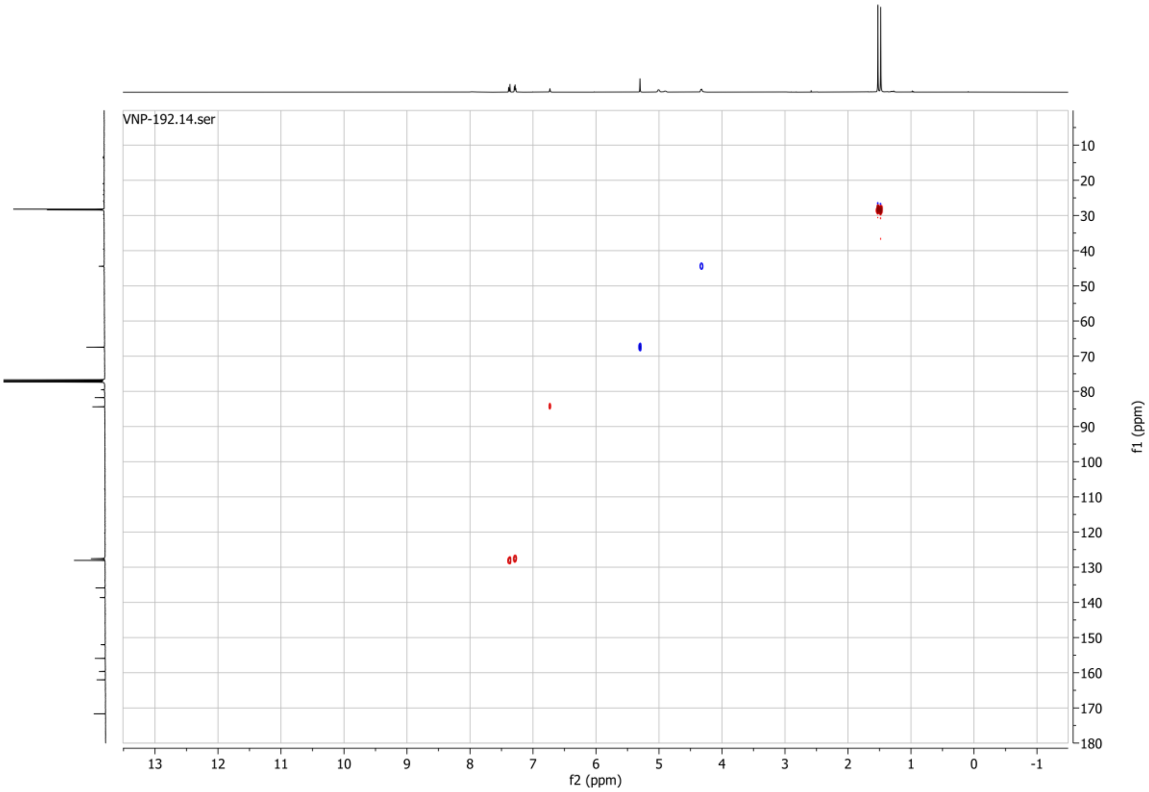

**1H spectrum of compound 119:**

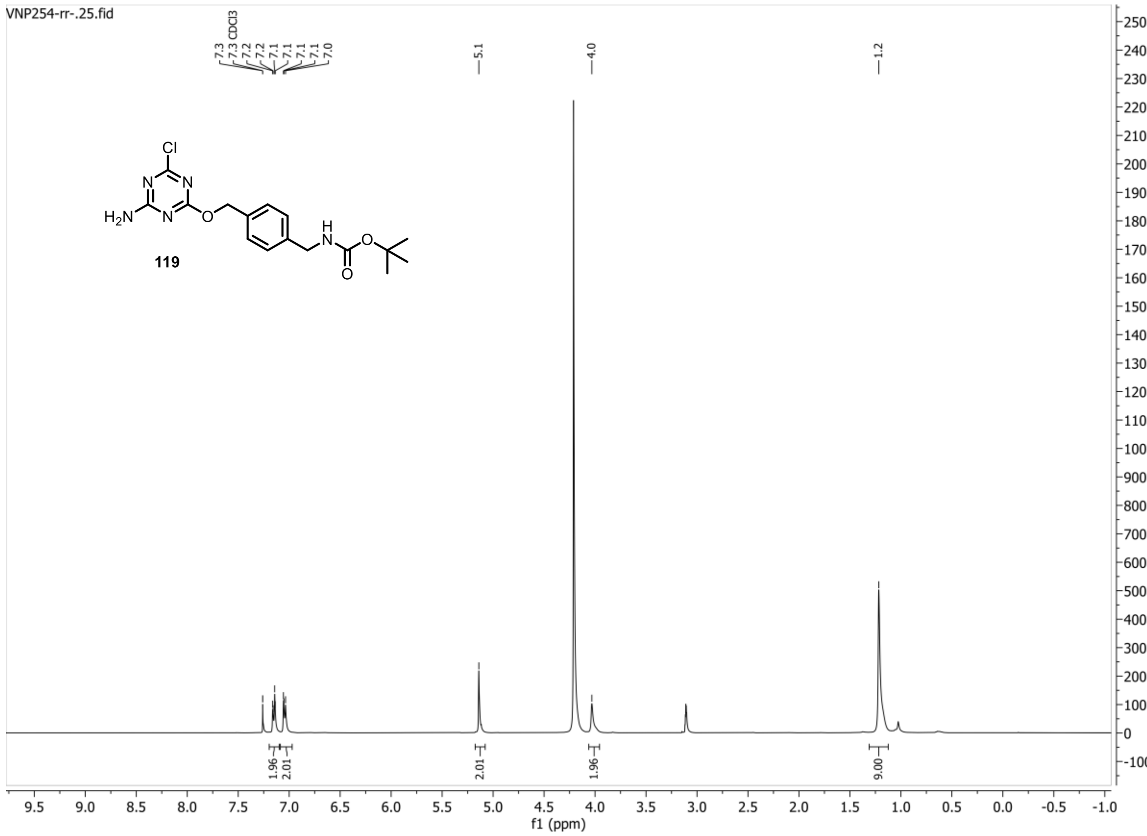


**13C{1H} spectrum of compound 119:**

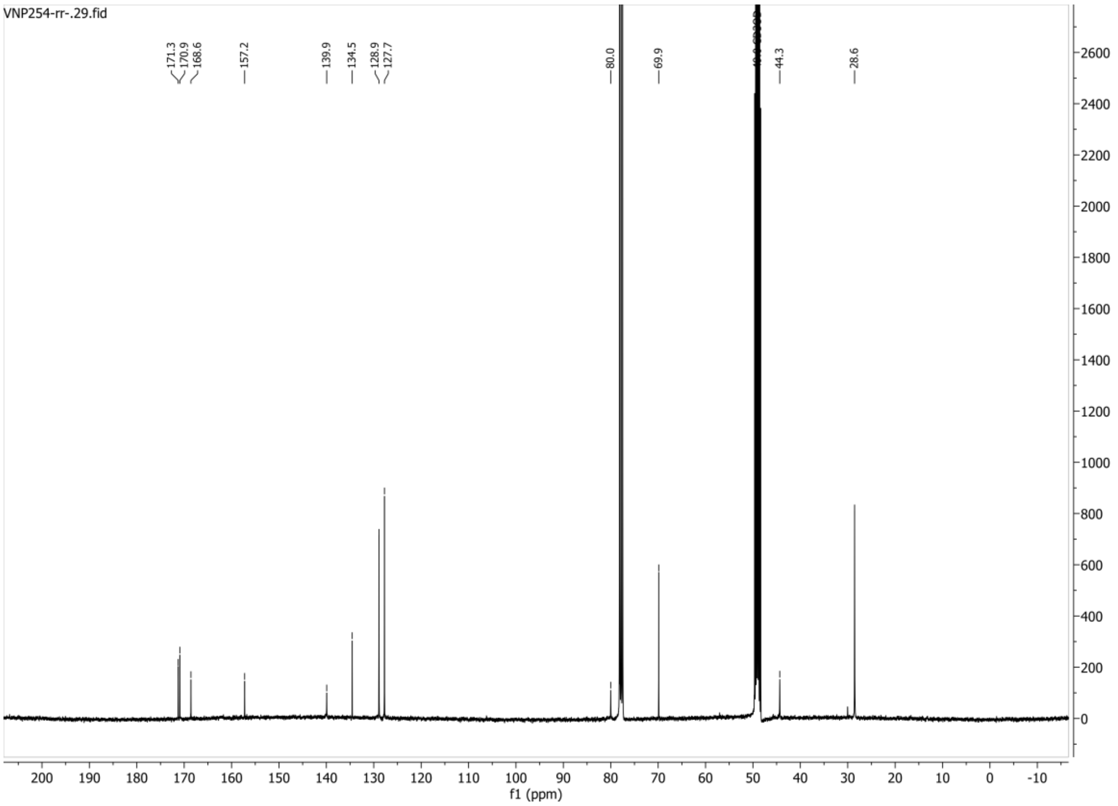

**HSQC-DEPT135 spectrum of compound 119:**

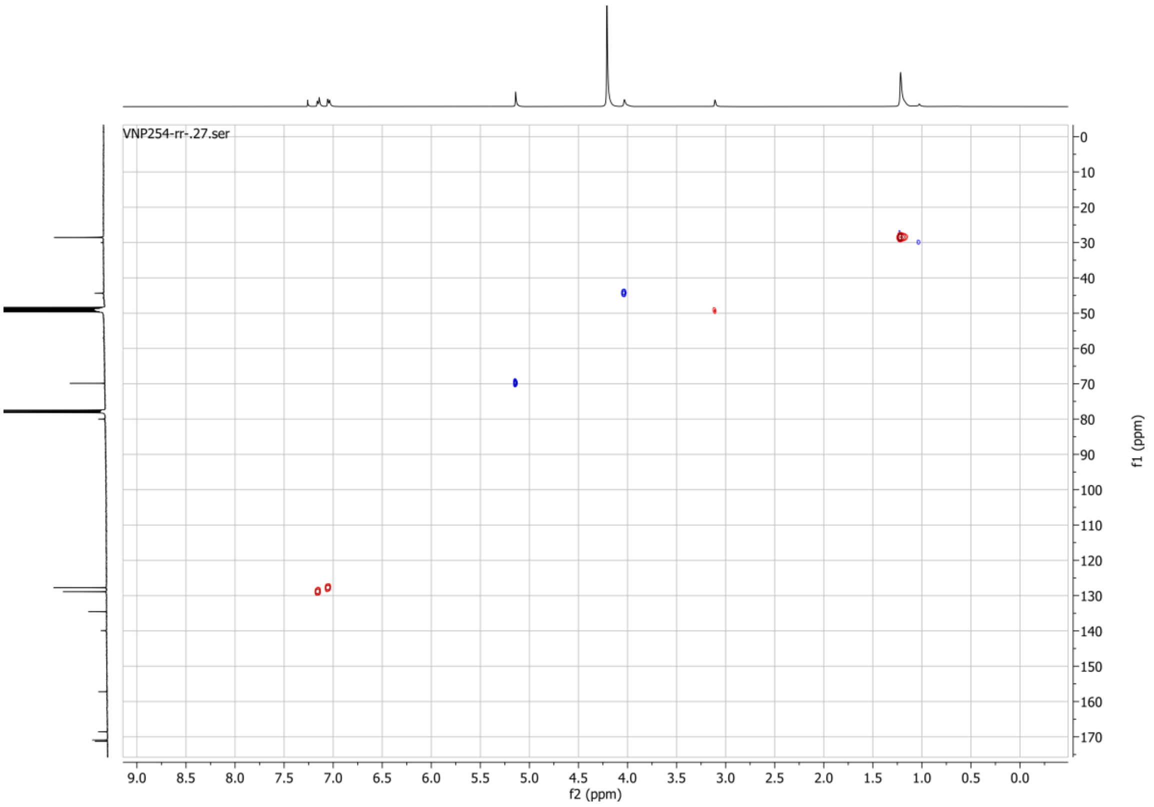

**1H spectrum of compound 122:**

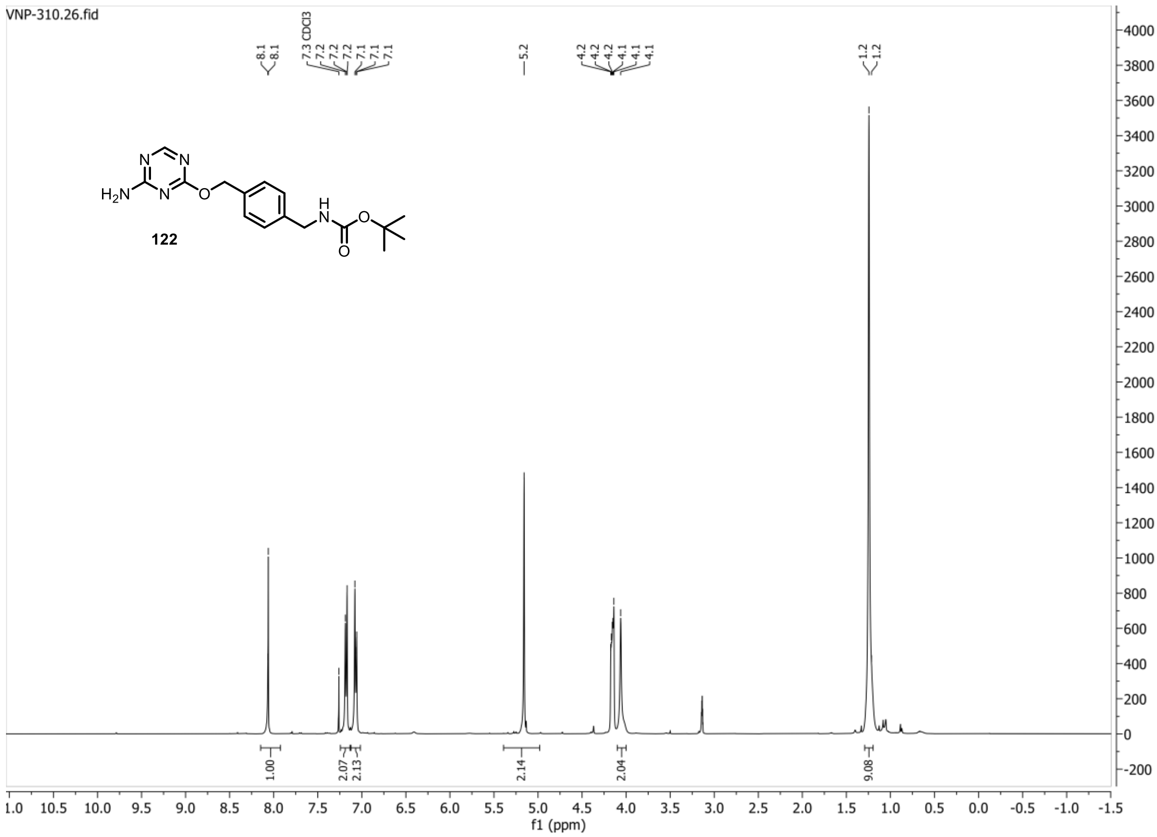


**13C{1H} spectrum of compound 122:**

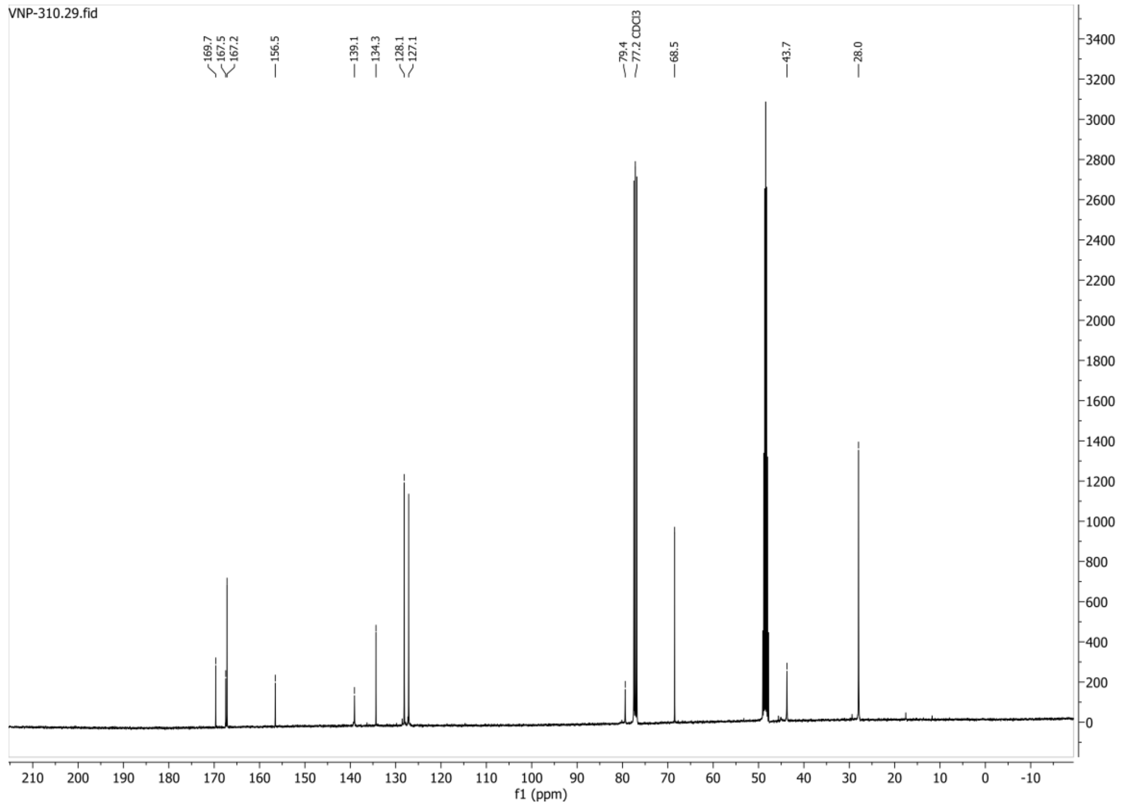

**HSQC-DEPT135 spectrum of compound 122:**

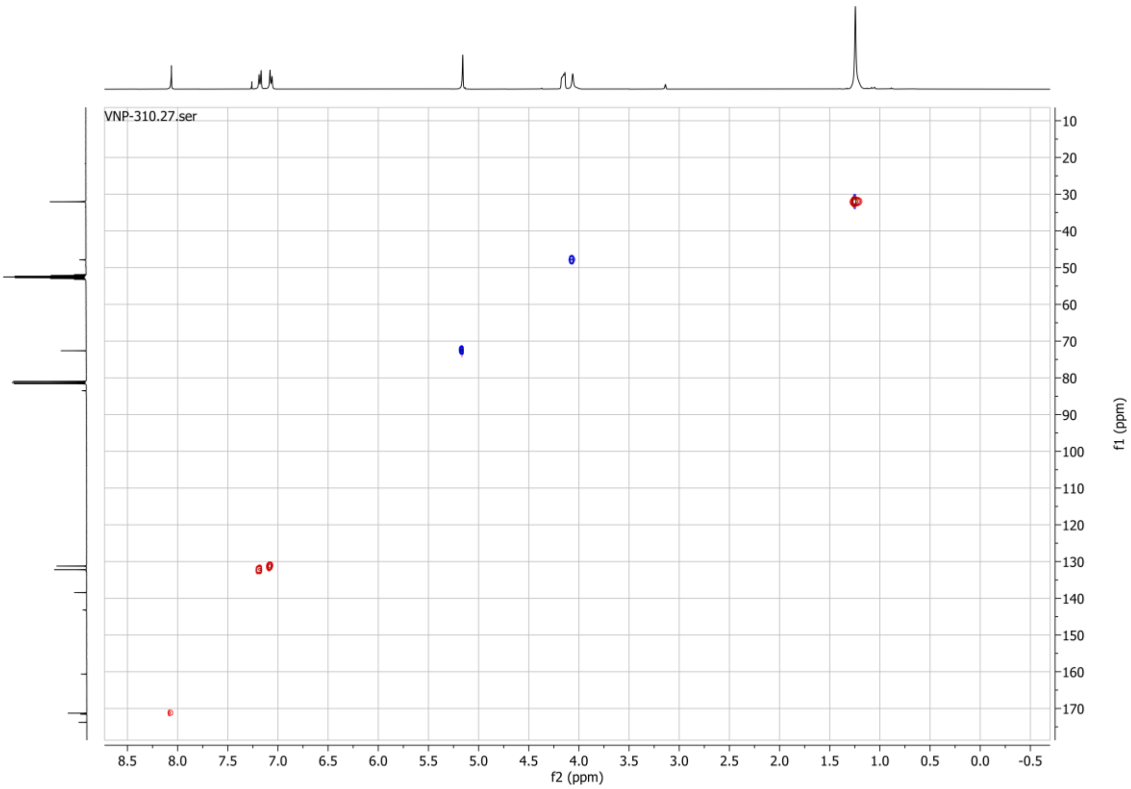

**1H spectrum of compound 123:**

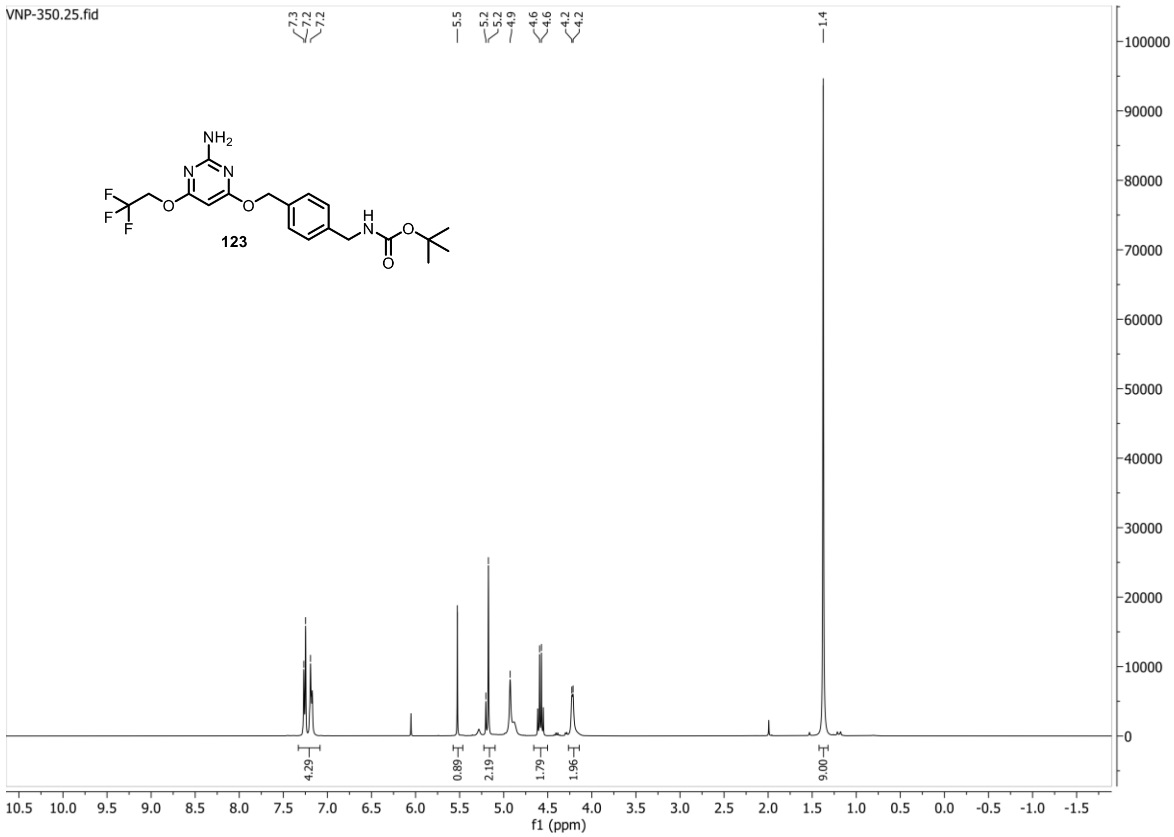


**13C{1H} spectrum of compound 123:**

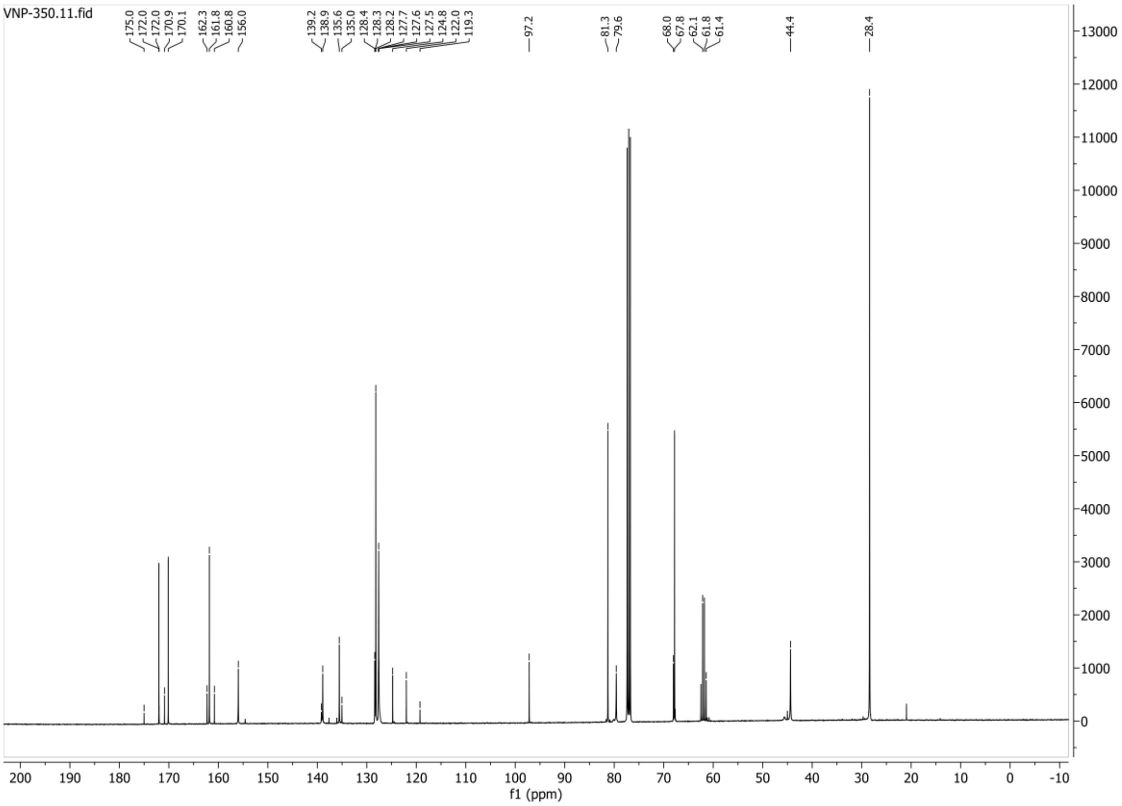

**HSQC-DEPT135 spectrum of compound 123:**

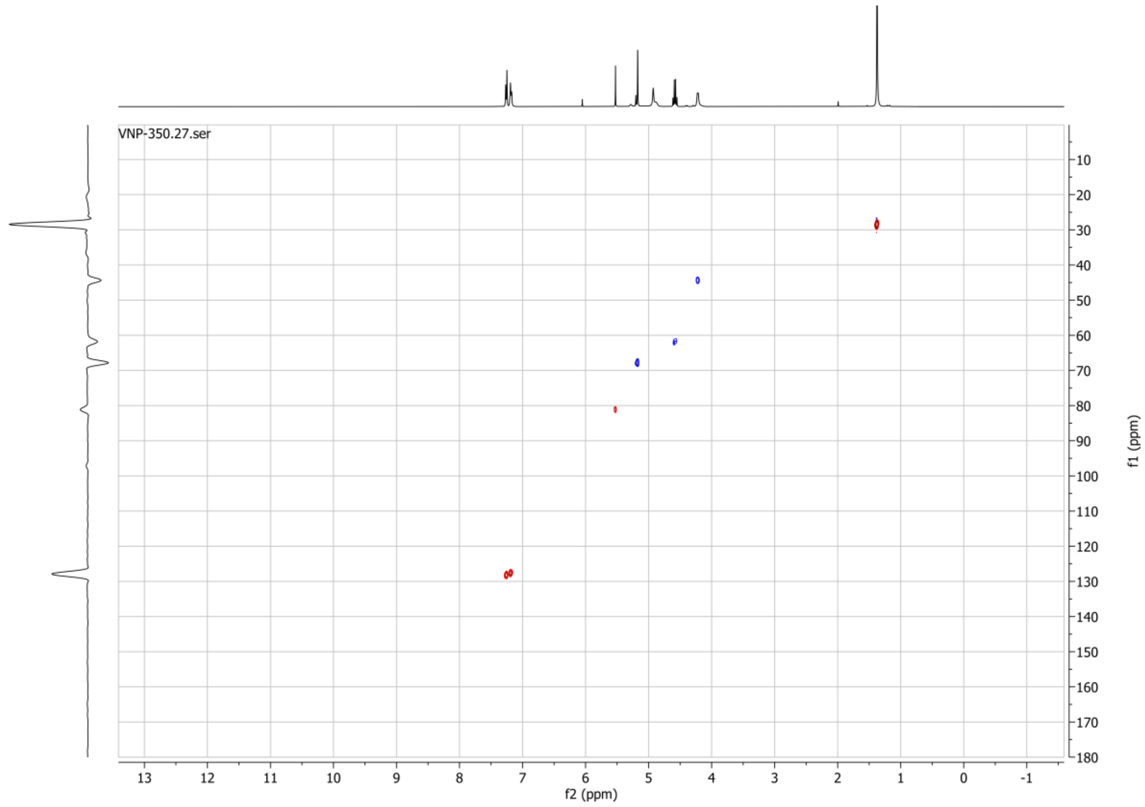


**19F spectrum of compound 123:**

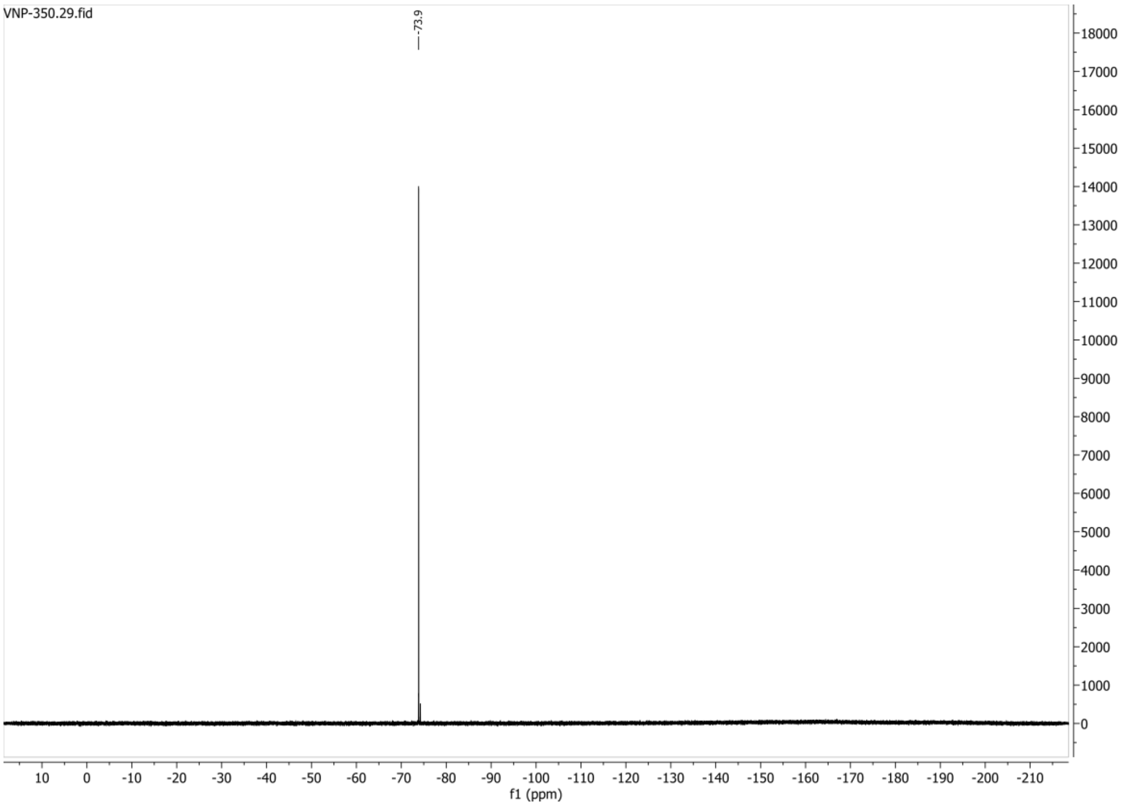

**1H spectrum of compound 124:**

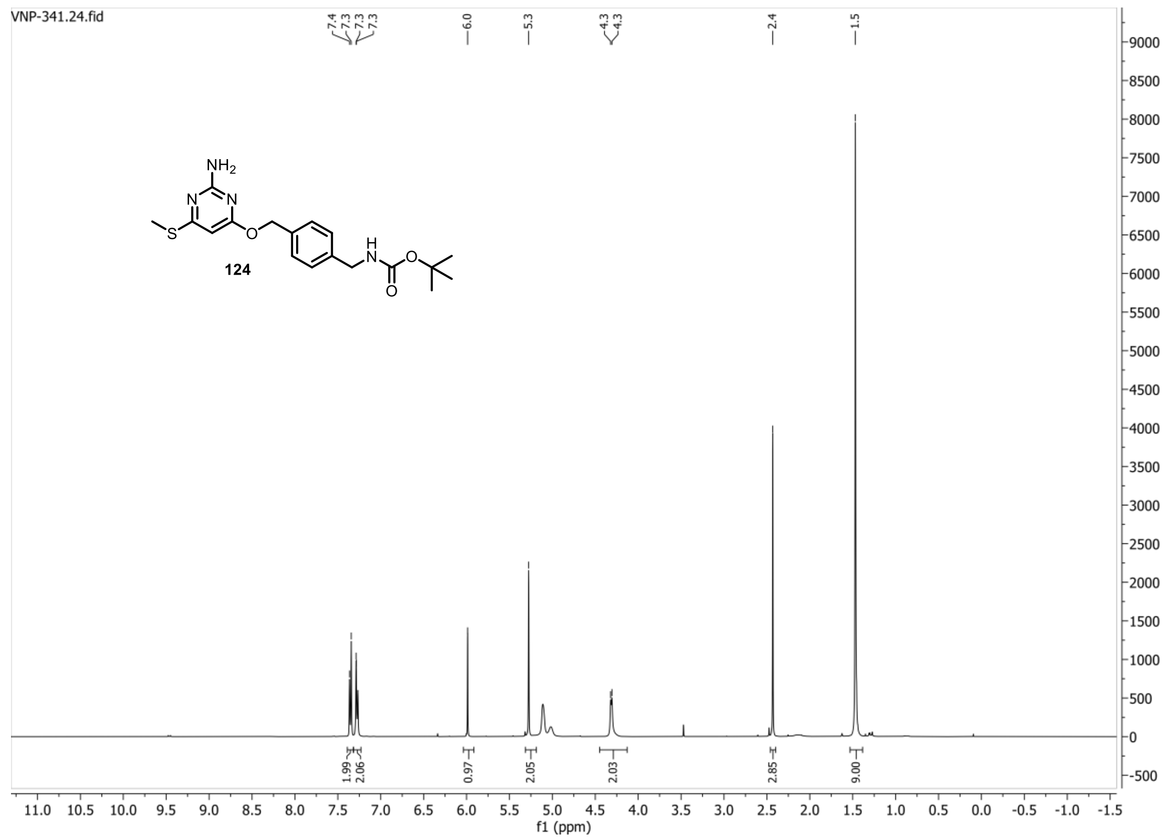


**13C{1H} spectrum of compound 124:**

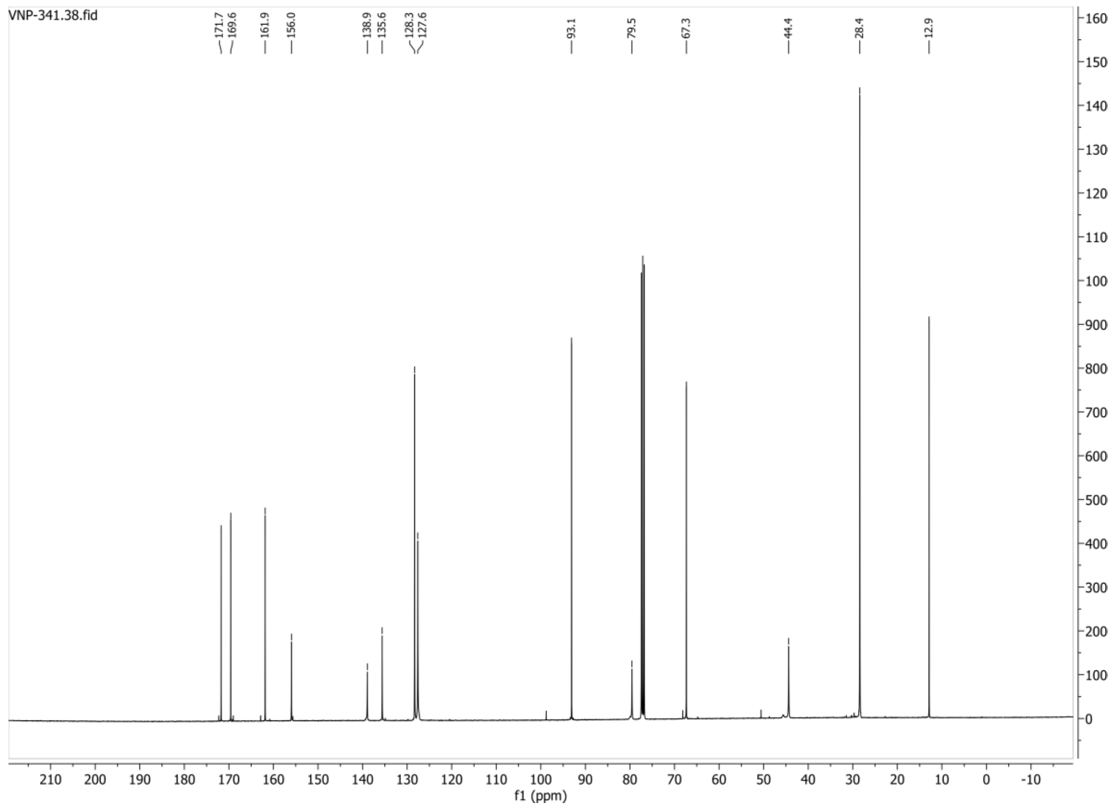

**HSQC-DEPT spectrum of compound 124:**

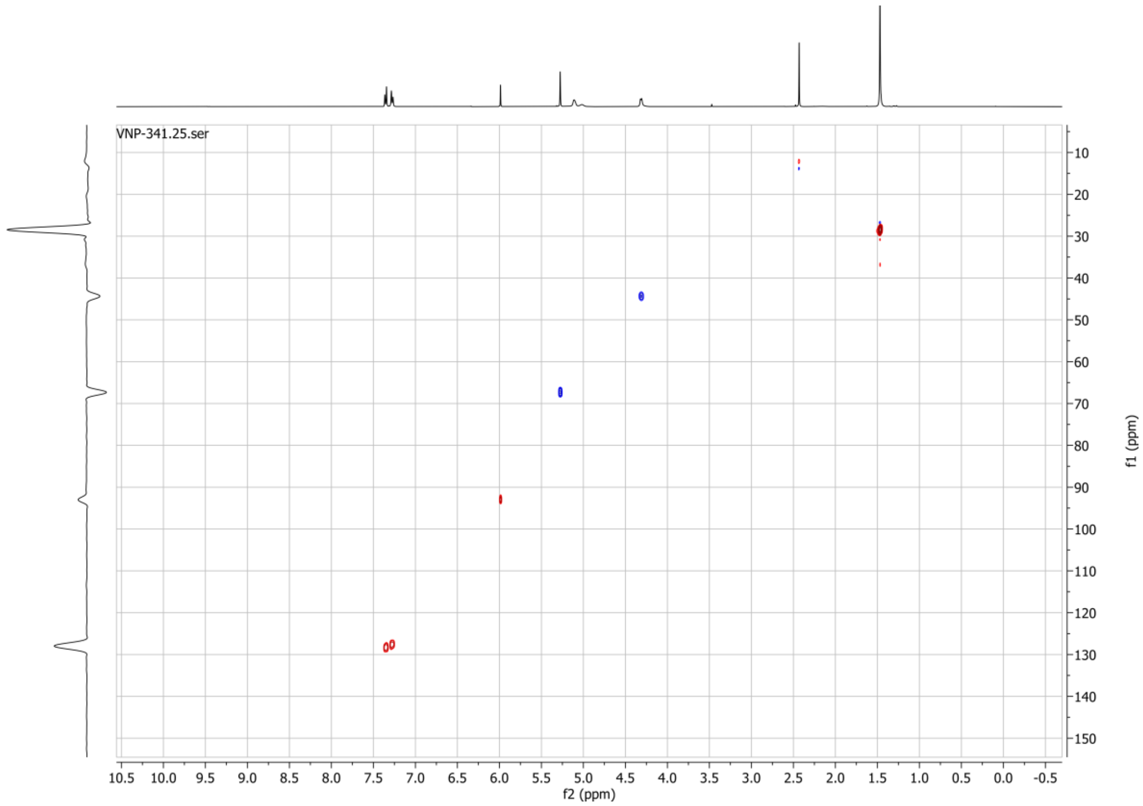

#### 1H spectrum of compound 125:

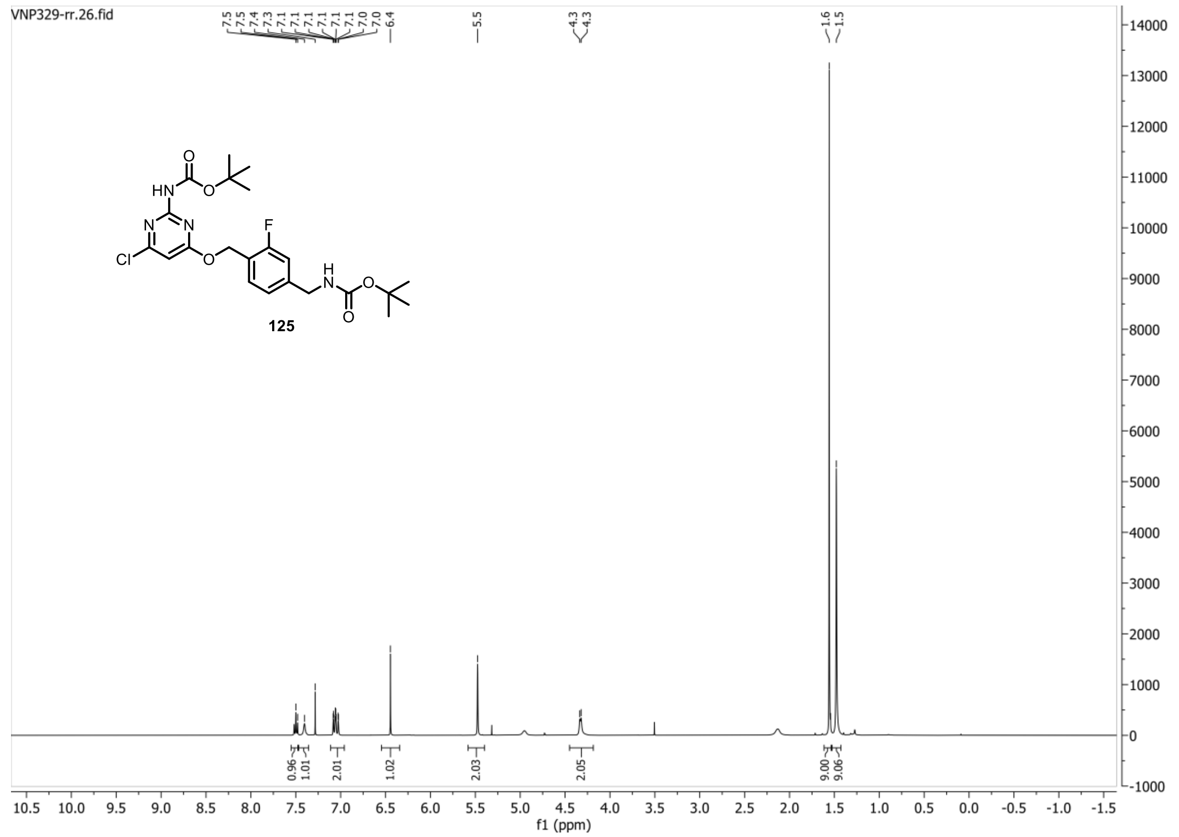


#### 13C{1H} spectrum of compound 125:

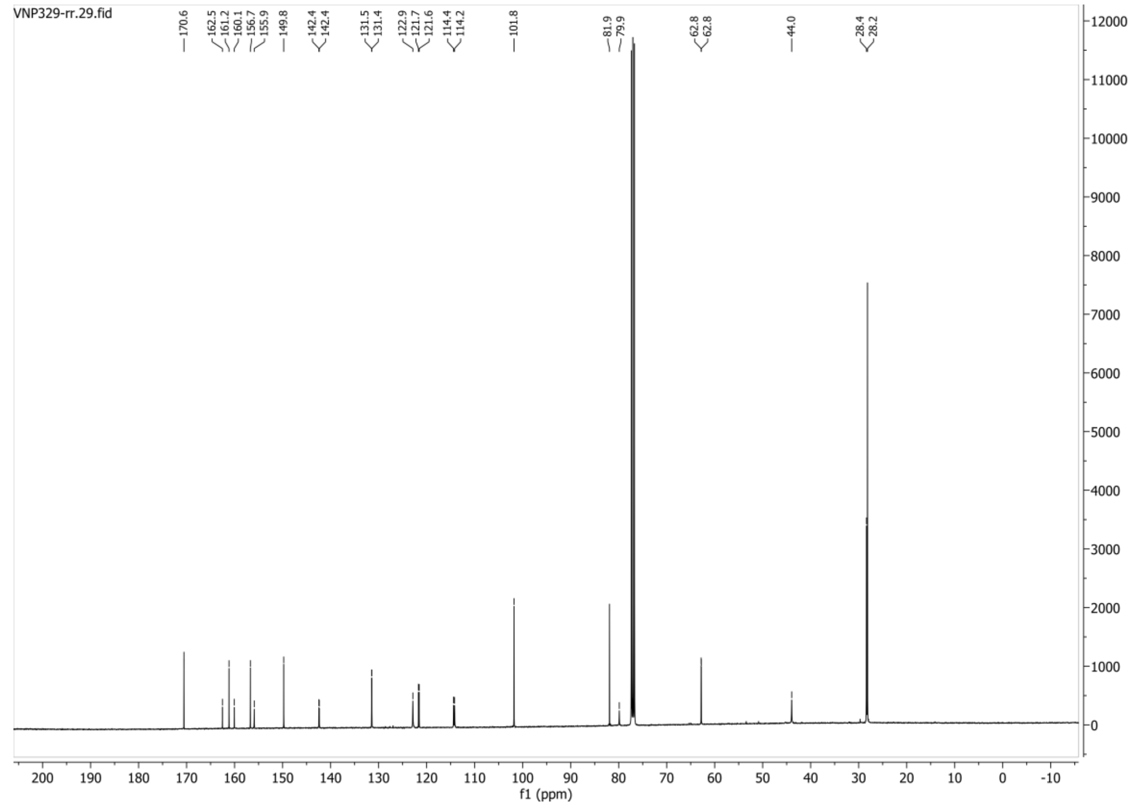

**HSQC-DEPT135 spectrum of compound 125:**

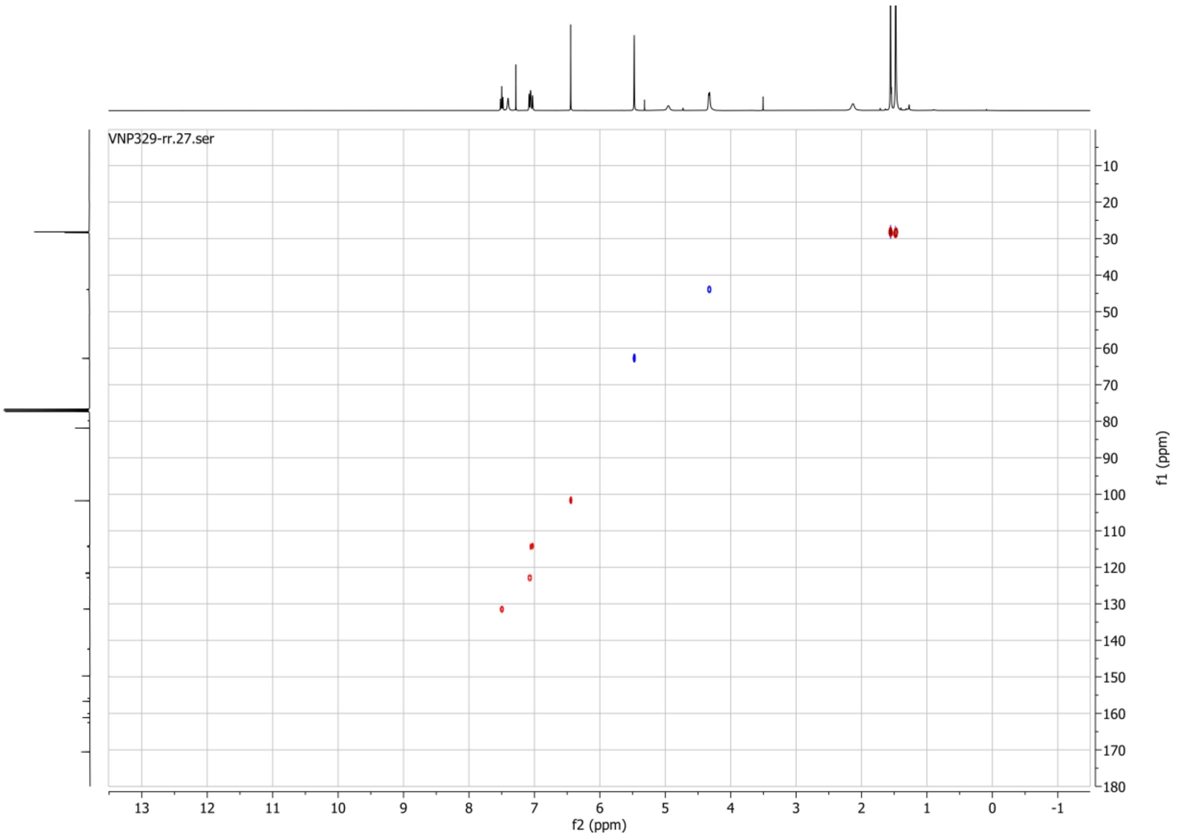


**19F spectrum of compound 125:**

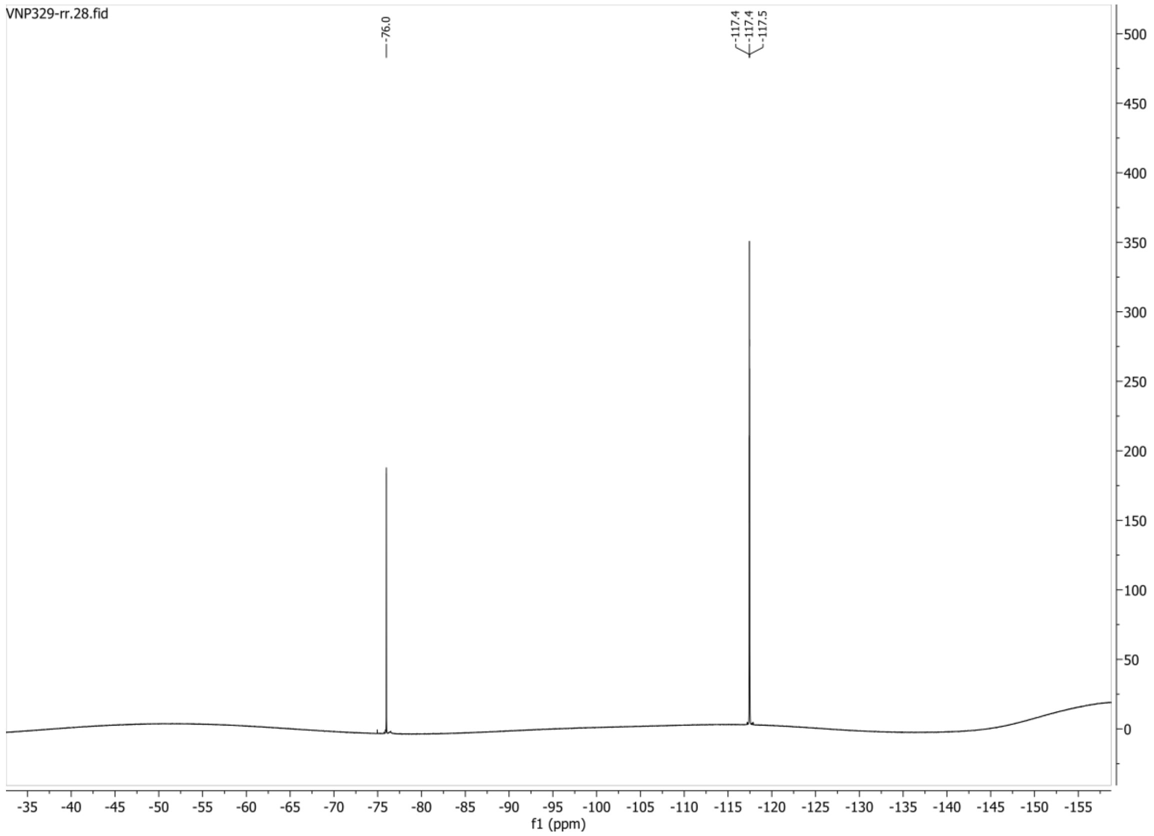

**1H spectrum of compound 126:**

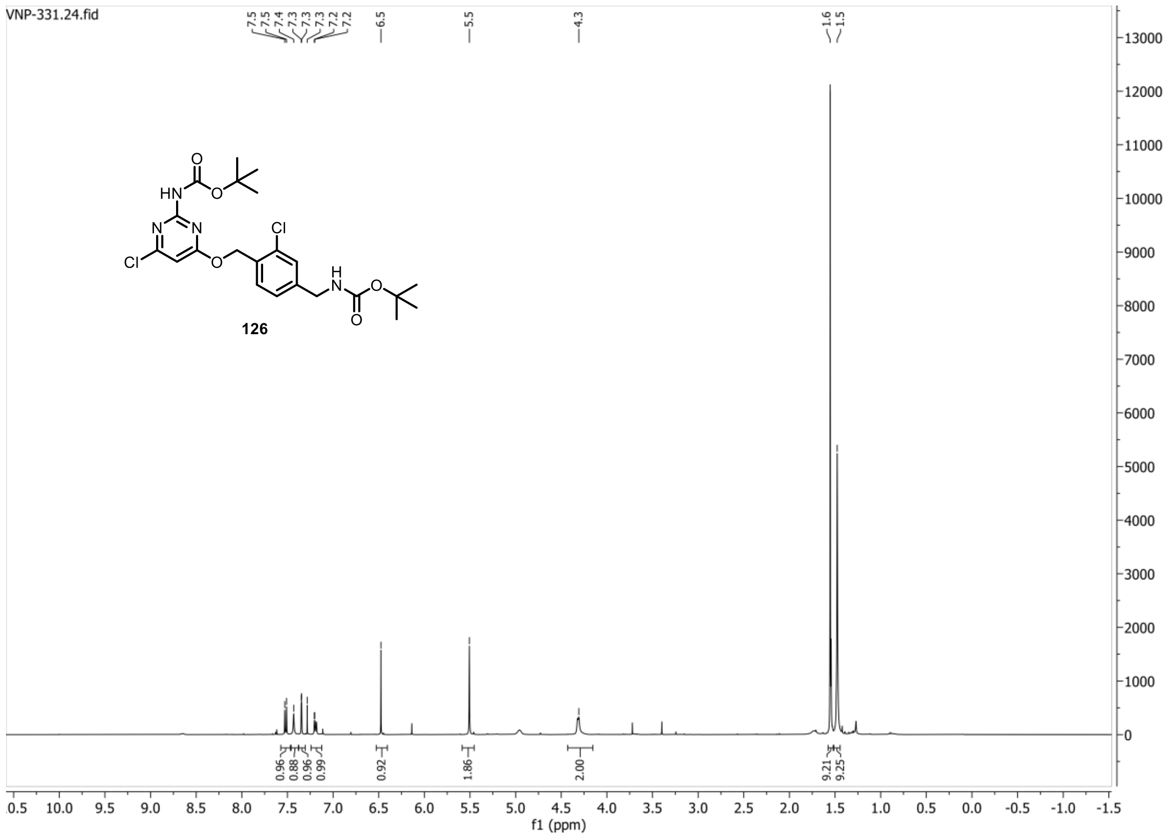


**13C{1H} spectrum of compound 126:**

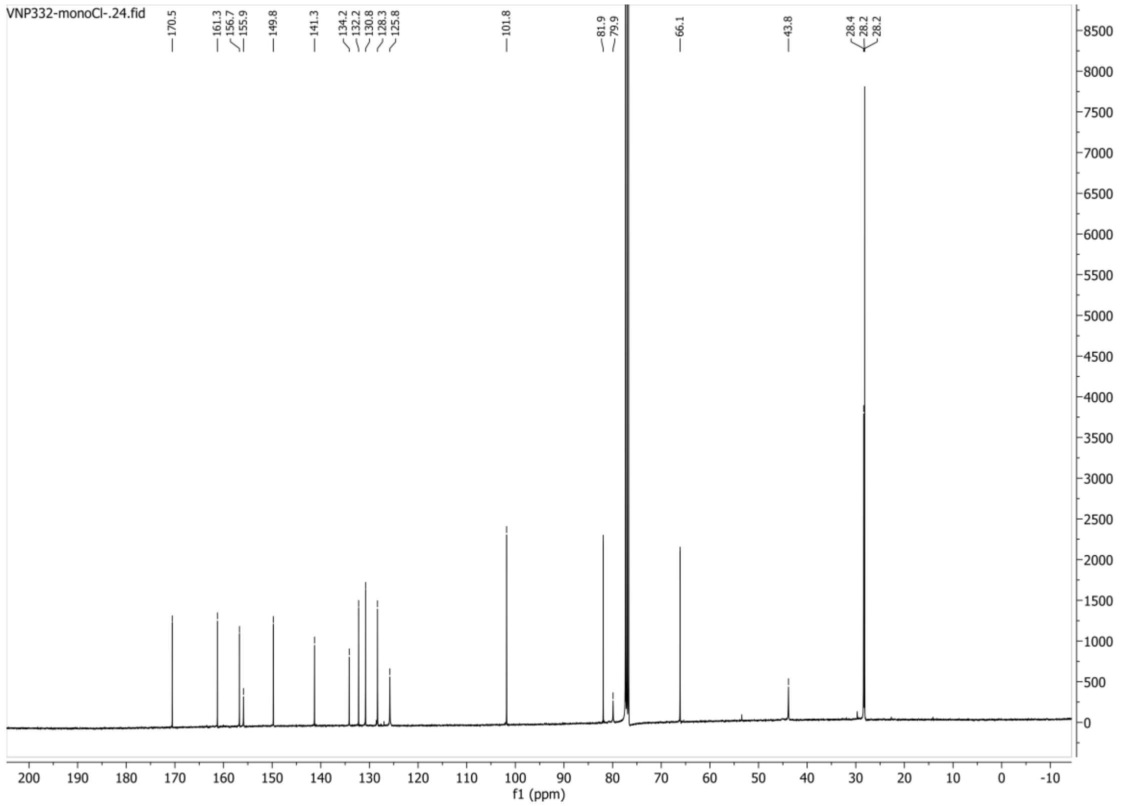

**HSQC-DEPT135 spectrum of compound 126:**

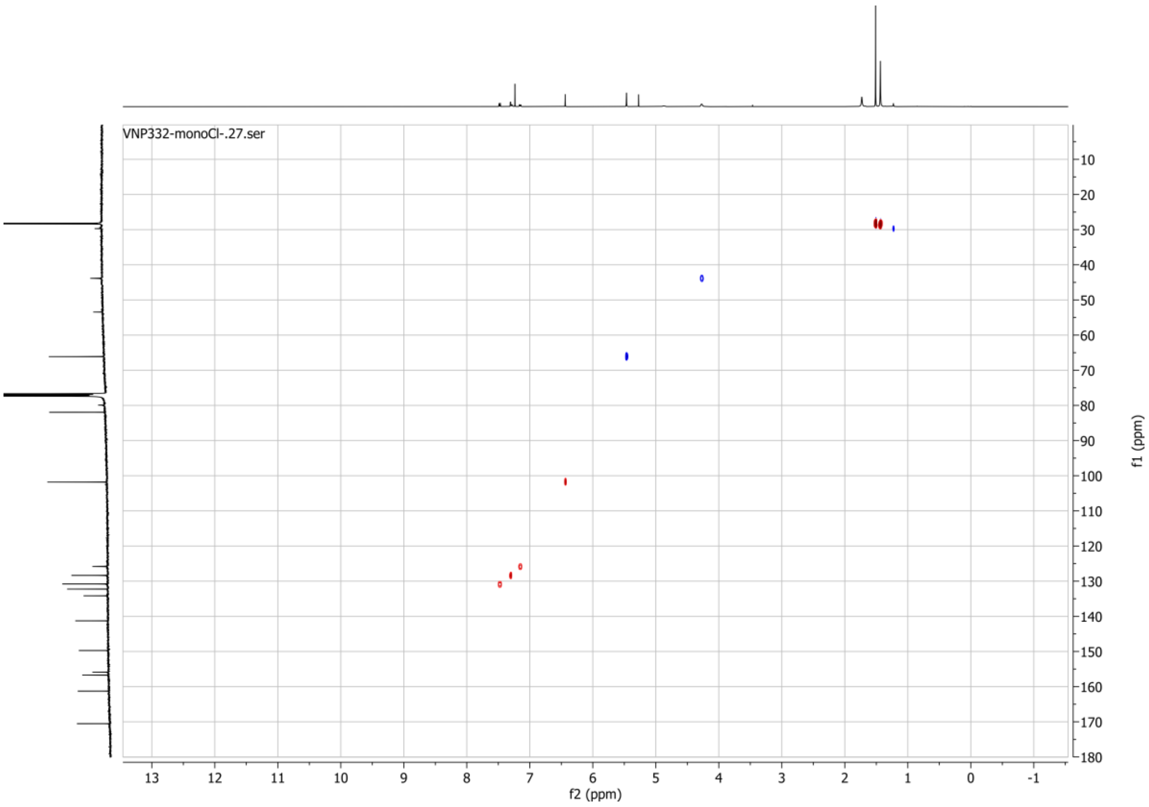

#### 1H spectrum of compound 130:

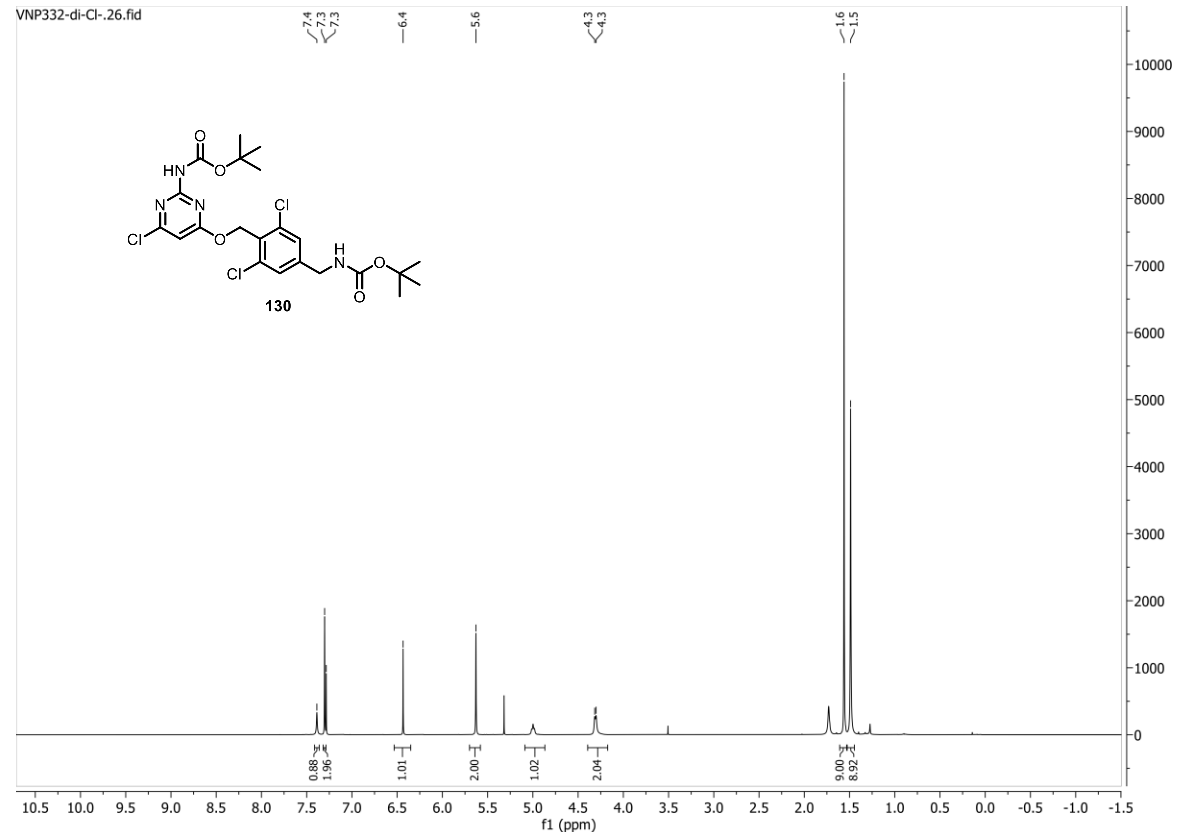


#### 13C{1H} spectrum of compound 130:

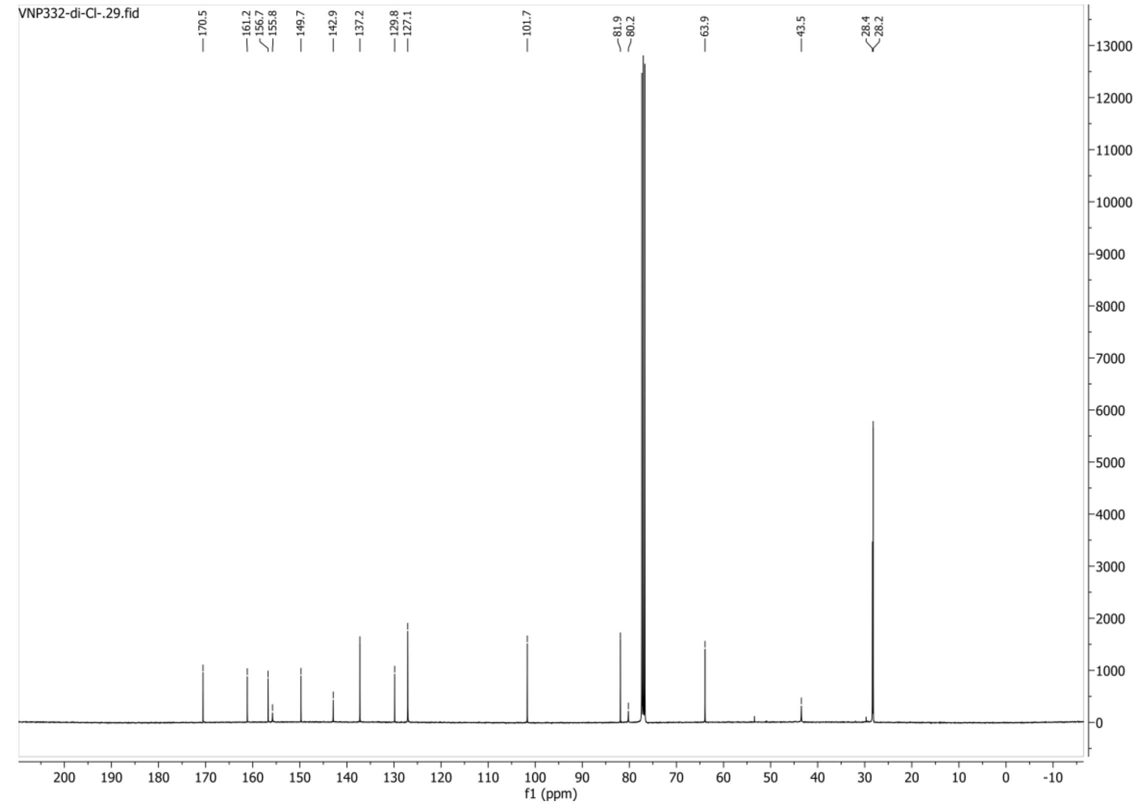

**HSQC-DEPT135 spectrum of compound 130:**

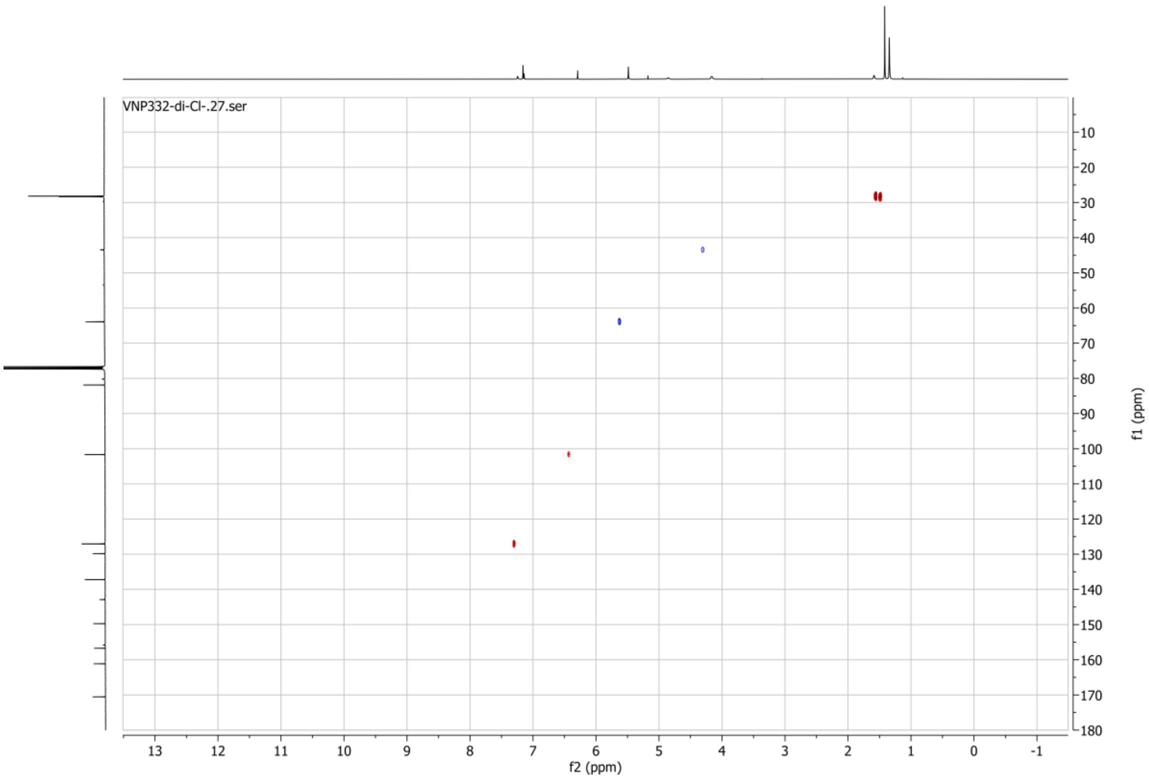

**1H spectrum of compound 134:**

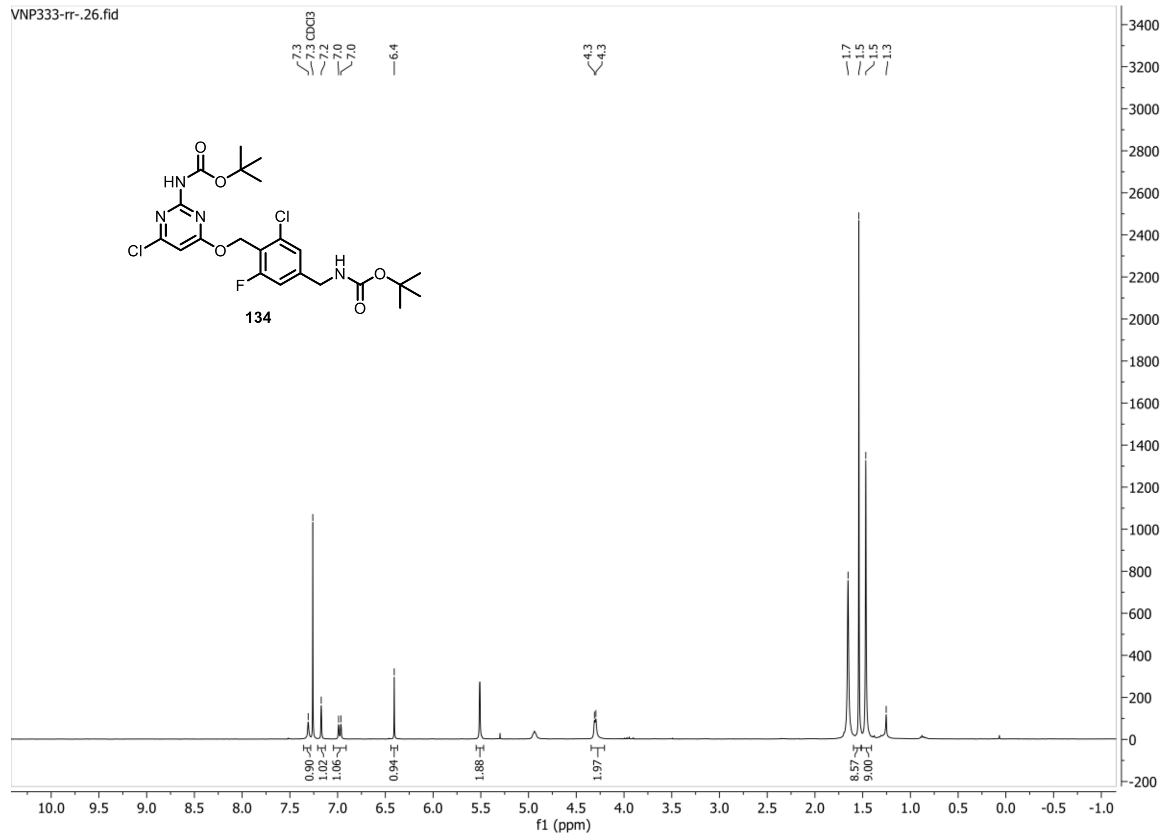


**13C{1H} spectrum of compound 134:**

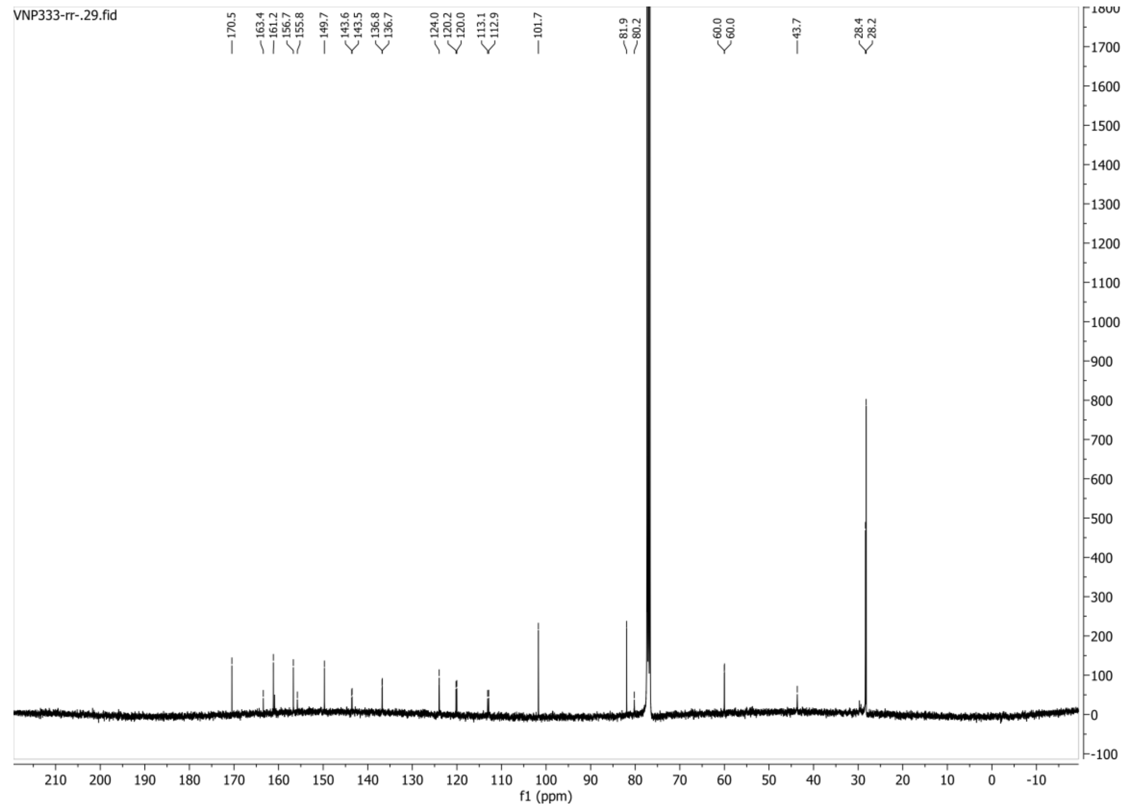

**HSQC-DEPT135 spectrum of compound 134:**

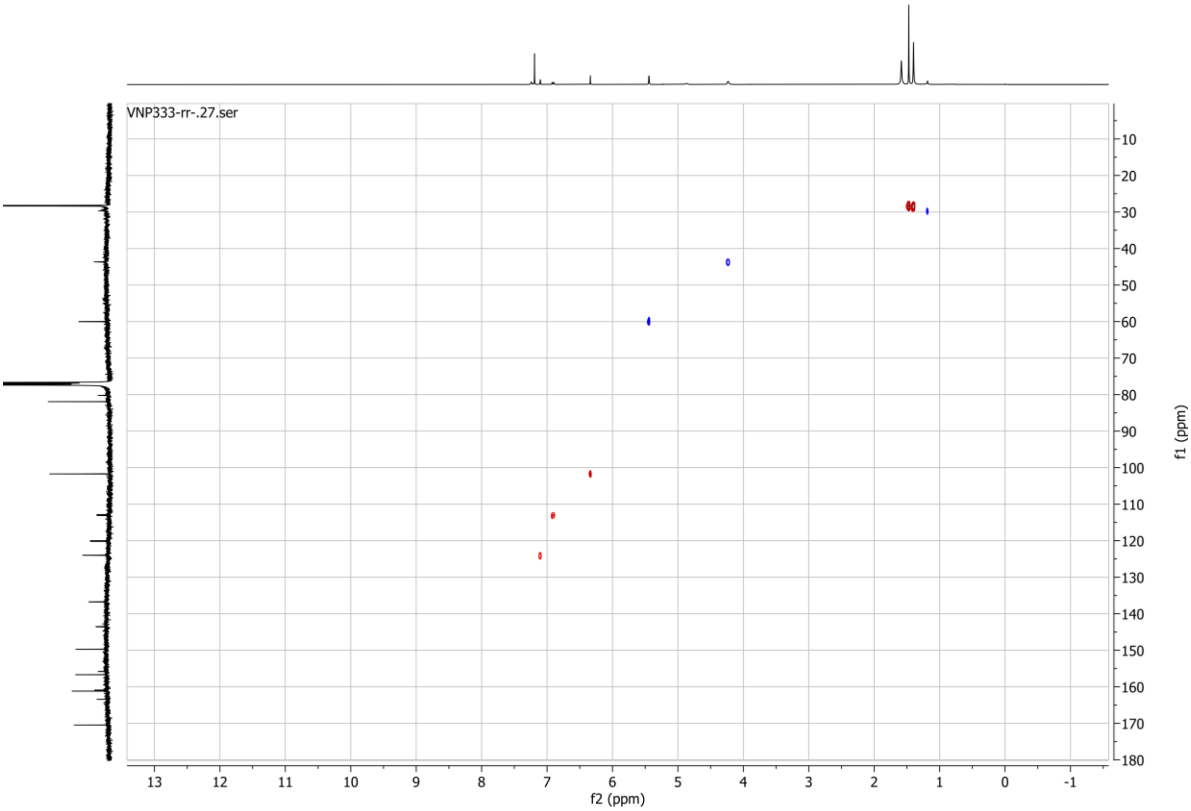


**19F spectrum of compound 134:**

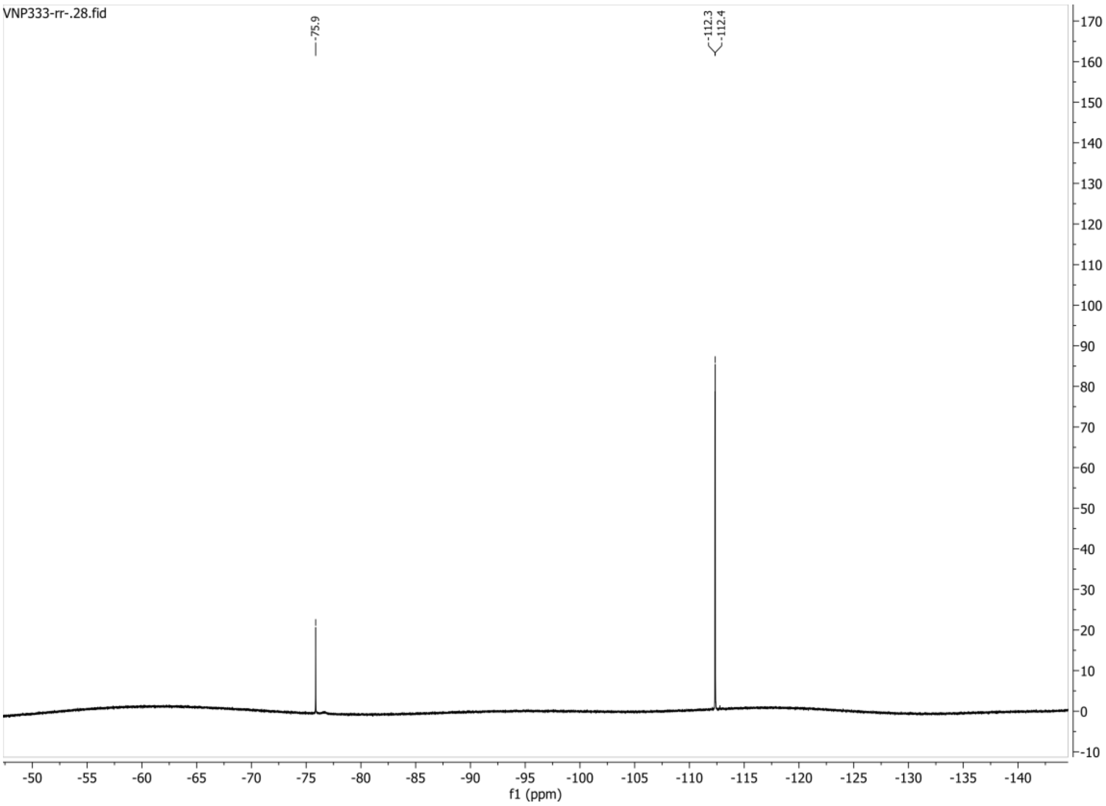

**1H spectrum of compound 135:**

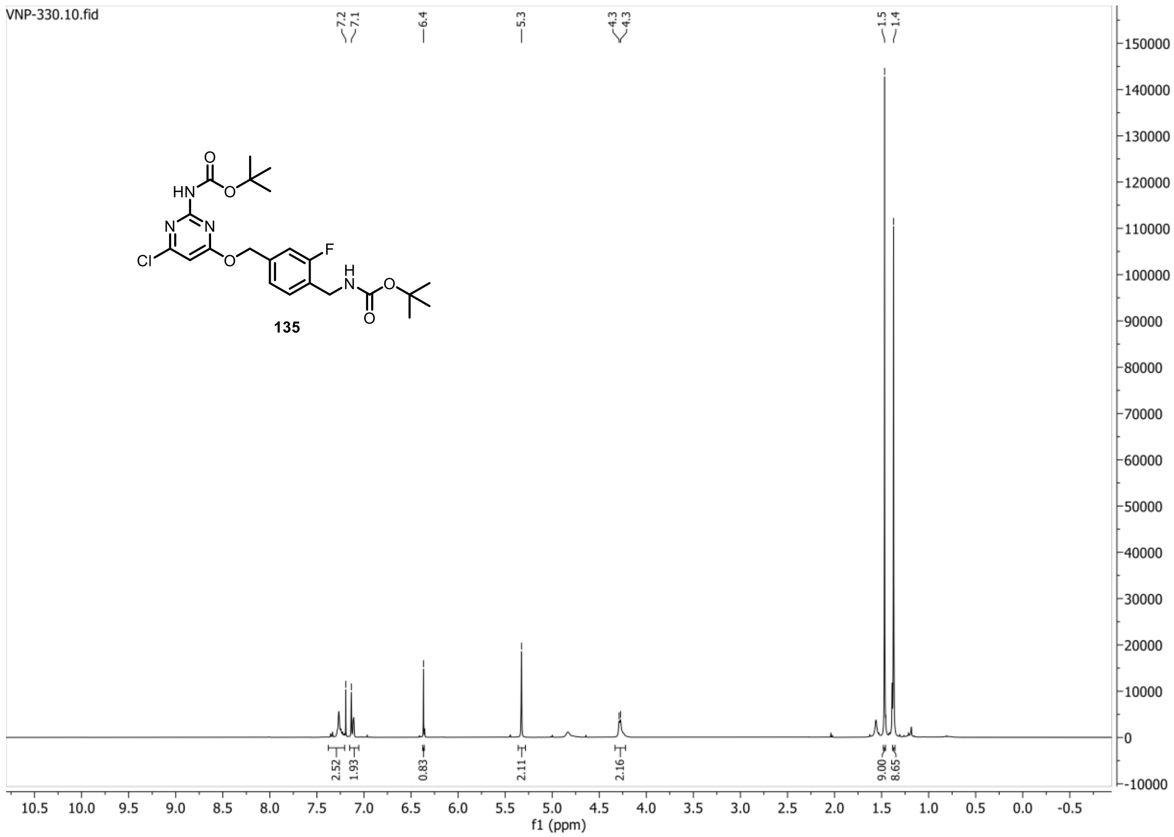


**13C{1H} spectrum of compound 135:**

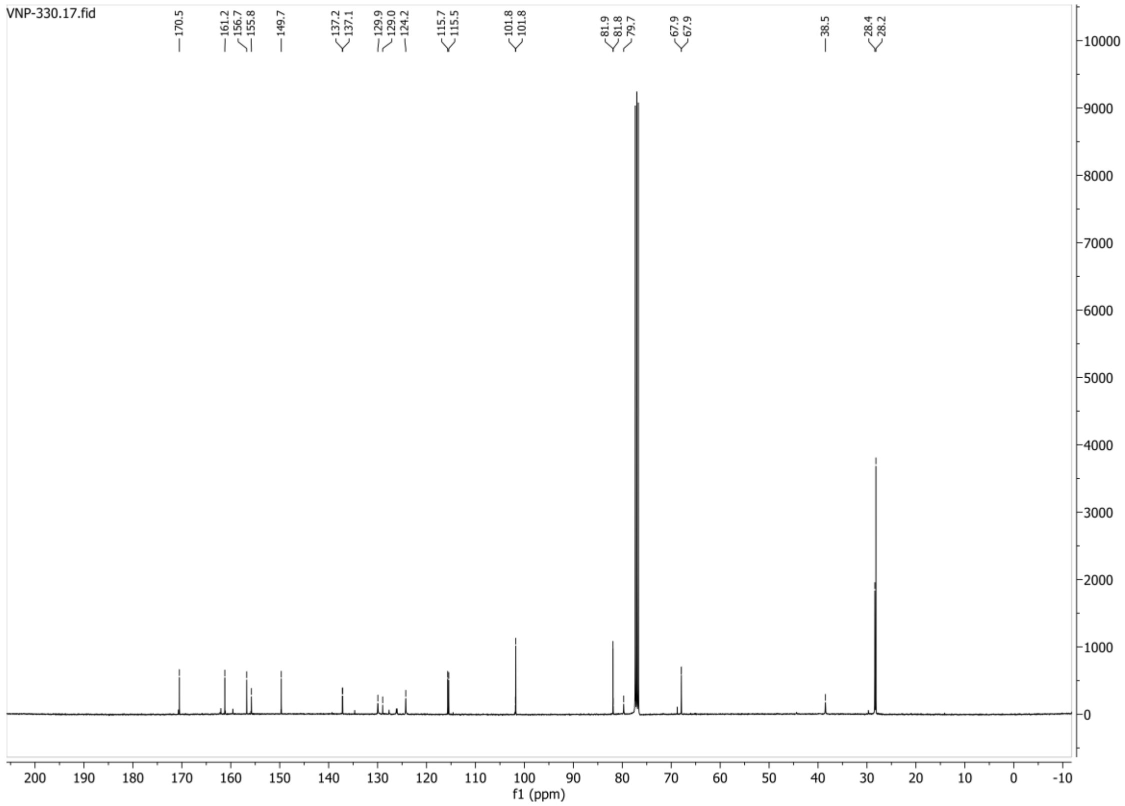

**HSQC-DEPT spectrum of compound 135:**

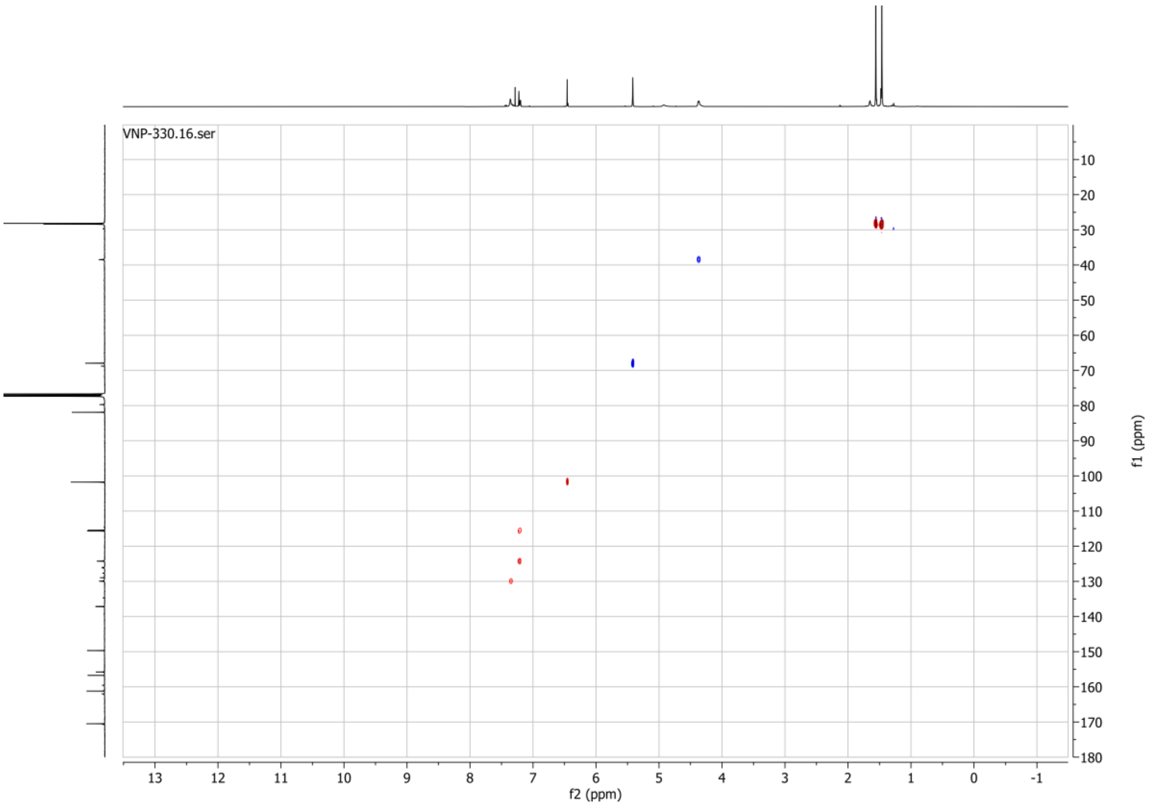


**19F spectrum of compound 135:**

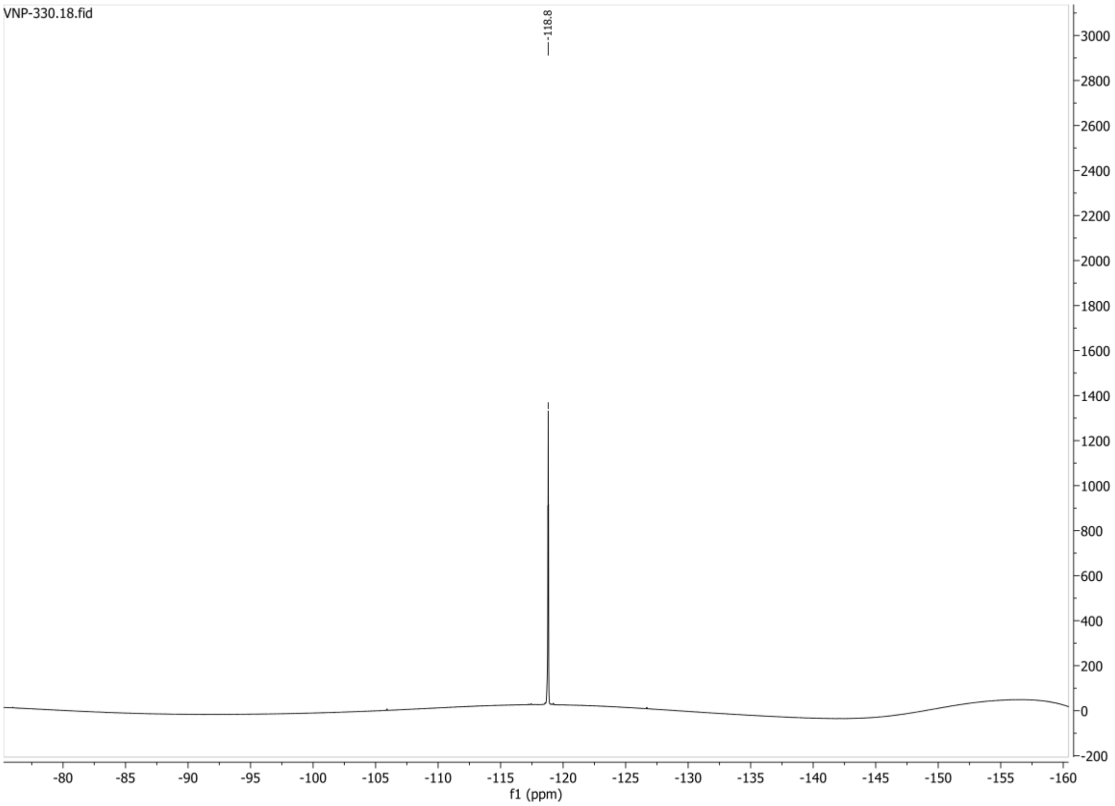

**1H spectrum of compound 140:**

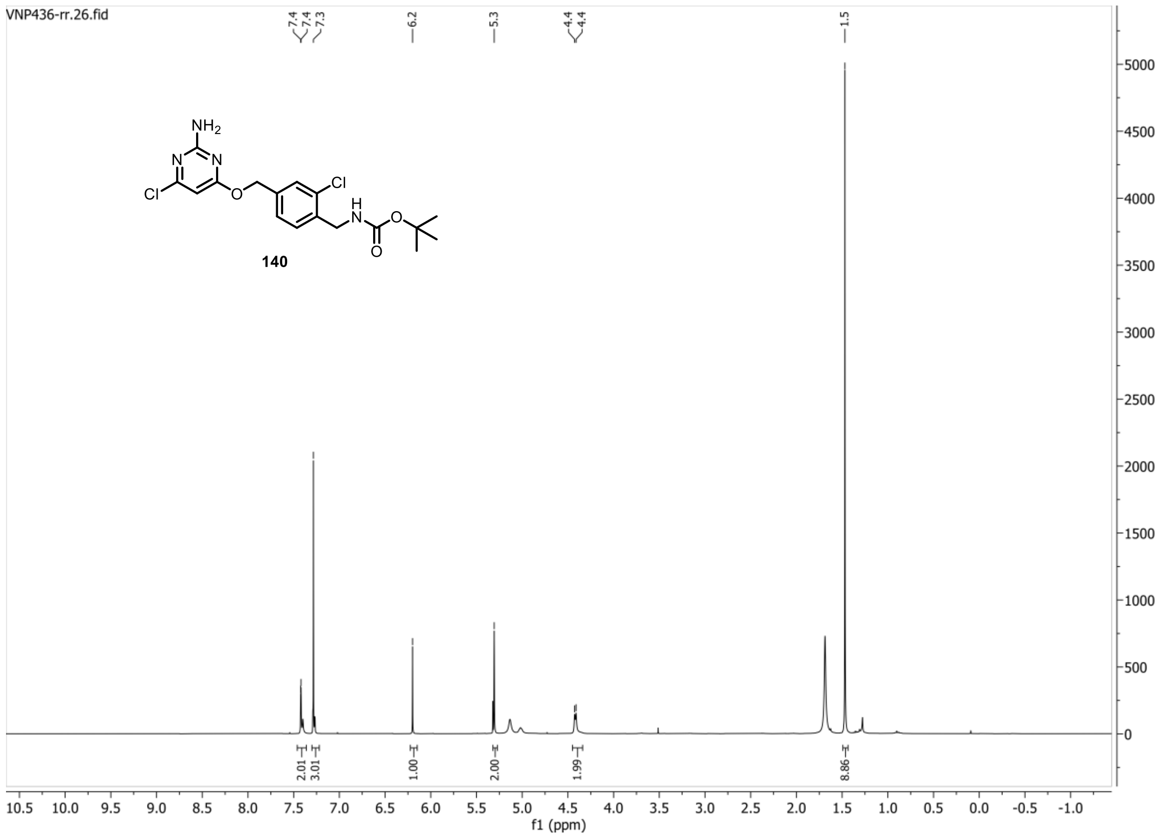


**13C{1H} spectrum of compound 140:**

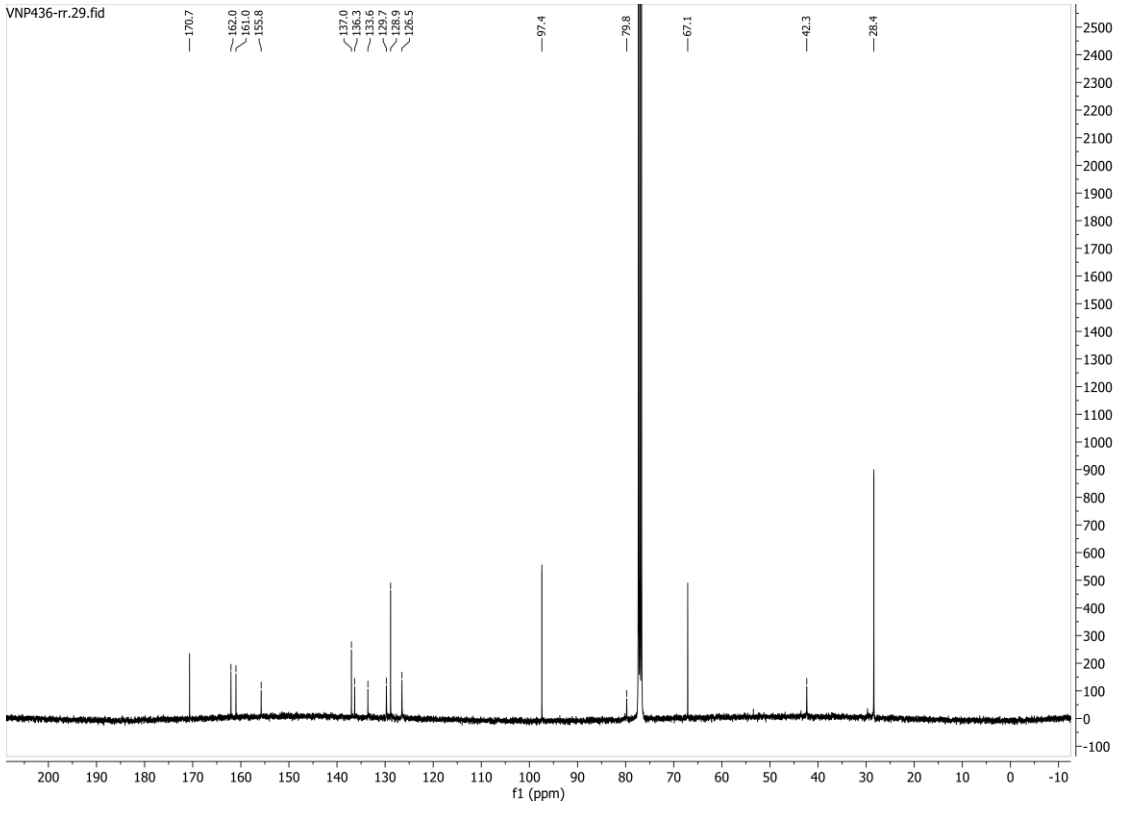

**HSQC-DEPT135 spectrum of compound 140:**

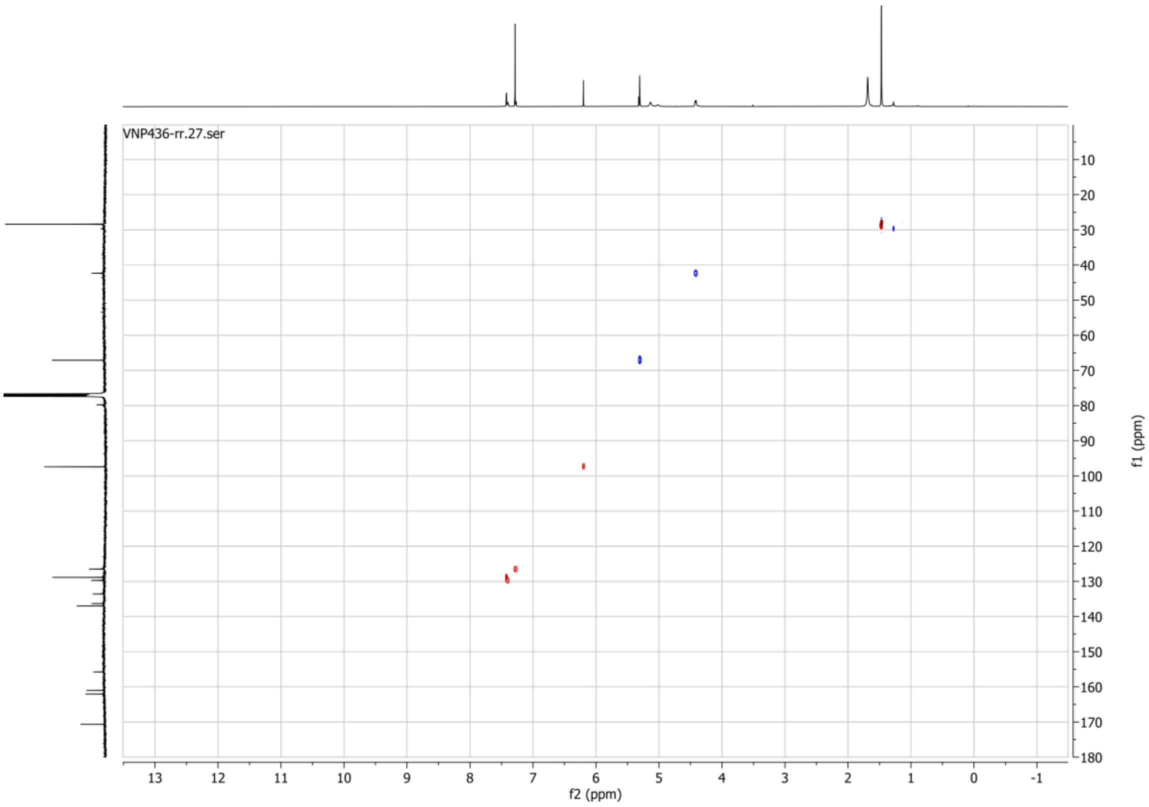

**1H spectrum of compound 144:**

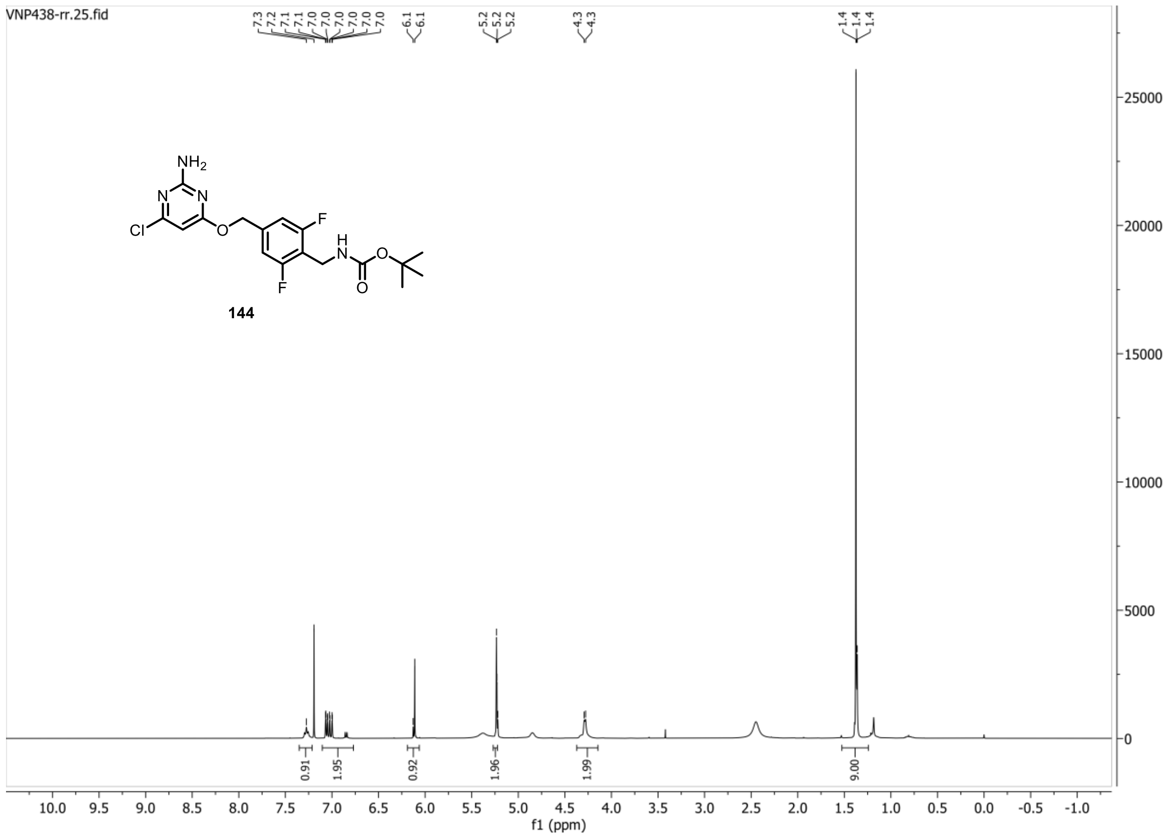


**13C{1H} spectrum of compound 144:**

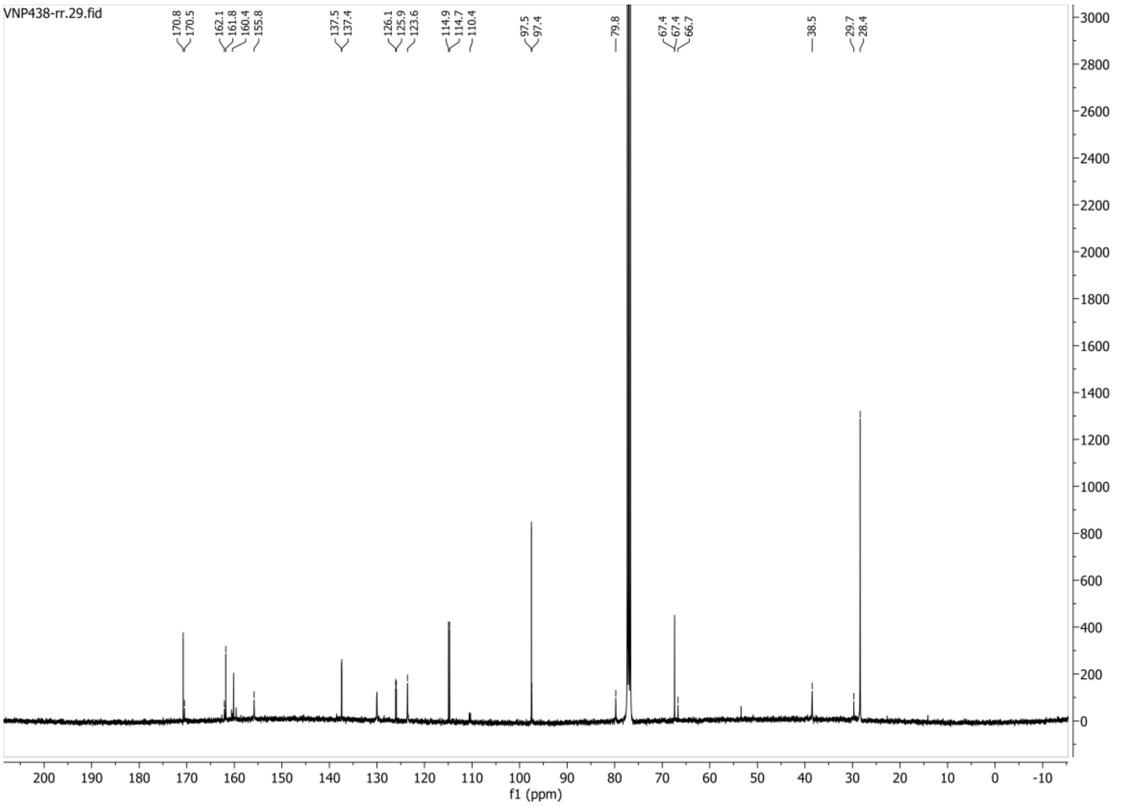

**HSQC-DEPT135 spectrum of compound 144:**

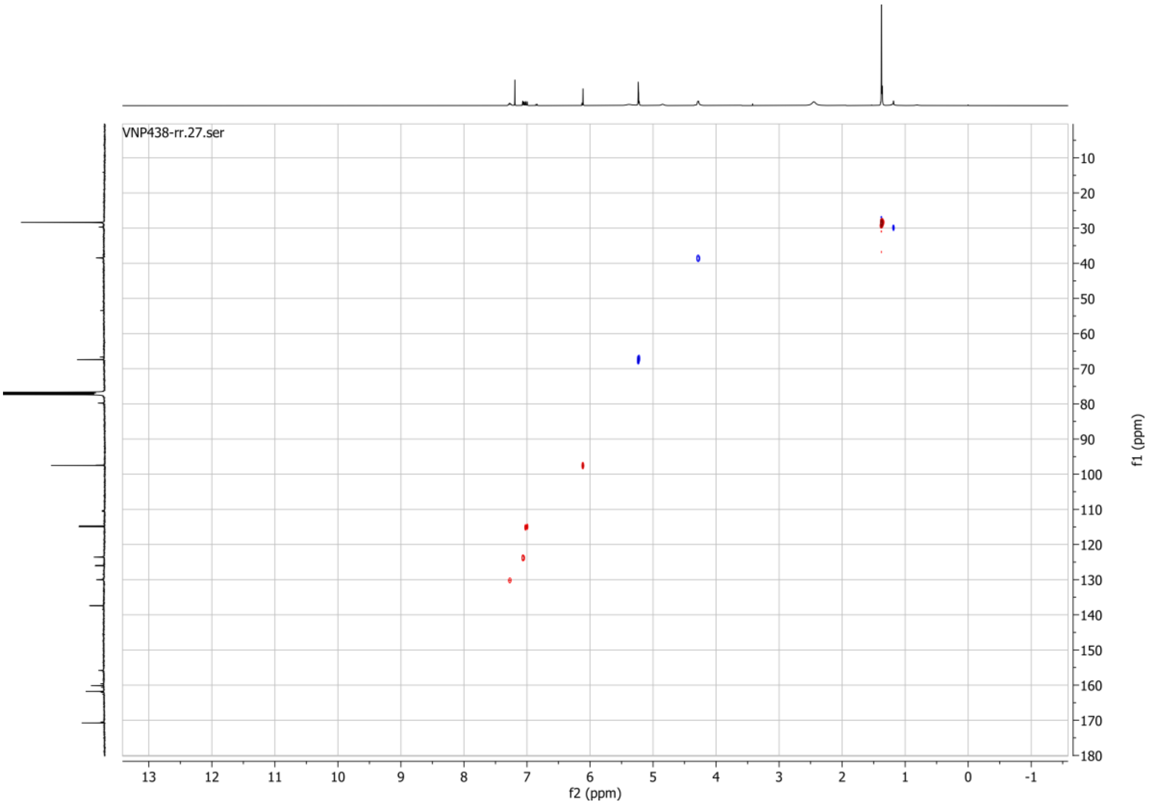


**19F spectrum of compound 144:**

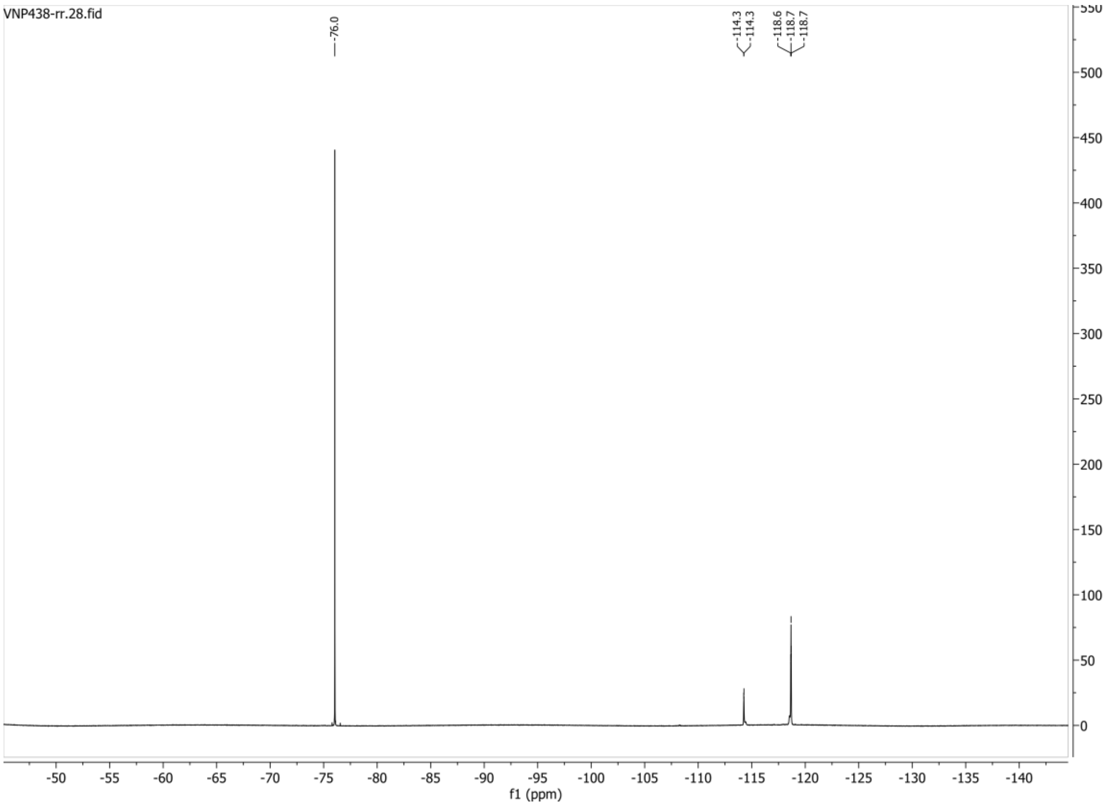

**1H spectrum of compound 145:**

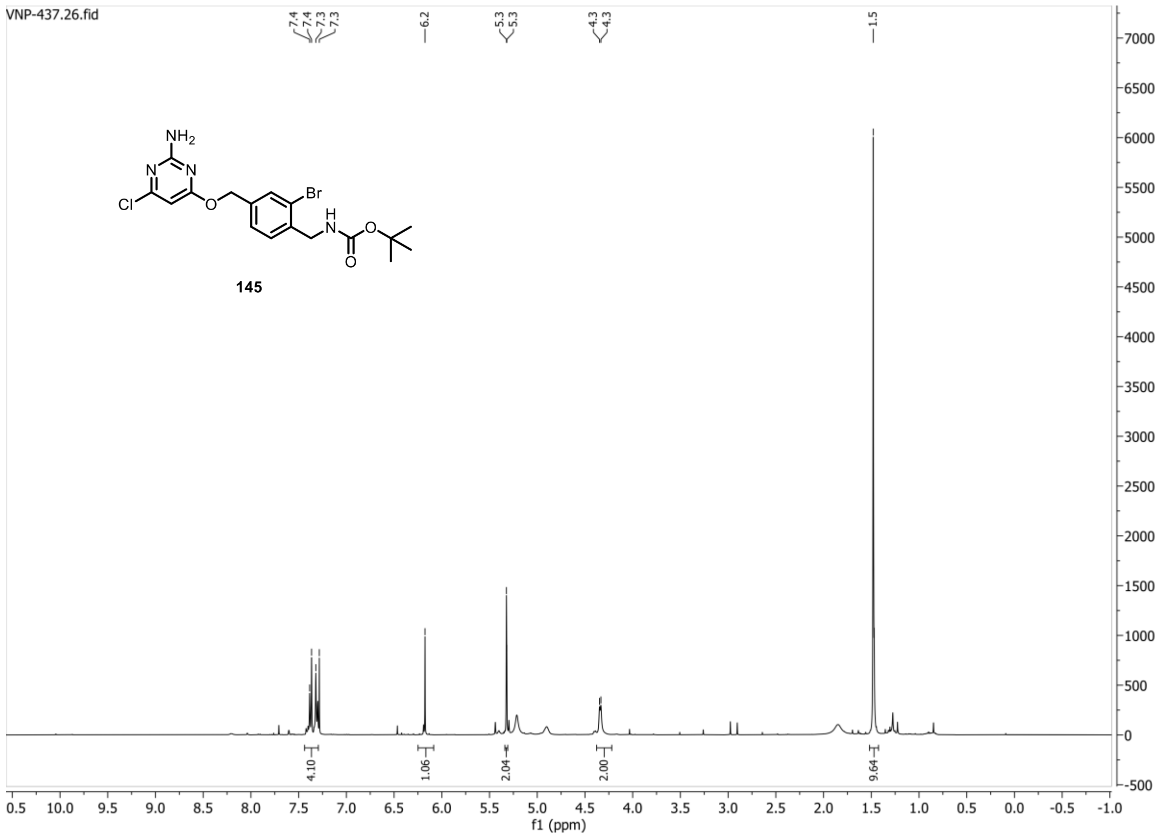


**13C{1H} spectrum of compound 145:**

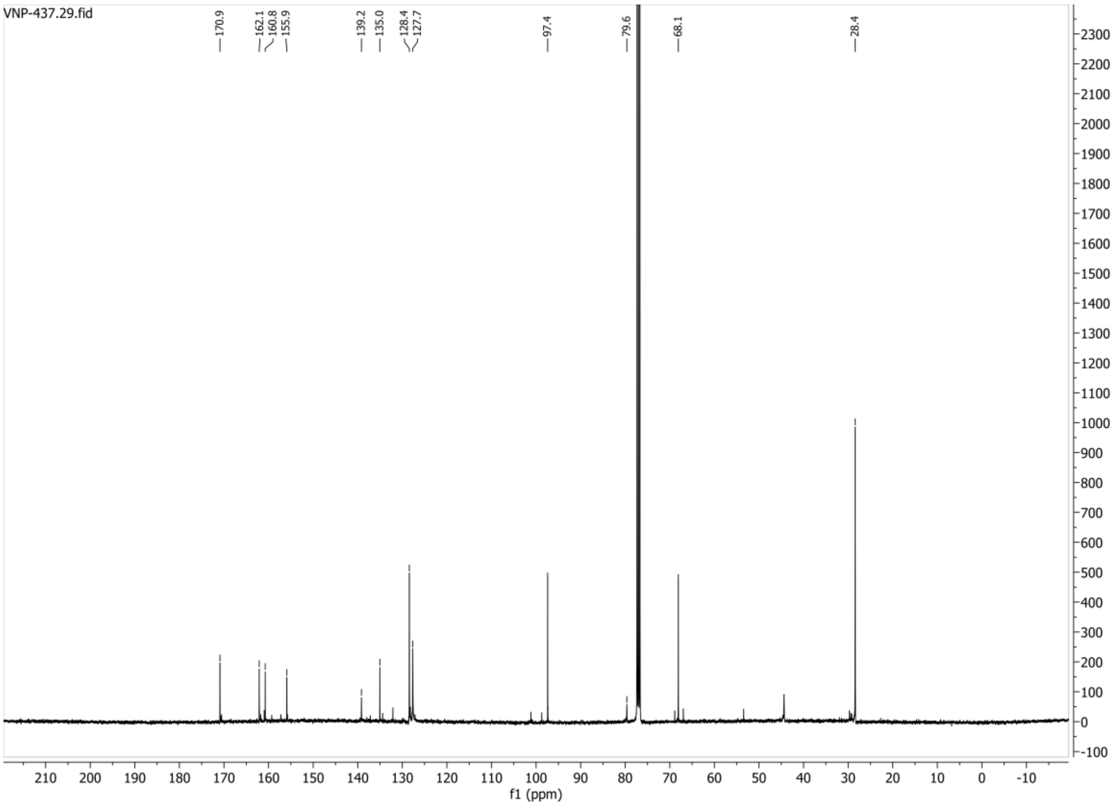

**HSQC-DEPT spectrum of compound 145:**

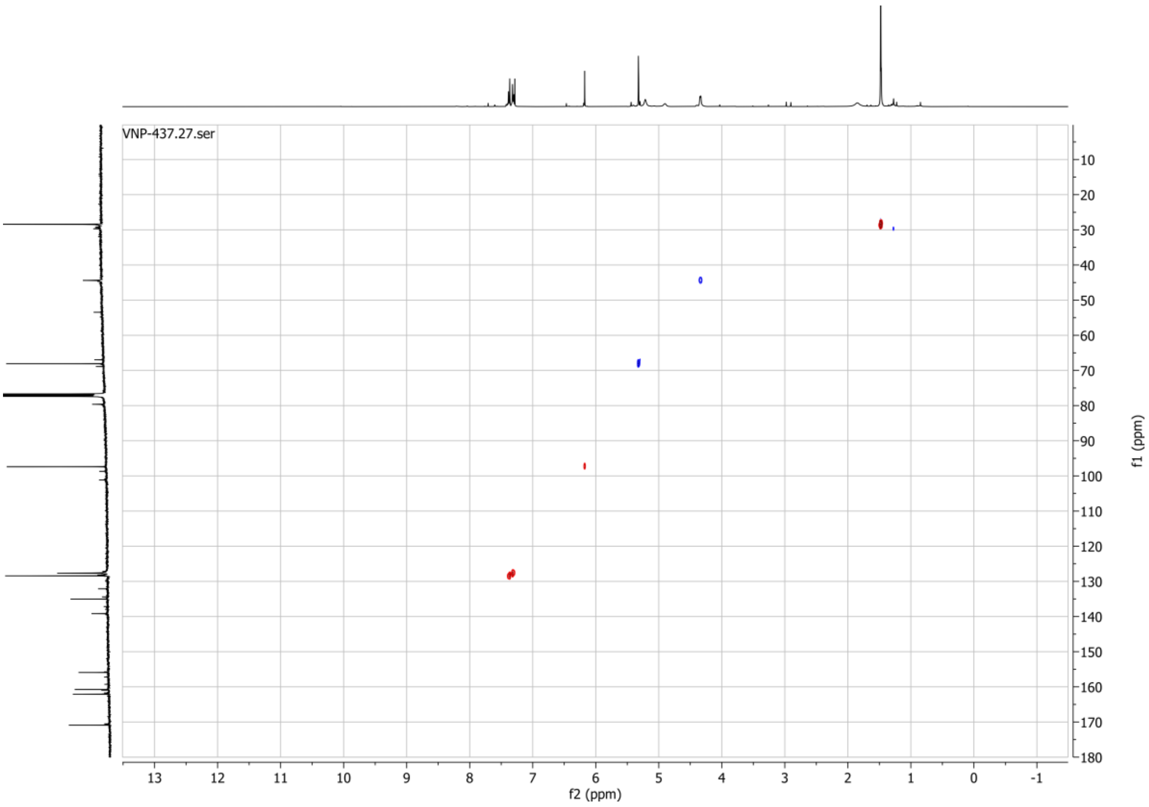

**1H spectrum of compound 146:**

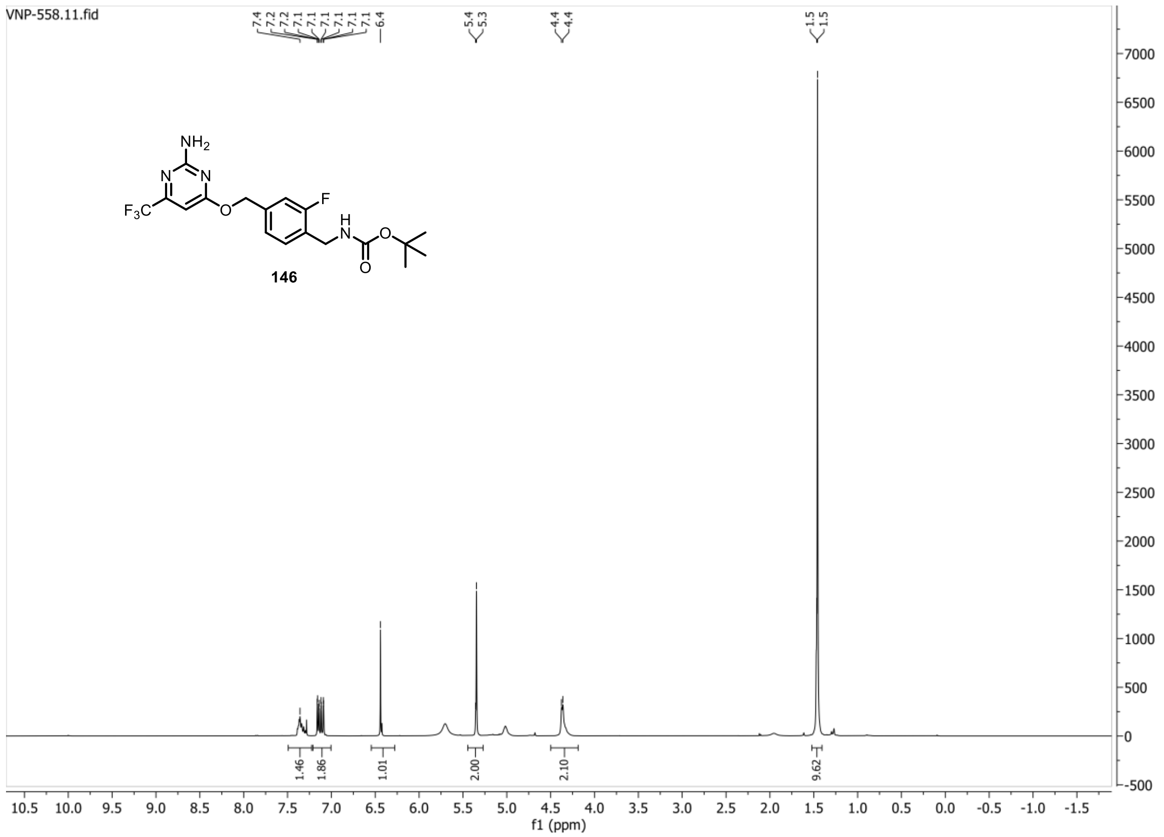


**13C{1H} spectrum of compound 146:**

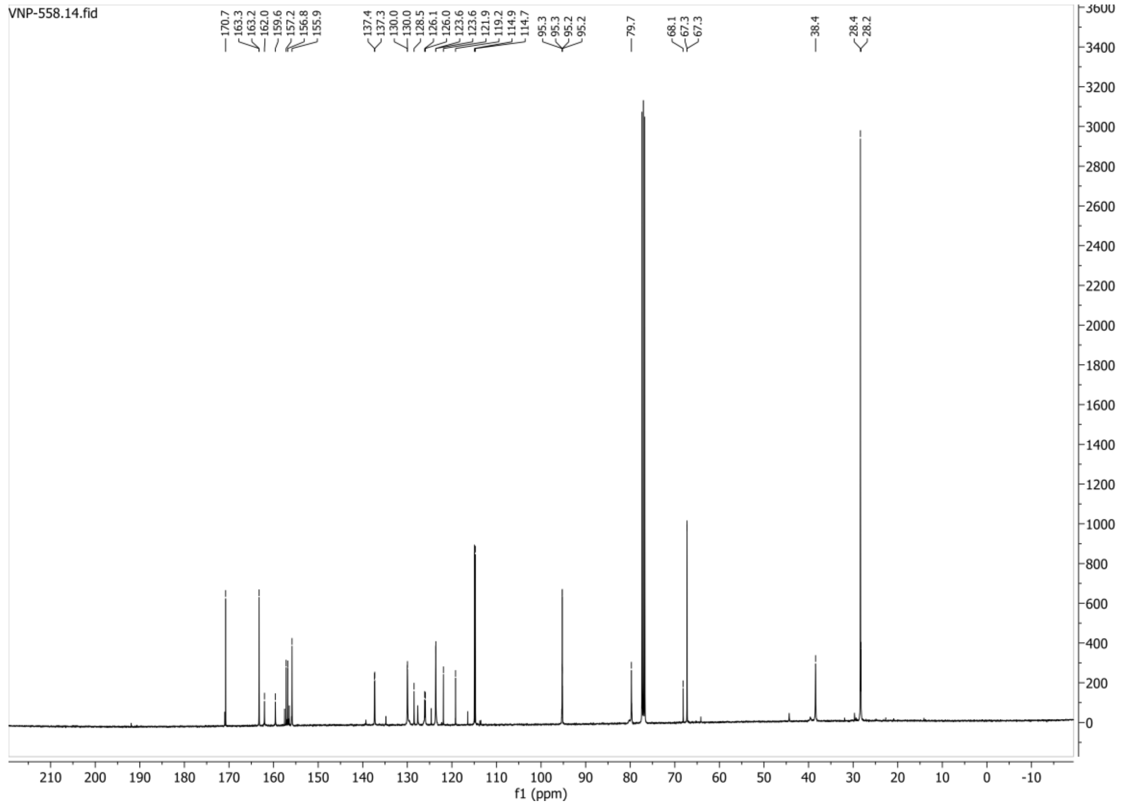

**HSQC-DEPT spectrum of compound 146:**

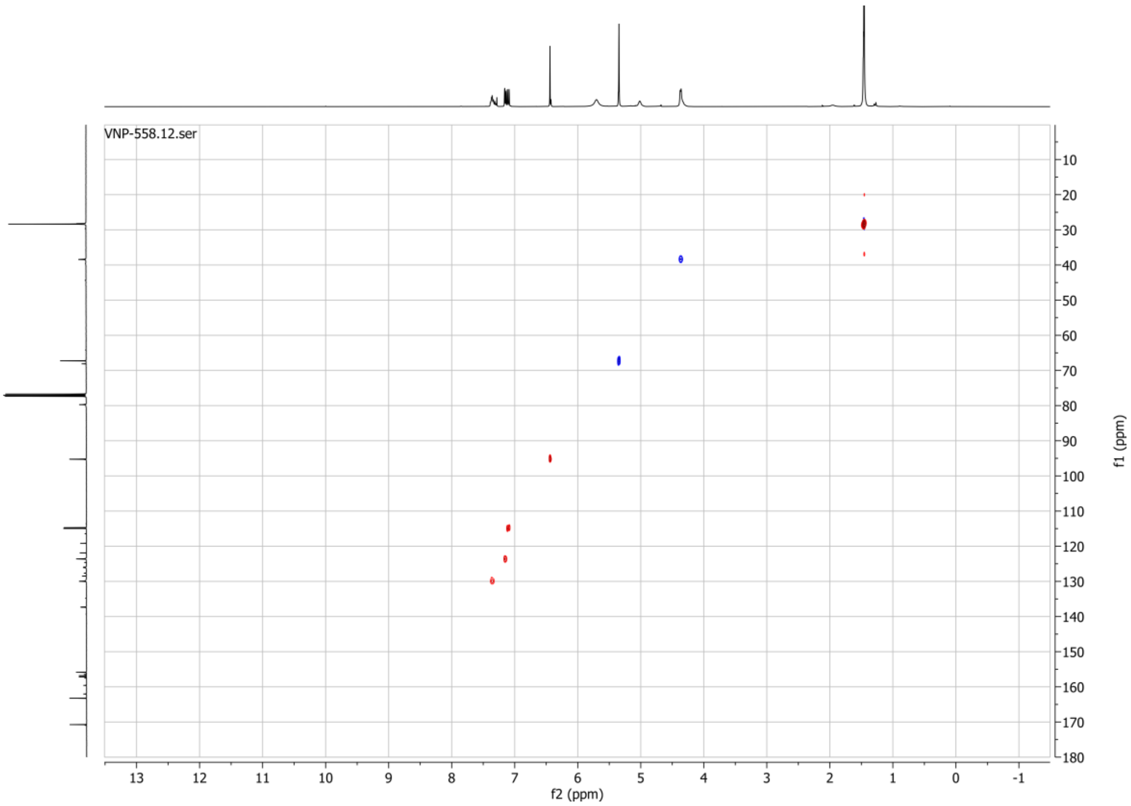


**19F spectrum of compound 146:**

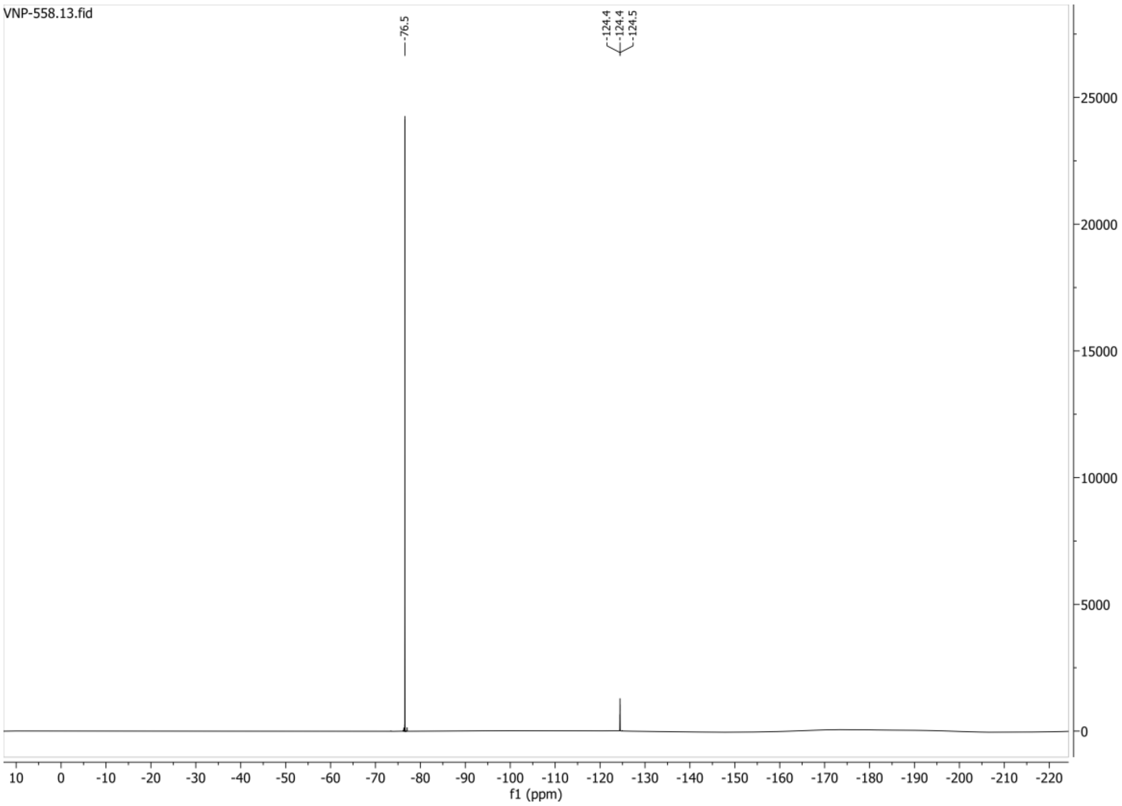

**1H spectrum of compound 151:**

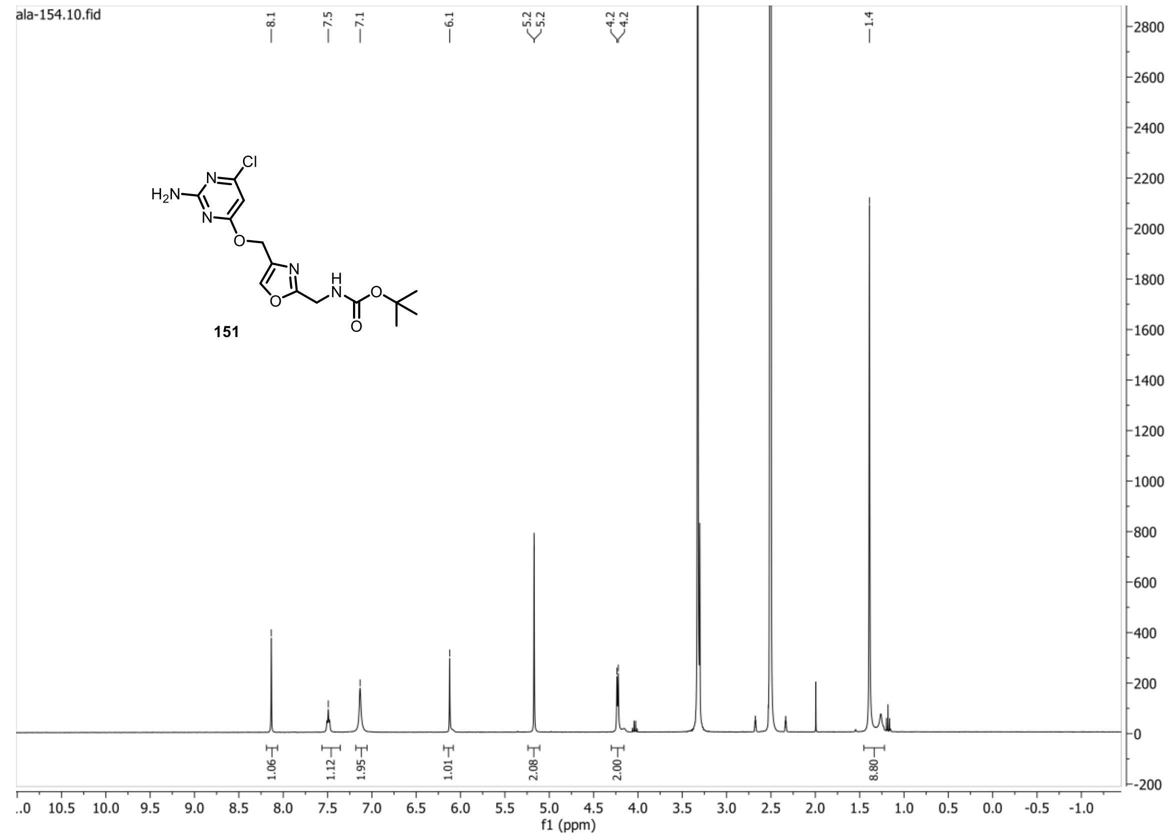


**13C{1H} spectrum of compound 151:**

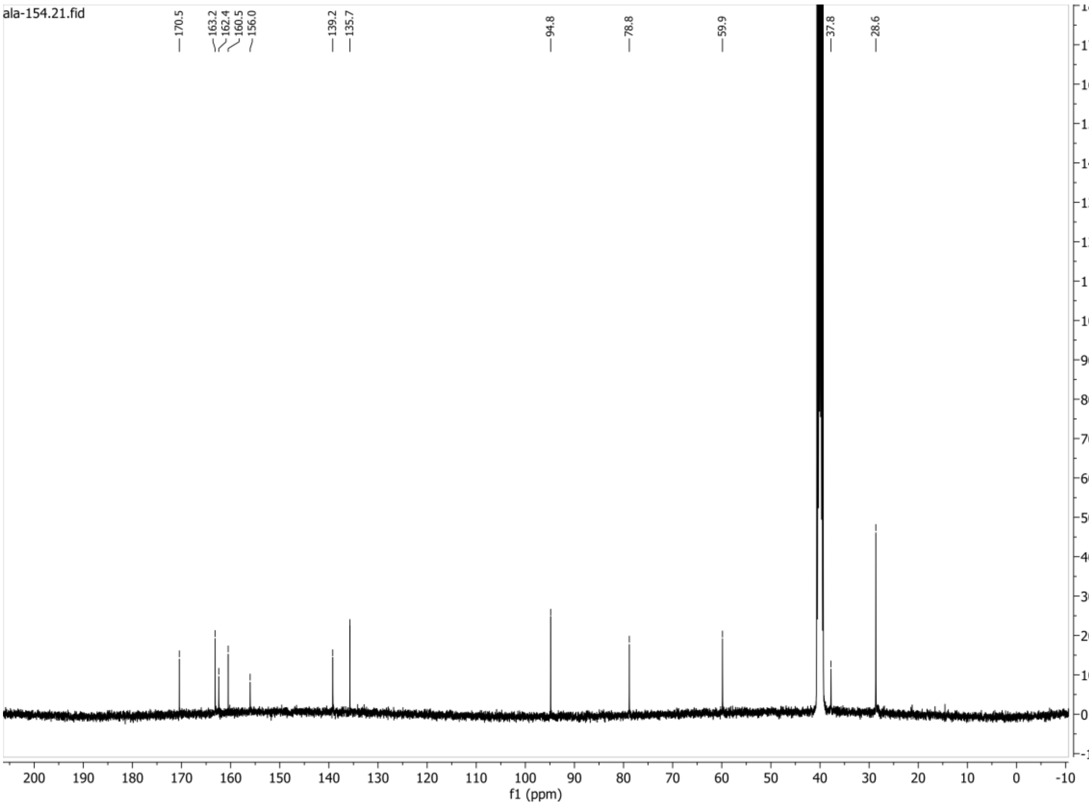

**1H spectrum of compound 157:**

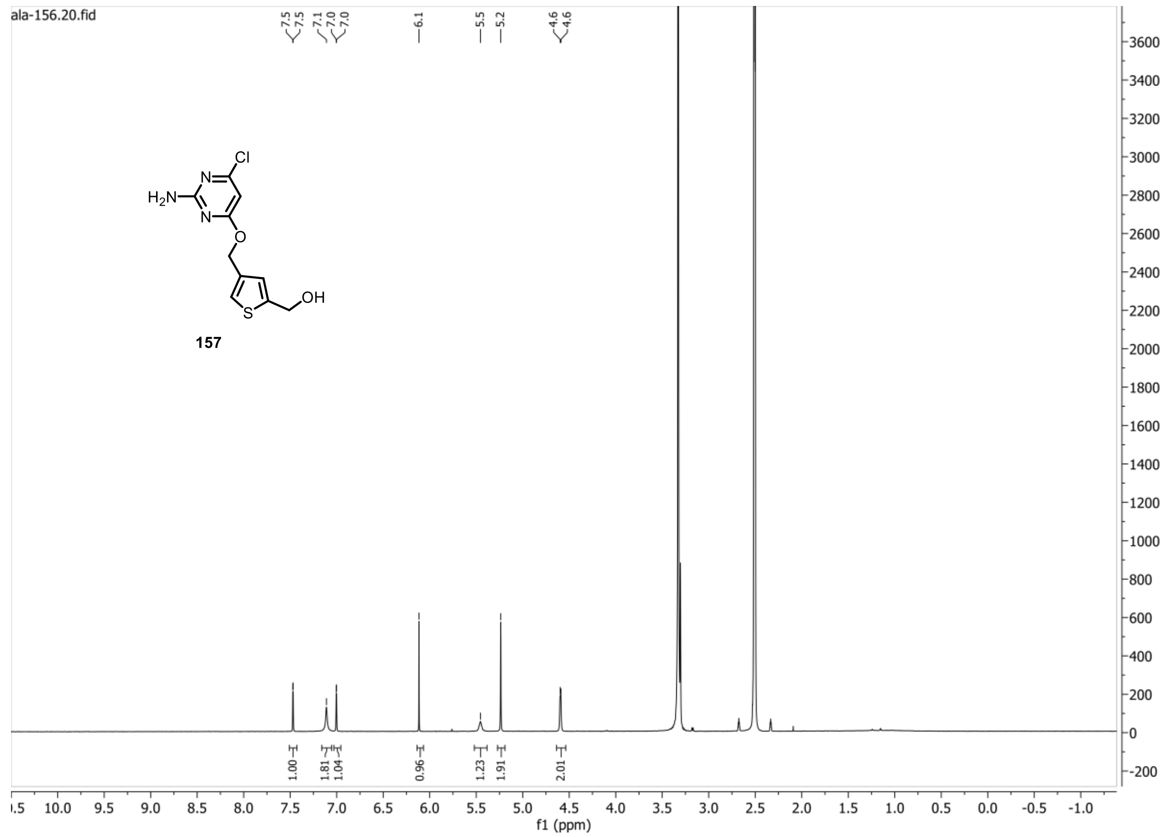


**13C{1H} spectrum of compound 157:**

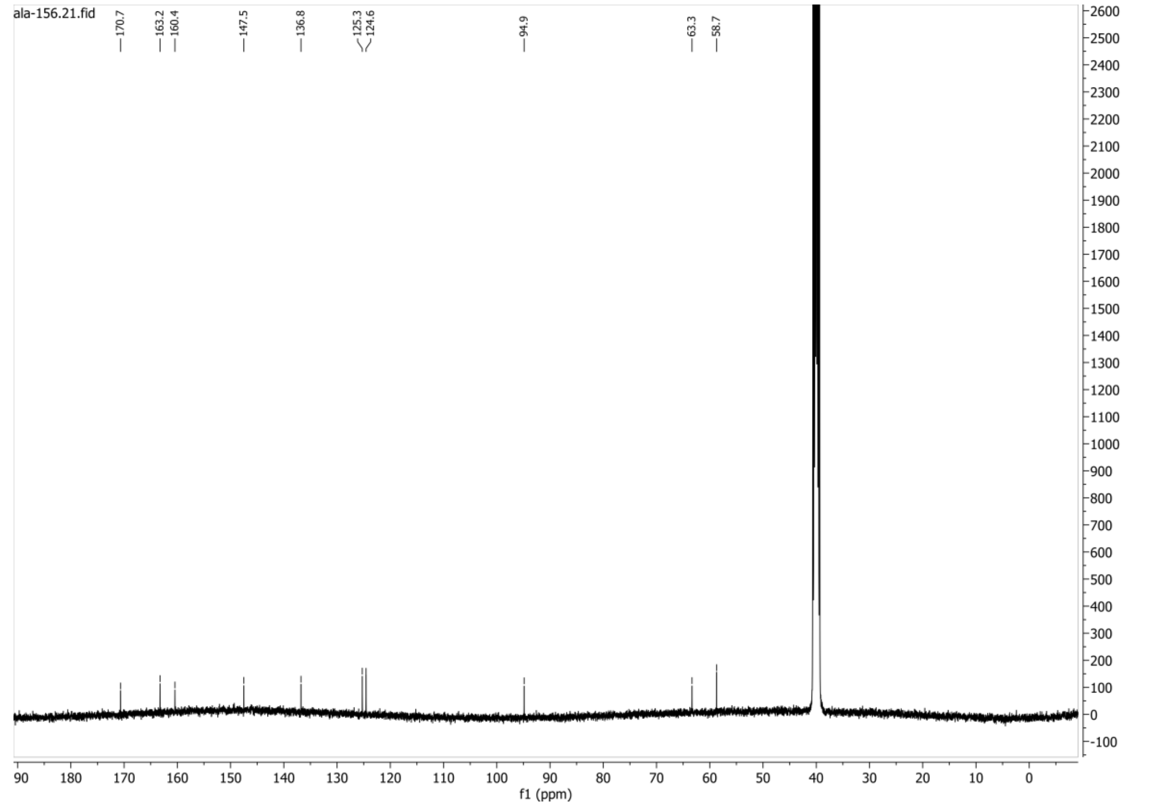

**1H spectrum of compound 164:**

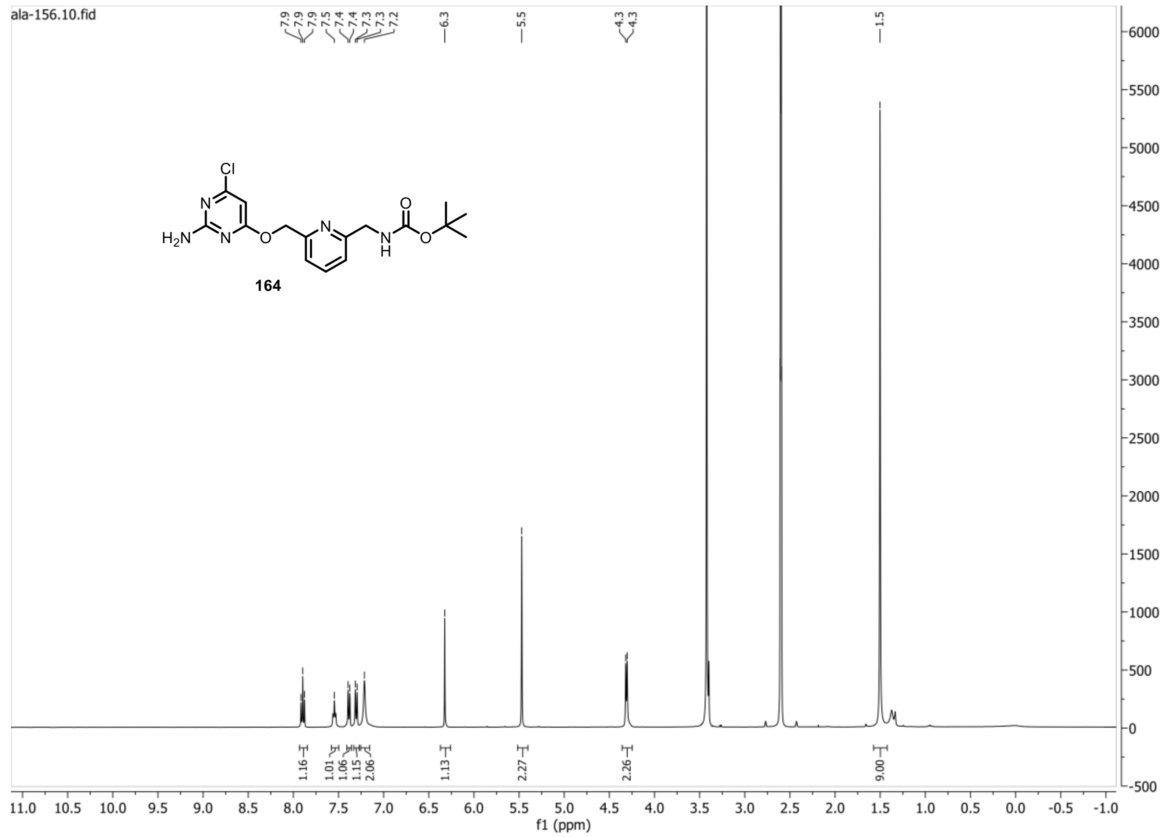


**13C{1H} spectrum of compound 164:**

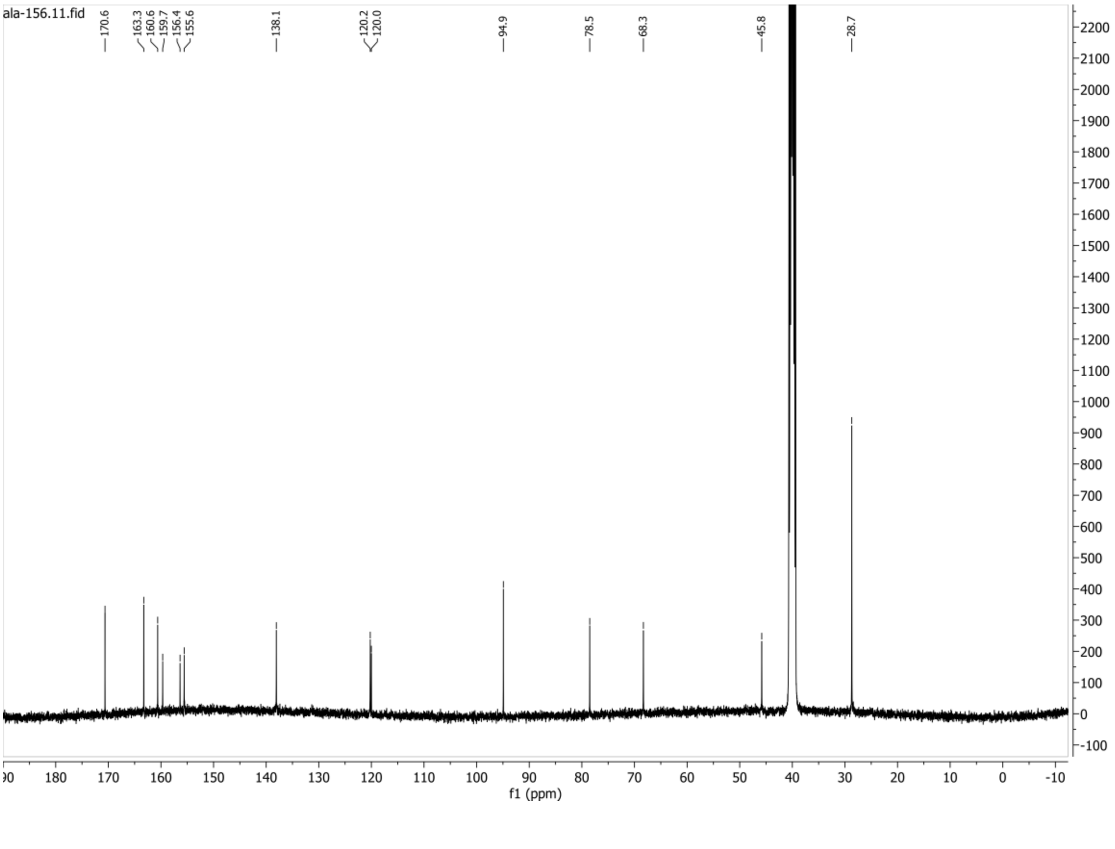

**1H spectrum of compound 169:**

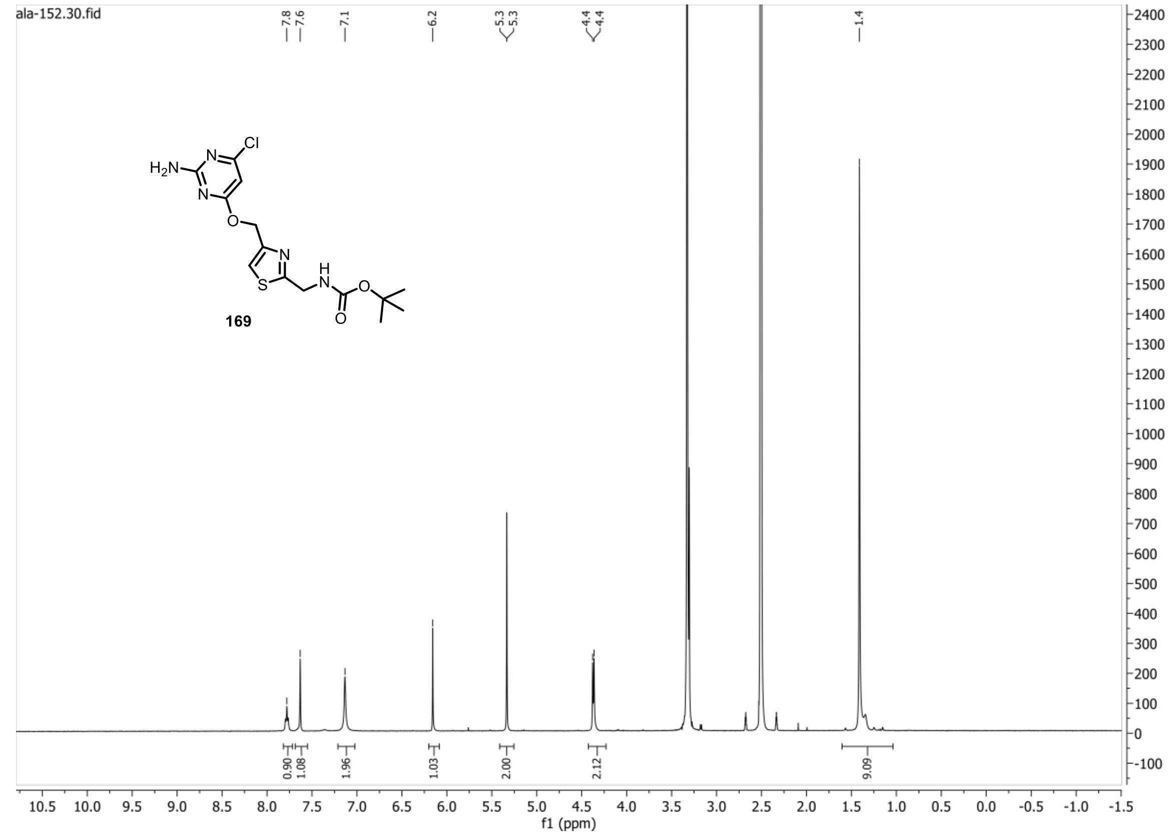


**13C{1H} spectrum of 169:**

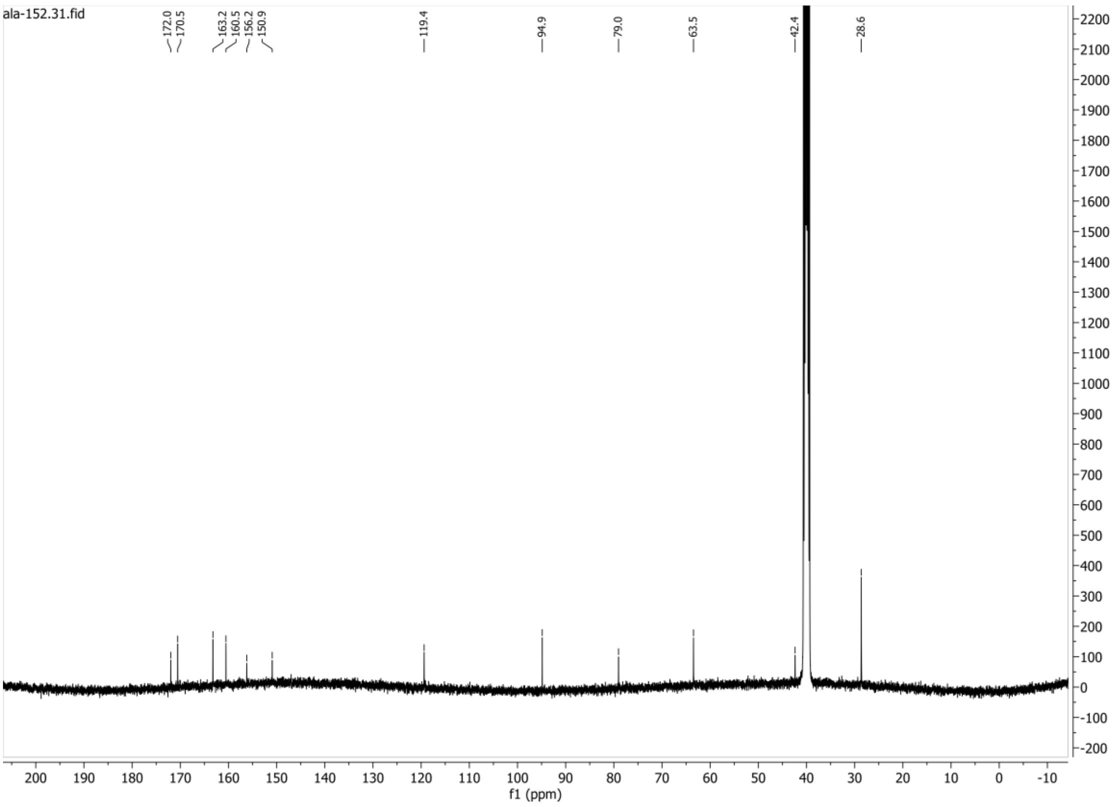
